# An experimental approach towards untangling the role of nature versus nurture in shaping the microbiome of social insects

**DOI:** 10.1101/2023.09.04.556269

**Authors:** Tali Magory Cohen, Levona Bodner, Sondra Turjeman, Efrat Sharon, Alisa Cohen, Sofia Bouchebti, Evgeny Tikhonov, Omry Koren, Eran Levin

## Abstract

The gut microbiota is intimately related to host wellbeing, in terms of physiology, immune function, and even social interactions. The strength of this relationship is dynamic, but the extent to which the microbiome is shaped by the identity of the host (nature) or its environment (nurture) remains largely unknown. Here we aimed to identify factors shaping the microbiomes of nursing workers and larvae of two Vespidae species, using a cross-species experimental design to control for effects of environment, host identity and their interactions. We found that the microbiome composition of adults depended principally on the environment. Conversely, larval microbiome composition differed more between host species, regardless of treatment. We also found distinct microbiota profiles between the two species, across life stages and independently. These findings further elucidate the complexity of the host-microbiome relationship shaped by the environment while retaining symbionts that benefit the host. These results suggest that holobiont evolution may have promoted the rise of social behavior in animals.

## Introduction

The microbiome is a dynamic ecosystem existing within its host, shaping many of its crucial functions and traits ^1^. High microbiome diversity and favorable composition have been suggested to expand the physiological potential of the host ^2^ and even affect fitness ^3^. The importance of the host-microbiome relationship extends interspecifically, as the composition of the gut microbiota can be used to differentiate between host species, including species with similar diets and some shared gut symbionts ^3^. A recent paradigm shift suggests that the association between host and microbiota is the result of a co-evolutionary process by which both evolve as a single entity (i.e., the holobiont approach ^4^). However, the strength of this relationship can change throughout an individual’s lifespan ^5^ or as a function of spatial dynamics ^4^. Therefore, the extent to which the microbiome composition and diversity are shaped by the identity of the host (i.e., nature) or its extrinsic environment (i.e., nurture) remains largely unknown.

Host genotype or evolutionary history is often associated with the core microbiome (i.e., the microbial community that is shared across multiple individuals from the same host ^6–8)^. While some evidence suggests that certain host loci are linked to variations in the composition of the bacterial community in humans ^5, 9–13^, evidence from genome-wide association studies (GWAS) suggests that only a small portion of the reported heritability of the microbiome can be attributed to host genetics in a classical familial inheritance pattern ^14^, emphasizing the need to understand the role of host identity in colonization and maintenance of gut microbiota.

Alternatively, the hypothesis that extrinsic factors (i.e., environmental) account for a larger proportion of the variance in the microbiome than genetics, is more widely supported ^15–17^. Berasategui et al. (2016 ^18^) showed that the microbiomes of several distinct species feeding on the same food source (conifer seedlings) were highly similar, suggesting that the environment played a bigger role in determining the gut microbial community. Similar results were found in beetles ^19^ and among army ants and their associated beetle species ^20^, for which foraging habits and habitat structured the microbial communities and even brood parasites ^21^. However, a large portion of the early evidence documenting environmental effects on the gut microbiome has been correlative. Recently, experimental approaches, with well-controlled environments, have been suggested to address causation in microbiome research ^22, 23^ as such methods have the power to contribute to our understanding of the basic evolutionary mechanism underlying host-microbiome development and change.

Social insects provide a unique opportunity to understand the relative roles of nature and nurture in determining the microbiome of individuals, where both factors can be controlled. In many species, individuals belonging to the same colony are very closely related, and therefore the makeup of the genetic contribution (i.e., nature) is relatively similar. This is especially important when testing the effect of the host genome on the microbiome across species, for example, by employing cross-fostering experiments. To date, such manipulations have been carried out across populations of the same species ^24–27^. Given that in social insects the gut environment and its symbionts play a significant role in nestmate recognition ^28–30^, colony labor division ^31, 32^ and maintained sociality ^28, 33–35^, it is reasonable to assume that the host-microbiome relationship is an important evolutionary feature that can attest to its relevance on a wider scale.

Therefore, to evaluate the role of nature vs. nurture in shaping the microbiome, we executed a cross-fostering experiment between two species of social wasps (family Vespidae), the Oriental hornet (*Vespa orientalis*) and the German wasp (*Vespula germanica*). We generated a uniform environment (i.e., nurture), with the two species representing two different “natures”. In these species, workers feed larvae with high-protein food and, in turn, obtain the majority of their own food from the oral secretions produced by larvae, through trophallaxis ^36^. By using young larvae (3rd instar) and newly emerged workers, we minimized the effect of previous environmental conditions on the microbiome. Because during metamorphosis, the larval gut content is expelled prior to pupation ^37^, newly emerged adults are thought to retain negligible gut microbiota ^38^. The absence of a microbiome at emergence negates effects of additional factors, such as life history traits or colony location, prior to microbiome establishment, leaving host identity as a robust proxy for measuring genetic effects.

We hypothesized that the significance of these factors will be reflected in differences in microbial community abundance and diversity. For example, if diet has the highest impact on the microbiome, most of the individuals will present with similar microbiome profiles regardless of their species or the identity of their counterpart (i.e., worker or larva). Inversely, if host identity affects the microbiome the most, microbial communities from individuals from the same species will be the most similar. Alternatively, if trophallaxis acts as a tool of mediating bacterial symbionts, workers and larvae from the same treatment (control or cross) will be more similar than their respective counterparts in other treatments. Similarly, the control treatments in this experimental design allow for identification of differences between life stages in *V. orientalis* and *V. germanica*.

## Methods

### Animal husbandry and collections

We collected wild colonies of two Vespid species, the Oriental hornet and the German wasp, at two locations (Oriental hornet – Kibbutz Beit Hashita, Israel, latitude: 32.550957, longitude: 35.438736; German wasp – the botanical garden at Tel Aviv University, Israel, latitude: 32.1141126, longitude: 34.8089881) using standard nest collection protocols ^39^. Each species was represented by a single colony to minimize the impact of neutral diversity and standardize experimental conditions. Both colonies included the founding queen, workers, and additional offspring (eggs, larvae, pupae). We reared the colonies in laboratory conditions (28°C, 70% relative humidity) in trapezoidal wooden boxes (14 L) with a Plexiglas front wall, supplied with *ad libitum* food (protein source and sugar solution (60% inverted sugar)) and water as well as cell building material (paper and soil). Three weeks prior to the beginning of the experiment, we standardized the feeding regime of both colonies by supplying the same type of protein daily (frozen bumble bees (*Bombus terrestris*) and raw organic chicken).

### Experimental design

To assess the contribution of species identity to the microbiome profile of larvae and workers, we conducted a cross-fostering experiment by which larvae of one species were nursed by workers of the second species. We also included controls (in which workers nursed their own larvae) (Figure 1). Accordingly, we carried out four treatment schemes: “control O” (*V. orientalis* workers nursing *V. orientalis* larvae), “control G” (*V. germanica* workers nursing *V. germanica* larvae), “cross O/G” (*V. orientalis* workers nursing *V. germanica* larvae) and “cross G/O” (*V. germanica* workers nursing *V. orientalis* larvae). Each treatment included four to six newly emerged workers and a piece of comb cut off from the wild colony that contained between 20-30 larvae in early stages (up to 3^rd^ instar). We selected only newly emerged workers as nurses because they emerge with a negligible microbiome, and can therefore be considered ‘tabula rasa’ for micr-obiota accumulation. Additionally, we selected only early-stage larvae to minimize the effect of previous interactions with nursing workers outside of the timeframe of the experiment and considered pre-treatment conditions in the statistical analysis by accounting for the number of days each larva had undergone treatment.

**Figure 1.**
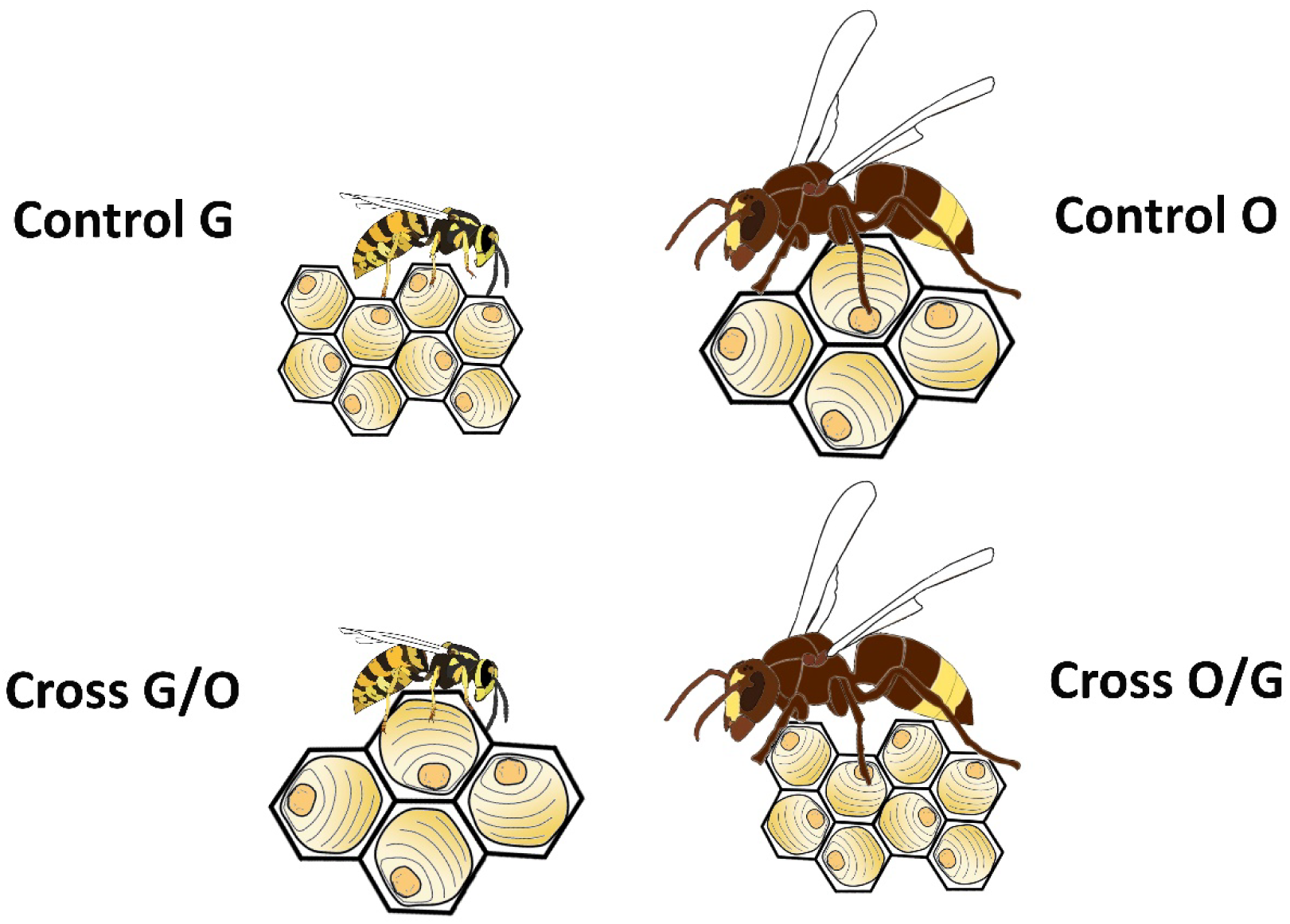
Experimental design of the four treatments in this study. ‘Control G’ included *V. germanica* workers and larvae; ‘Control O’ included *V. orientalis* workers and larvae; ‘Cross G/O’ included *V. germanica* workers and *V. orientalis* larvae; ‘Cross O/G’ included *V. orientalis* workers and *V. germanica* larvae. Sample size (n) is given for workers (‘W’) and larvae (‘LV’) separately as follows: Control G: n(W)=24, n(L)=12; Control O: n(W)=16, n(L)=18; Cross G/O: n(W)=40, n(L)=12; Cross O/G: n(W)=20, n(L)=5.

Environmental and dietary conditions on all four treatments were identical. We conducted all four treatments in small wooden boxes (14.5 × 12 × 10 cm) with glass doors, with a minimum of four replications for each treatment (additional replications were conducted if an experiment ended without collected larvae, e.g., if all larvae died). The combs containing the larvae were glued to the ceiling of the wooden box. We supplied the workers and larvae with a protein source (bumble bees and organic chicken – to avoid impact of antibiotics residues), a carbohydrate source (5ml tube containing sterilized sugar solution (60% inverted sugar)), and water (in a 5ml tube), all changed daily. We removed collected individuals from the treatment boxes daily and stored them at −20°C. We then replaced dead workers with other newly-emerged workers. We monitored surviving larvae and workers and egg hatching every day, and terminated experiments when all or most larvae had died (experiment length ranged between 5-39 days). Upon termination, we flash-froze all remaining workers and larvae and stored them at −20°C until further analysis. We only considered individuals for microbiome sequencing if they had survived four days or more in the treatment. We included workers even if no larvae had been collected from a particular treatment box, as long as nursing behavior had been observed.

### Microbiome analysis

We dissected the digestive tracts (mid- to hindgut) of the frozen workers and larvae in a sterile environment under a Plan S 1.0x FWD 81mm binocular (Zeiss, Discovery V12, Germany) and stored them at −80°C until further analysis. We extracted DNA from all samples (Table S1) using the PureLink Microbiome DNA Purification Kit (Invitrogen, Thermo Fisher; Waltham MA, USA) according to the manufacturer’s instructions and following mechanical crushing using sterile L-shaped spreaders and a 2 min bead beating step. We then PCR-amplified the variable V4 region of the 16S rRNA gene using the 515F-barcoded and 806R-non-barcoded primers ^40^. Each PCR reaction consisted of 25μL PrimeSTAR Max PCR mix (Takara Kusatsu, Shiga, Japan), 2 μM of each primer, 11 μL of ultra-pure water, and 10 μL DNA template. Thermal cycler conditions were as follows: 35 cycles of denaturation at 98°C for 10 sec, annealing at 55°C for 5 sec, and extension at 72°C for 20 sec, followed by a final elongation at 72°C for 1 min. We purified amplicons using Kapa Pure magnetic beads (Roche; Basel, Switzerland) and quantified them using the Picogreen dsDNA quantitation kit (Invitrogen, Thermo Fisher; Waltham, MA, USA). We then pooled equimolar amounts of DNA from individual samples and sequenced the pool using the Illumina MiSeq platform at the Genomic Center at the Bar-Ilan University Azrieli Faculty of Medicine in Tzfat, Israel. Appropriate negative and positive controls were included.

We initially processed the 16S rRNA gene sequence data with QIIME2 version 2020.8 ^41^ using default parameters. We used DADA2 ^42^ to filter noisy sequences, correct sequencing errors, remove chimeric sequences, remove singletons, and dereplicate sequences into amplicon sequence variants (ASVs). We assigned taxonomy using classify-sklearn naïve bayes classifier against GreenGenes ^43, 44^ and Silva138 databases ^45, 46^.

After the QIIME2 pipeline, we performed downstream analysis using phyloseq (version 1.34.0), R/bioconductor package for handling and analysis of high-throughput phylogenetic sequence data ^47^. We used only samples that were taken from wasps cross-fostered more than 4 days prior to collection for the analysis. First, we cleaned the taxonomy and filtered out empty taxa. We then rarefied the samples (using rarefy_even_depth function) to a minimum sequence depth of 1,000 and scaled by relative abundance; samples with fewer reads were removed.

We examined patterns of alpha-diversity (Faith’s Phylogenetic Diversity (PD) using the twbattaglia/btools R package ^48^) function) and beta-diversity (weighted UniFrac, phyloseq distance function) for different groups of samples using the Kruskal-Wallis test and post hoc Dunn’s multiple comparison test or FDR-corrected pair-wise PERMANOVAs, respectively. To assess gut microbiome differential abundance, we used DESeq2 (version 1.30.1), R/bioconductor package ^49^; significant taxa were those with an adjusted p-values<0.05 and a |log2foldchange| ≥0.58. We generated principal component analysis using the plotPCA function in the DESeq2 package and heatmaps using pheatmap (version 1.0.12^50^). When more than two conditions were compared, Kruskal-Wallis tests with Dun post-hocs were performed on the relative abundances of the multiple groups for taxa identified as differentially abundant overall by DESeq2. In addition, we analyzed the effect of the number of days in the experiment to test the change of the composition and diversity of the microbiome as a factor of time.

## Results

### Experimental and sequencing success

We conducted 20 replicates of the four treatments, in which a total of 98 workers and 47 larvae were collected (Table S1). The treatment in which workers of the bigger species, *V. orientalis*, nursed larvae of the smaller species, *V. germanica* (Cross O/G), yielded the smallest number of larvae (n=9). We posit that this occurred because the *V. germanica* comb cell size is much smaller than that of *V. orientalis*, which may have physically prevented Oriental hornets from properly reaching the early-stage, small German wasp larvae. In two out of the seven replicates of the cross treatment where German wasp workers nursed Oriental hornet larvae (Cross G/O), the workers destroyed the entire comb, leading to termination of the experimental replicate. Similarly, in one replicate of the cross treatment where Oriental hornet workers nursed German wasp larvae, no larvae survived, and the replicate was not included in the analysis.

Out of the 145 individual samples available for sequencing, 74 workers and 35 larvae were available for analysis, representing all four treatments. However, due to the lower number of available samples for *V. germanica* larvae from the cross treatment, only 3 individuals were successfully sequenced. Interestingly, the length of time each individual participated in the experiment (i.e., the factor of ‘Days’) was statistically significant in explaining differences in beta diversity in workers from both species, regardless of treatment (weighted UniFrac^51^: F = 6.01, R2 = 0.08, *P* = 0.006), but not in larvae (weighted UniFrac: F = 0.79, R2 = 0.02, *P* = 0.497), suggesting that microbiome diversity increased with time in the experiment for workers (Figure S1).

### Microbiome diversity of nursing workers

To test whether the microbiome of newly emerged nurses is shaped by nurture (i.e., diet) or nature (i.e., self-identity, identity of the nursed larvae), we examined whether workers from different treatments are characterized by different gut microbial communities. We found that alpha diversity did not differ significantly between workers from different treatment groups, regardless of their identity or the identity of the larvae they nursed (Faith PD^52^, *P* = 0.602, Figure 2A). Additionally, microbial beta diversity was not significantly different among workers (weighted UniFrac: F = 1.90, R^2^ = 0.08, *P* = 0.111, Figure 2B).

**Figure 2.**
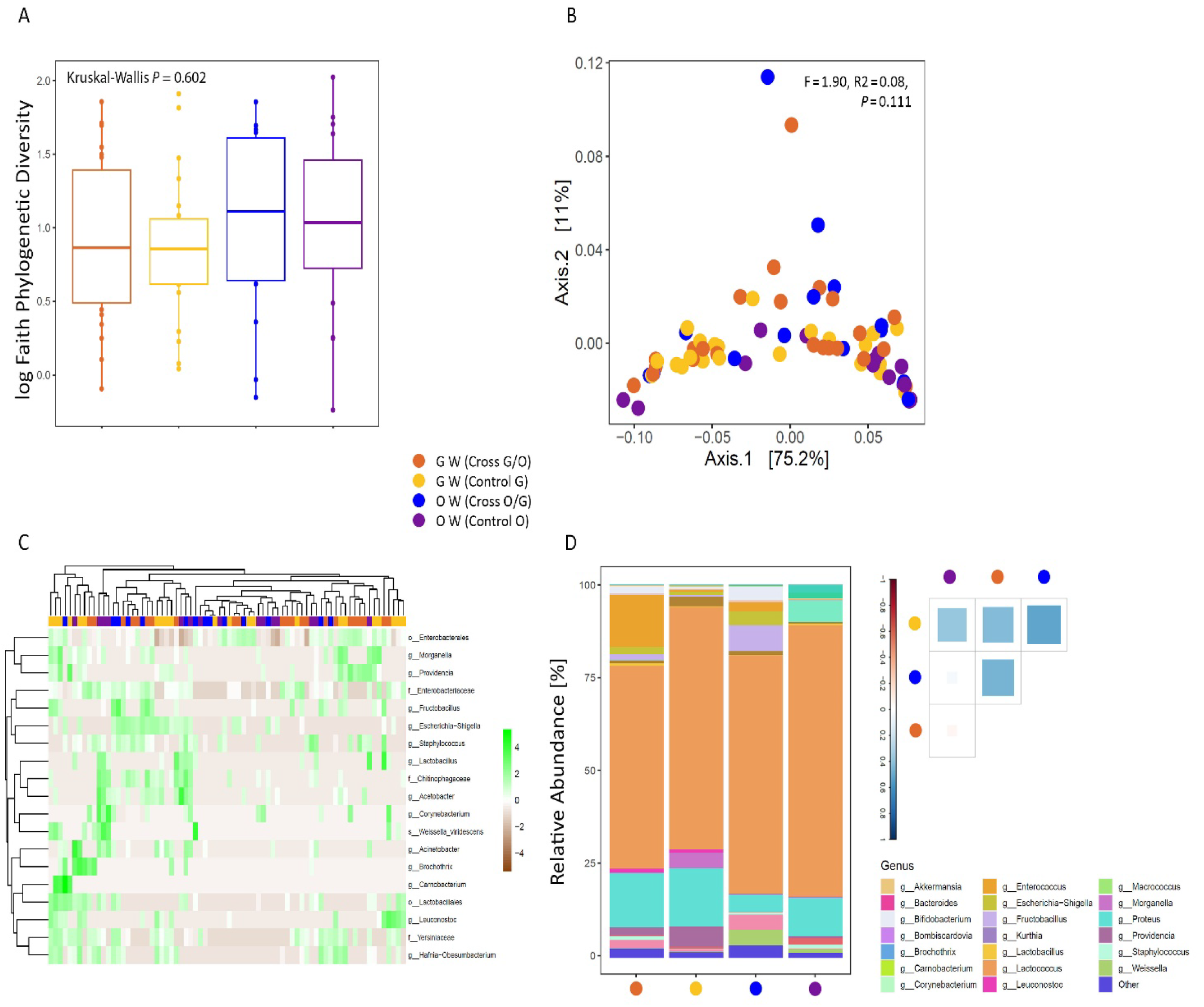
Microbiome diversity and composition in nursing workers subjected to each of the four treatments. (A) Faith’s phylogenetic (alpha) diversity (log). Differences between treatments, compared via the Kruskal-Wallis test, were statistically insignificant (P = 0.602). (B) Weighted UniFrac (beta) Principal Coordinate Analysis (PCoA) of the bacterial composition (P = 0.111). (C) Genus, Family, Order and Class-level shifts in workers from control and cross treatments. Heat map showing transformed (log2 base) normalized read counts of taxa found to be differentially abundant based on DESeq2 analysis (adjusted value of P < 0.05). (D) Relative abundance of the twenty most abundant genera and quantified similarity among the microbiome profiles. Taxonomic abundances from gut samples of the same treatment were averaged together. Spearman’s rank correlation coefficient was used to estimate similarity between treatments.

To better understand how the microbial community changes in response to intrinsic and extrinsic conditions, we identified altered abundance of specific microbial orders, families, genera and species as a function of treatment using DESeq2. A total of 19 taxa (2 orders, 3 families, 13 genera and 1 species) were found to be differentially abundant between workers of both species from the four different treatments when compared to German wasp workers from the control treatment (adjusted *P* < 0.05,Figure 2C, Table S2). Visual examination of heatmaps, however, did not reveal treatment-specific patterns of differential abundance (Figure 2C).

We further examined similarity patterns among workers. Similarity among workers, regardless of their treatment, was high (average Spearman’s rank correlation coefficient *r* = 0.30; Figure 2D). Relative abundance profiles of workers from different treatments showed similar patterns, dominated largely by the genera *Lactococcus* and *Proteus* (Figure 2D). At the phylum level, Firmicutes and Pseudomonadota dominated the taxa shared by workers from all four treatments, and of these, orders Lactobacillales and Enterobacterales were highly abundant (mean relative abundance >1% in at least one group, Table S3).

### Microbiome diversity of larvae

Similarly, we aimed to assess whether larvae from different treatments shared similar gut microbial communities. Alpha diversity did not differ significantly between larvae from different treatment groups (Faith PD, *P* = 0.193, Figure 3A). Inversely, beta diversity was significantly different among larvae from different treatments (weighted UniFrac: F = 3.36, R2 = 0.25, *P* = 0.009, Figure 3B). However, the degree of change varied between the groups: whereas Oriental hornet larvae did not significantly differ regardless of the identity of the nursing worker (i.e., ‘treatment’; F = 2.87, R2 = 0.14, *P* = 0.058), German wasp larvae nursed by German wasp workers differed significantly from German wasp larvae nursed by Oriental hornet workers (F = 5.26, R2 = 0.29, *P* = 0.025; but N=3). Additionally, although larvae of different species mostly differed significantly, German wasp larvae from the control treatment did not differ significantly from Oriental hornet larvae from the control treatment (Table S4).

**Figure 3.**
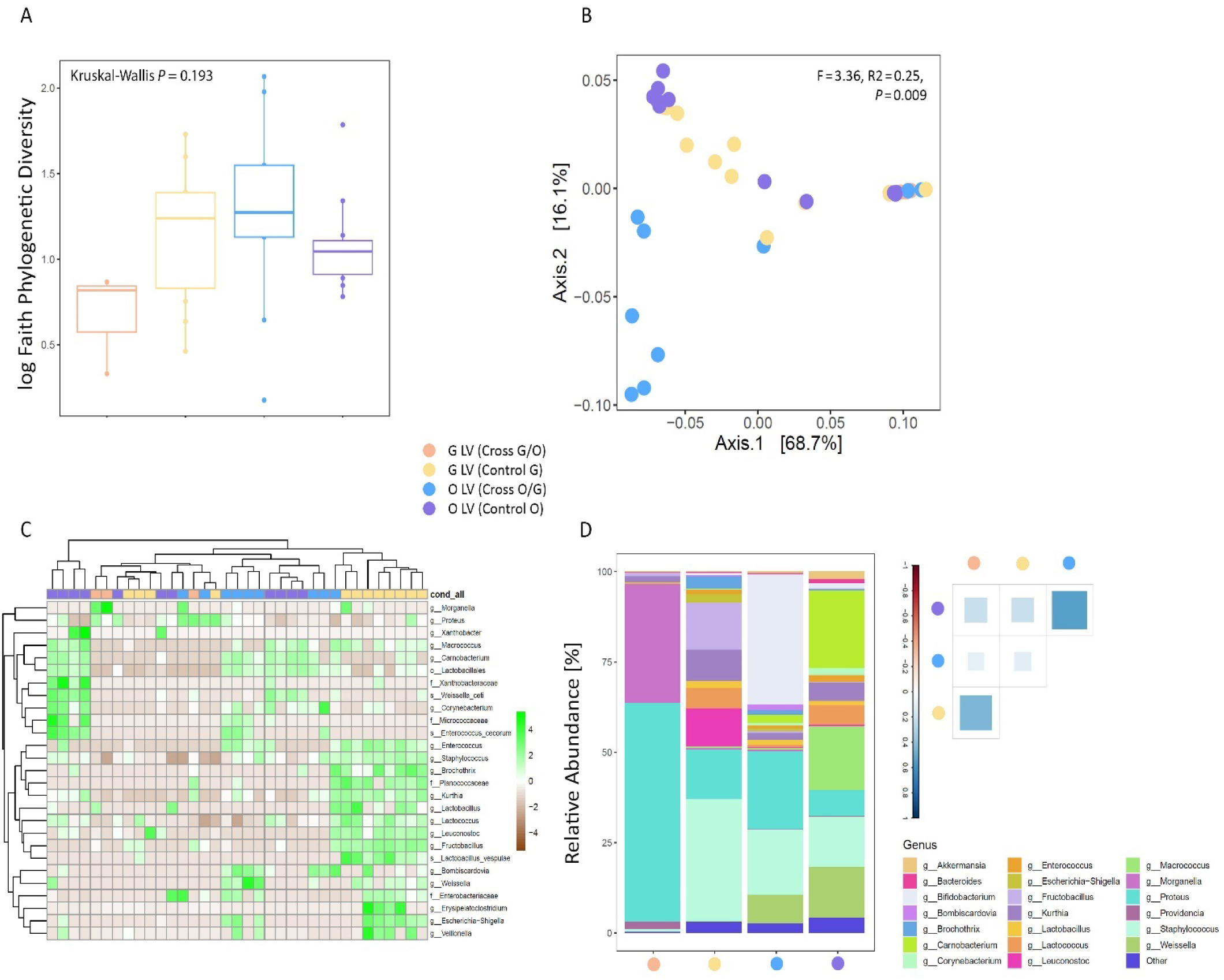
Microbiome diversity and composition in larvae subjected to each of the four treatments. (A) Faith’s phylogenetic (alpha) diversity (log). Differences between treatments, compared via the Kruskal-Wallis test, were statistically insignificant (P = 0.193). (B) Weighted UniFrac (beta) Principal Coordinate Analysis (PCoA) of the bacterial composition of larvae compared with PERMANOVA (P = 0.009). Results of post-hoc pairwise PERMANOVAs are in Table S4. (C) Genus, Family, Order and Class-level shifts in larvae from control and cross treatments. Heat map showing transformed (log2 base) normalized read counts of taxa found to be differentially abundant based on DESeq2 analysis (adjusted value of *P* < 0.05). (D) Relative abundance of the twenty most abundant genera and quantified similarity among the microbiome profiles. Taxonomic abundances from gut samples of the same treatment were averaged together. Spearman’s rank correlation coefficient was used to estimate similarity between treatments.

Differences among larvae from different treatments were also observed when examining differential abundance of bacterial taxa and reflected alterations relevant to each treatment. A total of 27 taxa (1 order, 4 families, 19 genera and 3 species) were found to be differentially abundant between larvae from the four different treatments when compared to German wasp larvae from the control treatment (adjusted *P* < 0.05; Figure 3C, Table S5). The vast majority of these taxa differed between Oriental hornet larvae from both treatments and German wasp larvae, and largely belonged to the phyla Firmicutes (16 taxa), Pseudomonadota (6 taxa) and Actinomycetota (3 taxa).

When we explored patterns of similarity among larvae from the four different treatments, a mixed pattern emerged. Whereas larvae from the same species tended to have more similar microbial communities (Spearman’s rank correlation coefficient between larvae from the same species in *V. orientalis* and *V. germanica r* = 0.54 and *r* = 0.46, respectively), some similarity was also found among larvae from the different species (Table S6; Figure 3D). Similar to nursing workers, Firmicutes and Pseudomonadota were the dominant phyla shared by larvae from all four treatments, and orders Lactobacillales, Bacillales and Enterobacterales were highly abundant (mean relative abundance >1% in at least one group, Table S7). Subsequently, we examined relative abundance patterns of specific taxa shared among larvae groups. Oriental hornet larvae shared a significantly elevated relative abundance of taxa from the Lactobacillales order (Kruskal-Wallis, *P* < 0.001, Figure 4A) and the genus *Carnobacterium* (Kruskal-Wallis, *P* < 0.001, Figure 4B). Likewise, the relative abundance of bacteria from the *Kurthia* genus was higher only in German wasp larvae (Kruskal-Wallis, *P* < 0.001, Figure 4C).

**Figure 4.**
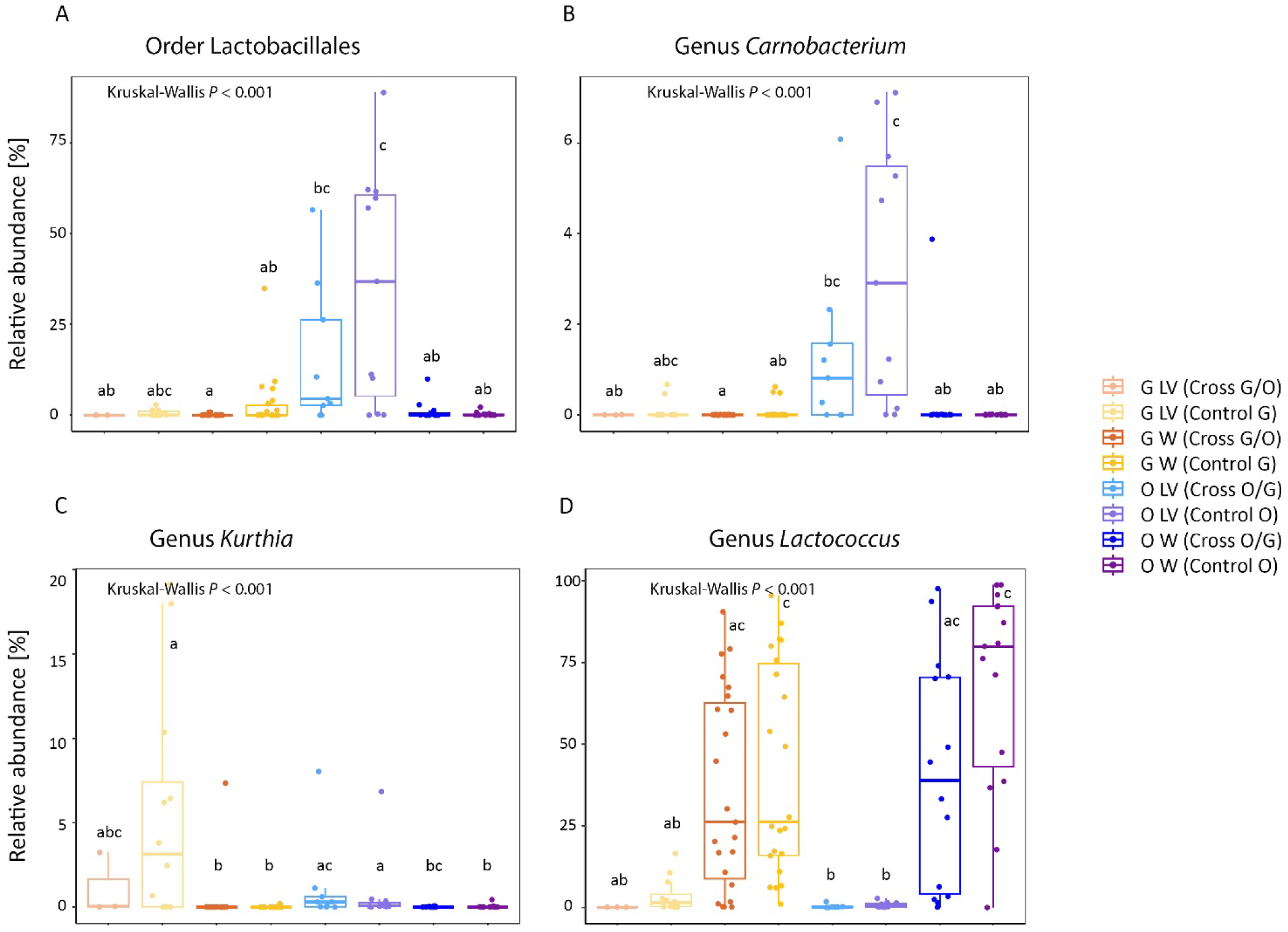
Bacterial taxa with a significantly higher relative abundance in larvae (‘LV’) of Oriental hornets (A, B), larvae of German wasps (C), workers (‘W’) of all four treatments (D) and workers and larvae of German wasps (E). Differences between treatments were compared with Kruskal-Wallis test and the post-hoc Dunn’s multiple comparison tests after DESeq2 identified significant taxa. Lowercase letters above the boxes show significant differences in relative abundance values between groups.

### Species core microbiota

To identify the core microbiota of *V. orientalis* and *V. germanica*, we searched for similarities among individuals of the same species, in each life stage. To minimize effects related to the experimental design, we analyzed only individuals from the control treatments. Similarly, because of the significant differences found between life stages, we examined each life stage separately. The microbiome composition of both species was fairly similar, particularly in nursing workers (Table 1). In nursing workers of both *V. orientalis* and *V. germanica*, the most relatively abundant taxon was the genus *Lactococcus* (52.8% and 32.3%, respectively), and both also had high abundances of the genus *Proteus* (8.1% and 7.8%, respectively) and an unclassified genus in the order Enterobacterales (3.1% and 19.0%, respectively). These taxa were also very prevalent (> 87%, Table 1). In German wasp workers, other relatively abundant taxa included unclassified genus in the order Lactobacillales (5.3%, prevalence 45%) and the families *Enterobacteriaceae* and *Yersiniaceae* (4.5% and 5.1%, with prevalence of 82% and 64%, respectively), whereas in Oriental hornet workers, they included the genera *Gilliamella* (11.9%, prevalence 33%), *Pseudomonas* (2.6%, prevalence 47%) and *Acetobacter* (3.2%, prevalence 47%).

**Table 1.**
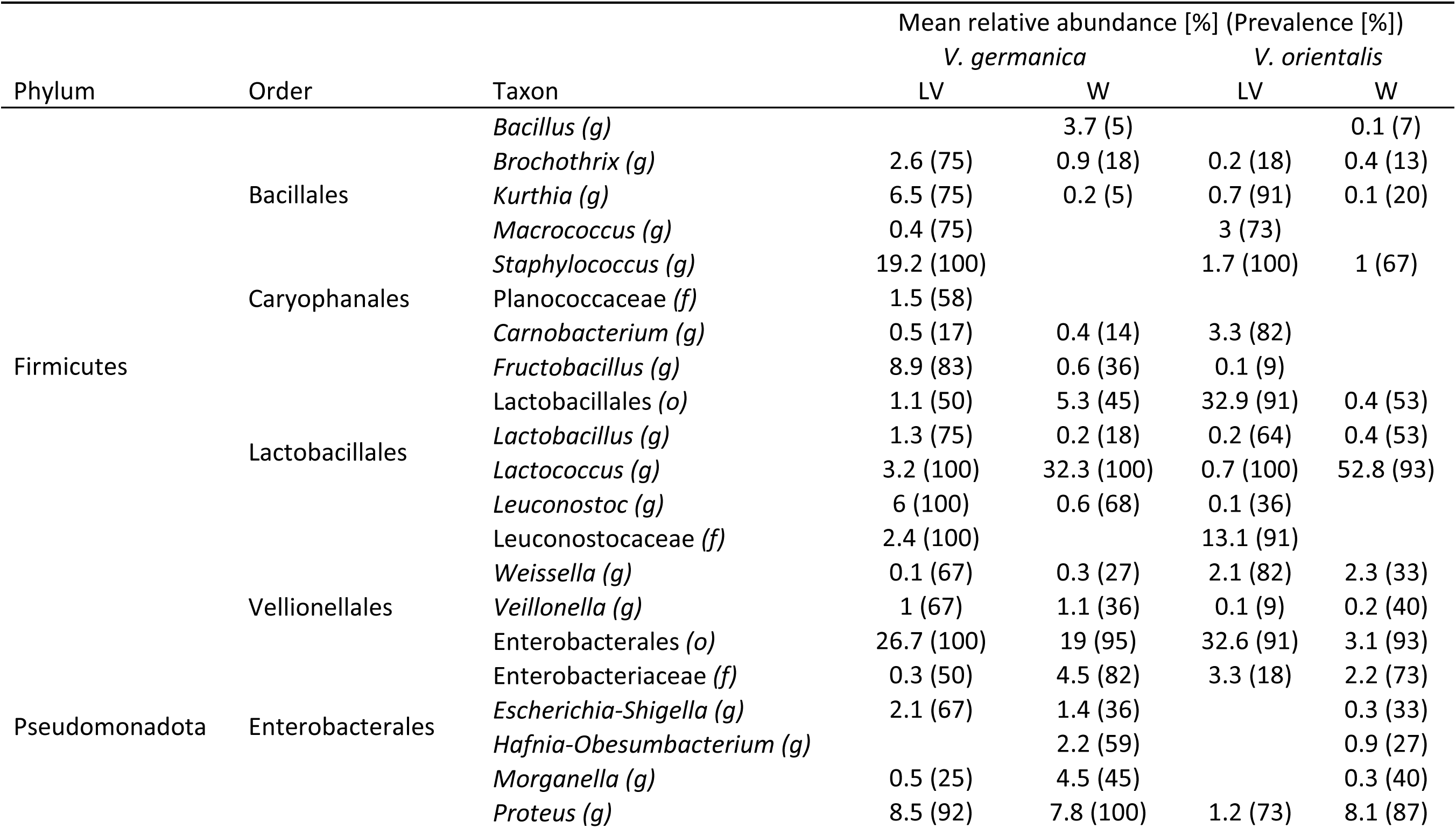
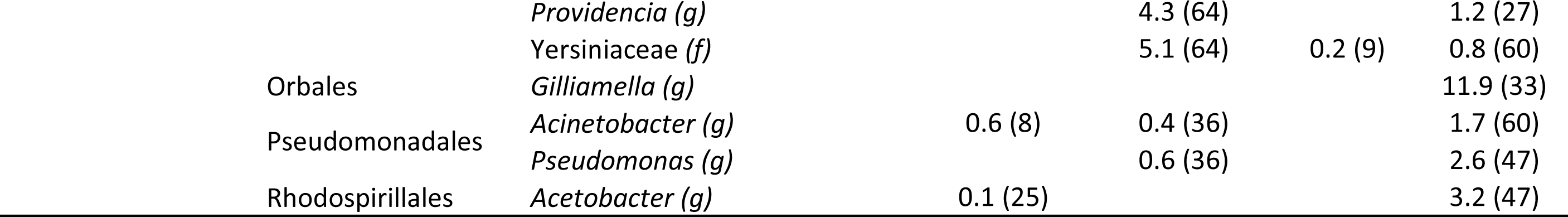
Microbiome composition of *V. germanica* and *V. orientalis* in larvae and in workers. Only taxa with mean relative abundance of 0.1% and above are shown. Prevalence in the group is shown in parentheses. Letters after the taxon in parentheses represent the taxonomic level at which that organism was identified (f – family, o – order, g – genus). LV-larvae, W-workers.

In larvae, the only abundant and prevalent (prevalence > 91%) taxon shared among both species was an unclassified genus in the order Enterobacterales (mean relative abundance 27.6% in *V. germanica*, 32.6% in *V. orientalis*). The remaining most relatively abundant taxa were genera *Staphylococcus* (19.2%, prevalence 100%), *Fructobacillus* (8.9%, prevalence 83%), *Proteus* (8.5%, prevalence 92%), *Kurthia* (6.5%, prevalence 75%), and *Leuconostoc* (6.0%, prevalence 100%) in the German wasp larvae, and order Lactobacillales (32.9%, prevalence 91%), family *Leuconostocaceae* (13.1%, prevalence 91%), and genus *Carnobacterium* (3.3%, prevalence 82%) in the Oriental hornet larvae.

### Microbiome transfer between workers and larvae

To characterize the microbiome profiles of larvae and workers and the role of trophallaxis in maintaining them, we analyzed Oriental hornets and German wasps separately to minimize variation stemming from the species-specific core microbiome. Beta diversity was significantly different between larvae and workers, both in Oriental hornets (weighted UniFrac: F = 4.51, R2 = 0.16, *P* = 0.021, Figure 5A) and in German wasps (F = 4.35, R2 = 0.12, *P* = 0.031, Figure 5B).

**Figure 5.**
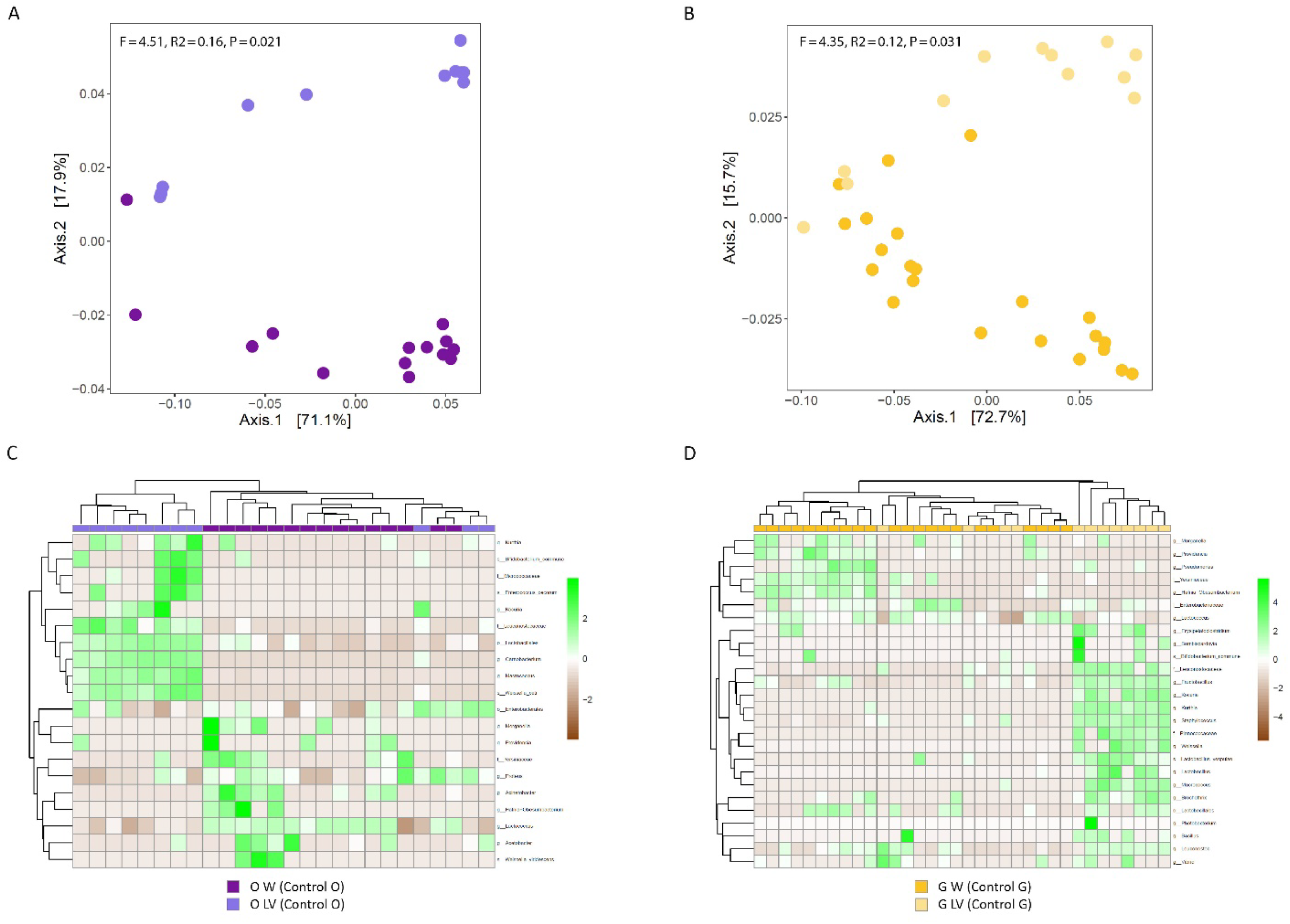
Differences between nursing workers and larvae in Oriental hornets (A, C) and German wasps (B, D). Weighted UniFrac Principal Coordinate Analysis (PCoA) of the bacterial composition of larvae and nursing workers of *V. orientalis* (*P* = 0.021; A) and *V. germanica* (*P* = 0.031; B). Heat map showing transformed (log2 base) normalized read counts of taxa found to be differentially abundant between larvae and nursing workers based on DESeq2 analysis (adjusted value of *P* < 0.05) are shown for *V. orientalis* (C) and *V. germanica* (D). Larvae (light) and workers (dark) are shown in purple and yellow shades in Oriental hornets and German wasps, respectively.

Differences between life stages were also evident in differential abundance patterns of specific microbial orders, families, genera and species. A total of 20 taxa (2 orders, 3 families, 11 genera and 4 species) and 26 taxa (1 order, 4 families, 19 genera and 2 species) were differentially abundant between workers and larvae in Oriental hornets (Figure 5C, Table S8) and German wasps (adjusted *P* < 0.05, Figure 5D, Table S9), respectively. In Oriental hornets, the majority of the taxa that were differentially more abundant in larvae belonged to classes Bacilli and Actinomycetia (10 out of 11 taxa), whereas in workers they belonged to the class Gammaproteobacteria (6 out of 9 taxa). In German wasps, taxa more abundant in larvae belonged mainly to class Bacilli (13 out of 18 taxa), whereas six out of eight taxa belonged to class Gammaproteobacteria in workers.

When analyzing relative abundance patterns of specific taxa shared among life stage, we found that the relative abundance of the genus *Lactococcus* was significantly higher in workers of both *V. orientalis* and *V. germanica* compared with larvae (Kruskal-Wallis, *P* < 0.001, Figure 4D).

## Discussion

### The role of the environment in shaping the gut microbiome of social insects

In this study we explored the role of the environment in shaping species microbiome composition in comparison with the host identity using a setting of cross-species nursing of larvae. Newly emerged workers, whose microbiome is depleted during metamorphosis ^53, 54^, provided a tabula rasa to test the establishment of a new gut microbiome. Worker microbiomes were largely shaped by diet, resulting in similar microbiome profiles for workers regardless of species and the identity of the larvae they were nursing. Many holometabolous insects show similar patterns of elimination of the larval gut and its contents as a meconium sac and subsequent acquisition of microbes from their environment, including mosquitos, honeybees, wasps and hornets ^37, 54–56^. The degree to which the gut community is acquired from the environment in each generation is largely thought to be associated with host niche specialty ^53^, but both facultative and obligatory bacteria can be acquired through horizontal transfer. For example, trophallaxis among siblings enables transmission of beneficial microbes in the honeybee (*Apis mellifera*) ^57, 58^, a mechanism that can plausibly occur in both *V. orientalis* and *V. germanica*. The immediate environment and its available microbes can also be shaped by members of the species, e.g., by defecating or regurgitating into food sources (e.g. vinegar flies ^59, 60^). We have observed worker-to-worker trophallaxis in both of our study species, suggesting a role for adult trophallaxis in maintaining a core or colony-specific microbiome, but directly testing adult saliva is necessary for determining this conclusively. By experimentally controlling their diet, we have shown that the microbiome composition of adult Oriental hornets and German wasps depended principally on the environment. Diet also had a clustering effect on the microbial communities of larvae, but to a lesser extent.

The main bacterial taxa found to be associated with the nursing worker microbiome were from the genera *Lactococcus* and *Proteus* and the order Enterobacterales, possibly playing an important role in assisting in the metabolism of nutrients in the workers’ carbohydrate-rich diet. *Lactococcus* bacteria produce lactic acid by fermenting glucose, which is one of the primary nutrients in the diet of adult hornets and wasps ^61^, whereas some *Proteus* species were shown to hydrolyze cellulose and xylan ^62–64^. The order Enterobacterales was also found to be one of the two most dominant orders in *Vespa mandarinia* and, to a lesser extent, in *Vespa simillima* ^65^. Although studies detailing hornet and wasp microbiomes are still scarce, our results correspond with taxa shown to be abundant in closely related species with similar diets, such as *Lactoccoccus, Gilliamella, Fructobacillus, and Leuconostoc* ^37, 65, 66^. Our results also confirm the contention put forward by Reeson et al. ^67^ that *V. germanica* is not dependent on a specific set of symbionts. However, it is likely that because of the substantial effect of diet on the microbiome of workers and the change in its composition we detected across time (Figure S1), the different environmental settings in these studies modified the microbial gut communities of these species, to some extent.

### Vertical transfer mediates host identity impact on microbiome diversity

Despite the imminent effect of diet on the microbiome environment in holometabolous insects, vertical (mother-to-offspring) transmission of gut microbes is crucial in some species, especially those with symbionts that confer niche-specializing abilities on their hosts ^38, 68^. Our results plausibly reflect evidence for this mechanism in larvae, where clustering was species-dependent regardless of the identity of the nursing worker in the majority of the samples (although variation indicating involvement of other factors manifested in a third, mixed-species clustering). One of the main caveats for this study was using larvae instead of eggs as the recipients for the nursing workers’ care. Had we been able to successfully carry out the experiments with eggs (which were expected to have a low bacterial load, if any ^65, 66^), we would have further minimized the effect of an already established microbiome and provided a cleaner slate to test colonization and establishment of gut symbionts. However, in our preliminary work, workers either ate the eggs or refrained from nursing them over the course of a few days, resulting in colony failure, thus leading us to use larvae. Having been previously fed at their original colonies, these larvae, however young, may have already carried a microbial community. Despite this, larval age did not explain the variation in microbiome diversity. Nevertheless, the clustering of same-species larvae suggests an involvement of an intra-specific vertical transfer of at least some symbionts.

Several mechanisms of vertical transfer have been suggested in holometabolous insects. In eusocial species, where overlap of generations in a shared space normally occurs, trophallaxis has been suggested to be the primary cause of maintenance of host-specific microbial communities in termites ^69, 70^, but not in bumble bees ^71^, and also extends to burying beetles (*Nicrophorus vespilloides*) ^72^. We did not find evidence to support trophallaxis transfer of microbiota between nursing workers and larvae in the context of this study, but it is possible that in a natural setting, where additional castes are present, some core microbiota are transferred to offspring in this manner. Particularly, in the context of the life history of both *V. orientalis* and *V. germanica* (both species have annual colonies initiated by a single overwintered queen), it is possible that the foundress queen transfers her microbiome via trophallaxis onto the first generation of workers (e.g., in bumblebees ^71^, who will in turn continue this inter-generational transmission.

Alternatively, but not mutually exclusively, vertical transfer can occur in the colony or nest environment via deposition of oral or anal secretions onto shared food substrate or comb material. In the burying beetle *Nicrophorus vespilloides*, adults deposit oral and anal secretion on a carcass used as a food source for its developing larvae ^72, 73^. In bumble bees, the hive environments (e.g., cocoon material and feces) are considered to be the primary sources for inter-generational transmission of microbiota ^71^. Other alternatives include association with the female oocytes (e.g., in weevils ^74^ and the thistle tortoise beetle (*Cassida rubiginosa)* ^75^), inoculation of the foam plug through which larvae pass while hatching (e.g., in desert locust, *Schistocerca gregaria* ^76^), and via glands (e.g., the milk glands of female tsetse flies (*Glossina)* ^74^). The direct mechanism of vertical transfer in both *V. orientalis* and *V. germanica* remains unclear, and further exploration of the colony components, life stages and castes will help elucidate this. However, differences stemming from the decoupling of life stages due to nutritional requirements will most likely impact the microbial interactions, even if some bacteria are transferred between the guts of workers and larvae reciprocally, resulting in selection for symbionts fit for their respective host.

### Microbiome importance in social insects

The seemingly conflicting degree of impact of the environment and host identity on the two life stages in Oriental hornets and German wasps suggests that their gut symbionts may have a wide range of host effects. Primarily, through reciprocal symbiotic nutrient metabolism, microbiomes specific to life stage or caste complement the division of labor in eusocial insects ^31^. Equally, host sociality enables the maintenance of stable microbial communities by providing reliable transmission routes between hosts of similar environments and nutritional requirements^77^. Division of labor in eusocial insects allows for optimal allocation of tasks between castes thus improving resource utilization, minimizing inter-individual competition, increasing colony fitness and possibly leading to their evolutionary success ^78^. Our results provide evidence of differences between life stages in two additional species of eusocial insects and support the hypothesis that microbial symbionts facilitate evolutionary behavioral adaptations such as adult and larval co-dependency on trophallaxis. It is unclear how much of their respective niche hosts can exploit using solely their own enzymes and what the extent of contribution their microbial symbionts provide is, but it is generally considered that gut symbionts allow their host to expand their potential niche ^79–81^. The high degree of shared taxa between these two species and the presence of various bacterial taxa in previously studied closely related wasps and hornets, however, suggests that this symbiotic relationship is not exclusive on either end.

Our results emphasize the importance of diet on gut microbial diversity in the adult stage, with some indication that host identity is relevant in the larval stage via vertical transfer. These findings contribute to further elucidation of the complexity of the host-microbiome relationship shaped, to a large extent, by the environment while retaining symbionts that benefit the host despite metamorphosis. By employing an experimental design that enabled us to control the environment and create direct cross-species nursing contact, we have shown this mechanism is in place in a multi-species setting, and therefore we suggest that this complex interplay likely shapes the gut microbiome of other eusocial species as well, especially those with a complete decoupling of life stages. Future work will likely shed light on modes of transfer as well as on the prevalence of the effect of extrinsic and intrinsic factors on different castes in the superorganism. These, and other evidence of host-microbiome symbiosis emerging from other species ^82, 83^, indicate that holobiont evolution (i.e., host-microbiome co-evolution) may have promoted the rise of social behavior in animals.

## Supporting information

metadata

## Funding

This research was supported by ISF grant #1538/18. TMC was supported by the Tel Aviv University’s rector’s emergency Corona fellowship and the Council for Higher Education (CHE) postdoctoral fellowship.

## Author contributions

TMC and EL conceptualized the idea for the study; TMC, LB, ST, EF, AC, SB and ET conceived and executed the experiments and subsequent analyses; EL and OK supervised; TMC, LB, ST, EF and AC wrote the original draft; TMC, LB, ST, EF, AC, SB, ET, OK and EL edited the draft.

## Competing interests

The authors declare that they have no competing interests.

## Materials & Correspondence

All data needed to evaluate the conclusions in the paper are present in the paper and/or the Supplementary Materials. Additional data related to this paper may be requested from the corresponding authors TMC and EL.

## Notes

https://biu365-my.sharepoint.com/:f:/g/personal/labkore1_biu_ac_il/ElkoVmUsaXhFstOCcOO_RBgBFsOVcw8rDjmg--cLgKTTgA?e=Ord1PI

### Competing Interest Statement

The authors have declared no competing interest.

## References

1. Foster, K. R., Schluter, J., Coyte, K. Z. & Rakoff-nahoum, S. The evolution of the host microbiome as an ecosystem on a leash. Nature 548, 43–51 (2017).

2. Goodrich, J. K., Davenport, E. R., Waters, J. L., Clark, A. G. & Ley, R. E. Cross-species comparisons of host genetic associations with the microbiome. Science 352, 532–535 (2016).

3. Hauffe, H. C. & Barelli, C. Conserve the germs: the gut microbiota and adaptive potential. Conservation Genetics 20, 19–27 (2019).

4. Berg, G. et al. Microbiome definition re-visited: old concepts and new challenges. Microbiome 8, 1– 22 (2020).

5. Spor, A., Koren, O. & Ley, R. Unravelling the effects of the environment and host genotype on the gut microbiome. Nat Rev Microbiol 9, 279–290 (2011).

6. Sanders, J. G. et al. Stability and phylogenetic correlation in gut microbiota: lessons from ants and apes. Molecular Ecology 23, 1268–1283 (2014).

7. Turnbaugh, P. J. et al. The Human Microbiome Project. Nature 449, 804–810 (2007).

8. Malacrinò, A. Host species identity shapes the diversity and structure of insect microbiota. Molecular Ecology 31, 723–735 (2022).

9. Davenport, E. R. et al. Genome-Wide Association Studies of the Human Gut Microbiota. PloS one 10, e0140301 (2015).

10. Knights, D. et al. Complex host genetics influence the microbiome in inflammatory bowel disease. Genome Med 6, 107 (2014).

11. Bonder, M. J. et al. The effect of host genetics on the gut microbiome. Nat Genet 48, 1407–1412 (2016).

12. Blekhman, R. et al. Host genetic variation impacts microbiome composition across human body sites. Genome Biol 16, 191 (2015).

13. Lewis, J. D. et al. Inflammation, Antibiotics, and Diet as Environmental Stressors of the Gut Microbiome in Pediatric Crohn’s Disease. Cell Host & Microbe 18, 489–500 (2015).

14. Douglas, G. M., Bielawski, J. P. & Langille, M. G. I. Re-evaluating the relationship between missing heritability and the microbiome. Microbiome 8, 1–8 (2020).

15. Rothschild, D. et al. Environment dominates over host genetics in shaping human gut microbiota. Nature 555, 210–215 (2018).

16. Zmora, N., Suez, J. & Elinav, E. You are what you eat: diet, health and the gut microbiota. Nat Rev Gastroenterol Hepatol 16, 35–56 (2019).

17. Grieneisen, L. et al. Gut microbiome heritability is nearly universal but environmentally contingent. Science 373, 181–186 (2021).

18. Berasategui, A. et al. The gut microbiota of the pine weevil is similar across Europe and resembles that of other conifer-feeding beetles. Mol Ecol 25, 4014–4031 (2016).

19. Kudo, R., Masuya, H., Endoh, R., Kikuchi, T. & Ikeda, H. Gut bacterial and fungal communities in ground-dwelling beetles are associated with host food habit and habitat. ISME J 13, 676–685 (2019).

20. Valdivia, C. et al. Microbial symbionts are shared between ants and their associated beetles. bioRxiv 2022.12.02.518891 (2023) doi:10.1101/2022.12.02.518891.

21. Ronchetti, F., Polidori, C., Schmitt, T., Steffan-Dewenter, I. & Keller, A. Bacterial gut microbiomes of aculeate brood parasites overlap with their aculeate hosts’, but have higher diversity and specialization. FEMS Microbiology Ecology 98, fiac137 (2022).

22. Surana, N. K. & Kasper, D. L. Moving beyond microbiome-wide associations to causal microbe identification. Nature 552, 244–247 (2017).

23. Teyssier, A. et al. Diet contributes to urban-induced alterations in gut microbiota: experimental evidence from a wild passerine. Proc. R. Soc. B. 287, 20192182 (2020).

24. Daft, J. G., Ptacek, T., Kumar, R., Morrow, C. & Lorenz, R. G. Cross-fostering immediately after birth induces a permanent microbiota shift that is shaped by the nursing mother. Microbiome 3, 17 (2015).

25. Teyssier, A., Lens, L., Matthysen, E. & White, J. Dynamics of Gut Microbiota Diversity During the Early Development of an Avian Host: Evidence From a Cross-Foster Experiment. Front. Microbiol. 9, 1524 (2018).

26. Bian, G. et al. Age, introduction of solid feed and weaning are more important determinants of gut bacterial succession in piglets than breed and nursing mother as revealed by a reciprocal cross-fostering model: Gut bacterial succession in piglets. Environ Microbiol 18, 1566–1577 (2016).

27. Mao, J., Zhang, Y., Liu, J. & Wang, H. Gut microbiota and growth performance of offspring are influenced by wet nurse in pigs using cross-fostering trial. J Sci Food Agric 103, 865–876 (2023).

28. Vernier, C. L. et al. The gut microbiome defines social group membership in honey bee colonies. Science Advances 6, eabd3431 (2020).

29. Matsuura, K. Nestmate recognition mediated by intestinal bacteria in a termite, *Reticulitermes speratus*. Oikos 92, 20–26 (2001).

30. Lizé, A., McKay, R. & Lewis, Z. Gut microbiota and kin recognition. Trends in Ecology & Evolution 28, 325–326 (2013).

31. Sinotte, V. M. et al. Synergies Between Division of Labor and Gut Microbiomes of Social Insects. Frontiers in Ecology and Evolution 7, 0–9 (2020).

32. Otani, S. et al. Gut microbial compositions mirror caste-specific diets in a major lineage of social insects. Environmental Microbiology Reports 11, 196–205 (2019).

33. Liberti, J. & Engel, P. The gut microbiota — brain axis of insects. Current Opinion in Insect Science 39, 6–13 (2020).

34. Sarkar, A. et al. The role of the microbiome in the neurobiology of social behaviour. Biol Rev 95, 1131–1166 (2020).

35. Montiel-Castro, A. J., González-Cervantes, R. M., Bravo-Ruiseco, G. & Pacheco-López, G. The microbiota-gut-brain axis: neurobehavioral correlates, health and sociality. Front. Integr. Neurosci. 7, (2013).

36. Matsuura, M. & Yamane, S. Biology of the vespine wasps. (Springer Verlag, 1990).

37. Cini, A. et al. Gut microbial composition in different castes and developmental stages of the invasive hornet Vespa velutina nigrithorax. Science of The Total Environment 745, 140873 (2020).

38. Hammer, T. J. & Moran, N. A. Links between metamorphosis and symbiosis in holometabolous insects. Phil. Trans. R. Soc. B 374, 20190068 (2019).

39. Volov, M. et al. The Effect of Climate and Diet on Body Lipid Composition in the Oriental Hornet (Vespa orientalis). Front. Ecol. Evol. 9, 755331 (2021).

40. Caporaso, J. G. et al. Ultra-high-throughput microbial community analysis on the Illumina HiSeq and MiSeq platforms. ISME Journal 6, 1621–1624 (2012).

41. Bolyen, E. et al. Reproducible, interactive, scalable and extensible microbiome data science using QIIME 2. Nat Biotechnol 37, 852–857 (2019).

42. Callahan, B. J. et al. DADA2: High-resolution sample inference from Illumina amplicon data. Nat Methods 13, 581–583 (2016).

43. DeSantis, T. Z. et al. Greengenes, a Chimera-Checked 16S rRNA Gene Database and Workbench Compatible with ARB. Appl Environ Microbiol 72, 5069–5072 (2006).

44. McDonald, D. et al. An improved Greengenes taxonomy with explicit ranks for ecological and evolutionary analyses of bacteria and archaea. ISME J 6, 610–618 (2012).

45. Quast, C. et al. The SILVA ribosomal RNA gene database project: improved data processing and web-based tools. Nucleic Acids Research 41, D590–D596 (2012).

46. Yilmaz, P. et al. The SILVA and “All-species Living Tree Project (LTP)” taxonomic frameworks. Nucl. Acids Res. 42, D643–D648 (2014).

47. McMurdie, P. J. & Holmes, S. Phyloseq: An R Package for Reproducible Interactive Analysis and Graphics of Microbiome Census Data. PLoS ONE 8, (2013).

48. Battaglia, T. W. "btools: A suite of R function for all types of microbial diversity analyses. (2018).

49. Love, M. I., Huber, W. & Anders, S. Moderated estimation of fold change and dispersion for RNA-seq data with DESeq2. Genome Biol 15, 550 (2014).

50. Kolde, R. & others. Pheatmap: pretty heatmaps. R package version 1, 726 (2012).

51. Lozupone, C. & Knight, R. UniFrac: a New Phylogenetic Method for Comparing Microbial Communities. Appl Environ Microbiol 71, 8228–8235 (2005).

52. Faith, D. P. Conservation evaluation and phylogenetic diversity. Biological Conservation 61, 1–10 (1992).

53. Engel, P. & Moran, N. A. The gut microbiota of insects – diversity in structure and function. FEMS Microbiol Rev 37, 699–735 (2013).

54. Moll, R. M., Romoser, W. S., Modrakowski, M. C., Moncayo, A. C. & Lerdthusnee, K. Meconial Peritrophic Membranes and the Fate of Midgut Bacteria During Mosquito (Diptera: Culicidae) Metamorphosis. J Med Entomol 38, 29–32 (2001).

55. Zheng, H., Steele, M. I., Leonard, S. P., Motta, E. V. S. & Moran, N. A. Honey bees as models for gut microbiota research. Lab Anim 47, 317–325 (2018).

56. Bağrıaçık, N. Extraction and Elemental Composition of Meconium in Polistes dominulus (Hymenoptera: Vespidae). Florida Entomologist 103, 206 (2020).

57. Martinson, V. G., Moy, J. & Moran, N. A. Establishment of Characteristic Gut Bacteria during Development of the Honeybee Worker. Appl Environ Microbiol 78, 2830–2840 (2012).

58. Powell, J. E., Martinson, V. G., Urban-Mead, K. & Moran, N. A. Routes of Acquisition of the Gut Microbiota of the Honey Bee Apis mellifera. Appl Environ Microbiol 80, 7378–7387 (2014).

59. Pais, I. S., Valente, R. S., Sporniak, M. & Teixeira, L. Drosophila melanogaster establishes a species-specific mutualistic interaction with stable gut-colonizing bacteria. PLoS Biol 16, e2005710 (2018).

60. Storelli, G. et al. Drosophila Perpetuates Nutritional Mutualism by Promoting the Fitness of Its Intestinal Symbiont Lactobacillus plantarum. Cell Metabolism 27, 362–377.e8 (2018).

61. Simpson, S. J. & Raubenheimer, D. The Nature of Nutrition: A Unifying Framework from Animal Adaptation to Human Obesity. (Princeton University Press, 2012). doi:10.1515/9781400842803.

62. Peristiwati, Natamihardja, Y. S. & Herlini, H. Isolation and identification of cellulolytic bacteria from termites gut (*Cryptotermes sp* .). J. Phys.: Conf. Ser. 1013, 012173 (2018).

63. Prem Anand, A. A. & Sripathi, K. Digestion of cellulose and xylan by symbiotic bacteria in the intestine of the Indian flying fox (Pteropus giganteus). Comparative Biochemistry and Physiology Part A: Molecular & Integrative Physiology 139, 65–69 (2004).

64. Anand, A. A. P. et al. Isolation and Characterization of Bacteria from the Gut of *Bombyx mori* that Degrade Cellulose, Xylan, Pectin and Starch and Their Impact on Digestion. Journal of Insect Science 10, 1–20 (2010).

65. Suenami, S., Konishi Nobu, M. & Miyazaki, R. Community analysis of gut microbiota in hornets, the largest eusocial wasps, Vespa mandarinia and V. simillima. Sci Rep 9, 9830 (2019).

66. Rothman, J. A., Loope, K. J., McFrederick, Q. S. & Wilson Rankin, E. E. Microbiome of the wasp Vespula pensylvanica in native and invasive populations, and associations with Moku virus. PLoS ONE 16, e0255463 (2021).

67. Reeson, A. F., Jankovic, T., Kasper, M. L., Rogers, S. & Austin, A. D. Application of 16S rDNA-DGGE to examine the microbial ecology associated with a social wasp Vespula germanica. Insect Mol Biol 12, 85–91 (2003).

68. Kucuk, R. A. Gut Bacteria in the Holometabola: A Review of Obligate and Facultative Symbionts. Journal of Insect Science 20, 22 (2020).

69. Abdul Rahman, N., et al. A molecular survey of Australian and North American termite genera indicates that vertical inheritance is the primary force shaping termite gut microbiomes. Microbiome 3, 5 (2015).

70. Nalepa, C. A. Origin of termite eusociality: trophallaxis integrates the social, nutritional, and microbial environments: Origin of termite eusociality. Ecol Entomol 40, 323–335 (2015).

71. Su, Q. et al. Strain-level analysis reveals the vertical microbial transmission during the life cycle of bumblebee. Microbiome 9, 216 (2021).

72. Shukla, S. P., Vogel, H., Heckel, D. G., Vilcinskas, A. & Kaltenpoth, M. Burying beetles regulate the microbiome of carcasses and use it to transmit a core microbiota to their offspring. Mol Ecol 27, 1980–1991 (2018).

73. Vogel, H. et al. The digestive and defensive basis of carcass utilization by the burying beetle and its microbiota. Nat Commun 8, 15186 (2017).

74. Zaidman-Rémy, A., Vigneron, A., Weiss, B. L. & Heddi, A. What can a weevil teach a fly, and reciprocally? Interaction of host immune systems with endosymbionts in Glossina and Sitophilus. BMC Microbiol 18, 150 (2018).

75. Salem, H. et al. Drastic Genome Reduction in an Herbivore’s Pectinolytic Symbiont. Cell 171, 1520–1531.e13 (2017).

76. 76. Lavy, O., et al. The maternal foam plug constitutes a reservoir for the desert locust’s bacterial symbionts. (2020).

77. Kwong, W. K. et al. Dynamic microbiome evolution in social bees. Science Advances 3, e1600513 (2017).

78. Robinson, G. E. Regulation of Division of Labor in Insect Societies. Annu. Rev. Entomol. 37, 637–665 (1992).

79. Lindsay, E. C., Metcalfe, N. B. & Llewellyn, M. S. The potential role of the gut microbiota in shaping host energetics and metabolic rate. J Anim Ecol. 89, 2415–2426 (2020).

80. Sudakaran, S., Kost, C. & Kaltenpoth, M. Symbiont Acquisition and Replacement as a Source of Ecological Innovation. Trends in Microbiology 25, 375–390 (2017).

81. Shapira, M. Gut Microbiotas and Host Evolution: Scaling Up Symbiosis. Trends in Ecology & Evolution 31, 539–549 (2016).

82. Lynch, J. B. & Hsiao, E. Y. Microbiomes as sources of emergent host phenotypes. Science 365, 1405– 1409 (2019).

83. Miller, E. T., Svanbäck, R. & Bohannan, B. J. M. Microbiomes as Metacommunities: Understanding Host-Associated Microbes through Metacommunity Ecology. Trends in Ecology & Evolution 33, 926– 935 (2018).

