## Supplementary material for "An experimental approach towards untangling the role of nature versus nurture in shaping the microbiome of social insects": Tables S1-S9; Figure S1

for

Tables S1-S9

Figure S1

**Table S1.** Details on samples used in this study. The column 'Sequenced' notates whether the sample was successfully sequenced (Y) or not (N). The column 'Included in analysis' notates whether the sample was used in the analysis (Y) or not (N).

| Sample ID | Treatment^ | Stage | Species | Larval instar | Box* | Days in experiment | Sequenced (Y/N) | Included in analysis (Y/N) |
|---|---|---|---|---|---|---|---|---|
| 58 | Control (G) | Mature worker | *V. germanica* | | CF1 | 15 | Y | Y |
| 59 | Control (G) | Mature worker | *V. germanica* | | CF1 | 15 | Y | N |
| 60 | Control (G) | Mature worker | *V. germanica* | | CF1 | 15 | Y | Y |
| 61 | Control (G) | Mature worker | *V. germanica* | | CF1 | 15 | Y | Y |
| 62 | Control (G) | Mature worker | *V. germanica* | | CF1 | 15 | Y | Y |
| 57 | Control (G) | Mature worker | *V. germanica* | | CF1 | 15 | Y | Y |
| 106 | Control (G) | Mature worker | *V. germanica* | | CF2 | 17 | Y | Y |
| 107 | Control (G) | Mature worker | *V. germanica* | | CF2 | 17 | Y | Y |
| 108 | Control (G) | Mature worker | *V. germanica* | | CF2 | 17 | Y | Y |
| 109 | Control (G) | Mature worker | *V. germanica* | | CF2 | 17 | Y | Y |
| 110 | Control (G) | Mature worker | *V. germanica* | | CF2 | 17 | Y | Y |
| 111 | Control (G) | Mature worker | *V. germanica* | | CF2 | 17 | Y | Y |
| 1 | Control (G) | Mature worker | *V. germanica* | | CF3 | 22 | Y | Y |
| 2 | Control (G) | Mature worker | *V. germanica* | | CF3 | 22 | Y | N |
| 3 | Control (G) | Mature worker | *V. germanica* | | CF3 | 22 | Y | Y |
| 64 | Control (G) | Mature worker | *V. germanica* | | CF3 | 13 | Y | Y |
| 65 | Control (G) | Mature worker | *V. germanica* | | CF3 | 13 | Y | Y |
| 143 | Control (G) | Mature worker | *V. germanica* | | CF3 | 7 | Y | Y |
| 5 | Control (G) | Mature worker | *V. germanica* | | CF4 | 22 | Y | Y |
| 6 | Control (G) | Mature worker | *V. germanica* | | CF4 | 22 | Y | Y |
| 7 | Control (G) | Mature worker | *V. germanica* | | CF4 | 22 | Y | Y |
| 8 | Control (G) | Mature worker | *V. germanica* | | CF4 | 22 | Y | Y |

| | | | | | | | | |
|---|---|---|---|---|---|---|---|---|
| 9 | Control (G) | Mature worker | *V. germanica* | | CF4 | 17 | Y | Y |
| 10 | Control (G) | Mature worker | *V. germanica* | | CF4 | 17 | Y | Y |
| 85 | Control (G) | Larva | *V. germanica* | 5th | CF1 | 6 | Y | Y |
| 86 | Control (G) | Larva | *V. germanica* | 5th | CF1 | 6 | Y | Y |
| 87 | Control (G) | Larva | *V. germanica* | 5th | CF1 | 6 | Y | Y |
| 56 | Control (G) | Larva | *V. germanica* | 5th | CF2 | 6 | Y | Y |
| 20 | Control (G) | Larva | *V. germanica* | 5th | CF3 | 6 | Y | Y |
| 68 | Control (G) | Larva | *V. germanica* | 4th-5th | CF4 | 11 | Y | Y |
| 69 | Control (G) | Larva | *V. germanica* | 4th-5th | CF4 | 11 | Y | Y |
| 70 | Control (G) | Larva | *V. germanica* | 4th-5th | CF4 | 4 | Y | Y |
| 71 | Control (G) | Larva | *V. germanica* | 4th-5th | CF4 | 4 | Y | Y |
| 72 | Control (G) | Larva | *V. germanica* | 4th-5th | CF4 | 4 | Y | Y |
| 12 | Control (G) | Larva | *V. germanica* | 4th | CF4 | 8 | Y | Y |
| 89 | Control (G) | Larva | *V. germanica* | 3rd-4th | CF4 | 7 | Y | Y |
| 101 | Control (O) | Mature worker | *V. orientalis* | | CF5 | 27 | Y | Y |
| 102 | Control (O) | Mature worker | *V. orientalis* | | CF5 | 27 | Y | Y |
| 103 | Control (O) | Mature worker | *V. orientalis* | | CF5 | 27 | Y | Y |
| 119 | Control (O) | Mature worker | *V. orientalis* | | CF5 | 17 | Y | Y |
| 48 | Control (O) | Mature worker | *V. orientalis* | | CF6 | 36 | Y | Y |
| 49 | Control (O) | Mature worker | *V. orientalis* | | CF6 | 36 | Y | Y |
| 50 | Control (O) | Mature worker | *V. orientalis* | | CF6 | 36 | Y | Y |
| 51 | Control (O) | Mature worker | *V. orientalis* | | CF6 | 36 | Y | Y |
| 32 | Control (O) | Mature worker | *V. orientalis* | | CF9 | 37 | Y | Y |
| 33 | Control (O) | Mature worker | *V. orientalis* | | CF9 | 37 | Y | Y |
| 34 | Control (O) | Mature worker | *V. orientalis* | | CF9 | 37 | Y | Y |
| 35 | Control (O) | Mature worker | *V. orientalis* | | CF9 | 37 | N | N |
| 40 | Control (O) | Mature worker | *V. orientalis* | | CF10 | 37 | Y | Y |
| 41 | Control (O) | Mature worker | *V. orientalis* | | CF10 | 37 | Y | Y |
| 42 | Control (O) | Mature worker | *V. orientalis* | | CF10 | 37 | Y | Y |
| 43 | Control (O) | Mature worker | *V. orientalis* | | CF10 | 37 | Y | Y |

| | | | | | | | | |
|---|---|---|---|---|---|---|---|---|
| 120 | Control (O) | Larva | *V. orientalis* | 5th | CF5 | 12 | Y | Y |
| 121 | Control (O) | Larva | *V. orientalis* | 5th | CF5 | 12 | N | N |
| 122 | Control (O) | Larva | *V. orientalis* | 5th | CF5 | 12 | Y | N |
| 123 | Control (O) | Larva | *V. orientalis* | 5th | CF5 | 12 | Y | N |
| 124 | Control (O) | Larva | *V. orientalis* | 5th | CF5 | 12 | Y | Y |
| 140 | Control (O) | Larva | *V. orientalis* | | CF5 | 8 | N | N |
| 142 | Control (O) | Larva | *V. orientalis* | 3rd-4th | CF5 | 8 | Y | N |
| 16 | Control (O) | Larva | *V. orientalis* | 2nd | CF6 | 32 | Y | Y |
| 26 | Control (O) | Larva | *V. orientalis* | 5th | CF6 | 17 | Y | Y |
| 129 | Control (O) | Larva | *V. orientalis* | 4th | CF6 | 12 | Y | Y |
| 130 | Control (O) | Larva | *V. orientalis* | 4th | CF6 | 12 | Y | Y |
| 135 | Control (O) | Larva | *V. orientalis* | 5th | CF6 | 10 | Y | Y |
| 136 | Control (O) | Larva | *V. orientalis* | 5th | CF6 | 10 | Y | Y |
| 137 | Control (O) | Larva | *V. orientalis* | 5th | CF6 | 10 | Y | Y |
| 138 | Control (O) | Larva | *V. orientalis* | 5th | CF6 | 10 | N | N |
| 139 | Control (O) | Larva | *V. orientalis* | 5th | CF6 | 10 | Y | Y |
| 29 | Control (O) | Larva | *V. orientalis* | 2nd | CF9 | 7 | Y | Y |
| 28 | Control (O) | Larva | *V. orientalis* | 1st-2nd | CF10 | 7 | N | N |
| 146 | Cross (G/O) | Mature worker | *V. germanica* | | CF7a | 15 | N | N |
| 147 | Cross (G/O) | Mature worker | *V. germanica* | | CF7a | 15 | N | N |
| 148 | Cross (G/O) | Mature worker | *V. germanica* | | CF7a | 15 | Y | N |
| 149 | Cross (G/O) | Mature worker | *V. germanica* | | CF7a | 15 | Y | N |
| 150 | Cross (G/O) | Mature worker | *V. germanica* | | CF7a | 15 | Y | N |
| 151 | Cross (G/O) | Mature worker | *V. germanica* | | CF7a | 15 | Y | N |
| 144 | Cross (G/O) | Mature worker | *V. germanica* | | CF7b | 12 | N | N |
| 145 | Cross (G/O) | Mature worker | *V. germanica* | | CF7b | 13 | Y | Y |
| 94 | Cross (G/O) | Mature worker | *V. germanica* | | CF7b | 19 | Y | Y |
| 95 | Cross (G/O) | Mature worker | *V. germanica* | | CF7b | 19 | Y | Y |
| 100 | Cross (G/O) | Mature worker | *V. germanica* | | CF7b | 14 | Y | Y |
| 105 | Cross (G/O) | Mature worker | *V. germanica* | | CF7b | 17 | Y | Y |

| | | | | | | | | |
|---|---|---|---|---|---|---|---|---|
| 152 | Cross (G/O) | Mature worker | *V. germanica* | | CF8a | 27 | Y | N |
| 153 | Cross (G/O) | Mature worker | *V. germanica* | | CF8a | 27 | Y | N |
| 154 | Cross (G/O) | Mature worker | *V. germanica* | | CF8a | 27 | Y | N |
| 155 | Cross (G/O) | Mature worker | *V. germanica* | | CF8a | 27 | Y | N |
| 156 | Cross (G/O) | Mature worker | *V. germanica* | | CF8a | 27 | Y | N |
| 157 | Cross (G/O) | Mature worker | *V. germanica* | | CF8a | 27 | Y | N |
| 96 | Cross (G/O) | Mature worker | *V. germanica* | | CF8b | 19 | Y | Y |
| 97 | Cross (G/O) | Mature worker | *V. germanica* | | CF8b | 19 | N | N |
| 98 | Cross (G/O) | Mature worker | *V. germanica* | | CF8b | 19 | Y | N |
| 99 | Cross (G/O) | Mature worker | *V. germanica* | | CF8b | 19 | Y | Y |
| 118 | Cross (G/O) | Mature worker | *V. germanica* | | CF8b | 8 | Y | Y |
| 73 | Cross (G/O) | Mature worker | *V. germanica* | | CF11 | 14 | Y | Y |
| 74 | Cross (G/O) | Mature worker | *V. germanica* | | CF11 | 14 | Y | Y |
| 75 | Cross (G/O) | Mature worker | *V. germanica* | | CF11 | 14 | Y | Y |
| 76 | Cross (G/O) | Mature worker | *V. germanica* | | CF11 | 14 | Y | Y |
| 77 | Cross (G/O) | Mature worker | *V. germanica* | | CF11 | 14 | Y | Y |
| 78 | Cross (G/O) | Mature worker | *V. germanica* | | CF11 | 14 | Y | Y |
| 36 | Cross (G/O) | Mature worker | *V. germanica* | | CF12 | 12 | Y | Y |
| 37 | Cross (G/O) | Mature worker | *V. germanica* | | CF12 | 12 | Y | Y |
| 38 | Cross (G/O) | Mature worker | *V. germanica* | | CF12 | 12 | Y | Y |
| 39 | Cross (G/O) | Mature worker | *V. germanica* | | CF12 | 12 | Y | Y |
| 112 | Cross (G/O) | Mature worker | *V. germanica* | | CF12 | 7 | Y | Y |
| 131 | Cross (G/O) | Mature worker | *V. germanica* | | CF18 | 23 | Y | Y |
| 132 | Cross (G/O) | Mature worker | *V. germanica* | | CF18 | 23 | Y | Y |
| 133 | Cross (G/O) | Mature worker | *V. germanica* | | CF18 | 23 | Y | Y |
| 134 | Cross (G/O) | Mature worker | *V. germanica* | | CF18 | 23 | Y | Y |
| 25 | Cross (G/O) | Larva | *V. orientalis* | 5th | CF7b | 9 | Y | Y |
| 79 | Cross (G/O) | Larva | *V. orientalis* | 4th | CF7b | 12 | N | N |
| 84 | Cross (G/O) | Larva | *V. orientalis* | 5th | CF8b | 19 | Y | Y |
| 17 | Cross (G/O) | Larva | *V. orientalis* | 4th | CF11 | 10 | Y | Y |

| | | | | | | | | |
|---|---|---|---|---|---|---|---|---|
| 18 | Cross (G/O) | Larva | *V. orientalis* | 4th | CF11 | 10 | Y | Y |
| 80 | Cross (G/O) | Larva | *V. orientalis* | 4th | CF11 | 6 | Y | Y |
| 81 | Cross (G/O) | Larva | *V. orientalis* | 4th | CF11 | 6 | Y | Y |
| 27 | Cross (G/O) | Larva | *V. orientalis* | 4th | CF12 | 10 | Y | Y |
| 30 | Cross (G/O) | Larva | *V. orientalis* | 5th | CF12 | 8 | Y | Y |
| 141 | Cross (G/O) | Larva | *V. orientalis* | 4th-5th | CF12 | 6 | N | N |
| 82 | Cross (G/O) | Larva | *V. orientalis* | 3rd-4th | CF18 | 3 | Y | N |
| 88 | Cross (G/O) | Larva | *V. orientalis* | 4th-5th | CF18 | 6 | Y | Y |
| 44 | Cross (O/G) | Mature worker | *V. orientalis* | | CF13 | 6 | N | N |
| 45 | Cross (O/G) | Mature worker | *V. orientalis* | | CF13 | 6 | Y | Y |
| 46 | Cross (O/G) | Mature worker | *V. orientalis* | | CF13 | 6 | Y | Y |
| 47 | Cross (O/G) | Mature worker | *V. orientalis* | | CF13 | 6 | N | N |
| 90 | Cross (O/G) | Mature worker | *V. orientalis* | | CF14 | 9 | Y | Y |
| 91 | Cross (O/G) | Mature worker | *V. orientalis* | | CF14 | 9 | Y | Y |
| 92 | Cross (O/G) | Mature worker | *V. orientalis* | | CF14 | 9 | Y | Y |
| 93 | Cross (O/G) | Mature worker | *V. orientalis* | | CF14 | 9 | Y | Y |
| 52 | Cross (O/G) | Mature worker | *V. orientalis* | | CF15 | 5 | Y | Y |
| 53 | Cross (O/G) | Mature worker | *V. orientalis* | | CF15 | 5 | Y | Y |
| 54 | Cross (O/G) | Mature worker | *V. orientalis* | | CF15 | 5 | Y | Y |
| 55 | Cross (O/G) | Mature worker | *V. orientalis* | | CF15 | 5 | Y | Y |
| 113 | Cross (O/G) | Mature worker | *V. orientalis* | | CF16 | 7 | Y | Y |
| 114 | Cross (O/G) | Mature worker | *V. orientalis* | | CF16 | 7 | Y | Y |
| 115 | Cross (O/G) | Mature worker | *V. orientalis* | | CF16 | 7 | Y | Y |
| 117 | Cross (O/G) | Mature worker | *V. orientalis* | | CF16 | 4 | Y | Y |
| 125 | Cross (O/G) | Mature worker | *V. orientalis* | | CF17 | 12 | Y | N |
| 126 | Cross (O/G) | Mature worker | *V. orientalis* | | CF17 | 12 | Y | N |
| 127 | Cross (O/G) | Mature worker | *V. orientalis* | | CF17 | 12 | Y | N |
| 128 | Cross (O/G) | Mature worker | *V. orientalis* | | CF17 | 12 | Y | N |
| 21 | Cross (O/G) | Larva | *V. germanica* | 3rd-4th | CF13 | 6 | N | N |
| 22 | Cross (O/G) | Larva | *V. germanica* | 3rd-4th | CF13 | 6 | Y | Y |

| | | | | | | | | |
|---|---|---|---|---|---|---|---|---|
| 23 | Cross (O/G) | Larva | *V. germanica* | 3rd-4th | CF13 | 6 | Y | Y |
| 24 | Cross (O/G) | Larva | *V. germanica* | 3rd-4th | CF13 | 6 | Y | Y |
| 63 | Cross (O/G) | Larva | *V. germanica* | 5th | CF16 | 3 | Y | N |

* *Boxes with identical names in which experiments were repeated are marked with a or b.*

^ *Treatments were as follows: Control (G) - V. germanica workers nursing V. germanica larvae; Control (O) - V. orientalis workers nursing V. orientalis larvae; Cross (G/O) - V. germanica workers nursing V. orientalis larvae; and Cross (O/G) - V. orientalis workers nursing V. germanica larvae.*

**Table S2.** Differentially abundant taxa between workers of both species from the four different treatments when compared to German wasp workers from the control treatment. Adjusted P < 0.05, Wald test followed by Benjamini–Hochberg correction for multiple testing.

| Taxon | Phylum | Class | Order | Family | Taxa level | Base Mean | log2 Fold Change | lfcSE | stat | *P* | padj |
|---|---|---|---|---|---|---|---|---|---|---|---|
| Chitinophagaceae | Bacteroidota | Chitinophagia | Chitinophagales | Chitinophagaceae | Family | 3.44 | 1.73 | 0.56 | 3.10 | 0.002 | 0.012 |
| Enterobacteriaceae | Pseudomonadota | Gammaproteobacteria | Enterobacterales | Enterobacteriaceae | Family | 223.59 | -3.41 | 0.90 | -3.80 | 0.000 | 0.001 |
| Yersiniaceae | Pseudomonadota | Gammaproteobacteria | Enterobacterales | Yersiniaceae | Family | 99.61 | -2.62 | 0.88 | -2.98 | 0.003 | 0.014 |
| Acetobacter | Pseudomonadota | Alphaproteobacteria | Rhodospirillales | Acetobacteraceae | Genus | 6.38 | 3.35 | 0.61 | 5.49 | 0.000 | 0.000 |
| Acinetobacter | Pseudomonadota | Gammaproteobacteria | Pseudomonadales | Moraxellaceae | Genus | 3.33 | 1.65 | 0.51 | 3.25 | 0.001 | 0.008 |
| Brochothrix | Bacillota | Bacilli | Bacillales | Listeriaceae | Genus | 2.66 | -1.96 | 0.64 | -3.06 | 0.002 | 0.012 |
| Carnobacterium | Bacillota | Bacilli | Lactobacillales | Carnobacteriaceae | Genus | 1.94 | -2.02 | 0.55 | -3.66 | 0.000 | 0.002 |
| Corynebacterium | Actinomycetota | Actinomycetia | Mycobacteriales | Corynebacteriaceae | Genus | 1.98 | 2.02 | 0.48 | 4.20 | 0.000 | 0.000 |
| Escherichia-Shigella | Gammaproteobacteria | Enterobacterales | Enterobacteriaceae | | Genus | 46.77 | -2.27 | 0.87 | -2.61 | 0.009 | 0.043 |
| Fructobacillus | Bacillota | Bacilli | Lactobacillales | Lactobacillaceae | Genus | 17.77 | -3.56 | 0.86 | -4.12 | 0.000 | 0.000 |
| Hafnia-Obesumbacterium | Pseudomonadota | Gammaproteobacteria | Enterobacterales | Hafniaceae | Genus | 38.20 | -2.23 | 0.88 | -2.55 | 0.011 | 0.049 |
| Lactobacillus | Bacillota | Bacilli | Lactobacillales | Lactobacillaceae | Genus | 4.27 | 3.45 | 0.63 | 5.52 | 0.000 | 0.000 |
| Leuconostoc | Bacillota | Bacilli | Lactobacillales | Lactobacillaceae | Genus | 9.46 | -4.48 | 0.75 | -6.00 | 0.000 | 0.000 |
| Morganella | Pseudomonadota | Gammaproteobacteria | Enterobacterales | Morganellaceae | Genus | 37.11 | -5.80 | 0.76 | -7.62 | 0.000 | 0.000 |
| Providencia | Pseudomonadota | Gammaproteobacteria | Enterobacterales | Morganellaceae | Genus | 19.86 | -4.30 | 0.79 | -5.41 | 0.000 | 0.000 |
| Staphylococcus | Bacillota | Bacilli | Bacillales | Staphylococcaceae | Genus | 9.42 | 3.73 | 0.66 | 5.70 | 0.000 | 0.000 |
| Enterobacterales | Pseudomonadota | Gammaproteobacteria | Enterobacterales | | Order | 893.91 | -2.55 | 0.79 | -3.21 | 0.001 | 0.009 |
| Lactobacillales | Bacillota | Bacilli | Lactobacillales | | Order | 54.29 | -3.49 | 0.85 | -4.11 | 0.000 | 0.000 |
| Weissella viridescens | Bacillota | Bacilli | Lactobacillales | Lactobacillaceae | Species | 3.72 | 1.80 | 0.58 | 3.09 | 0.002 | 0.012 |

**Table S3.** Microbiome composition of *V. germanica* and *V. orientalis* in nursing workers per treatment. Only taxa with mean relative abundance of 1% and above in at least one group is shown. Prevalence in the group is shown in parenthesis. Letters before the taxonomic name represent the taxonomic level at which that organism was identified (f – family, o – order, g – genus, s – species).

| | | | Mean relative abundance [%] (Prevalence [%]) | | | |
|---|---|---|---|---|---|---|
| | | | *V. germanica* | | *V. orientalis* | |
| Phylum | Order | Taxon | Cross | Control | Cross | Control |
| Actinomycetota | Bifidobacteriales | *Bifidobacterium (g)* | 2.0 (48) | 0.8 (36) | 2.9 (57) | 0.1 (33) |
| Firmicutes | Bacillales | *Bacillus (g)* | 0.1 (9) | 3.6 (5) | 1.5 (14) | 0.1 (7) |
| Firmicutes | Bacillales | *Kurthia (g)* | 5.2 (4) | 0.2 (5) | 0.0 (21) | 0.1 (20) |
| Firmicutes | Lactobacillales | *Carnobacterium (g)* | | 0.4 (14) | 2.8 (7) | |
| Firmicutes | Lactobacillales | *Enterococcus (g)* | 7.4 (87) | 0.5 (50) | 1.3 (86) | 0.2 (47) |
| Firmicutes | Lactobacillales | *Fructobacillus (g)* | 1.7 (43) | 0.6 (36) | 9.0 (36) | 0.0 (13) |
| Firmicutes | Lactobacillales | *Lactobacillales (o)* | 0.3 (17) | 5.3 (45) | 1.8 (43) | 0.3 (53) |
| Firmicutes | Lactobacillales | *Lactobacillus (g)* | 1.1 (22) | 0.0 (9) | 0.1 (14) | 0.3 (53) |
| Firmicutes | Lactobacillales | *Lactococcus (g)* | 25.2 (100) | 32.2 (100) | 29.1 (100) | 48.7 (93) |
| Firmicutes | Lactobacillales | *Leuconostoc (g)* | 2.7 (17) | 0.6 (68) | 0.1 (21) | |
| Firmicutes | Lactobacillales | *Leuconostocaceae (f)* | 0.2 (52) | 0.1 (32) | 1.4 (93) | 0.0 (33) |
| Firmicutes | Lactobacillales | *Weissella viridescens(s)* | 0.4 (9) | 0.3 (27) | 8.8 (21) | 2.1 (33) |
| Firmicutes | Vellionellales | *Veillonella (g)* | 2.4 (43) | 1.0 (36) | 2.9 (64) | 0.2 (40) |
| Pseudomonadota | Enterobacterales | *Enterobacterales (o)* | 16.6 (96) | 18.9 (95) | 20.4 (100) | 2.9 (93) |
| Pseudomonadota | Enterobacterales | *Enterobacteriaceae (f)* | 10.8 (70) | 4.5 (82) | 1.2 (64) | 2.0 (73) |
| Pseudomonadota | Enterobacterales | *Escherichia-Shigella (g)* | 2.0 (43) | 1.3 (36) | 2.6 (64) | 0.2 (33) |
| Pseudomonadota | Enterobacterales | *Hafnia-Obesumbacterium (g)* | 0.9 (43) | 2.2 (59) | 1.3 (36) | 0.9 (27) |
| Pseudomonadota | Enterobacterales | *Morganella (g)* | 0.2 (26) | 4.5 (45) | 0.1 (21) | 0.3 (40) |
| Pseudomonadota | Enterobacterales | *Proteus (g)* | 7.1 (96) | 7.8 (100) | 2.1 (100) | 7.5 (87) |
| Pseudomonadota | Enterobacterales | *Providencia (g)* | 3.9 (26) | 4.3 (64) | 0.0 (7) | 1.1 (27) |
| Pseudomonadota | Enterobacterales | *Serratia symbiotica (s)* | 0.0 (4) | | | 18.3 (20) |
| Pseudomonadota | Enterobacterales | *Yersiniaceae (f)* | 1.3 (48) | 5.1 (64) | 3.4 (64) | 0.8 (60) |
| Pseudomonadota | Pseudomonadales | *Acinetobacter (g)* | | 0.4 (36) | 0.1 (43) | 1.6 (60) |
| Pseudomonadota | Pseudomonadales | *Pseudomonas (g)* | 0.1 (30) | 0.6 (36) | 0.1 (21) | 2.4 (47) |

| | | | | | | |
|---|---|---|---|---|---|---|
| Pseudomonadota | Rhodospirillales | *Acetobacter (g)* | 0.1 (35) | 0.1 (36) | 0.1 (57) | 2.9 (47) |

**Table S4.** Weighted UniFrac Beta diversity between larvae from different treatments. P values are listed in the lower diagonal, F values and summarized $R2$ are listed in the upper diagonal.

| | *V. germanica* (Control) | *V. germanica* (Cross) | *V. orientalis* (Control) | *V. orientalis* (Cross) |
|---|---|---|---|---|
| *V. germanica* (Control) | | F=5.26, R2=0.29 | F=1.03, R2=0.05 | F=3.07, R2=0.14 |
| *V. germanica* (Cross) | 0.025 | | F=5.31, R2=0.31 | F=5.02, R2=0.33 |
| *V. orientalis* (Control) | 0.309 | 0.039 | | F=2.87, R2=0.14 |
| *V. orientalis* (Cross) | 0.044 | 0.043 | 0.058 | |

**Table S5.** Differentially abundant taxa between larvae of both species from the four different treatments when compared to German wasp larvae from the control treatment. Adjusted P < 0.05, Wald test followed by Benjamini–Hochberg correction for multiple testing.

| Taxon | Phylum | Class | Order | Family | Taxa level | base Mean | log2 Fold Change | lfcSE | stat | *P* | padj |
|---|---|---|---|---|---|---|---|---|---|---|---|
| Enterobacteriaceae | Pseudomonadota | Gammaproteobacteria | Enterobacterales | Enterobacteriaceae | Family | 4.70 | -2.50 | 0.72 | -3.48 | 0.001 | 0.002 |
| Micrococcaceae | Actinomycetota | Actinomycetia | Micrococcales | Micrococcaceae | Family | 2.87 | 2.64 | 0.68 | 3.87 | 0.000 | 0.001 |
| Planococcaceae | Firmicutes | Bacilli | Caryophanales | Planococcaceae | Family | 18.14 | -5.45 | 0.94 | -5.81 | 0.000 | 0.000 |
| Xanthobacteraceae | Pseudomonadota | Alphaproteobacteria | Hyphomicrobiales | Xanthobacteraceae | Family | 1.67 | 1.47 | 0.58 | 2.54 | 0.011 | 0.031 |
| Bombiscardovia | Actinomycetota | Actinomycetia | Bifidobacteriales | Bifidobacteriaceae | Genus | 11.36 | -2.37 | 0.84 | -2.83 | 0.005 | 0.016 |
| Brochothrix | Firmicutes | Bacilli | Bacillales | Listeriaceae | Genus | 43.74 | -5.09 | 1.04 | -4.92 | 0.000 | 0.000 |
| Carnobacterium | Firmicutes | Bacilli | Lactobacillales | Carnobacteriaceae | Genus | 60.46 | 4.81 | 0.86 | 5.57 | 0.000 | 0.000 |
| Corynebacterium | Actinomycetota | Actinomycetia | Mycobacteriales | Corynebacteriaceae | Genus | 8.88 | 2.98 | 0.80 | 3.72 | 0.000 | 0.001 |
| Enterococcus | Firmicutes | Bacilli | Lactobacillales | Enterococcaceae | Genus | 18.37 | -2.27 | 0.85 | -2.67 | 0.008 | 0.023 |
| Erysipelatoclostridium | Firmicutes | Erysipelotrichia | Erysipelotrichales | Erysipelotrichaceae | Genus | 1.90 | -1.83 | 0.62 | -2.93 | 0.003 | 0.014 |
| Escherichia-Shigella | Pseudomonadota | Gammaproteobacteria | Enterobacterales | Enterobacteriaceae | Genus | 30.27 | -5.40 | 0.97 | -5.56 | 0.000 | 0.000 |
| Fructobacillus | Firmicutes | Bacilli | Lactobacillales | Lactobacillaceae | Genus | 132.10 | -7.96 | 0.93 | -8.58 | 0.000 | 0.000 |
| Kurthia | Firmicutes | Bacilli | Bacillales | Planococcaceae | Genus | 110.71 | -2.90 | 1.03 | -2.82 | 0.005 | 0.016 |
| Lactobacillus | Firmicutes | Bacilli | Lactobacillales | Lactobacillaceae | Genus | 4.16 | -2.91 | 0.74 | -3.95 | 0.000 | 0.001 |
| Lactococcus | Firmicutes | Bacilli | Lactobacillales | Streptococcaceae | Genus | 69.98 | -2.19 | 0.76 | -2.88 | 0.004 | 0.016 |
| Leuconostoc | Firmicutes | Bacilli | Lactobacillales | Lactobacillaceae | Genus | 15.77 | -3.68 | 0.69 | -5.32 | 0.000 | 0.000 |
| Macrococcus | Firmicutes | Bacilli | Bacillales | Staphylococcaceae | Genus | 44.59 | 2.84 | 0.89 | 3.18 | 0.001 | 0.007 |
| Morganella | Pseudomonadot | Gammaproteobacteria | Enterobacterales | Morganellaceae | Genus | 80.70 | -2.87 | 1.01 | -2.85 | 0.00 | 0.01 |

| | | | | | | | | | | | |
|---|---|---|---|---|---|---|---|---|---|---|---|
| | a | | | | | | | | | 4 | 6 |
| Proteus | Pseudomonadota | Gammaproteobacteria | Enterobacterales | Enterobacteriaceae | Genus | 432.81 | -3.09 | 1.19 | -2.59 | 0.010 | 0.028 |
| Staphylococcus | Firmicutes | Bacilli | Bacillales | Staphylococcaceae | Genus | 472.15 | -3.42 | 0.92 | -3.71 | 0.000 | 0.001 |
| Veillonella | Firmicutes | Negativicutes | Vellionellales | Veillonellaceae | Genus | 3.70 | -2.04 | 0.73 | -2.79 | 0.005 | 0.017 |
| Weissella | Firmicutes | Bacilli | Lactobacillales | Lactobacillaceae | Genus | 9.14 | -2.09 | 0.85 | -2.47 | 0.014 | 0.036 |
| Xanthobacter | Pseudomonadota | Alphaproteobacteria | Hyphomicrobiales | Xanthobacteraceae | Genus | 1.76 | 1.75 | 0.62 | 2.85 | 0.004 | 0.016 |
| Lactobacillales | Firmicutes | Bacilli | Lactobacillales | | Order | 667.84 | 5.85 | 0.97 | 6.05 | 0.000 | 0.000 |
| Enterococcus cecorum | Firmicutes | Bacilli | Lactobacillales | Enterococcaceae | Species | 3.39 | 2.72 | 0.70 | 3.92 | 0.000 | 0.001 |
| Lactobacillus vespulae | Firmicutes | Bacilli | Lactobacillales | Lactobacillaceae | Species | 6.15 | -3.98 | 0.72 | -5.51 | 0.000 | 0.000 |
| Weissella ceti | Firmicutes | Bacilli | Lactobacillales | Lactobacillaceae | Species | 28.50 | 6.02 | 0.78 | 7.74 | 0.000 | 0.000 |

**Table S6.** Spearman's rank correlation coefficient between larvae from different treatments. Average *r* values are listed in the upper diagonal, and P values are listed in the upper diagonal.

| | *V. germanica* (Control) | *V. germanica* (Cross) | *V. orientalis* (Control) | *V. orientalis* (Cross) |
|---|---|---|---|---|
| *V. germanica* (Control) | | 0.03 | 0.57 | 0.74 |
| *V. germanica* (Cross) | 0.46 | | 0.56 | 0.76 |
| *V. orientalis* (Control) | 0.22 | 0.23 | | 0.01 |
| *V. orientalis* (Cross) | 0.13 | 0.12 | 0.54 | |

**Table S7.** Microbiome composition of *V. germanica* and *V. orientalis* in larvae per treatment. Only taxa with mean relative abundance of 1% and above in at least one group is shown. Prevalence in the group is shown in parenthesis. Letters before the taxonomic name represent the taxonomic level at which that organism was identified (f – family, o – order, g – genus, s – species).

| | | | Mean relative abundance [%] (Prevalence [%]) | | | |
|---|---|---|---|---|---|---|
| | | | *V. germanica* | | *V. orientalis* | |
| Phylum | Order | Taxon | Cross | Control | Cross | Control |
| Actinomycetota | Bifidobacteriales | *Bifidobacterium commune (s)* | 0.0 (33) | 0.9 (25) | 15.9 (89) | 0.2 (64) |
| Firmicutes | Bacillales | *Brochothrix (g)* | 0.1 (33) | 2.6 (75) | 5.1 (11) | 0.2 (18) |
| Firmicutes | Bacillales | *Kurthia (g)* | 1.5 (67) | 6.4 (75) | 1.1 (78) | 0.7 (91) |
| Firmicutes | Bacillales | *Macrococcus (g)* | 0.0 (33) | 0.4 (75) | 0.5 (44) | 3.0 (73) |
| Firmicutes | Bacillales | *Staphylococcus (g)* | 0.6 (67) | 18.9 (100) | 8.3 (89) | 1.7 (100) |
| Firmicutes | Caryophanales | *Planococcaceae (f)* | 0.6 (33) | 1.5 (58) | 0.4 (22) | |
| Firmicutes | Lactobacillales | *Carnobacterium (g)* | | 0.5 (17) | 1.4 (67) | 3.2 (82) |
| Firmicutes | Lactobacillales | *Fructobacillus (g)* | 0.7 (67) | 8.8 (83) | 0.6 (33) | 0.1 (9) |
| Firmicutes | Lactobacillales | *Lactobacillales (o)* | | 1.0 (50) | 14.1 (78) | 32.5 (91) |
| Firmicutes | Lactobacillales | *Lactobacillus (g)* | 0.3 (33) | 1.3 (50) | 0.8 (11) | 0.3 (36) |
| Firmicutes | Lactobacillales | *Lactococcus (g)* | 0.1 (100) | 3.2 (100) | 0.2 (100) | 0.7 (100) |
| Firmicutes | Lactobacillales | *Leuconostoc (g)* | 0.0 (33) | 5.9 (100) | 1.0 (11) | 0.1 (36) |
| Firmicutes | Lactobacillales | *Leuconostocaceae (f)* | 0.1 (100) | 2.4 (100) | 7.4 (89) | 12.9 (91) |
| Firmicutes | Lactobacillales | *Weissella (g)* | | 0.1 (67) | 5.8 (56) | 0.3 (18) |
| Firmicutes | Lactobacillales | *Weissella ceti (s)* | 0.1 (33) | 0.1 (8) | 0.0 (33) | 2.1 (82) |
| Firmicutes | Vellionellales | *Veillonella (g)* | | 1.0 (67) | 0.2 (22) | 0.1 (9) |
| Pseudomonadota | Enterobacterales | *Enterobacterales (o)* | 34.9 (100) | 26.3 (100) | 6.1 (100) | 32.2 (91) |
| Pseudomonadota | Enterobacterales | *Enterobacteriaceae (f)* | | 0.2 (50) | 2.5 (56) | 3.2 (18) |
| Pseudomonadota | Enterobacterales | *Escherichia-Shigella (g)* | | 2.1 (67) | 1.1 (33) | 0.1 (18) |
| Pseudomonadota | Enterobacterales | *Morganella (g)* | 26.3 (67) | 0.5 (25) | 0.4 (33) | |
| Pseudomonadota | Enterobacterales | *Proteus (g)* | 32.2 (100) | 8.4 (92) | 10.0 (89) | 1.2 (73) |
| Pseudomonadota | Enterobacterales | *Providencia (g)* | 1.7 (67) | 0.0 (8) | 0.1 (22) | 0.1 (9) |
| Pseudomonadota | Enterobacterales | *Yersiniaceae (f)* | 0.4 (33) | 0.1 (8) | 6.5 (56) | 0.2 (9) |

**Table S8.** Differentially abundant taxa between larvae and workers of the oriental hornet (*V. orientalis*) from the control treatment. Adjusted P < 0.05, Wald test followed by Benjamini–Hochberg correction for multiple testing.

| Taxon | Phylum | Class | Order | Family | Taxa level | baseMean | log2FoldChange | lfcSE | stat | pvalue | padj |
|---|---|---|---|---|---|---|---|---|---|---|---|
| Leuconostocaceae | Firmicutes | Bacilli | Lactobacillales | Leuconostocaceae | Family | 122.59 | -8.83 | 0.88 | -10.02 | 0.000 | 0.000 |
| Micrococcaceae | Actinomycetota | Actinomycetia | Micrococcales | Micrococcaceae | Family | 1.66 | -2.64 | 0.52 | -5.04 | 0.000 | 0.000 |
| Yersiniaceae | Pseudomonadota | Gammaproteobacteria | Enterobacterales | Yersiniaceae | Family | 87.64 | 4.19 | 1.04 | 4.05 | 0.000 | 0.001 |
| Carnobacterium | Firmicutes | Bacilli | Lactobacillales | Carnobacteriaceae | Genus | 22.61 | -7.09 | 0.82 | -8.66 | 0.000 | 0.000 |
| Macrococcus | Firmicutes | Bacilli | Bacillales | Staphylococcaceae | Genus | 15.54 | -6.80 | 0.73 | -9.29 | 0.000 | 0.000 |
| Kurthia | Firmicutes | Bacilli | Bacillales | Planococcaceae | Genus | 40.19 | -3.79 | 0.89 | -4.25 | 0.000 | 0.000 |
| Kocuria | Actinomycetota | Actinomycetia | Micrococcales | Micrococcaceae | Genus | 2.36 | -2.31 | 0.57 | -4.07 | 0.000 | 0.001 |
| Morganella | Pseudomonadota | Gammaproteobacteria | Enterobacterales | Morganellaceae | Genus | 52.27 | 2.93 | 0.99 | 2.96 | 0.003 | 0.023 |
| Acinetobacter | Pseudomonadota | Gammaproteobacteria | Pseudomonadales | Moraxellaceae | Genus | 2.91 | 3.00 | 0.69 | 4.37 | 0.000 | 0.000 |
| Proteus | Pseudomonadota | Gammaproteobacteria | Enterobacterales | Enterobacteriaceae | Genus | 398.33 | 3.19 | 1.05 | 3.04 | 0.002 | 0.019 |
| Acetobacter | Pseudomonadota | Alphaproteobacteria | Rhodospirillales | Acetobacteraceae | Genus | 5.46 | 3.32 | 0.72 | 4.63 | 0.000 | 0.000 |
| Providencia | Pseudomonadota | Gammaproteobacteria | Enterobacterales | Morganellaceae | Genus | 50.60 | 3.87 | 1.04 | 3.73 | 0.000 | 0.002 |
| Hafnia-Obesumbacterium | Pseudomonadota | Gammaproteobacteria | Enterobacterales | Hafniaceae | Genus | 26.64 | 3.94 | 1.01 | 3.89 | 0.000 | 0.001 |
| Lactococcus | Firmicutes | Bacilli | Lactobacillales | Streptococcaceae | Genus | 1382.70 | 6.40 | 0.67 | 9.58 | 0.000 | 0.000 |
| Lactobacillales | Firmicutes | Bacilli | Lactobacillales |  | Order | 254.85 | -6.97 | 0.98 | -7.13 | 0.000 | 0.000 |
| Enterobacterales | Pseudomonadota | Gammaproteobacteria | Enterobacterales |  | Order | 1000.51 | -3.12 | 0.94 | -3.31 | 0.001 | 0.008 |
| Weissella ceti | Firmicutes | Bacilli | Lactobacillales | Lactobacillaceae | Species | 10.02 | -6.43 | 0.59 | -10.99 | 0.000 | 0.000 |
| Enterococcus cecorum | Firmicutes | Bacilli | Lactobacillales | Enterococcaceae | Species | 1.81 | -2.72 | 0.53 | -5.19 | 0.000 | 0.000 |
| Bifidobacterium commune | Actinomycetota | Actinomycetia | Bifidobacteriales | Bifidobacteriaceae | Species | 76.31 | -2.24 | 0.73 | -3.08 | 0.002 | 0.017 |
| Weissella viridescens | Firmicutes | Bacilli | Lactobacillales | Lactobacillaceae | Species | 3.18 | 3.40 | 0.75 | 4.52 | 0.000 | 0.000 |

**Table S9.** Differentially abundant taxa between larvae and workers of the German wasp (*V. germanica*) from the control treatment. Adjusted P < 0.05, Wald test followed by Benjamini–Hochberg correction for multiple testing.

| Taxon | Phylum | Class | Order | Family | Taxa level | baseMean | log2FoldChange | lfcSE | stat | pvalue | padj |
|---|---|---|---|---|---|---|---|---|---|---|---|
| Leuconostocaceae | Firmicutes | Bacilli | Lactobacillales | Leuconostocaceae | Family | 122.59 | -6.14 | 0.78 | -7.88 | 0.000 | 0.000 |
| Planococcaceae | Firmicutes | Bacilli | Caryophanales | Planococcaceae | Family | 6.83 | -5.46 | 0.60 | -9.05 | 0.000 | 0.000 |
| Enterobacteriaceae | Pseudomonadota | Gammaproteobacteria | Enterobacterales | Enterobacteriaceae | Family | 175.47 | 4.97 | 0.99 | 5.03 | 0.000 | 0.000 |
| Yersiniaceae | Pseudomonadota | Gammaproteobacteria | Enterobacterales | Yersiniaceae | Family | 87.64 | 7.20 | 0.96 | 7.51 | 0.000 | 0.000 |
| Staphylococcus | Firmicutes | Bacilli | Bacillales | Staphylococcaceae | Genus | 168.73 | -9.69 | 0.82 | -11.89 | 0.000 | 0.000 |
| Kurthia | Firmicutes | Bacilli | Bacillales | Planococcaceae | Genus | 40.19 | -7.46 | 0.82 | -9.14 | 0.000 | 0.000 |
| Lactobacillus | Firmicutes | Bacilli | Lactobacillales | Lactobacillaceae | Genus | 7.79 | -4.94 | 0.74 | -6.70 | 0.000 | 0.000 |
| Fructobacillus | Firmicutes | Bacilli | Lactobacillales | Lactobacillaceae | Genus | 81.36 | -4.86 | 0.87 | -5.57 | 0.000 | 0.000 |
| Macrococcus | Firmicutes | Bacilli | Bacillales | Staphylococcaceae | Genus | 15.54 | -3.96 | 0.65 | -6.05 | 0.000 | 0.000 |
| Leuconostoc | Firmicutes | Bacilli | Lactobacillales | Lactobacillaceae | Genus | 45.98 | -3.73 | 0.80 | -4.65 | 0.000 | 0.000 |
| Brochothrix | Firmicutes | Bacilli | Bacillales | Listeriaceae | Genus | 18.18 | -3.30 | 0.80 | -4.12 | 0.000 | 0.000 |
| Kocuria | Actinomycetota | Actinomycetia | Micrococcales | Micrococcaceae | Genus | 2.36 | -2.99 | 0.49 | -6.15 | 0.000 | 0.000 |
| Bombiscardovia | Actinomycetota | Actinomycetia | Bifidobacteriales | Bifidobacteriaceae | Genus | 4.45 | -2.90 | 0.55 | -5.23 | 0.000 | 0.000 |
| Weissella | Firmicutes | Bacilli | Lactobacillales | Lactobacillaceae | Genus | 4.06 | -2.21 | 0.60 | -3.67 | 0.000 | 0.001 |
| Photobacterium | Pseudomonadota | Gammaproteobacteria | Vibrionales | Vibrionaceae | Genus | 1.32 | -1.83 | 0.47 | -3.91 | 0.000 | 0.000 |
| Vibrio | Pseudomonadota | Gammaproteobacteria | Vibrionales | Vibrionaceae | Genus | 4.77 | -1.69 | 0.66 | -2.58 | 0.010 | 0.033 |
| Erysipelatoclostridium | Firmicutes | Bacilli | Erysipelotrichales | Coprobacillaceae | Genus | 2.23 | -1.38 | 0.52 | -2.66 | 0.008 | 0.028 |
| Bacillus | Firmicutes | Bacilli | Bacillales | Bacillaceae | Genus | 1.70 | -1.33 | 0.51 | -2.62 | 0.009 | 0.030 |
| Pseudomonas | Pseudomonadota | Gammaproteobacteria | Pseudomonadales | Pseudomonadaceae | Genus | 3.52 | 2.02 | 0.63 | 3.21 | 0.001 | 0.005 |
| Lactococcus | Firmicutes | Bacilli | Lactobacillales | Streptococcaceae | Genus | 1382.70 | 3.52 | 0.60 | 5.85 | 0.000 | 0.000 |
| Morganella | Pseudomonadota | Gammaproteobacteria | Enterobacterales | Morganellaceae | Genus | 52.27 | 3.97 | 0.81 | 4.90 | 0.000 | 0.000 |
| Hafnia-Obesumbacterium | Pseudomonadota | Gammaproteobacteria | Enterobacterales | Hafniaceae | Genus | 26.64 | 6.18 | 0.92 | 6.69 | 0.000 | 0.000 |
| Providencia | Pseudomonadota | Gammaproteobacteria | Enterobacterales | Morganellaceae | Genus | 50.60 | 7.27 | 0.97 | 7.53 | 0.000 | 0.000 |
| Lactobacillales | Firmicutes | Bacilli | Lactobacillales | | Order | 254.85 | 2.36 | 0.88 | 2.67 | 0.008 | 0.028 |
| Bifidobacterium commune | Actinomycetota | Actinomycetia | Bifidobacteriales | Bifidobacteriaceae | Species | 76.31 | -2.97 | 0.61 | -4.85 | 0.000 | 0.000 |
| Lactobacillus vespulae | Firmicutes | Bacilli | Lactobacillales | Lactobacillaceae | Species | 3.94 | -2.53 | 0.56 | -4.52 | 0.000 | 0.000 |

**Figure S1.** Weighted UniFrac Beta diversity among workers of *V. orientalis* and *V. germanica* collected after participating for a various number of days in the experiment (treatments).

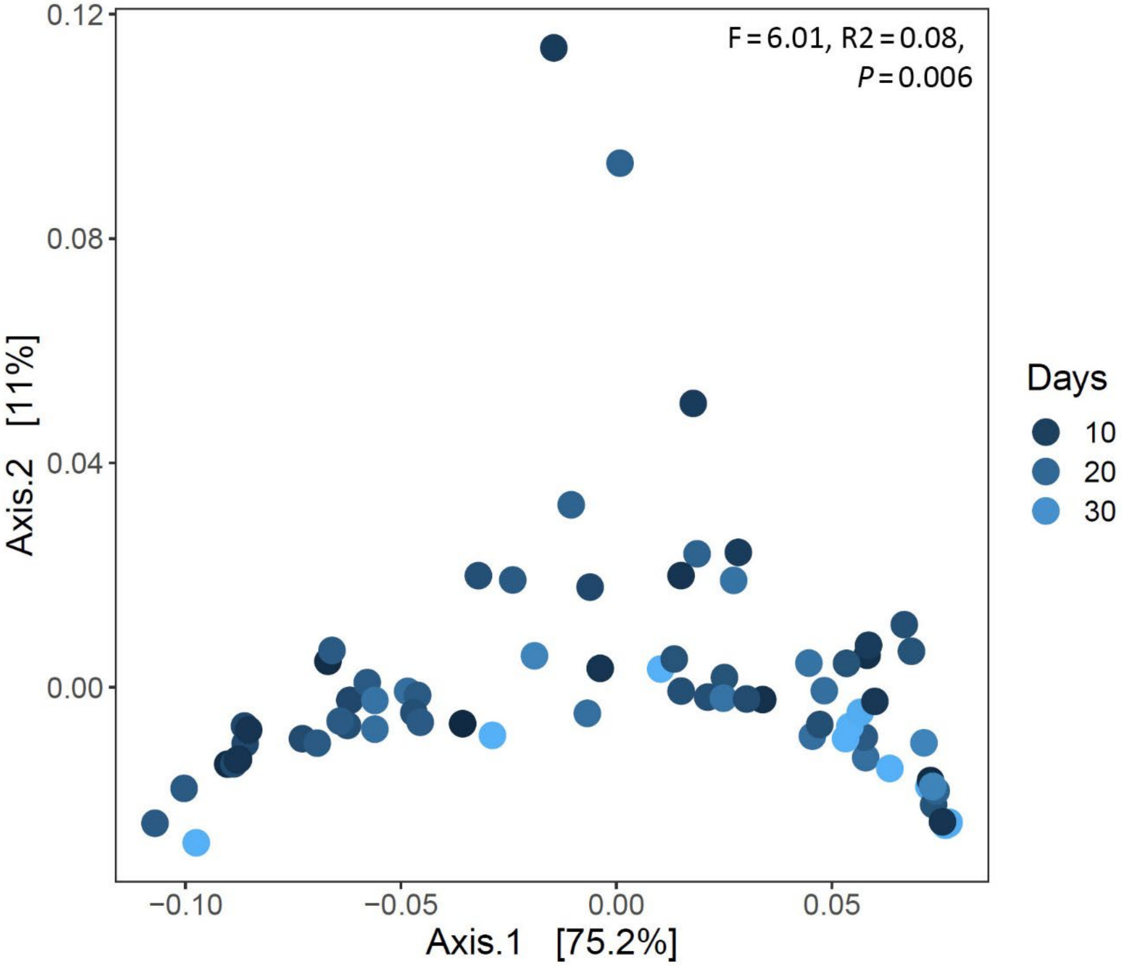
