## Supplementary material for "An experimental approach towards untangling the role of nature versus nurture in shaping the microbiome of social insects": R code: 16S_wasp.Rmd

````
---
title: "16S Binge"
author: "Efrat Sharon"
date:  "`r format(Sys.time(), 'Last updated: %d/%m/%y')`"
output: html_document
---
## load
```{r setup, include=FALSE}
knitr::opts_chunk$set(echo = F)

#BiocManager::install("metagenomeSeq")
#BiocManager::install("phyloseq")
#BiocManager::install("DESeq2")
#BiocManager::install("ram")
#BiocManager::install('EnhancedVolcano')
#BiocManager::install('microbiome')
#remotes::install_github("twbattaglia/btools")

library(readr)
library(plyr)
library(dplyr)
library(tidyr)
library(forcats) #fct_reorder

library(taxa)
#library(metagenomeSeq)
library(readxl)
#library(car)
library(tibble)
library(stringr)
library(ggplot2)
library(ggthemes)
library(ggrepel)
library(forcats)
library(ggridges) #geom_density_ridges
library(ggVennDiagram)
library(table1)
#library(caret)
#library( DataExplorer)

library(phyloseq)
library(vegan)
library(rstatix) #kruskal_test
library(lmerTest) #lmer + anova
library(emmeans) # post hoc of lmer
library(pairwiseAdonis)
library(psych)
library(RColorBrewer)
library(DESeq2)
library(randomcoloR)
# library(vsn)
library(openxlsx)
library(factoextra)
library(PairedData)
#library(EnhancedVolcano)
library(reshape2)
library(RColorBrewer) # Load a package giving more colors
library(pheatmap) # load a package for making heatmaps
library(gridExtra)
library(grid)
library(ggpubr)
library(btools)

source("RNAseq_functions.R")
source("phyloseq_functions.R")
```

##environment
remove the first line & replace "#" with \t before!!!!!!!!!!!!!!! FOR otu & taxa

```{r}
same_dir<-T

set<-"days4" #days4  (filter boxes)
proj <- "Wasp" 
classif<-"" #
RNA_corr<-F
thresh<-0.58
use_padj<- T


metadata_dir <- paste0("C:/Users/efrat/Google Drive/Omry LAB/",proj,"/16S_",proj)
if (grepl("efratsh",getwd())) 
  metadata_dir <- paste0("C:/Users/efratsh/Google Drive/Omry LAB/",proj,"/16S_",proj)
if (grepl("amirv",getwd())) 
  metadata_dir<- paste0("C:/Users/amirv/Google Drive/Omry LAB/",proj,"/16S_",proj)

if(!same_dir){
  result_dir <- paste0(metadata_dir,"/",proj,"_results_", set, classif,"_sig_",if (use_padj) "PADJ." else "PVAL.",thresh,"_",  
                       Sys.Date(), "_",format(Sys.time(), "%s"))
  if (!file.exists(result_dir)) {dir.create(result_dir)}

}else{
  dirs <- file.info(list.dirs(path=metadata_dir,full.names = TRUE,recursive = TRUE)) %>%
                    rownames_to_column("dir") %>%
                    dplyr::filter(str_detect(dir, paste0(proj,"_results_", set))&
                                  str_detect(dir,  if (use_padj) "PADJ." else "PVAL.")) %>%
                    dplyr::filter(mtime==max(mtime)) #%>% 
  result_dir <- dirs$dir
}


```


#phyloseq

## shortcut
```{r}
#if phyloseq object already exists, load it
load(paste0(metadata_dir,"/physeq_rarefied_and_scaled",classif,"_",set,".Rdata"),verbose=T)
# openxlsx::write.xlsx(meta_16S,file=paste0(result_dir,"/meta_16S_",proj,"_",set,".xlsx"))

```

## preperations
https://joey711.github.io/phyloseq/index.html 
https://gist.github.com/erictleung/eda33bceceebb190eb9a44c93a077d32 
build phyloseq object that includes ASV+TAX+META+tree
```{r fig.height=25, fig.width=12}
Sondra_color<-list("cond_all"=c("G_MW_CTRL-G" = "#FAC31F","G_LV_CTRL-G" = "#FBE090",
                                "O_MW_CTRL-O"= "#78149b","O_LV_CTRL-O"=  "#8772E6",
                                "G_MW_Cross-G-O"= "#E1692C","G_LV_Cross-O-G"=  "#f9b693",
                                "O_MW_Cross-O-G"= "#0000ff","O_LV_Cross-G-O"=  "#55aaff"),
               Stage=c("MW"= "#800020","LV"= "#ff80a0"))

#remove the first line & replace "#" with \t before!!!!!!!!!!!!!!! FOR otu & taxa
taxa <- read.table(paste0(metadata_dir,"/exports",classif,"/taxonomy.tsv"), header = T,  sep = "\t")
taxa2 <- read.table(paste0(metadata_dir,"/exports",classif,"/taxonomy_ES.tsv"), header = T,  sep = "\t")
taxaSL<- read.table(paste0(metadata_dir,"/exports",classif,"/taxonomy_SL.tsv"), header=T,  sep="\t")
phy_tree <- read_tree(paste0(metadata_dir,"/exports",classif,"/tree.nwk"))

# split taxon column and create short_title, and get the best from 2 classifiers
taxa_16S_merged<-prep_taxa(taxa) 
taxa_16S_df2<-prep_taxa(taxa2)
taxa_16S_df_SL<-prep_taxa(taxaSL,"SL")
nrow(taxa_16S_merged)
for (r in seq_along( rownames(taxa_16S_merged))){
  #r<-2
  #print(paste(taxa_16S_merged[r,]$mark , taxa_16S_df2[r,]$mark))
  if (rownames(taxa_16S_merged[r,])== rownames(taxa_16S_df2[r,]) &
      taxa_16S_merged[r,]$mark < taxa_16S_df2[r,]$mark) {
    print(paste(rownames(taxa_16S_merged[r,]),"taxa_16S_merged:",taxa_16S_merged[r,]$short_title,
                rownames(taxa_16S_df2[r,]),"taxa_16S_df2:",taxa_16S_df2[r,]$short_title))
      print("replacing 2...")
      taxa_16S_merged[r,]<-taxa_16S_df2[r,]
  }
  if (rownames(taxa_16S_merged[r,])== rownames(taxa_16S_df_SL[r,]) &
      taxa_16S_merged[r,]$mark < taxa_16S_df_SL[r,]$mark) {
    print(paste(rownames(taxa_16S_merged[r,]),"taxa_16S_merged:",taxa_16S_merged[r,]$short_title,
                rownames(taxa_16S_df_SL[r,]),"taxa_16S_df2:",taxa_16S_df_SL[r,]$short_title))
      print("replacing SL...")
      taxa_16S_merged[r,]<-taxa_16S_df_SL[r,]
  }
}


#--------------------OTU table

#clean sample with less than 1000
otu_df_full<- read.table(paste0(metadata_dir,"/exports",classif,"/otu_table.txt"), 
                         header= T,sep="\t",check.names = F) %>%
                column_to_rownames("OTU ID")
ncol(otu_df_full)
head(table(colSums(otu_df_full)),20)
otu_df_full[,colSums(otu_df_full)<1000]

otu_df_full<-otu_df_full[,colSums(otu_df_full)> 1000 ]
ncol(otu_df_full)
head(table(colSums(otu_df_full)),20)

#read & filter according to OTU and boxes failed
meta_16S_org <- read.table(paste0(metadata_dir,"/meta-data.tsv"), header = T,  sep = "\t")
str(meta_16S_org)

meta_16S<-as.data.frame(meta_16S_org,stringsAsFactors = FALSE) %>%
              dplyr::filter(SampleID %in% colnames(otu_df_full))  %>%
              dplyr::filter(!(Box %in% c("CF17","CF7old","CF8old")) & SampleID !=59)  %>%
              mutate(cond_all=as.factor(cond_all))  %>%
              mutate(Wasp=as.factor(Wasp))  %>%
              mutate(Box=as.factor(Box))  %>%
              mutate(Stage=as.factor(Stage))  %>%
              mutate(Condition=as.factor(Condition))  #%>%

openxlsx::write.xlsx(as.data.frame(table1(~ Wasp +  Stage+ Partner + Box| Condition,
                                             data=meta_16S)),
          file=paste0(result_dir,"/Condition_table1_",proj,"_",set,".xlsx"))
nrow(meta_16S)
str(meta_16S)


#--------------------------OTU
otu_df_org <- otu_df_full %>%
                rownames_to_column("OTU.ID") %>%
                dplyr::select("OTU.ID",as.character(meta_16S$SampleID)) %>% #correlate OTU&meta
                column_to_rownames("OTU.ID")
table(colSums(otu_df_org))

#clean ASVs
head(table(rowSums(otu_df_org != 0) ))
nrow(otu_df_org)
otu_df<-otu_df_org[rowSums(otu_df_org != 0) >1, ] 
nrow(otu_df)
colnames(otu_df)[match(meta_16S$SampleID, colnames(otu_df))] = meta_16S$sample_name


#-------------------------------------------------crrolate taxa & OTU
nrow(taxa_16S_merged)
taxa_16S_df<-taxa_16S_merged %>%
            rownames_to_column("ASV") %>% 
            dplyr::filter(ASV %in% rownames(otu_df) &
                          !str_detect(Order ,"Chloroplast") &
                          !str_detect(Family ,"Mitochondria") ) %>%
            column_to_rownames("ASV")
nrow(taxa_16S_df)
#-------------------------------------------------Tree
phy_tree <- phyloseq::read_tree(paste0(metadata_dir,"/exports",classif,"/tree.nwk"))

# -----------------------change names to sample_name in OTU & META
meta_16S<-meta_16S %>%
            column_to_rownames("sample_name")
#            column_to_rownames("SampleID")


meta_16S_MW<- meta_16S %>%
                dplyr::filter(Stage=="MW")
ggplot(meta_16S_MW, aes(x=Days)) + 
  geom_density()+ 
  geom_vline(aes(xintercept=mean(Days)),
            color="blue", linetype="dashed", size=1)+
  geom_histogram(aes(y=..density..))+
  facet_wrap(~Condition, ncol=1,scale="fixed") +
  ggtitle("MW Only-Days in experiment Density Plot")
ggsave(paste0(result_dir,"/Days_in_experiment_Density_Plot_MW_",proj,"_",set,".jpg"), width=6, height=7)

meta_16S_LV<- meta_16S %>%
                dplyr::filter(Stage=="LV")
ggplot(meta_16S_LV, aes(x=Days)) + 
  geom_density()+ 
  geom_vline(aes(xintercept=mean(Days)),
            color="blue", linetype="dashed", size=1)+
  geom_histogram(aes(y=..density..))+
  facet_wrap(~Condition, ncol=1,scale="fixed") +
  ggtitle("LV Only-Days in experiment Density Plot")
ggsave(paste0(result_dir,"/Days_in_experiment_Density_Plot_LV_",proj,"_",set,".jpg"), width=6, height=7)


```

## build phyloseq object

```{r}
OTU = otu_table(as.matrix(otu_df), taxa_are_rows = TRUE)
TAX = tax_table(as.matrix(taxa_16S_df))
META <- sample_data(meta_16S)


#sanity taxa names
length(taxa_names(OTU))
length(taxa_names(TAX))
length(taxa_names(phy_tree))

head(taxa_names(OTU))
head(taxa_names(TAX))

#sanity sample names
head(sample_names(OTU))
head(sample_names(META))

physeq = phyloseq(OTU, TAX, META, phy_tree)

sample_data(physeq)
rank_names(physeq)
sample_variables(physeq)
nsamples(physeq)
physeq
```
## clean the empty, white-space, NA values taxas
not relevant as all the rest is done baset on short_title

```{r}
taxa_mtrx <- as(access(physeq, "tax_table"), "matrix")
names(taxa_mtrx)<-taxa_names(physeq)

bad_empty=c(NA, "", " ", "\t", "signed")
print(length(names(taxa_mtrx)))
keep_species <- names(taxa_mtrx)[ !(taxa_mtrx %in% bad_empty) ]
print(length(keep_species))
length(names(taxa_mtrx)[ (taxa_mtrx %in% bad_empty) ])

physeq_clean <- prune_taxa(keep_species, physeq)
physeq_clean

taxa_mtrx_clean <- as(access(physeq_clean, "tax_table"), "matrix")
names(taxa_mtrx_clean)<-taxa_names(physeq_clean)

```


##rarefy
take same number of ASVs with rarefy_even_depth
https://micca.readthedocs.io/en/latest/phyloseq.html 
```{r fig.height=8, fig.width=10}
head(table(sample_sums(physeq_clean)))
physeq_rarefied<-rarefy(physeq_clean) #physeq_filtered
str(phy_tree(physeq_rarefied))
phy_tree(physeq_rarefied) <- ape::multi2di(phy_tree(physeq_rarefied))
str(phy_tree(physeq_rarefied))
sample_names(physeq_rarefied)

```


##Scale taxa to relative abundance (of defined rank)

```{r}
#transform to relative abundance
physeq_rarefied_scaled = 
  transform_sample_counts(physeq_rarefied, function(y) 100 * y/sum(y))
physeq_rarefied_scaled

#``melted'' phyloseq data is stored much less efficiently, usually for producing graphics
physeq_rarefied_scaled_data <- psmelt(physeq_rarefied_scaled)
str(physeq_rarefied_scaled_data)

openxlsx::write.xlsx (physeq_rarefied_scaled_data, paste0(metadata_dir,"/physeq_rarefied_scaled_data",classif,".xlsx"),
                              sheetName ="physeq_rarefied_scaled_data" ,
                              row.names = TRUE, col.names = TRUE, append = FALSE) 
## Split to MW & LV
physeq_rarefied_MW <-subset_samples(physeq_rarefied, Stage=="MW")
physeq_rarefied_MW
physeq_rarefied_LV <-subset_samples(physeq_rarefied, Stage=="LV")
physeq_rarefied_LV
physeq_rarefied_scaled_MW <-subset_samples(physeq_rarefied_scaled, Stage=="MW")
physeq_rarefied_scaled_LV <-subset_samples(physeq_rarefied_scaled, Stage=="LV")
physeq_rarefied_scaled_data_MW<- physeq_rarefied_scaled_data %>%
                      dplyr::filter(Stage=="MW")
physeq_rarefied_scaled_data_LV<- physeq_rarefied_scaled_data %>%
                      dplyr::filter(Stage=="LV")
#--------------------CTRL
physeq_rarefied_ctrlG <-subset_samples(physeq_rarefied, Condition=="CTRL-G")
physeq_rarefied_ctrlG
physeq_rarefied_ctrlO <-subset_samples(physeq_rarefied, Condition=="CTRL-O")
physeq_rarefied_ctrlO
physeq_rarefied_scaled_ctrlG <-subset_samples(physeq_rarefied_scaled, Condition=="CTRL-G")
physeq_rarefied_scaled_ctrlO <-subset_samples(physeq_rarefied_scaled, Condition=="CTRL-O")
physeq_rarefied_scaled_data_ctrlG<- physeq_rarefied_scaled_data %>%
                      dplyr::filter(Condition=="CTRL-G")
physeq_rarefied_scaled_data_ctrlO<- physeq_rarefied_scaled_data %>%
                      dplyr::filter(Condition=="CTRL-O")
#--------------------Cross
physeq_rarefied_crossGO <-subset_samples(physeq_rarefied, Condition=="Cross-G-O")
physeq_rarefied_crossGO
physeq_rarefied_crossOG <-subset_samples(physeq_rarefied, Condition=="Cross-O-G")
physeq_rarefied_crossOG
physeq_rarefied_scaled_crossGO <-subset_samples(physeq_rarefied_scaled, Condition=="Cross-G-O")
physeq_rarefied_scaled_crossOG <-subset_samples(physeq_rarefied_scaled, Condition=="Cross-O-G")
physeq_rarefied_scaled_data_crossGO<- physeq_rarefied_scaled_data %>%
                      dplyr::filter(Condition=="Cross-G-O")
physeq_rarefied_scaled_data_crossOG<- physeq_rarefied_scaled_data %>%
                      dplyr::filter(Condition=="Cross-O-G")


save(meta_16S,
     taxa_16S_df,
     physeq_rarefied,
     physeq_rarefied_MW,
     physeq_rarefied_LV,
     physeq_rarefied_ctrlG,
     physeq_rarefied_ctrlO,
     physeq_rarefied_crossGO,
     physeq_rarefied_crossOG,
     physeq_rarefied_scaled_MW,
     physeq_rarefied_scaled_LV,
     physeq_rarefied_scaled_ctrlG,
     physeq_rarefied_scaled_ctrlO,
     physeq_rarefied_scaled_crossGO,
     physeq_rarefied_scaled_crossOG,
     physeq_rarefied_data,
     physeq_rarefied_scaled,
     physeq_rarefied_scaled_data,
     physeq_rarefied_scaled_data_MW,
     physeq_rarefied_scaled_data_LV,
     physeq_rarefied_scaled_data_ctrlO,
     physeq_rarefied_scaled_data_ctrlG,
     physeq_rarefied_scaled_data_crossGO,
     physeq_rarefied_scaled_data_crossOG,
     Sondra_color,
    file=paste0(metadata_dir,"/physeq_rarefied_and_scaled",classif,"_",set,".Rdata"))

qplot(log10(rowSums(otu_table(physeq_rarefied_scaled)))) +
  xlab("Logged scaled counts-per-sample")

ggplot(data = physeq_rarefied_scaled_data, 
       mapping = aes(x = Abundance, y = short_title,color=Stage,shape=Condition)) +
    geom_point()+
    ggtitle(paste(proj,"Short_title Relative Abundance"))
ggsave(paste0(result_dir,"/freq_Relative_Abund_",proj,"_",set,".jpg"), width=15, height=10)


```
### plot network
```{r fig.height=12, fig.width=12}
theme_set(theme_bw())
pdf(file = paste0(result_dir,"/Networks_", set,classif, ".pdf"))

#taxa
# ig <- make_network(physeq_rarefied_scaled, "taxa", max.dist=0.7)
# plot_network(ig, physeq_rarefied_scaled, "taxa", color = "Phylum", line_weight = 0.4, label = NULL)

#sample
ig = make_network(physeq_rarefied, type = "samples", distance = "bray", max.dist = 0.55)
plot_network(ig, physeq_rarefied, color = "cond_all",  line_weight = 0.4, point_size = 2, #shape="sex",
    label = NULL) #+ scale_color_manual(values = wes_palette("Cavalcanti1", n = 5))

ig = make_network(physeq_rarefied_MW, type = "samples", distance = "bray", max.dist = 0.55)
plot_network(ig, physeq_rarefied, color = "cond_all",  line_weight = 0.4, point_size = 2, #shape="sex",
    label = NULL) #+ scale_color_manual(values = wes_palette("Cavalcanti1", n = 5))

ig = make_network(physeq_rarefied_LV, type = "samples", distance = "bray", max.dist = 0.55)
plot_network(ig, physeq_rarefied, color = "cond_all",  line_weight = 0.4, point_size = 2, #shape="sex",
    label = NULL) #+ scale_color_manual(values = wes_palette("Cavalcanti1", n = 5))

dev.off()


```

### merge the ASVs at the each rank=level and plot. 
Given a taxonomic rank , the phyloseq function tax_glom merges the ASVs with the same taxonomy, summing the abundances.
Usually, there will be multiple ASVs with the same taxonomy assignment. These may be spurious (perpetuated errors, chimeras) or could be different species where the sequence identity is less than 97% but the genus-level taxonomy is the same. Some ASVs may also contain multiple species if they are more than 97% identical. 
```{r fig.height=8, fig.width=10}
options(getClass.msg=FALSE)
pdf(file = paste0(result_dir,"/merge_plot_taxa_Order_",set, ".pdf"),paper = "USr",width=11, height=8.5)
mpt <- merge_plot_taxa(physeq_rarefied_scaled,
                       physeq_rarefied_scaled_data,
                       field= "cond_all",
                       ord=T,
                       output_format = "",
                       rank_name = "Order")

mpt <- merge_plot_taxa(physeq_rarefied_scaled_MW,
                       physeq_rarefied_scaled_data_MW,
                       field= "cond_all",
                       ord=T,
                       set_name="MW",
                       output_format = "",
                       rank_name = "Order")

mpt <- merge_plot_taxa(physeq_rarefied_scaled_LV,
                       physeq_rarefied_scaled_data_LV,
                       field= "cond_all",
                       ord=T,
                       set_name="LV",
                       output_format = "",
                       rank_name = "Order")

dev.off()

mycolors <- distinctColorPalette(length(unique(taxa_16S_df$GenusAgg))+1)
names(mycolors) <- sample(c(unique(taxa_16S_df$GenusAgg),"Other"))

pdf(file = paste0(result_dir,"/merge_plot_taxa_Genus_",set, ".pdf"),paper = "USr",width=11, height=8.5)
mpt <- merge_plot_taxa(physeq_rarefied_scaled_MW,
                       physeq_rarefied_scaled_data_MW,
                       field= "cond_all",
                       # ord=T,
                       top_taxa = 20,
                       set_name="MW",
                       spec_colors = mycolors,
                       output_format = "",
                       rank_name = "Genus")

mpt <- merge_plot_taxa(physeq_rarefied_scaled_LV,
                       physeq_rarefied_scaled_data_LV,
                       field= "cond_all",
                       # ord=T,
                       top_taxa = 20,
                       set_name="LV",
                       spec_colors = mycolors,
                       output_format = "",
                       rank_name = "Genus")

dev.off()

```

### Prevalence Genus(cross filtered)
```{r}


getPalette = colorRampPalette(brewer.pal(9, "Set1"))

pdf(file = paste0(result_dir,"/Stacked_barplot_Genus_top20_",set, ".pdf"),paper = "USr",width=11, height=8.5)
for (stg in levels(physeq_rarefied_scaled_data$Stage)) {
  print(stg)
  genus_data <- physeq_rarefied_data %>%
                  # dplyr::filter(
                  dplyr::filter(!str_detect(Sample,"Cross") & 
                                  Abundance>0 &
                                  Stage == stg) %>%
                  mutate(Stage_Wasp=interaction(Stage,Wasp)) %>%
                  dplyr::select(Abundance,Sample,GenusAgg,Stage_Wasp) %>%
                  group_by(Stage_Wasp,Sample,GenusAgg) %>%
                  dplyr::summarise(abund = sum(Abundance)) #%>%
  genus_samples<-genus_data %>%
                  dplyr::select(Stage_Wasp,Sample) %>%
                  group_by(Stage_Wasp) %>%
                  summarise(sample_n = n_distinct(Sample))
  genus_grouped<- genus_data%>%
                  group_by(Stage_Wasp,GenusAgg) %>%
                  dplyr::summarise(mean_abund = mean(abund),
                                   taxa_n = n()) %>%
                  mutate(rel_mean_abund = mean_abund / sum(mean_abund)*100)

  data_4prev<- genus_grouped%>%
                  left_join(genus_samples)   %>%
                  mutate(prevalence=taxa_n/sample_n*100)   %>%
                  dplyr::select(-taxa_n,-sample_n,-mean_abund)   %>%
                  pivot_wider(names_from = "Stage_Wasp", values_from =c( "rel_mean_abund","prevalence"))
  assign(paste0("data_4prev_",stg),data_4prev)
  
  top<-genus_grouped %>%
        ungroup() %>%
        arrange(desc(rel_mean_abund)) %>%
        top_n(20,mean_abund) %>%
        pull(GenusAgg)
  genus_grouped_top20<- genus_grouped %>%
                          dplyr::select(Stage_Wasp,GenusAgg,rel_mean_abund)   %>%
                          mutate(GenusAgg:=ifelse(GenusAgg %in% top,GenusAgg,"Other"))   %>%
                          arrange(Stage_Wasp)
  print(ggplot(genus_grouped_top20, aes(x = as.factor(Stage_Wasp), y = rel_mean_abund, fill = GenusAgg))+ 
              labs(x="Stage_Wasp (Cross filtered)",y="Relative Abundance (%)")+ 
              ggtitle(paste("Microbiome Composition Scaled Genus top20")) + 
              geom_bar(stat = "identity", position = "stack") + 
              #facet_wrap(~eval(field), scales = "free_x") + 
              scale_fill_manual(values = getPalette(length(unique(genus_grouped_top20$GenusAgg)))) + 
              theme(axis.text.x = element_text(angle = 90),
                    panel.grid.major = element_blank(), 
                    panel.grid.minor = element_blank(),
                    panel.background = element_blank(), 
                    legend.text=element_text(size=10),
                    text=element_text(size = 14,  family="sans")) +
              guides(fill=guide_legend(ncol=2)))

}
dev.off()
openxlsx::write.xlsx (list(data_4prev_MW,data_4prev_LV), paste0(result_dir,"/Prevalence_cross_filtered_",proj,".xlsx"))
# openxlsx::write.xlsx (list(data_4prev_MW,data_4prev_LV), paste0(result_dir,"/Prevalence_W-cross_",proj,".xlsx"))

```

### Mean Rel abund short_title excel
```{r}

for (stg in levels(physeq_rarefied_scaled_data$Stage)) {
  print(stg)
  taxa_data <- physeq_rarefied_data %>%
                  dplyr::filter(Abundance>0 &   Stage == stg) %>%
                  dplyr::select(Abundance,Sample,short_title,cond_all) %>%
                  group_by(cond_all,Sample,short_title) %>%
                  dplyr::summarise(abund = sum(Abundance)) #%>%
  taxa_samples<-taxa_data %>%
                  dplyr::select(cond_all,Sample) %>%
                  group_by(cond_all) %>%
                  summarise(sample_n = n_distinct(Sample))
  taxa_grouped<- taxa_data%>%
                  group_by(cond_all,short_title) %>%
                  dplyr::summarise(mean_abund = mean(abund),
                                   taxa_n = n()) %>%
                  mutate(rel_mean_abund = mean_abund / sum(mean_abund)*100)

  data_4prev<- taxa_grouped%>%
                  left_join(taxa_samples)   %>%
                  mutate(prevalence=taxa_n/sample_n*100)   %>%
                  dplyr::select(-taxa_n,-sample_n,-mean_abund)   %>%
                  pivot_wider(names_from = "cond_all", values_from =c( "rel_mean_abund","prevalence"))
  assign(paste0("data_4prev_",stg),data_4prev)
  
}
openxlsx::write.xlsx (list(data_4prev_MW,data_4prev_LV), paste0(result_dir,"/Prevalence_short_title_",proj,".xlsx"))
```


## Alpha diversity
calculated on rarified but not scaled data
```{r}

wb <- openxlsx::createWorkbook()
pdf(file = paste0(result_dir,"/Adiversity_statistics_", set,classif, ".pdf"))
ad<-Adiver(wb_in=wb, phy_rarefied=physeq_rarefied, sheet_a= "cond_all",plot_jpg=F,
            field1="cond_all")
ad<-Adiver(wb_in=wb, phy_rarefied=physeq_rarefied_MW, sheet_a= "MW cond_all",plot_jpg=F,spec_colors=Sondra_color[[1]],
            field1="cond_all")
ad<-Adiver(wb_in=wb, phy_rarefied=physeq_rarefied_LV, sheet_a= "LV cond_all",plot_jpg=F,spec_colors=Sondra_color[[1]],
            field1="cond_all")
ad<-Adiver(wb_in=wb, phy_rarefied=physeq_rarefied, sheet_a= "Box",plot_jpg=F,
            field1="Box")
ad<-Adiver(wb_in=wb, phy_rarefied=physeq_rarefied, sheet_a= "cond_box",plot_jpg=F,
            field1="cond_all",field2="Box")

# ad<-Adiver(wb_in=wb, phy_rarefied=physeq_rarefied, sheet_a= "wasp_stage",
#             field1="Wasp",field2="Stage")
# ad<-Adiver(wb_in=wb, phy_rarefied=physeq_rarefied, sheet_a= "stage_wasp",
#             field1="Stage",field2="Wasp")
# ad<-Adiver(wb_in=wb, phy_rarefied=physeq_rarefied, sheet_a= "wasp_Partner_day", plus_f3=F,
#             field1="Wasp",field2="Partner", field3="Days")
# ad<-Adiver(wb_in=wb, phy_rarefied=physeq_rarefied_MW, sheet_a= "Only_MW_cond_box",
#             field1="Condition",field2="Box")
# ad<-Adiver(wb_in=wb, phy_rarefied=physeq_rarefied_LV, sheet_a= "Only_LV_cond_box",
#             field1="Condition",field2="Box")
# ad<-Adiver(wb_in=wb, phy_rarefied=physeq_rarefied_ctrlG, sheet_a= "Only_ctrlG_box_stg_day",plus_f3=F,
#             field1="Stage", field2="Box", field3="Days")
# ad<-Adiver(wb_in=wb, phy_rarefied=physeq_rarefied_ctrlO, sheet_a= "Only_ctrlO_box_stg_day",plus_f3=F,
#             field1="Stage", field2="Box", field3="Days")
# ad<-Adiver(wb_in=wb, phy_rarefied=physeq_rarefied_crossGO,sheet_a= "Only_crssGO_stg_box_day",plus_f3=F,
#             field1="Stage", field2="Box", field3="Days")
# ad<-Adiver(wb_in=wb, phy_rarefied=physeq_rarefied_crossOG, sheet_a= "Only_crssOG_stg_box_day",plus_f3=F,
#             field1="Stage", field2="Box", field3="Days")
#openxlsx::saveWorkbook(wb_crossOG,file=paste0(result_dir,"/diversity_statistics_crossOG_", set,classif,".xlsx"),overwrite = TRUE)
openxlsx::saveWorkbook(wb,file=paste0(result_dir,"/Adiversity_statistics_", set,classif,".xlsx"),  overwrite = TRUE)
dev.off()

```


## Beta diversity 
Plot PCoA bray as distance & calculate ANOVA
ANOVA-Test whether the phases differ significantly from each other using the permutational ANOVA (PERMANOVA) analysis
ANOVA significance tests are performed using F-tests based on sequential sums of squares from permutations of the raw data.

```{r}
ord_meths = c( "PCoA") #"DPCoA", "NMDS"
pdf(file = paste0(result_dir,"/Bdiversity_statistics_", set,classif, ".pdf"))
wbB <- openxlsx::createWorkbook()
#-----------------all
distW <- phyloseq::distance(physeq_rarefied, method="unifrac", weighted=T) #
dist_unw <- phyloseq::distance(physeq_rarefied, method="unifrac", weighted=F) #
dist_bray <- phyloseq::distance(physeq_rarefied, method="bray", weighted=T) #
md_rarefied<-data.frame(sample_data(physeq_rarefied))

# strata_col<-"Box"
# perm <- how(nperm = 1000)
# setBlocks(perm) <- with(md_rarefied, get(strata_col))

sheet_name<-paste0("Stage no strata") #,strata_col
anova_Stage <- adonis2(distW ~ Stage, data=md_rarefied) #, permutations = perm
bd2<-Bdiver(wb_in=wbB, phy_rarefied=physeq_rarefied,Bdiver_dist=F,  weight = T, dist_in = distW, 
           sheet_b= sheet_name,field1= "Stage", ellipse=F,spec_colors = Sondra_color[[2]],
           ord_meths=ord_meths, plot_jpg=F, set_name = "unifrac_w",anova_in=anova_Stage) 
anova_Stage <- adonis2(dist_unw ~ Stage, data=md_rarefied)
bd2<-Bdiver(wb_in=wbB, phy_rarefied=physeq_rarefied,Bdiver_dist=F,  weight = F,dist_in = dist_unw,
              sheet_b=sheet_name,field1="Stage",ellipse=F,
              ord_meths=ord_meths, plot_jpg=F, set_name = "unifrac_unw",anova_in=anova_Stage)  
anova_Stage <- adonis2(dist_bray ~ Stage, data=md_rarefied)
bd2<-Bdiver(wb_in=wbB, phy_rarefied=physeq_rarefied,Bdiver_dist=F,  weight = T, dist_in = dist_bray,
           sheet_b= sheet_name,field1= "Stage",ellipse=F,
           ord_meths=ord_meths, plot_jpg=F, set_name = "bray", anova_in=anova_Stage)  
  
#-----------------MW
distW <- phyloseq::distance(physeq_rarefied_MW, method="unifrac", weighted=T) #
dist_unw <- phyloseq::distance(physeq_rarefied_MW, method="unifrac", weighted=F) #
dist_bray <- phyloseq::distance(physeq_rarefied_MW, method="bray", weighted=T) #
md_rarefied<-data.frame(sample_data(physeq_rarefied_MW))

sheet_name<-paste0("MW cond no strata") #,strata_col
anova_cond <- adonis2(distW ~ cond_all, data=md_rarefied) #, permutations = perm
anova_pair_cond <- pairwiseAdonis::pairwise.adonis2(distW ~ cond_all, data=md_rarefied) #, strata = strata_col
bd2<-Bdiver(wb_in=wbB, phy_rarefied=physeq_rarefied_MW,Bdiver_dist=F,  weight = T, dist_in = distW, 
           sheet_b= sheet_name,field1= "cond_all", ellipse=F,spec_colors = Sondra_color[[1]], #to_label="SampleID",
           ord_meths=ord_meths, plot_jpg=F, set_name = "unifrac_w",anova_in=anova_cond,#ellipse=F,
           anova_pair=anova_pair_cond) 
anova_cond <- adonis2(dist_unw ~ cond_all, data=md_rarefied)
anova_pair_cond <- pairwiseAdonis::pairwise.adonis2(dist_unw ~ cond_all, data=md_rarefied)
bd2<-Bdiver(wb_in=wbB, phy_rarefied=physeq_rarefied_MW,Bdiver_dist=F,  weight = F,dist_in = dist_unw,
              sheet_b=sheet_name,field1="cond_all",ellipse=F,
              ord_meths=ord_meths, plot_jpg=F, set_name = "unifrac_unw",anova_in=anova_cond,
           anova_pair=anova_pair_cond)  
anova_cond <- adonis2(dist_bray ~ cond_all, data=md_rarefied)
anova_pair_cond <- pairwiseAdonis::pairwise.adonis2(dist_bray ~ cond_all, data=md_rarefied)
bd2<-Bdiver(wb_in=wbB, phy_rarefied=physeq_rarefied_MW,Bdiver_dist=F,  weight = T, dist_in = dist_bray,
           sheet_b= sheet_name,field1= "cond_all",ellipse=F,
           ord_meths=ord_meths, plot_jpg=F, set_name = "bray", anova_in=anova_cond,
           anova_pair=anova_pair_cond)  
  
sheet_name<-paste0("MW Days") #,strata_col
anova_cond <- adonis2(distW ~ Days, data=md_rarefied) #, permutations = perm
bd2<-Bdiver(wb_in=wbB, phy_rarefied=physeq_rarefied_MW,Bdiver_dist=F,  weight = T, dist_in = distW, 
           sheet_b= sheet_name,field1= "Days", #spec_colors = Sondra_color[[1]],ellipse=F,
           ord_meths=ord_meths, plot_jpg=F, set_name = "unifrac_w",anova_in=anova_cond) 
anova_cond <- adonis2(dist_unw ~ Days, data=md_rarefied)
bd2<-Bdiver(wb_in=wbB, phy_rarefied=physeq_rarefied_MW,Bdiver_dist=F,  weight = F,dist_in = dist_unw,
              sheet_b=sheet_name,field1="Days",
              ord_meths=ord_meths, plot_jpg=F, set_name = "unifrac_unw",anova_in=anova_cond)  
anova_cond <- adonis2(dist_bray ~ Days, data=md_rarefied)
bd2<-Bdiver(wb_in=wbB, phy_rarefied=physeq_rarefied_MW,Bdiver_dist=F,  weight = T, dist_in = dist_bray,
           sheet_b= sheet_name,field1= "Days",
           ord_meths=ord_meths, plot_jpg=F, set_name = "bray", anova_in=anova_cond)  

sheet_name<-paste0("MW Box no strata")
anova_cond <- adonis2(distW ~ Box, data=md_rarefied)#, permutations = perm
bd2<-Bdiver(wb_in=wbB, phy_rarefied=physeq_rarefied_MW,Bdiver_dist=F,  weight = T, dist_in = distW, 
           sheet_b= sheet_name,field1= "Box", ellipse=F,#spec_colors = Sondra_color[[1]],
           ord_meths=ord_meths, plot_jpg=F, set_name = "unifrac_w",anova_in=anova_cond) 
anova_cond <- adonis2(dist_unw ~ Box, data=md_rarefied)
bd2<-Bdiver(wb_in=wbB, phy_rarefied=physeq_rarefied_MW,Bdiver_dist=F,  weight = F,dist_in = dist_unw,
              sheet_b=sheet_name,field1="Box", ellipse=F,
              ord_meths=ord_meths, plot_jpg=F, set_name = "unifrac_unw",anova_in=anova_cond)  
anova_cond <- adonis2(dist_bray ~ Box, data=md_rarefied)
bd2<-Bdiver(wb_in=wbB, phy_rarefied=physeq_rarefied_MW,Bdiver_dist=F,  weight = T, dist_in = dist_bray,
           sheet_b= sheet_name,field1= "Box", ellipse=F,
           ord_meths=ord_meths, plot_jpg=F, set_name = "bray", anova_in=anova_cond)  

  

#-----------------LV
distW <- phyloseq::distance(physeq_rarefied_LV, method="unifrac", weighted=T) #
dist_unw <- phyloseq::distance(physeq_rarefied_LV, method="unifrac", weighted=F) #
dist_bray <- phyloseq::distance(physeq_rarefied_LV, method="bray", weighted=T) #
md_rarefied<-data.frame(sample_data(physeq_rarefied_LV))


sheet_name<-paste0("LV cond no strata")
anova_cond <- adonis2(distW ~ cond_all, data=md_rarefied) #, permutations = perm
anova_pair_cond <- pairwiseAdonis::pairwise.adonis2(distW ~ cond_all, data=md_rarefied) #, strata = strata_col
bd2<-Bdiver(wb_in=wbB, phy_rarefied=physeq_rarefied_LV,Bdiver_dist=F,  weight = T, dist_in = distW, 
           sheet_b= sheet_name,field1= "cond_all",ellipse=F, spec_colors = Sondra_color[[1]],
           ord_meths=ord_meths, plot_jpg=F, set_name = "unifrac_w",anova_in=anova_cond,#ellipse=F,
           anova_pair=anova_pair_cond) 
anova_cond <- adonis2(dist_unw ~ cond_all, data=md_rarefied) #, permutations = perm
anova_pair_cond <- pairwiseAdonis::pairwise.adonis2(dist_unw ~ cond_all, data=md_rarefied)#, strata = strata_col
bd2<-Bdiver(wb_in=wbB, phy_rarefied=physeq_rarefied_LV,Bdiver_dist=F,  weight = F,dist_in = dist_unw,
              sheet_b=sheet_name,field1="cond_all",ellipse=F,
              ord_meths=ord_meths, plot_jpg=F, set_name = "unifrac_unw",anova_in=anova_cond,
           anova_pair=anova_pair_cond)  
anova_cond <- adonis2(dist_bray ~ cond_all, data=md_rarefied) #, permutations = perm
anova_pair_cond <- pairwiseAdonis::pairwise.adonis2(dist_bray ~ cond_all, data=md_rarefied) #, strata = strata_col
bd2<-Bdiver(wb_in=wbB, phy_rarefied=physeq_rarefied_LV,Bdiver_dist=F,  weight = T, dist_in = dist_bray,
           sheet_b= sheet_name,field1= "cond_all",ellipse=F,
           ord_meths=ord_meths, plot_jpg=F, set_name = "bray", anova_in=anova_cond,
           anova_pair=anova_pair_cond)  
  
sheet_name<-paste0("Days no strata")
anova_cond <- adonis2(distW ~ Days, data=md_rarefied)#, permutations = perm
bd2<-Bdiver(wb_in=wbB, phy_rarefied=physeq_rarefied_LV,Bdiver_dist=F,  weight = T, dist_in = distW, 
           sheet_b= sheet_name,field1= "Days", #spec_colors = Sondra_color[[1]],ellipse=F,
           ord_meths=ord_meths, plot_jpg=F, set_name = "unifrac_w",anova_in=anova_cond) 
anova_cond <- adonis2(dist_unw ~ Days, data=md_rarefied)
bd2<-Bdiver(wb_in=wbB, phy_rarefied=physeq_rarefied_LV,Bdiver_dist=F,  weight = F,dist_in = dist_unw,
              sheet_b=sheet_name,field1="Days",
              ord_meths=ord_meths, plot_jpg=F, set_name = "unifrac_unw",anova_in=anova_cond)  
anova_cond <- adonis2(dist_bray ~ Days, data=md_rarefied)
bd2<-Bdiver(wb_in=wbB, phy_rarefied=physeq_rarefied_LV,Bdiver_dist=F,  weight = T, dist_in = dist_bray,
           sheet_b= sheet_name,field1= "Days",
           ord_meths=ord_meths, plot_jpg=F, set_name = "bray", anova_in=anova_cond)  
  
sheet_name<-paste0("LV Box no strata")
anova_cond <- adonis2(distW ~ Box, data=md_rarefied)#, permutations = perm
bd2<-Bdiver(wb_in=wbB, phy_rarefied=physeq_rarefied_LV,Bdiver_dist=F,  weight = T, dist_in = distW, 
           sheet_b= sheet_name,field1= "Box", ellipse=F,#spec_colors = Sondra_color[[1]],
           ord_meths=ord_meths, plot_jpg=F, set_name = "unifrac_w",anova_in=anova_cond) 
anova_cond <- adonis2(dist_unw ~ Box, data=md_rarefied)
bd2<-Bdiver(wb_in=wbB, phy_rarefied=physeq_rarefied_LV,Bdiver_dist=F,  weight = F,dist_in = dist_unw,
              sheet_b=sheet_name,field1="Box",ellipse=F,
              ord_meths=ord_meths, plot_jpg=F, set_name = "unifrac_unw",anova_in=anova_cond)  
anova_cond <- adonis2(dist_bray ~ Box, data=md_rarefied)
bd2<-Bdiver(wb_in=wbB, phy_rarefied=physeq_rarefied_LV,Bdiver_dist=F,  weight = T, dist_in = dist_bray,
           sheet_b= sheet_name,field1= "Box",ellipse=F,
           ord_meths=ord_meths, plot_jpg=F, set_name = "bray", anova_in=anova_cond)  


for (ord in c("_ctrlG","_ctrlO","_crossGO","_crossOG")) { 
  #ord<-"_ctrlG"
  print(paste("---------------->",ord))
  phseq<-get(paste0("physeq_rarefied",ord))
  md_rarefied<-data.frame(sample_data(phseq))
  perm <- how(nperm = 1000)

  distW <- phyloseq::distance(phseq, method="unifrac", weighted=T) #
  dist_unw <- phyloseq::distance(phseq, method="unifrac", weighted=F) #
  dist_bray <- phyloseq::distance(phseq, method="bray", weighted=T) #
  
  sheet_name<-paste0("Stage",ord)
  anova_cond <- adonis2(distW ~ Stage, data=md_rarefied)#, permutations = perm
  bd2<-Bdiver(wb_in=wbB, phy_rarefied=phseq,Bdiver_dist=F,  weight = T, dist_in = distW, 
             sheet_b= sheet_name,field1= "Stage", ellipse=F,spec_colors = Sondra_color[[2]],
             ord_meths=ord_meths, plot_jpg=F, set_name = "unifrac_w",anova_in=anova_cond) 
  anova_cond <- adonis2(dist_unw ~ Stage, data=md_rarefied)
  bd2<-Bdiver(wb_in=wbB, phy_rarefied=phseq,Bdiver_dist=F,  weight = F,dist_in = dist_unw,
                sheet_b=sheet_name,field1="Stage",ellipse=F,
                ord_meths=ord_meths, plot_jpg=F, set_name = "unifrac_unw",anova_in=anova_cond)  
  anova_cond <- adonis2(dist_bray ~ Stage, data=md_rarefied)
  bd2<-Bdiver(wb_in=wbB, phy_rarefied=phseq,Bdiver_dist=F,  weight = T, dist_in = dist_bray,
             sheet_b= sheet_name,field1= "Stage",ellipse=F,
             ord_meths=ord_meths, plot_jpg=F, set_name = "bray", anova_in=anova_cond)  

}
openxlsx::saveWorkbook(wbB,file=paste0(result_dir,"/Bdiversity_statistics_", set,classif,".xlsx"),  overwrite = TRUE)
dev.off()


# sheet_b<-"cond_wsp_d_OnlyMW"
# anova_Only_MW <- adonis2(dist ~ sample_data(physeq_rarefied_MW)$Condition*
#                             sample_data(physeq_rarefied_MW)$Wasp*
#                             sample_data(physeq_rarefied_MW)$Days)
# ad<-Bdiver(wb_in=wbB, phy_rarefied=physeq_rarefied_MW, sheet_b= sheet_b,anova=anova_Only_MW, plus_f3 = F,
#             field1="Condition",field2="Wasp", field3="Days",set_name = "unifrac_w", ellipse = F)
# anova_Only_MW <- adonis2(dist_unw ~ sample_data(physeq_rarefied_MW)$Condition*
#                             sample_data(physeq_rarefied_MW)$Wasp*
#                             sample_data(physeq_rarefied_MW)$Days)
# if (!ad) ad2<-Bdiver(wb_in=wbB, phy_rarefied=physeq_rarefied_MW,weight = F, sheet_b= sheet_b,anova=anova_Only_MW,plus_f3 = F,
#             field1="Condition",field2="Wasp", field3="Days",set_name = "unifrac_unw", ellipse = F)
# anova_Only_MW <- adonis2(dist_bray ~ sample_data(physeq_rarefied_MW)$Condition*
#                             sample_data(physeq_rarefied_MW)$Wasp*
#                             sample_data(physeq_rarefied_MW)$Days)
# if (!ad) ad2<-Bdiver(wb_in=wbB, phy_rarefied=physeq_rarefied_MW,weight = F, sheet_b= sheet_b,anova=anova_Only_MW,plus_f3 = F,
#             field1="Condition",field2="Wasp", field3="Days",set_name = "bray", ellipse = F)
# 
# 
# sheet_b<-"cond_wsp_OnlyMW"
# anova_Only_MW <- adonis2(dist ~ sample_data(physeq_rarefied_MW)$Condition*
#                             sample_data(physeq_rarefied_MW)$Wasp)
# ad<-Bdiver(wb_in=wbB, phy_rarefied=physeq_rarefied_MW, sheet_b= sheet_b,anova=anova_Only_MW, 
#             field1="Condition",field2="Wasp", set_name = "unifrac_w", ellipse = F)
# anova_Only_MW <- adonis2(dist_unw ~ sample_data(physeq_rarefied_MW)$Condition*
#                             sample_data(physeq_rarefied_MW)$Wasp)
# if (!ad) ad2<-Bdiver(wb_in=wbB, phy_rarefied=physeq_rarefied_MW,weight = F, sheet_b= sheet_b,anova=anova_Only_MW,
#             field1="Condition",field2="Wasp", set_name = "unifrac_unw", ellipse = F)
# anova_Only_MW <- adonis2(dist_bray ~ sample_data(physeq_rarefied_MW)$Condition*
#                             sample_data(physeq_rarefied_MW)$Wasp)
# if (!ad) ad2<-Bdiver(wb_in=wbB, phy_rarefied=physeq_rarefied_MW,weight = F, sheet_b= sheet_b,anova=anova_Only_MW,
#             field1="Condition",field2="Wasp", set_name = "bray", ellipse = F)

# dist <- phyloseq::distance(physeq_rarefied_LV, method="unifrac", weighted=T) #
# dist_unw <- phyloseq::distance(physeq_rarefied_LV, method="unifrac", weighted=F) #
# dist_bray <- phyloseq::distance(physeq_rarefied_LV, method="bray", weighted=T) #
# 
# anova_Only_LV <- adonis2(dist ~ sample_data(physeq_rarefied_LV)$Condition*
#                             sample_data(physeq_rarefied_LV)$Wasp*
#                             sample_data(physeq_rarefied_LV)$Days)
# ad<-Bdiver(wb_in=wbB, phy_rarefied=physeq_rarefied_LV, sheet_b= "Only_LV",anova=anova_Only_LV,plus_f3 = F,
#             field1="Condition",field2="Wasp", field3="Days",set_name = "unifrac_w", ellipse = F)
# anova_Only_LV <- adonis2(dist_unw ~ sample_data(physeq_rarefied_LV)$Condition*
#                             sample_data(physeq_rarefied_LV)$Wasp*
#                             sample_data(physeq_rarefied_LV)$Days)
# if (!ad) ad2<-Bdiver(wb_in=wbB, phy_rarefied=physeq_rarefied_LV,weight = F, sheet_b= "Only_LV",anova=anova_Only_LV,plus_f3 = F,
#             field1="Condition",field2="Wasp", field3="Days",set_name = "unifrac_unw", ellipse = F)
# anova_Only_LV <- adonis2(dist_bray ~ sample_data(physeq_rarefied_LV)$Condition*
#                             sample_data(physeq_rarefied_LV)$Wasp*
#                             sample_data(physeq_rarefied_LV)$Days)
# if (!ad) ad2<-Bdiver(wb_in=wbB, phy_rarefied=physeq_rarefied_LV,weight = F, sheet_b= "Only_LV",anova=anova_Only_LV,plus_f3 = F,
#             field1="Condition",field2="Wasp", field3="Days",set_name = "bray", ellipse = F)
# 

# dist <- phyloseq::distance(physeq_rarefied_ctrlG, method="unifrac", weighted=T) #
# dist_unw <- phyloseq::distance(physeq_rarefied_ctrlG, method="unifrac", weighted=F) #
# dist_bray <- phyloseq::distance(physeq_rarefied_ctrlG, method="bray", weighted=T) #
# 
# anova_Only_ctrlG <- adonis2(dist ~ sample_data(physeq_rarefied_ctrlG)$Stage*
#                             sample_data(physeq_rarefied_ctrlG)$Box*
#                             sample_data(physeq_rarefied_ctrlG)$Days)
# ad<-Bdiver(wb_in=wbB, phy_rarefied=physeq_rarefied_ctrlG,sheet_b= "Only_ctrlG",anova=anova_Only_ctrlG,plus_f3 = F,
#             field1="Stage", field2="Days", field3="Box",set_name = "unifrac_w", ellipse = F)
# anova_Only_ctrlG <- adonis2(dist_unw ~ sample_data(physeq_rarefied_ctrlG)$Stage*
#                             sample_data(physeq_rarefied_ctrlG)$Box*
#                             sample_data(physeq_rarefied_ctrlG)$Days)
# if (!ad) ad<-Bdiver(wb_in=wbB, phy_rarefied=physeq_rarefied_ctrlG,sheet_b= "Only_ctrlG",anova=anova_Only_ctrlG,plus_f3 = F,
#             field1="Stage", field2="Days",field3="Box",set_name = "unifrac_unw", ellipse = F)
# anova_Only_ctrlG <- adonis2(dist_bray ~ sample_data(physeq_rarefied_ctrlG)$Stage*
#                             sample_data(physeq_rarefied_ctrlG)$Box*
#                             sample_data(physeq_rarefied_ctrlG)$Days)
# if (!ad) ad<-Bdiver(wb_in=wbB, phy_rarefied=physeq_rarefied_ctrlG,sheet_b= "Only_ctrlG",anova=anova_Only_ctrlG,plus_f3 = F,
#             field1="Stage", field2="Days",field3="Box",set_name = "bray", ellipse = F)
# 
# 
# dist <- phyloseq::distance(physeq_rarefied_ctrlO, method="unifrac", weighted=T) #
# dist_unw <- phyloseq::distance(physeq_rarefied_ctrlO, method="unifrac", weighted=F) #
# dist_bray <- phyloseq::distance(physeq_rarefied_ctrlO, method="bray", weighted=T) #
# 
# anova_Only_ctrlO <- adonis2(dist ~ sample_data(physeq_rarefied_ctrlO)$Stage*
#                             sample_data(physeq_rarefied_ctrlO)$Box*
#                             sample_data(physeq_rarefied_ctrlO)$Days)
# ad<-Bdiver(wb_in=wbB, phy_rarefied=physeq_rarefied_ctrlO,sheet_b= "Only_ctrlO",anova=anova_Only_ctrlO,plus_f3 = F,
#             field1="Stage", field2="Days",field3="Box",set_name = "unifrac_w", ellipse = F)
# anova_Only_ctrlO <- adonis2(dist_unw ~ sample_data(physeq_rarefied_ctrlO)$Stage*
#                             sample_data(physeq_rarefied_ctrlO)$Box*
#                             sample_data(physeq_rarefied_ctrlO)$Days)
# if (!ad) ad<-Bdiver(wb_in=wbB, phy_rarefied=physeq_rarefied_ctrlO,sheet_b= "Only_ctrlO",anova=anova_Only_ctrlO,plus_f3 = F,
#             field1="Stage", field2="Days",field3="Box",set_name = "unifrac_unw", ellipse = F)
# anova_Only_ctrlO <- adonis2(dist_bray ~ sample_data(physeq_rarefied_ctrlO)$Stage*
#                             sample_data(physeq_rarefied_ctrlO)$Box*
#                             sample_data(physeq_rarefied_ctrlO)$Days)
# if (!ad) ad<-Bdiver(wb_in=wbB, phy_rarefied=physeq_rarefied_ctrlO,sheet_b= "Only_ctrlO",anova=anova_Only_ctrlO,plus_f3 = F,
#             field1="Stage", field2="Days",field3="Box",set_name = "bray", ellipse = F)
# 
# 
# dist <- phyloseq::distance(physeq_rarefied_crossGO, method="unifrac", weighted=T) #
# dist_unw <- phyloseq::distance(physeq_rarefied_crossGO, method="unifrac", weighted=F) #
# dist_bray <- phyloseq::distance(physeq_rarefied_crossGO, method="bray", weighted=T) #
# 
# anova_Only_crossGO <- adonis2(dist ~ sample_data(physeq_rarefied_crossGO)$Stage*
#                             sample_data(physeq_rarefied_crossGO)$Box*
#                             sample_data(physeq_rarefied_crossGO)$Days)
# ad<-Bdiver(wb_in=wbB, phy_rarefied=physeq_rarefied_crossGO,sheet_b= "Only_crossGO",anova=anova_Only_crossGO,plus_f3 = F,
#             field1="Stage", field2="Days",field3="Box",set_name = "unifrac_w", ellipse = F)
# anova_Only_crossGO <- adonis2(dist_unw ~ sample_data(physeq_rarefied_crossGO)$Stage*
#                             sample_data(physeq_rarefied_crossGO)$Box*
#                             sample_data(physeq_rarefied_crossGO)$Days)
# if (!ad) ad<-Bdiver(wb_in=wbB, phy_rarefied=physeq_rarefied_crossGO,sheet_b= "Only_crossGO",anova=anova_Only_crossGO,plus_f3 = F,
#             field1="Stage", field2="Days",field3="Box",set_name = "unifrac_unw", ellipse = F)
# anova_Only_crossGO <- adonis2(dist_bray ~ sample_data(physeq_rarefied_crossGO)$Stage*
#                             sample_data(physeq_rarefied_crossGO)$Box*
#                             sample_data(physeq_rarefied_crossGO)$Days)
# if (!ad) ad<-Bdiver(wb_in=wbB, phy_rarefied=physeq_rarefied_crossGO,sheet_b= "Only_crossGO",anova=anova_Only_crossGO,plus_f3 = F,
#             field1="Stage", field2="Days",field3="Box",set_name = "bray", ellipse = F)
# 
# 
# dist <- phyloseq::distance(physeq_rarefied_crossOG, method="unifrac", weighted=T) #
# dist_unw <- phyloseq::distance(physeq_rarefied_crossOG, method="unifrac", weighted=F) #
# dist_bray <- phyloseq::distance(physeq_rarefied_crossOG, method="bray", weighted=T) #
# 
# anova_Only_crossOG <- adonis2(dist ~ sample_data(physeq_rarefied_crossOG)$Stage*
#                             sample_data(physeq_rarefied_crossOG)$Box*
#                             sample_data(physeq_rarefied_crossOG)$Days)
# ad<-Bdiver(wb_in=wbB, phy_rarefied=physeq_rarefied_crossOG,sheet_b= "Only_crossOG",anova=anova_Only_crossOG,plus_f3 = F,
#             field1="Stage", field2="Days",field3="Box",set_name = "unifrac_w", ellipse = F)
# anova_Only_crossOG <- adonis2(dist_unw ~ sample_data(physeq_rarefied_crossOG)$Stage*
#                             sample_data(physeq_rarefied_crossOG)$Box*
#                             sample_data(physeq_rarefied_crossOG)$Days)
# if (!ad) ad<-Bdiver(wb_in=wbB, phy_rarefied=physeq_rarefied_crossOG,sheet_b= "Only_crossOG",anova=anova_Only_crossOG,plus_f3 = F,
#             field1="Stage", field2="Days",field3="Box",set_name = "unifrac_unw", ellipse = F)
# anova_Only_crossOG <- adonis2(dist_bray ~ sample_data(physeq_rarefied_crossOG)$Stage*
#                             sample_data(physeq_rarefied_crossOG)$Box*
#                             sample_data(physeq_rarefied_crossOG)$Days)
# if (!ad) ad<-Bdiver(wb_in=wbB, phy_rarefied=physeq_rarefied_crossOG,sheet_b= "Only_crossOG",anova=anova_Only_crossOG,plus_f3 = F,
#             field1="Stage", field2="Days",field3="Box",set_name = "bray", ellipse = F)


# openxlsx::saveWorkbook(wbB,file=paste0(result_dir,"/Bdiversity_statistics_", set,classif,".xlsx"),  overwrite = TRUE)

```

### Condition distances heatmap unifrac 

```{r}
distW <- phyloseq::distance(physeq_rarefied, method="unifrac", weighted=T) #
dh<-dist_heatmap(phy_rarefied=physeq_rarefied,field="cond_all",dist_in=distW,w=T, set_name = "all")
distW_MW <- phyloseq::distance(physeq_rarefied_MW, method="unifrac", weighted=T) #
dh<-dist_heatmap(phy_rarefied=physeq_rarefied_MW,field="cond_all",dist_in=distW_MW,w=T, set_name = "MW")
distW_LV <- phyloseq::distance(physeq_rarefied_LV, method="unifrac", weighted=T) #
# md_before<-data.frame(sample_data(physeq_rarefied_LV))%>%
#               arrange(cond_all)
# pairwiseAdonis::pairwise.adonis2(distW_LV ~ cond_all,data=md_before)
md_dh<-dist_heatmap(phy_rarefied=physeq_rarefied_LV,field="cond_all",dist_in=distW_LV,w=T, set_name = "LV")
# pairwiseAdonis::pairwise.adonis2(dist_4heatmap ~ cond,data=md_dh)


```
### top bacteria PCoA
```{r}
  tb<- top_bacteria(phylo=physeq_rarefied, phylo_data=physeq_rarefied_data, min_samples=20,field="cond_all") 
  taxa_lvl<-prep_count_levels(physeq_rarefied_scaled_data,taxa_level=c("GenusAgg"),rel = T)

  sb<-sig_boxplot(count_df=taxa_count_GenusAgg_rel,res_sig_list=tb,
                  res_dir=result_dir,pair_name = "cond_all", stats= F, 
                  spec_colors = Sondra_color[[1]] ,pnt_size = 1.5,
                field="cond_all",field2= "cond_all") #,group_field = "PointTime"

```

### compare cond_all rel taxa 

```{r}
library("ggradar")

abund_thresh<-1

for (stg in levels(meta_16S$Stage)) {
  wb <- openxlsx::createWorkbook()
  wb_asv <- openxlsx::createWorkbook()
  venn_list<-list()
  venn_asv_list<-list()
  i<-1
  for (cnd in unique(subset(meta_16S,Stage==stg)$cond_all)) {
    #cnd<-levels(meta_16S$cond_all)[1]
    print(cnd)
    cnd_df<-subset(physeq_rarefied_scaled_data,cond_all==cnd) %>%
                dplyr::select(Abundance,Sample,GenusAgg,cond_all) %>%
                group_by(Sample,GenusAgg,cond_all) %>%
                dplyr::summarise(rel_abund = sum(Abundance))  %>%
                dplyr::filter(rel_abund>abund_thresh) %>%
                group_by(cond_all,GenusAgg) %>%
                dplyr::summarise(mean_rel_abund = mean(rel_abund))  #%>%
    # cnd_asv_df<-subset(physeq_rarefied_scaled_data,cond_all==cnd) %>%
    #             dplyr::select(Abundance,Sample,cond_all,short_title,OTU) %>%
    #             dplyr::filter(Abundance>abund_thresh) %>%
    #             group_by(cond_all,short_title,OTU) %>%
    #             dplyr::summarise(mean_rel_abund = mean(Abundance))  %>%
    #             ungroup()
    venn_list[i]<-c(unique(subset(cnd_df,select=GenusAgg)))
    names(venn_list)[i]<-cnd
    # venn_asv_list[i]<-c(unique(subset(cnd_asv_df,select=OTU)))
    # names(venn_asv_list)[i]<-cnd
    # assign(cnd,cnd_df)
    if (i==1) taxa_4heatmap<-cnd_df else {
      taxa_4heatmap<- taxa_4heatmap %>%
                        bind_rows(cnd_df)
      for (j in 1:(i-1)){
        print(j)
        sn<-str_remove_all(paste0("comn_",names(venn_list)[i],".",names(venn_list)[j]),"-")
        addWorksheet(wb, sn)
        writeData(wb,sn,intersect(unlist(venn_list[i]),unlist(venn_list[j]))) 
        # addWorksheet(wb_asv, sn)
        # common_asv<-data.frame(OTU=intersect(unlist(venn_asv_list[i]),unlist(venn_asv_list[j]))) %>%
        #               left_join(dplyr::select(cnd_asv_df,OTU,short_title),by="OTU")
        # writeData(wb_asv,sn,common_asv) 
      }
    }
    i<-i+1
  }
  sn<-"common_all"
  addWorksheet(wb, sn)
  writeData(wb,sn,Reduce(intersect, venn_list)) 
  openxlsx::saveWorkbook(wb,file=paste0(result_dir,"/common_taxa_cond_all_Genus_rel_abund_", stg,".xlsx"),
                         overwrite= T)
  # addWorksheet(wb_asv, sn)
  # writeData(wb_asv,sn,Reduce(intersect, venn_asv_list)) 
  # openxlsx::saveWorkbook(wb_asv,file=paste0(result_dir,"/common_taxa_cond_all_ASV_rel_abund_", stg,".xlsx"),
  #                        overwrite= T)
  
  
  
  taxa_4heatmap<- taxa_4heatmap%>%
                  pivot_wider(names_from = cond_all, values_from = mean_rel_abund)%>%
                  replace(is.na(.), 0) %>% 
                  column_to_rownames("GenusAgg")
  taxa_4radar <- as.data.frame(t(taxa_4heatmap)) %>% 
          rownames_to_column("group") %>% 
          mutate_each(funs(rescale), -group) #%>%
    taxa_4radar<-taxa_4radar[ ,colSums(taxa_4radar != 0) >2 ]


  taxa_cor = corr.test(taxa_4heatmap,method = "spearman")
  taxa_cor_res<-merge(melt(taxa_cor$r),melt(taxa_cor$p), by=c("Var1","Var2")) %>%
                  dplyr::select(Var1,Var2,r=value.x,p=value.y)
  openxlsx::write.xlsx(taxa_cor_res,paste0(result_dir,"/corr_taxa_cond_all_rel_abund_",stg, ".xlsx"))
  pdf(file=paste0(result_dir,"/common_taxa_cond_all_rel_abund_",stg, ".pdf"))
  
  #correlation
  corrplot(taxa_cor$r, method = 'square', order = 'FPC', type = 'lower', diag = FALSE,
           p.mat = taxa_cor$p, sig.level = c(0.001, 0.01, 0.05), pch.cex = 0.9,insig = 'label_sig')
  
  #radar
 print(ggradar(taxa_4radar,
               base.size = 5,
               grid.label.size = 5,
               axis.label.size = 4.5,
               group.point.size = 1,
               legend.text.size = 8,
               legend.position="bottom"))


  #Make venn plot
  print(ggVennDiagram(venn_list, label_alpha = 0,set_size=3)+
      theme(legend.position = "none")+
      ggtitle(paste("compare cond_all Genus",stg,"Abundance (Thresh",abund_thresh,")")) +
      #scale_fill_gradientn(colours = terrain.colors(7))
      #paletteer::scale_fill_paletteer_c("viridis::plasma")
      scale_fill_distiller(palette = "RdBu"))
  
  pheatmap(taxa_4heatmap, cluster_cols = F, scale = "row", fontsize=6, fontsize_row = 5,
           main=paste("cond_all Genus",stg,"Abundance (Thresh",abund_thresh,") heatmap"))
  
  # ggVennDiagram(venn_asv_list, label_alpha = 0,set_size=3)+
  #   theme(legend.position = "none")+
  #   ggtitle(paste("compare cond_all ASV",stg,"Abundance (Thresh",abund_thresh,")")) +
  #   #scale_fill_gradientn(colours = terrain.colors(7))
  #   #paletteer::scale_fill_paletteer_c("viridis::plasma")
  #   scale_fill_distiller(palette = "RdBu")

  dev.off()
}


```
### compare Stage+Wasp rel taxa 

```{r}
abund_thresh<-1

physeq_rarefied_scaled_data <-physeq_rarefied_scaled_data %>%
                              mutate(Stage_Wasp=interaction(Stage,Wasp))
wb <- openxlsx::createWorkbook()
venn_list<-list()
i<-1
for (stg in levels(physeq_rarefied_scaled_data$Stage_Wasp)) {
  #stg<-levels(meta_16S$Stage_Wasp)[1]
  print(stg)
  stg_df<-subset(physeq_rarefied_scaled_data,Stage_Wasp==stg) %>%
              dplyr::select(Abundance,Sample,GenusAgg,Stage_Wasp) %>%
              group_by(Sample,GenusAgg,Stage_Wasp) %>%
              dplyr::summarise(rel_abund = sum(Abundance))  %>%
              dplyr::filter(rel_abund>abund_thresh) %>%
              group_by(Stage_Wasp,GenusAgg) %>%
              dplyr::summarise(mean_rel_abund = mean(rel_abund))  #%>%
  venn_list[i]<-c(unique(subset(stg_df,select=GenusAgg)))
  names(venn_list)[i]<-stg
  # assign(stg,stg_df)
  if (i==1) taxa_4heatmap<-stg_df else {
    taxa_4heatmap<- taxa_4heatmap %>%
                      bind_rows(stg_df)
    for (j in 1:(i-1)){
      print(j)
      sn<-str_remove_all(paste0("comn_",names(venn_list)[i],".",names(venn_list)[j]),"-")
      addWorksheet(wb, sn)
      writeData(wb,sn,intersect(unlist(venn_list[i]),unlist(venn_list[j]))) 
    }
  }
  i<-i+1
}
sn<-"common_all"
addWorksheet(wb, sn)
writeData(wb,sn,Reduce(intersect, venn_list)) 
openxlsx::saveWorkbook(wb,file=paste0(result_dir,"/common_taxa_Stage_Wasp_Genus_rel_abund_",".xlsx"),
                       overwrite= T)

taxa_4heatmap<- taxa_4heatmap%>%
                pivot_wider(names_from = Stage_Wasp, values_from = mean_rel_abund)%>%
                replace(is.na(.), 0) %>% 
                column_to_rownames("GenusAgg")

  
pdf(file=paste0(result_dir,"/common_taxa_Stage_Wasp_rel_abund_", ".pdf"))

#Make venn plot
print(ggVennDiagram(venn_list, label_alpha = 0,set_size=3)+
    theme(legend.position = "none")+
    ggtitle(paste("compare Stage_Wasp Genus","Abundance (Thresh",abund_thresh,")")) +
    #scale_fill_gradientn(colours = terrain.colors(7))
    #paletteer::scale_fill_paletteer_c("viridis::plasma")
    scale_fill_distiller(palette = "RdBu"))

pheatmap(taxa_4heatmap, cluster_cols = F, scale = "row", fontsize=6, fontsize_row = 5,
         main=paste("Stage_Wasp Genus","Abundance (Thresh",abund_thresh,") heatmap"))


dev.off()

```
<!-- ### correlation of condition -->
<!-- ```{r} -->
<!-- taxa_lvl<-prep_count_levels(physeq_rarefied_scaled_data,taxa_level=c("GenusAgg"),rel=T) -->
<!-- taxa_4cor<-taxa_count_GenusAgg_rel  %>% -->
<!--                       rownames_to_column("taxa")  %>% -->
<!--                       pivot_longer(cols=-taxa) %>% -->
<!--                       # dplyr::filter(str_detect(name,stg)) %>% -->
<!--                       separate(name, c(NA, "id"),  "_", extra = "merge") %>% -->
<!--                       dplyr::group_by(taxa,id) %>% -->
<!--                       dplyr::summarize(Mean = mean(value, na.rm=TRUE)) %>% -->
<!--                       pivot_wider(names_from = id, values_from=Mean)   %>% -->
<!--                       ungroup() %>% -->
<!--                       column_to_rownames("taxa") -->
<!-- pdf(file=paste0(result_dir,"/corr_taxa_genus_rel_abund", ".pdf")) -->
<!-- taxa_cor = corr.test(taxa_4cor,method = "spearman") -->
<!-- taxa_cor_res<-merge(melt(taxa_cor$r),melt(taxa_cor$p), by=c("Var1","Var2")) %>% -->
<!--                 dplyr::select(Var1,Var2,r=value.x,p=value.y) -->
<!--   #correlation -->
<!-- corrplot(taxa_cor$r, method = 'square', order = 'FPC', type = 'lower', diag = FALSE, -->
<!--            p.mat = taxa_cor$p, sig.level = c(0.001, 0.01, 0.05), pch.cex = 0.9,insig = 'label_sig') -->

<!-- for (stg in levels(physeq_rarefied_scaled_data$Stage)) { -->
<!--   #stg<-"LV" -->
<!--   print(stg) -->
<!--   taxa_4cor_p <- taxa_4cor %>% -->
<!--                   dplyr::select(contains(stg)) -->
<!--   taxa_cor = corr.test(taxa_4cor_p,method = "spearman") -->
<!--   taxa_cor_res_p<-merge(melt(taxa_cor$r),melt(taxa_cor$p), by=c("Var1","Var2")) %>% -->
<!--                   dplyr::select(Var1,Var2,r=value.x,p=value.y) -->
<!--   assign(paste0("taxa_cor_res_",stg),taxa_cor_res_p) -->
<!--   #correlation -->
<!--   corrplot(taxa_cor$r, method = 'square', order = 'FPC', type = 'lower', diag = FALSE, -->
<!--            p.mat = taxa_cor$p, sig.level = c(0.001, 0.01, 0.05), pch.cex = 0.9,insig = 'label_sig') -->
<!-- } -->
<!-- openxlsx::write.xlsx(list(taxa_cor_res=taxa_cor_res,taxa_cor_res_LV=taxa_cor_res_LV,taxa_cor_res_MW=taxa_cor_res_MW), -->
<!--                      file=paste0(result_dir,"/corr_taxa_genus_rel_abund", ".xlsx")) -->
<!-- dev.off() -->

<!-- ``` -->

#Radar chart
```{r}
library(fmsb)
#devtools::install_github("ricardo-bion/ggradar")

taxa_4heatmap_f<-taxa_4heatmap[rowSums(taxa_4heatmap != 0) >3, ]
taxa_4radar<-as.data.frame(rbind( t(taxa_4heatmap_f),max=rep(150,nrow(taxa_4heatmap_f)),min=rep(0,nrow(taxa_4heatmap_f))))
radarchart(taxa_4radar)


df <- as.data.frame(t(taxa_4heatmap_f)) %>% 
  rownames_to_column("taxa")

ggradar(  df,values.radar = c("0", "50", "100"),
  grid.min = 0, grid.mid = 50, grid.max = 100,
  group.colours = c("#00AFBB", "#E7B800", "#FC4E07","#00AFAA"))
```
# Proximity RandomForest
```{r}
library(randomForest)
set.seed(123)
x<-taxa_count_GenusAgg_rel %>%
    rownames_to_column("taxa") %>%
    mutate(taxa=str_replace_all(str_remove_all(taxa,"\\[|\\]"),"-","_")) %>%
    pivot_longer(-taxa, names_to = "sample_name", values_to = "Abundance") %>%
    left_join(rownames_to_column(meta_16S,"sample_name")%>% dplyr::select(sample_name,cond_all)) %>%
    pivot_wider(names_from = "taxa",values_from =  "Abundance") %>%
    column_to_rownames("sample_name")

unq_classes <- levels(meta_16S$cond_all)
n_classes   <- length(unq_classes)

genus_rf <- randomForest(cond_all ~ ., data=x, 
                        importance=TRUE,
                        proximity=TRUE)
importance(genus_rf)
var_imp <- importance(genus_rf)


prx <- genus_rf$proximity
mean_prx <- matrix(NA, n_classes, n_classes)
for (i in 1:(n_classes-1)){
  for (j in (i+1):n_classes){
    cls_i <- meta_16S$cond_all == unq_classes[i]
    cls_j <- meta_16S$cond_all == unq_classes[j]
    mean_prx[j,i] <- mean(prx[cls_i, cls_j])
  }
}

mean_prx <- mean_prx/max(mean_prx, na.rm=TRUE)
rownames(mean_prx) <- unq_classes
colnames(mean_prx) <- unq_classes
mean_prx_long<-melt(mean_prx) %>%
                dplyr::filter(!is.na(value))

pdf(file=paste0(result_dir,"/RF_importance", ".pdf"))
ggplot(mean_prx_long, aes(x=Var1,y=Var2,fill=value)) + 
    geom_tile(color = "white")+ #aes(fill = value),
    geom_text(aes(label = sprintf("%1.2f",value)), vjust = 1) + 
    theme_minimal() +
    theme(axis.text.x = element_text(angle = 90)) +
    labs(x="",y="", fill="Proximity")+
    ggtitle("Random Forest Proximity")

varImpPlot(genus_rf)
dev.off()

```


# DESeq2- taxa differential abundance 
###prep

```{r fig.height=10, fig.width=20}

md<- meta_16S %>%
      rownames_to_column("sample_name") %>%
      #dplyr::select(Wasp,Condition,Stage,sample_name,Days) %>%
      mutate(cond_day=paste0(Condition,"_",Days)) %>%
      mutate_if(is.numeric,as.factor) %>%
      arrange(sample_name) %>%
      column_to_rownames("sample_name")

taxa_count<-physeq_rarefied_data %>%
            dplyr::select(Sample,Abundance,short_title) %>%
    # only samples that has RNAseq
            #dplyr::filter(Sample %in% rownames(md)) %>%
            group_by(short_title,Sample) %>% 
            dplyr::summarise(total_abund = sum(Abundance))  %>%
            tidyr::pivot_wider(names_from =Sample, values_from =total_abund)%>%
            replace(is.na(.), 0)%>% 
 #estimateSizeFactorsForMatrix: every row contains at least one zero, cannot compute log geometric means
            replace(.==0, 1)%>% 
            dplyr::filter(!str_detect(short_title,"k__") & !str_detect( short_title,"d__")) %>%
            column_to_rownames("short_title")
ncol(taxa_count)
#filter the rows empty taxa
nrow(taxa_count)
taxa_count<-taxa_count[apply(taxa_count>=0, 1, any), ]
nrow(taxa_count)
#filter zero columns
taxa_count <- taxa_count[, colSums(taxa_count != 0) > 0]
ncol(taxa_count)
md<-md %>%
      rownames_to_column("sample_name") %>%
      dplyr::filter(sample_name %in% colnames(taxa_count)) %>%
      column_to_rownames("sample_name")
nrow(md)

#------------------Stage
for (stg in levels(md$Stage)) {
  md_p<- subset(md,Stage==stg)
  assign(paste0("md_",stg),md_p)
  assign(paste0("taxa_count_",stg),taxa_count[,rownames(md_p)])
}

```
###results+PCA
```{r fig.height=10, fig.width=20}

pdf(file=paste0(result_dir,"/Deseq_pairs_",proj,"_",set, "_short_title.pdf"),paper = "USr",width=11, height=8.5) 
for (cnd in c("Stage","Wasp","cond_all")){
  set_name<-cnd
  dsn<-as.formula(paste0(" ~ ",cnd))
  taxa_res<-deseq_part(md,taxa_count,
                          set_name=set_name,
                          value="y",
                          field="dummy",
                          dds_design=dsn,#
                          to_label=T,
                          pca_text=F,
                          out_format="",
                          res_for=cnd)
  sig_taxa_grp<-taxa_sig_pairs(taxa_thresh=thresh, taxa_all_results=taxa_res,
                               out_format="",an_colors = Sondra_color, 
                               # special_color = colorRampPalette(c("#3F831A", "white", "#B6006A"))(50),
                               special_color = colorRampPalette(c("saddlebrown", "white", "green"))(50),
                               set_name=set_name, corr=RNA_corr,use_padj = use_padj)
  save_list<-c("md",
       "taxa_count",
       taxa_res,
       "taxa_res",
       str_replace(taxa_res,"taxa","vst"),
       str_replace(taxa_res,"taxa","anno"),
       paste0("taxa_norm_abund_",set_name))
  save(list=save_list,file=paste0(metadata_dir,"/Deseq_files_",cnd,"_",set_name,"_",set,"_",proj,".Rdata")     )
}


#load(paste0(metadata_dir,"/Deseq_files_",set_name,"_",set,"_",proj,".Rdata") ,verbose = T  )

for (stg in levels(md$Stage)) {
  set_name<-paste0("Days+Wasp_",stg,"only")

  taxa_res<-deseq_part(md,taxa_count,set_name=set_name,
                  value=stg,
                  field="Stage",
                  dds_design=formula(" ~ Days +  Wasp"),
                  pca_text=F,
                  out_format="",
                  to_label=F,
                  res_for="Wasp")
  sig_taxa<-taxa_sig_pairs(taxa_thresh=thresh, taxa_all_results=taxa_res,
                               out_format="",
                               # special_color = colorRampPalette(c("#3F831A", "white", "#B6006A"))(50),
                               special_color = colorRampPalette(c("saddlebrown", "white", "green"))(50),
                               set_name=set_name, corr=RNA_corr,use_padj = use_padj)
}


# set_name<-"Wasp_LVonly"
# taxa_res_LV<-deseq_part(md,taxa_count,
#                         set_name=set_name,
#                         value="LV",
#                         field="Stage",
#                         dds_design=formula(" ~ Wasp"),#
#                         to_label=T,
#                         pca_text=F,
#                         out_format="",
#                         res_for="Wasp")
# # taxa_res_filtered<-c("taxa_y_O_MW_CTRL-O_vs_O_LV_CTRL-O","taxa_y_G_MW_CTRL-G_vs_G_LV_CTRL-G")
# sig_taxa_grp<-taxa_sig_pairs(taxa_thresh=thresh, taxa_all_results=taxa_res_LV,
#                              out_format="",an_colors = Sondra_color,
#                              set_name=set_name, corr=RNA_corr,use_padj = use_padj)


#---------------all Vs O_MW_CTRL
  set_name<-paste0("all_vs_O_MW_CTRL-O")
  all_vs_ref_res<- all_vs_ref(ref="O_MW_CTRL-O",
             ref_col="cond_all",
             design_formula=dsn, 
             res_dir = result_dir,
             md=md,
             count_data = taxa_count,
             set_name=set_name)
  sig_taxa<-taxa_sig_pairs(taxa_thresh=thresh,
                            taxa_all_results=all_vs_ref_res,
                            anno_df = cond_df,
                            an_colors = Sondra_color,
                            # special_color = colorRampPalette(c("#3F831A", "white", "#B6006A"))(50),
                            special_color = colorRampPalette(c("saddlebrown", "white", "green"))(50),
                            set_name=set_name, 
                            out_format="",
                            font_size=10,
                            use_padj = use_padj)

  #---------------subsets
#pdf(file=paste0(result_dir,"/Deseq_colors3_",proj,"_",set, "_short_title.pdf"),paper = "USr",width=11, height=8.5) 

for (stg in levels(md$Stage)) {
  for (wsp in levels(md$Wasp)) {

    set_name<-paste0(stg,"_vs_",wsp,"_",stg,"_CTRL-",wsp)
    ref<-paste0(wsp,"_",stg,"_CTRL-",wsp)
    print(ref)
    all_vs_ref_res<- all_vs_ref(ref=ref,
               ref_col="cond_all",
               design_formula=formula(" ~ cond_all"), 
               res_dir = result_dir,
               md=get(paste0("md_",stg)),
               count_data = get(paste0("taxa_count_",stg)),
               set_name=set_name)
    sig_taxa<-taxa_sig_pairs(taxa_thresh=thresh,
                              taxa_all_results=all_vs_ref_res,
                              anno_df = cond_df,
                              an_colors = Sondra_color,
                              special_color = colorRampPalette(c("saddlebrown", "white", "green"))(50),
                              # special_color = colorRampPalette(c("#3F831A", "white", "#B6006A"))(50),
                              # special_color = colorRampPalette(c("#060606", "#8D8D8D", "white"))(50),
                              set_name=set_name, 
                              out_format="",
                              font_size=10,
                              use_padj = use_padj)
  }
}
dev.off()


#----------only specific Stage
# for (stg in levels(md$Stage)) {
#   set_name<-paste0("day_cond_",stg,"_only")
# 
#   taxa_MW_res<-deseq_part(md,taxa_count,set_name=set_name,
#                   value=stg,
#                   field="Stage",
#                   dds_design=formula(" ~ Days +  Condition"),#Wasp +Box+
#                   to_label=F,
#                   res_for="Condition")
#   sig_taxa_MW<-taxa_sig_pairs(taxa_thresh=thresh, taxa_all_results=taxa_MW_res,
#                                set_name=set_name, corr=RNA_corr,use_padj = use_padj)
# }
# 

#----------only Condition=CTRL-G
# set_name<-"CTRL-G_only_day_Box_Stage"
# taxa_ctrlG_res<-deseq_part(md,taxa_count, set_name=set_name,
#                 value="CTRL-G",
#                 field="Condition",
#                 dds_design=formula(" ~ Days + Box+ Stage"),
#                 to_label=F,
#                 res_for="Stage")
# sig_taxa_ctrlG<-taxa_sig_pairs(taxa_thresh=thresh, taxa_all_results=taxa_ctrlG_res,
#                              set_name=set_name, corr=RNA_corr,use_padj = use_padj)

#----------only Condition=CTRL-O
# set_name<-"CTRL-O_only_day_Stage"
# taxa_ctrlO_res<-deseq_part(md,taxa_count, set_name=set_name,
#                 value="CTRL-O",
#                 field="Condition",
#                 dds_design=formula(" ~ Days + Stage"),  #Box+
#                 to_label=F,
#                 res_for="Stage")
# sig_taxa_ctrlO<-taxa_sig_pairs(taxa_thresh=thresh, taxa_all_results=taxa_ctrlO_res,
#                              set_name=set_name, corr=RNA_corr,use_padj = use_padj)
# 
#----------only Condition=Cross-O-G
# set_name<-"Cross-OG_only_day_Stage"
# taxa_crossOG_res<-deseq_part(md,taxa_count,set_name=set_name,
#                 value="Cross-O-G",
#                 field="Condition",
#                 dds_design=formula(" ~ Days + Stage"),#Box+
#                 to_label=F,
#                 res_for="Stage")
# sig_taxa_crossOG<-taxa_sig_pairs(taxa_thresh=thresh, taxa_all_results=taxa_crossOG_res,
#                              set_name=set_name, corr=RNA_corr,use_padj = use_padj)
#----------only Condition=Cross-G-O
# set_name<-"Cross-GO_only_day_Box_Stage"
# taxa_crossGO_res<-deseq_part(md,taxa_count,set_name=set_name,
#                 value="Cross-G-O",
#                 field="Condition",
#                 dds_design=formula(" ~ Days + Box+ Stage"),
#                 to_label=F,
#                 res_for="Stage")
# sig_taxa_crossGO<-taxa_sig_pairs(taxa_thresh=thresh, taxa_all_results=taxa_crossGO_res,
#                              set_name=set_name, corr=RNA_corr,use_padj = use_padj)
# 
#----------only specific Wasp

```


#test LRT
```{r}
dds_ignore_taxa <- DESeqDataSetFromMatrix(countData = taxa_count,
                                colData = md, ignoreRank=T,
                                design= ~ Stage+ Wasp+ Condition +Wasp:Condition) 


dds_lrt_taxa <- DESeq(dds_taxa2, test="LRT", reduced = ~1 )
resultsNames(dds_lrt_taxa)
res_lrt_taxa<-results(dds_lrt_taxa)
sig_padj_lrt<-as.data.frame(subset(res_lrt_taxa,padj<0.05))
pv <- plot_volcano(res_lrt_taxa,paste0(proj,"_",compared,"_LRT"),
                        res_dir = res_dir,
                        lfc=taxa_thresh)
colData(dds_lrt_taxa)

lrt_taxa_abund_long<-as.data.frame(counts(dds_lrt_taxa, normalized=TRUE)) %>%
                    rownames_to_column("short_title") %>%
                    dplyr::filter(short_title %in% rownames(sig_padj_lrt)) %>%
                    melt( variable.name ="sample_name",value.name = "lrt_norm_abund") %>%
                    left_join(rownames_to_column(md_all_cond,"sample_name"),by="sample_name")%>%
                    dplyr::select(-sample_name) 
ggplot(lrt_taxa_abund_long,
  aes(x = all_cond, y = lrt_norm_abund, color = short_title, group = short_title)) + 
  geom_point() + stat_summary(fun.y=mean, geom="line") +
  scale_y_log10()
ggsave(paste0(result_dir,"/LRT_",compared,"_",proj,"_",set,"_padj_sig.jpg"), width=20, height=10)

ggplot(lrt_taxa_abund_long,
  aes(x = all_cond, y = lrt_norm_abund, color = Stage, group = Stage)) + 
  geom_point() + stat_summary(fun.y=mean, geom="line") +
  scale_y_log10()
ggsave(paste0(result_dir,"/LRT_Stage_",proj,"_",set,"_padj_sig.jpg"), width=20, height=10)

```


#clustering
#### samples
```{r}
taxa_lvl<-prep_count_levels(physeq_rarefied_scaled_data,taxa_level=c("GenusAgg"),rel=T) 

count_dst <- vegdist( t(taxa_count_GenusAgg_rel))
dst <- as.matrix(count_dst)
hclst<-hclust(count_dst, method="ward.D2")
plot(hclst,hang=-1, ylab="Similarity", axes=FALSE, cex=0.5)
axis(2, seq(0, 18, by=2), seq(18, 0, by=-2))

```

#### choosing K

```{r}
set.seed(123)

wss <- (nrow(dds_ASV_abund_scaled)-1)*sum(apply(dds_ASV_abund_scaled,2,var))
for (i in 2:10) wss[i] <- sum(kmeans(dds_ASV_abund_scaled,
                                     centers=i)$withinss)
plot(1:10, wss, type="b", xlab="Number of Clusters",
     ylab="Within groups sum of squares")
```

k=6?
elbow plot to Determining Optimal Clusters
#### plot diffrernt k clustering
```{r}
count_sig_scaled <- t(dds_ASV_abund_scaled) # Centers and scales data.

pdf(file=paste0(result_dir,"/kmeans_cluster_",proj,"_",set, ".pdf"))  #,width=7, height=7

for (k in 3:6) {
  #k=6
  kmeans_cluster <- kmeans(count_sig_scaled, k)
  kmeans_cluster$centers
  print(fviz_cluster(kmeans_cluster, data = count_sig_scaled, 
                     main = paste0(proj," kmeans k=",k)))

  for (clusterNum in 1:k){
    plot(kmeans_cluster$centers[clusterNum,],	type='n',xlab="Samples",  ylab="Relative	Expression") #,	ylim=c(-4,4)
    axis(1, at = 1:ncol(count_sig_scaled),colnames(kmeans_cluster$centers))
    matlines(y=t(count_sig_scaled[kmeans_cluster$cluster==clusterNum,	]),	col='grey')
    points(kmeans_cluster$centers[clusterNum,],type='l')
    points(kmeans_cluster$centers[clusterNum,],	pch=20) #col=meta$treat,
  }
}

dev.off()
```


#Feature Selection

```{r}
set.seed(123)
#taxa_count in Genus level
taxa_lvl<-prep_count_levels(physeq_rarefied_data,taxa_level=c("GenusAgg"))
count_df_logged <- log10(taxa_count_GenusAgg+1) %>% #log transform data.
                          t() %>%
                          as.data.frame(stringsAsFactors=FALSE) %>%
                          rownames_to_column("sample_name") %>%
                          left_join(rownames_to_column(subset(meta_16S,select=c(Stage,Wasp)),"sample_name"),
                                    by="sample_name") %>%
                          mutate(Stage_Wasp=interaction(Stage,Wasp)) %>%
                          dplyr::select(-c(Stage,Wasp)) %>%
                          column_to_rownames("sample_name")

trainRowNumbers <- createDataPartition(count_df_logged$Stage_Wasp, p=0.6, list=FALSE) 
wasp_train_pre <- count_df_logged[trainRowNumbers,]
wasp_test_pre <- count_df_logged[-trainRowNumbers,]
wasp_train <- wasp_train_pre %>%
              dplyr::select(-Stage_Wasp)
wasp_test <- wasp_test_pre %>%
              dplyr::select(-Stage_Wasp)

#check that each dataset has the equivalent number of healthy and sick

prop.table(table(wasp_train_pre$Stage_Wasp))
prop.table(table(wasp_test_pre$Stage_Wasp))


#---------------------------------------Recursive feature elimination- RFE 
#select the most important features. A Random Forest algorithm is used on each iteration to evaluate the model.
RFE_ctrl <- rfeControl(functions = rfFuncs,
                   method = "repeatedcv",
                   repeats = 5,
                   verbose = FALSE)

model_RFE <- rfe(x=wasp_train, y=wasp_train_pre$Stage_Wasp, sizes=c(1:ncol(wasp_train)), rfeControl = RFE_ctrl)
model_RFE
pred_RFE<- predict(model_RFE, wasp_test_pre)
#CM_RFE <- confusionMatrix(reference =wasp_test_pre$Stage_Wasp, pred_RFE)
length(wasp_test_pre$Stage_Wasp)
length(pred_RFE$pred)
levels(wasp_test_pre$Stage_Wasp)
levels(pred_RFE$pred)

print(model_RFE$fit$confusion)
ggplot(data = model_RFE, metric = "Accuracy") + theme_bw()

g1<-ggplot(model_RFE, type=c("g", "o"))+ 
     ggtitle(paste("taxas Accuracy with RFE-Recursive Feature Elimination",
      "\n ",model_RFE$bestSubset,"taxas contribute to accuracy"))
g2<-ggplot(data = model_RFE, metric = "Kappa") + theme_bw()
if (length(model_RFE$optVariables)> 20) {
  text1<-tableGrob(data.frame(g=model_RFE$optVariables[1:(model_RFE$bestSubset/2)]),
                   theme = ttheme_minimal(base_size = 10))
  text2<-tableGrob(data.frame(g=model_RFE$optVariables[(model_RFE$bestSubset/2+1):model_RFE$bestSubset]),
                   theme = ttheme_minimal(base_size = 10))
  together <- arrangeGrob(g1,g2,text1,text2,  ncol = 2,nrow = 2)#,widths=c(1,1,3)
}else{
  text1<-tableGrob(data.frame(g=model_RFE$optVariables),
                   theme = ttheme_minimal(base_size = 10))
  text2<-tableGrob(data.frame(model_RFE$fit$confusion),
                   theme = ttheme_minimal(base_size = 10))
  together <- arrangeGrob(g1, g2,text1,text2,  ncol = 2,nrow = 2) #,widths=c(1,2,3)

}
plot(together)
dev.copy(jpeg,filename=paste0(result_dir,"/RFE_importance_",set, ".jpg"), 
           width=2000, height=700)
dev.off()

#------------------------------------rf
set.seed(123)
trctrl <- trainControl(method = "repeatedcv", number = 10, repeats = 3)
model_rf = train(Stage_Wasp ~ ., data=wasp_train_pre, method='rf', trControl=trctrl)
model_rf
pred_rf<- predict(model_rf, wasp_test_pre)
CM_rf <- confusionMatrix(reference =wasp_test_pre$Stage_Wasp, pred_rf)
ggplot(model_rf, type=c("g", "o"))+ 
     ggtitle(paste("taxas Accuracy with Random Forest",
      "\n ",model_rf$results[model_rf$results$mtry==model_rf$bestTune$mtry,]$Accuracy,"Accuracy results"))
dev.copy(jpeg,filename=paste0(result_dir,"/RF_importance_",set, ".jpg"), 
           width=1000, height=700)
dev.off()

plot(varImp(model_rf), 10, main=paste("Random Forest Variable Importance Accuracy:",
                                      round(as.numeric(CM_rf$overall['Accuracy'])*100,2),"Kappa:",
                                      round(as.numeric(CM_rf$overall['Kappa'])*100,2)))
dev.copy(jpeg,filename=paste0(result_dir,"/rf_importance_",set, ".jpg"), 
           width=900, height=500)
dev.off()


#------------------------------------------lvq
set.seed(123)
TG = expand.grid(k=1:3,size=5)
model_lvq <- caret::train(Stage_Wasp ~., data=wasp_train_pre, method="lvq", #preProcess="scale",
                          trControl=trctrl,tuneGrid=TG) 
pred_lvq<- predict(model_lvq, wasp_test_pre)
CM_lvq <- confusionMatrix(reference =wasp_test_pre$Stage_Wasp, pred_lvq)
tbl<-tableGrob(as.data.frame(CM_lvq$table),theme=ttheme_default(base_size = 4) )
g1<-varImp(model_lvq, scale=FALSE)
pdf(file = paste0(result_dir,"/lvq_importance_",set, ".pdf"))
par(mfrow =c (2,1))
plot(g1, main=paste("lvq Variable Importance Accuracy:",round(as.numeric(CM_lvq$overall['Accuracy'])*100,2)),
     cex.axis = 0.1)
plot(tbl)
par( mfrow=c(1,1))

# ROCRpred = prediction(as.numeric(pred_lvq), wasp_test_pre$Stage_Wasp)
# plot(performance(ROCRpred,"tpr","fpr"),colorize=TRUE)
dev.off()


# train the GBM model
set.seed(123)
model_gbm <- train(Stage_Wasp ~., data=wasp_train_pre, method="gbm", trControl=trctrl, verbose=FALSE)
pred_gbm<- predict(model_gbm, wasp_test_pre)
CM_gbm <- confusionMatrix(reference =wasp_test_pre$Stage_Wasp, pred_gbm)
plot(varImp(model_gbm), 10, main=paste("gbm Variable Importance Accuracy:",round(as.numeric(CM_gbm$overall['Accuracy'])*100,2)))
dev.copy(jpeg,filename=paste0(result_dir,"/gbm_importance_",set, ".jpg"), 
           width=900, height=500)
dev.off()


# train the SVM model
set.seed(123)
model_svm <- train(Stage_Wasp ~., data=wasp_train_pre, method="svmRadial", trControl=trctrl)
pred_svm<- predict(model_svm, wasp_test_pre)
CM_svm <- confusionMatrix(reference =wasp_test_pre$Stage_Wasp, pred_svm)
tbl<-tableGrob(as.data.frame(CM_svm$table),theme=ttheme_default(base_size = 4) )
pdf(file = paste0(result_dir,"/svm_importance_",set, ".pdf"))
plot(varImp(model_svm), 10, main=paste("svm Variable Importance Accuracy:",round(as.numeric(CM_svm$overall['Accuracy'])*100,2)))
plot(tbl)
dev.off()

results <- resamples(list( RF = model_rf, LVQ= model_lvq, SVM=model_svm)) # GLMNET= model_glmnet,  GBM=modelGbm

bwplot(results)
dev.copy(jpeg,filename=paste0(result_dir,"/compare_importance_",set, ".jpg"), 
           width=900, height=500)
dev.off()

```


#Picrust2
```{r}
func_abund<- openxlsx::read.xlsx(paste0(metadata_dir,"/exports/path_abun_unstrat_descrip.xlsx"))
rownames(func_abund)<-NULL
md<- meta_16S %>%
      dplyr::select(Condition,mouseID,sample_name) %>%
      mutate_if(is.numeric,as.factor) %>%
      arrange(sample_name) %>%
      column_to_rownames("sample_name")
func_all_df <- md %>%
                  dplyr::select(Condition)

func_abund_df <- func_abund %>% 
              as.data.frame(stringsAsFactors=FALSE)  %>%
              dplyr::select(-description)   %>%
              column_to_rownames("pathway") %>% 
              t()  %>% 
              as.data.frame(stringsAsFactors=FALSE)  %>%
              rownames_to_column("SampleID")  %>%
              mutate(SampleID=str_replace(SampleID,"351316-","")) %>% 
            #+Condition+Wasp
              left_join(dplyr::select(meta_16S,c(SampleID,sample_name)),by="SampleID") %>% 
              dplyr::select(-SampleID) %>% 
              dplyr::mutate_if(is.numeric, as.integer)   %>% #deseq2 works with integers
              arrange(factor(sample_name,levels=rownames(md))) %>%
              column_to_rownames("sample_name") %>% 
              t()  %>% 
              as.data.frame(stringsAsFactors=FALSE)  


dds_func <- DESeqDataSetFromMatrix(countData = func_abund_df,
                                colData = md,
                                design=~ mouseID+Condition) #
dds_func$Condition <- relevel(dds_func$Condition, ref = "before") 
dds_func<- DESeq(dds_func)
dds_func<- estimateSizeFactors(dds_func)
plotDispEsts(dds_func)
resultsNames(dds_func)

func_after_vs_before <- results(dds_func, name="Condition_after_vs_before") # , alpha=alpha
vst_funcbefore <- varianceStabilizingTransformation (dds_func, blind=FALSE)

DESeq2::summary(func_after_vs_before)

pc<- plot_PCA(dds_func,group=c("Condition"), set_name="func_Condition", dds_res = DESeq2::summary(func_after_vs_before))

ph<- plot_dds_heatmap(dds_func,group=c("Condition"), set_name="func_Condition")

func_all_res <- "func_after_vs_before"
#only func_after_vs_before
sig_func_all<-taxa_sig_pairs(taxa_thresh=1, 
                              taxa_all_results=func_all_res,set_name=set, use_padj=F, corr=F)
#---------------------------------------
sig_func_all_desc<-sig_func_all %>%
                      dplyr::rename(pathway=short_title) %>%
                      left_join(dplyr::select(func_abund,c(pathway,description)),
                                              by="pathway") %>%
                      arrange(pathway)
ggtexttable(unique(dplyr::select(sig_func_all_desc,c(pathway,description))), rows = NULL)
dev.copy(jpeg,filename=paste0(result_dir,"/Function_description_sig_lfc.",taxa_thresh,"_all_pairs_",set, ".jpg"), 
            width=800, height=nrow(sig_func_all_desc)*25)
dev.off()

```
````
