## Supplementary material for "An experimental approach towards untangling the role of nature versus nurture in shaping the microbiome of social insects": R code: Phyloseq_functions.R

```
###########################################Phyloseq_functions.R#########################
plot_PCA <- function(dds,group, set_name,dds_res,res_dir=result_dir,to_label=T,out_format="jpeg"){
  # print(group)
  vstcounts <- varianceStabilizingTransformation(dds, blind=TRUE)
  p<-DESeq2::plotPCA(vstcounts, intgroup= group) + 
    labs(title=paste('PCA', group, set_name))
  if (to_label)     p<-p+geom_text_repel(aes(label=colnames(dds)), size=3, max.overlaps=Inf, #vjust=2, 
                                         point.padding = 0.1)
  if (!missing(dds_res)) {
    if (!grepl("efratsh",getwd())){
      res_txt<-capture.output(summary(dds_res))
    } else {
      res_txt<-capture.output(DESeq2::summary(dds_res))
    }
    
    text <- paste(res_txt,collapse="\n ")
    text<-text_grob(text)
    
    together <- arrangeGrob(p, text, ncol = 2,nrow = 1)
    plot(together)
  } else print(p)
  if (out_format=="jpeg"){
    dev.copy(jpeg,filename=paste0(res_dir,"/plot_deseq_",set_name,"_PCA", ".jpg"), 
             width=1000, height=600)
    dev.off()
  }
}

plot_dds_heatmap <- function(dds,group, set_name, res_dir=result_dir,out_format="jpeg"){
  #rld = rlogTransformation(dds)
  vsd <- varianceStabilizingTransformation (dds)
  distsRL <- dist(t(assay(vsd))) # Calculate distances using transformed &normalized counts
  mat <- as.matrix(distsRL) # convert to matrix
  #mat <- mat[, order(colnames(mat))]
  colors <- colorRampPalette( rev(brewer.pal(9, "Blues")) )(255) # set colors
  print(length(group))
  if (length(group)==1){
    colnames(mat)<-paste0(colData(vsd)[,group],"_",colnames(mat))
    rownames(mat)<-paste0(colData(vsd)[,group],"_",rownames(mat))
    pheatmap(mat, legend = TRUE,
             clustering_distance_rows=distsRL,
             clustering_distance_cols=distsRL,
             col=colors, main = paste(set_name,"Sample-to-sample distances (vsd)"))
    
  } else {
    df = as.data.frame(colData(vsd)[,group]) # dataframe with a column groups
    pheatmap(mat, annotation_col=df, annotation_row = df, legend = TRUE,
             clustering_distance_rows=distsRL,
             clustering_distance_cols=distsRL,
             col=colors, main = paste(set_name,"Sample-to-sample distances (vsd)"))
    
  }
  if (out_format=="jpeg"){
    dev.copy(jpeg,filename=paste0(res_dir,"/heatmap_samples_",set_name, ".jpg"), 
             width=600, height=600)
    dev.off()
  }
  
}

plot_lfc <- function(res_df, set_name, no_phylum=FALSE, res_dir=result_dir){
  print(nrow(res_df))
  
  if (no_phylum){
    print(ggplot(res_df, aes(x=short_title, y=log2FoldChange)) +
            geom_jitter(size=3, width = 0.2) + xlab("Taxa") +
            ggtitle(paste("Taxa log2FoldChange",set_name ,"(Alpha=",alpha,")",nrow(res_df))) +
            theme(axis.text.x = element_text(angle = -90, hjust = 0, vjust=0.5, size=14)) +
            coord_flip())
    
  } else{
    print(ggplot(res_df, aes(x=short_title, y=log2FoldChange, color=Phylum)) +
            geom_jitter(size=3, width = 0.2) + xlab("Taxa") +
            ggtitle(paste("Taxa log2FoldChange",set_name ,"(Alpha=",alpha,")",nrow(res_df))) +
            theme(axis.text.x = element_text(angle = -90, hjust = 0, vjust=0.5, size=14)) +
            coord_flip())
  }
  dev.copy(jpeg,filename=paste0(res_dir,"/LFC_",set_name, "_plot.jpg"), 
             width=1100, height=600)
  dev.off()
}


# phyloseq_to_lefse
# this script convert phyloseq object into lefse recognized file format
phyloseq_to_lefse <- function(physeq, st_field="TimeW", nd_field="PatientID",short=F){
  #physeq<-physeq_rarefied     #physeq_rarefied_scaled
  #st_field="TimeW" #day
  #nd_field="PatientID"   #mouseID
  
  if (!short){
    # aggregate to one level
    ps_lefse <- physeq %>% 
  				tax_glom(taxrank = 'Species', NArm = F)
    
    # extract taxonomic data from phyloseq object and then stored in a vector called lefse_tax
    #lefse_tax <- ps_lefse %>% 
    #  tax_table %>% 
    #  data.frame(stringsAsFactors=FALSE) %>% 
    #  dplyr::select(short_title) %>%
    #  mutate(short_title=ifelse(substr(short_title,1,1)==" ",substr(short_title,2,str_length(short_title)),short_title))
    to_clean<- c("Unassigned" ,"unidentified","uncultured","gut")
    lefse_tax <- ps_lefse %>% 
              tax_table %>% 
              data.frame(stringsAsFactors=FALSE) %>% 
              rownames_to_column("OTU") %>%
              dplyr::select(-short_title,-mark)  %>%
      mutate_all(~replace(.,str_detect(.,paste(to_clean,collapse = "|")),"")) %>%
              unite(col= "lefse_title",  Domain:Species, na.rm=TRUE, sep= "|")  #%>%
              #dplyr::filter(lefse_title!="||||||")  %>%
    #take off the last |
    for (i in 1:7){
      lefse_tax<-lefse_tax %>%
                  mutate(lefse_title=ifelse(substr(lefse_title,str_length(lefse_title),str_length(lefse_title))=="|",
                                substr(lefse_title,1,(str_length(lefse_title)-1)),lefse_title))  #%>%
        
    }
  }else{
    lefse_tax <- physeq %>% 
                  tax_table %>% 
                  data.frame(stringsAsFactors=FALSE) %>% 
                  rownames_to_column("OTU") %>%
                  dplyr::select(OTU,short_title)  %>%
                  rename(lefse_title=short_title)
      
  }
  lefse_tax<-lefse_tax %>%
              mutate(lefse_title=str_replace_all(lefse_title,"-","_"))  %>%
              column_to_rownames("OTU")  
  
  # extract otu abundance matrix from phyloeq object and annotated with tax information
  lefse_abundance <- otu_table(ps_lefse) %>% 
						as.data.frame(stringsAsFactors = F) %>% 
            rownames_to_column %>% 
            right_join(rownames_to_column(lefse_tax),by="rowname")%>% 
            column_to_rownames("rowname") %>% 
						t %>% 
						data.frame(stringsAsFactors = F)

  # extract sample matadata and order sample same in lefse_matrix
  lefse_md <- sample_data(ps_lefse)%>% 
          data.frame(stringsAsFactors = F) %>%
          rownames_to_column("sample_name") %>%
					dplyr::select("sample_name",eval(st_field),eval(nd_field)) #%>%  
					#mutate_if(is.factor,as.character())  %>%
					#column_to_rownames("sample_name") #SampleID
  
  # merge 
  lefse_table <- right_join(lefse_md, rownames_to_column(lefse_abundance,"sample_name"), 
                           by = "sample_name") %>% 
                  column_to_rownames("sample_name") %>% 
                  t%>% 
                  data.frame(stringsAsFactors = F)
  lefse_table<- cbind(lefse_table[,ncol(lefse_table)], lefse_table[,1:ncol(lefse_table)-1],stringsAsFactors = F)
  lefse_table[1,1]<-st_field
  lefse_table[2,1]<-nd_field
  
  return(lefse_table)
}

my_rarefy <- function(physeq, min_sum){
  #physeq<-physeq_filtered
  set.seed(123)
#browser()
  # Plot the rarefaction curves using vegan function rarecurve()
  otu<-otu_table(physeq)
  class(otu) <- "matrix" # as.matrix() will do nothing
  rarecurve(t(otu), step=50, cex=0.5)
  dev.copy(jpeg,filename=paste0(result_dir,"/unrarefied_",set, ".jpg"), 
           width=1000, height=700)
  dev.off()
  #rarecurve(t(otu_table(physeq_clean)), step=50, cex=0.5)
  
  #Rarefy the samples without replacement. Rarefaction is used to simulate even number of reads per sample. In this example, the rarefaction depth chosen is the 80% of the minimum sample depth in the dataset (in this case ? reads per sample).
  
  sample_sums(physeq)
  # take the same number of speciman from each sample
  if (missing(min_sum)){
    physeq_rarefied <- rarefy_even_depth(physeq, rngseed=123,
                                      sample.size=0.99*min(sample_sums(physeq)), replace=T)
  } else {
    physeq_rarefied <- rarefy_even_depth(physeq, rngseed=123,
                                        sample.size=min_sum, replace=T)
  }
  physeq_rarefied
  sample_sums(physeq_rarefied)
  otu<-otu_table(physeq_rarefied)
  class(otu) <- "matrix" # as.matrix() will do nothing
  rarecurve(t(otu), step=50, cex=0.5)
  dev.copy(jpeg,filename=paste0(result_dir,"/rarefied_",set, ".jpg"), 
           width=1000, height=700)
  dev.off()
  
  sample_data(physeq_rarefied)
  rank_names(physeq_rarefied)
  sample_variables(physeq_rarefied)
  
  #phylo_data <- psmelt(physeq_rarefied)
  assign("physeq_rarefied_data",psmelt(physeq_rarefied), envir = .GlobalEnv)
  
  str(physeq_rarefied_data)
  
  #openxlsx::write.xlsx (physeq_rarefied_data,
  #                      paste0(metadata_dir,"/physeq_rarefied_data_",set,classif,".xlsx"),
  #                      sheetName ="physeq_rarefied_data" ,col.names = TRUE) 
  
  #openxlsx::write.xlsx (otu_table(physeq_rarefied),paste0(metadata_dir,"/physeq_rarefied_otu_table.xlsx"),
  #                      sheetName ="physeq_rarefied_otu_table.xlsx" ,col.names = TRUE,row.names=TRUE) 
  
  return(physeq_rarefied)
}

  prep_taxa <- function(tax,classifier="GG"){
  #tax<-taxaGT
  # split taxon column
  taxa_16S <- data.frame(stringr::str_split_fixed(tax$Taxon,";",7),stringsAsFactors=FALSE)
  colnames(taxa_16S) <- c("Domain", "Phylum", "Class", "Order", "Family", "Genus", "Species")
  rownames(taxa_16S) <- tax$OTU.ID

  #add short title
  if (classifier!="GG"){
    taxa_16S <- taxa_16S %>% 
      rownames_to_column("OTU")  %>% 
      mutate(Species=str_trim(Species,side = "left"),
             Species=str_replace_all(Species," ","_"),
             Species=ifelse(str_trim(Species)=="s__","",str_trim(Species)),
             Genus=ifelse(str_trim(Genus)=="g__","",str_trim(Genus)),
             Family=ifelse(str_trim(Family)=="f__","",str_trim(Family)),
             Order=ifelse(str_trim(Order)=="o__","",str_trim(Order)),
             Phylum=ifelse(str_trim(Phylum)=="p__","",str_trim(Phylum)))  %>%
      # add short taxa title=last rank
      mutate(short_title=if_else(!endsWith(Species,"_")& 
                                   Species!=""& 
                                   !str_detect(Species,"uncultured") & 
                                   !str_detect(Species,"_gut") & 
                                   !str_detect(Species,"metagenome") & 
                                   !str_detect(Species,"unidentified"),Species, #only GT
                                 if_else(!endsWith(Genus,"_")& 
                                           !str_detect(Genus,"uncultured") & 
                                           Genus!="",Genus,
                                         if_else(!endsWith(Family,"_")& 
                                                   !str_detect(Family,"uncultured") & 
                                                   Family!="",Family,
                                                 if_else(!endsWith(Order,"_")& 
                                                           !str_detect(Order,"uncultured") & 
                                                           Order!="",Order,
                                                         if_else(!endsWith(Class,"_")& Class!="",Class,
                                                                 if_else(!endsWith(Phylum,"_")& Phylum!="",Phylum,
                                                                         Domain))))))) %>%
      # add short taxa Genus level
      mutate(GenusAgg=if_else(!endsWith(Genus,"_")& 
                                           !str_detect(Genus,"uncultured") & 
                                           Genus!="",Genus,
                                         if_else(!endsWith(Family,"_")& 
                                                   !str_detect(Family,"uncultured") & 
                                                   Family!="",Family,
                                                 if_else(!endsWith(Order,"_")& 
                                                           !str_detect(Order,"uncultured") & 
                                                           Order!="",Order,
                                                         if_else(!endsWith(Class,"_")& Class!="",Class,
                                                                 if_else(!endsWith(Phylum,"_")& Phylum!="",Phylum,
                                                                         Domain)))))) %>%
      #add mark for level
      mutate(mark=case_when(str_detect(short_title,"k__")~1,
                            str_detect(short_title,"p__")~2,
                            str_detect(short_title,"c__")~3,
                            str_detect(short_title,"o__")~4,
                            str_detect(short_title,"f__")~5,
                            str_detect(short_title,"g__")~6,
                            str_detect(short_title,"s__")~7,
                            TRUE ~  0))   %>%
      column_to_rownames("OTU")
  }else{
    taxa_16S <- taxa_16S %>% 
      rownames_to_column("OTU")  %>% 
      # add short taxa title=last rank
      mutate(short_title=str_trim(if_else(!endsWith(Species,"_")& 
                                   Species!=""& 
                                   !str_detect(Species,"uncultured") & 
                                   !str_detect(Species,"_gut") & 
                                   !str_detect(Species,"metagenome") & 
                                   !str_detect(Species,"unidentified"),
                                 str_replace(str_replace(paste0(Genus,"_",Species)," s__",""),"g__","s__"),
                                 if_else(!endsWith(Genus,"_")& 
                                           !str_detect(Genus,"uncultured") & 
                                           Genus!="",Genus,
                                         if_else(!endsWith(Family,"_")& 
                                                   !str_detect(Family,"uncultured") & 
                                                   Family!="",Family,
                                                 if_else(!endsWith(Order,"_")& 
                                                           !str_detect(Order,"uncultured") & 
                                                           Order!="",Order,
                                                         if_else(!endsWith(Class,"_")& Class!="",Class,
                                                                 if_else(!endsWith(Phylum,"_")& Phylum!="",Phylum,
                                                                         Domain)))))))) %>%
      # add short taxa Genus level
      mutate(GenusAgg=str_trim(if_else(!endsWith(Genus,"_")& 
                                 !str_detect(Genus,"uncultured") & 
                                 Genus!="",Genus,
                               if_else(!endsWith(Family,"_")& 
                                         !str_detect(Family,"uncultured") & 
                                         Family!="",Family,
                                       if_else(!endsWith(Order,"_")& 
                                                 !str_detect(Order,"uncultured") & 
                                                 Order!="",Order,
                                               if_else(!endsWith(Class,"_")& Class!="",Class,
                                                       if_else(!endsWith(Phylum,"_")& Phylum!="",Phylum,
                                                               Domain))))))) %>%
      #add mark for level
      mutate(mark=case_when(str_detect(short_title,"k__")~1,
                            str_detect(short_title,"p__")~2,
                            str_detect(short_title,"c__")~3,
                            str_detect(short_title,"o__")~4,
                            str_detect(short_title,"f__")~5,
                            str_detect(short_title,"g__")~6,
                            str_detect(short_title,"s__")~7,
                            TRUE ~  0))   %>%
      column_to_rownames("OTU")
  }
    #remove the level letter
    #mutate_at(vars(!contains("short") & !contains("OTU")),funs(substring(.,5))) %>%
    #dplyr::filter(short_title!="Unassigned") %>%
							 
  print(paste("nrow taxa:",nrow(taxa_16S)))
  return(taxa_16S)
}

corr_gene_tax <- function(in_corr_p, in_corr_r, in_gene_res_pval_sig , in_res_sig_taxa, 
                          in_wb, in_sheet, in_tl_size=.6, pval_thresh=0.05) {
  #in_corr_p<-corr_p
  #in_corr_r<-corr_r
  #in_gene_res_pval_sig<-gene_sig_lfc
  #in_res_sig_taxa<-res_sig_taxa
  #in_tl_size=.45
  #in_sheet<-sheet_name
  
  print(paste("sheet:",in_sheet))
  #write sig cor genes per OTU to excel
  corr_p_sig<-subset(in_corr_p,pval<=pval_thresh)
  print(paste("no of cor pval<",pval_thresh,":",nrow(corr_p_sig)))
  if (nrow(corr_p_sig)>0){
    colno<-1
    for (t in unique(corr_p_sig$taxa)){
      #t<-" s__subflava" 
#browser(expr = {t == "s__unknown.1359.3321.4393"})
      corr_r_sig_t<-subset(in_corr_r,taxa==t,select=c(gene,r)) 
      corr_p_sig_t<-subset(corr_p_sig,taxa==t,select=c(gene,pval)) %>%
        left_join(corr_r_sig_t,by="gene")
      if (nrow(corr_p_sig_t)>2){
        result = tryCatch({
          
          anno_t<-
            getBM(attributes=c('external_gene_name','description', 'gene_biotype'), 
                  filters = 'mgi_symbol', #external_gene_name
                  values = corr_p_sig_t[,1], 
                  mart = mouse_ensembl,useCache = FALSE)
          #setNames(anno_t$external_gene_name,t)
          anno_t<-
            left_join(anno_t,corr_p_sig_t, by=c("external_gene_name"= "gene")) %>%
            data.table::setnames(c("external_gene_name","r"),c(eval(t),
                                                               paste(eval(t),"pval"),
                                                               paste(eval(t),"r")))
          print(colnames(anno_t)[1])
          writeData(in_wb, sheet = in_sheet,anno_t , rowNames = F, startCol = colno)
															  
        },
        error = function (condition) {
          colnames(corr_p_sig_t)<-c(t,paste(t,"pval"),paste(t,"r"))
          writeData(in_wb, sheet = in_sheet,corr_p_sig_t , rowNames = F, startCol = colno)
															  
        })
        colno=colno+5
      }
    }
    setColWidths(in_wb, in_sheet, cols = 1:colno, widths = "auto")
    
    
    #for(t in unique(in_corr_r$taxa)){
    # corr_r_4plot<-in_corr_r %>%                   
    #    mutate(gene=as.factor(gene)) %>%
    #    dplyr::filter(taxa==t) %>%
    #    arrange( r)
    #  print(ggplot(corr_r_4plot, aes(x=reorder(gene,r), y=r)) + 
    #          geom_point() +
    #          scale_y_continuous()+
    #          ggtitle(paste(in_sheet," correlation for", t)))
    #}
    
    #re-order gene by LFC
    gene_corr<- in_gene_res_pval_sig %>%
                  rownames_to_column() %>%
                  dplyr::filter(rowname %in% unique(in_corr_p$gene)) %>%
                  dplyr::arrange(log2FoldChange) %>%
                  column_to_rownames()
    nrow(gene_corr)
    taxa_corr<- in_res_sig_taxa %>%
                  rownames_to_column() %>%
                  dplyr::filter(rowname %in% unique(in_corr_p$taxa)) %>%
                  arrange(desc(log2FoldChange))  %>%
                  column_to_rownames()
    nrow(taxa_corr)
    
    #pivot_wider (move rows into columns).
    corr_p_spread<- pivot_wider(in_corr_p, names_from=gene, values_from=pval) %>% 
                      column_to_rownames("taxa") %>%
                      relocate(rownames(gene_corr))  %>%
                      as.matrix()
    nrow(corr_p_spread)
    ncol(corr_p_spread)
    
    #re-order taxa accordong to LFC

    if (nrow(corr_p_spread)>1) {
      corr_p_spread<-corr_p_spread[rownames(taxa_corr),]
    }
    corr_r_spread<- pivot_wider(in_corr_r, names_from =gene, values_from =r) %>% 
                      column_to_rownames("taxa")  %>%
                      relocate(rownames(gene_corr))  %>%
                      as.matrix()
    nrow(corr_p_spread)
    ncol(corr_p_spread)
    #re-order taxa accordong to LFC
    if (nrow(corr_r_spread)>1) {
      corr_r_spread<-corr_r_spread[rownames(taxa_corr),]
    }
#browser()
    #filter for sig correlations
    corr_p_sig_spread <-corr_p_spread%>%
      as.data.frame%>%
      select_if(~ any(.<=pval_thresh))%>%
      filter_all(any_vars(. <=pval_thresh))%>%
      as.matrix()

    #corr_p_sig_spread <-corr_p_spread[,apply(corr_p_spread<=0.05, 2, any) ]
#    if (nrow(corr_p_spread)>1) corr_p_sig_spread <-corr_p_sig_spread[apply(corr_p_sig_spread<=0.05, 1, any) ,]
    nrow(corr_p_sig_spread)
    ncol(corr_p_sig_spread)
    
    corr_r_sig_spread<-corr_r_spread%>%
                        as.data.frame%>%
                        rownames_to_column("taxa") %>%
                        dplyr::filter(taxa %in% rownames(corr_p_sig_spread))%>%
                        dplyr::select(taxa,colnames(corr_p_sig_spread)) %>%
                        column_to_rownames("taxa") %>%
                        as.matrix()
    nrow(corr_r_sig_spread)
    ncol(corr_r_sig_spread)
    
    print(paste("no of cor genes pval<",pval_thresh,"(corr_p_sig_spread):",ncol(corr_p_sig_spread)))
    print(paste("no of cor taxa pval<",pval_thresh,"(corr_p_sig_spread):",nrow(corr_p_sig_spread)))
#browser()    
    
    gene_corr_sig<- gene_corr %>%
                      rownames_to_column() %>%
                      dplyr::filter(rowname %in% colnames(corr_p_sig_spread)) #%>%
                      #column_to_rownames()
    save(gene_corr_sig,file=paste0(metadata_dir,"/sig_genes_in_sig_16S_corr_",in_sheet,".Rdata"))
    taxa_corr_sig<- taxa_corr %>%
                      rownames_to_column() %>%
                      dplyr::filter(rowname %in% rownames(corr_p_sig_spread)) #%>%
                      #column_to_rownames()
#browser()    
    #par(mfrow=c(2,1))
    #corrplot::corrplot(corr_r_spread, p.mat = corr_p_spread, method = "square", is.corr=FALSE,
    #                   insig = "label_sig", sig.level = c(.001, .01, .05), 
    #                   col = colorRampPalette(c("blue","white","red"))(100),
    #                   title=paste("\n\n\nTaxa-significant genes correlation", in_sheet,
    #                               "\nno of genes:",ncol(corr_r_spread),
    #                               "\nno of TAXAs:",nrow(corr_r_spread)-1),
    #                   cl.cex=.7, cl.ratio = 0.08,#cl.pos = 'b', cl.offset = 1,
    #                   tl.cex=in_tl_size, tl.srt=30, tl.col="black",
    #                   pch.cex = .7, pch.col = "black") #, order="FPC"
    
					  
																							  
																		   
																			
																						
																		
																		   
																			  
																	
																	  
	 
    
 
      #corrplot only pval<0.05
      
      corrplot::corrplot(corr_r_sig_spread, p.mat = corr_p_sig_spread, method = "square", is.corr=FALSE,
                         insig = "label_sig", sig.level = c(.001, .01, .05), 
                         col = colorRampPalette(c("blue","white","red"))(100),
																						 
                         #title=paste("\nno of genes:",ncol(corr_r_sig_spread),
                        #             "no of TAXAs:",nrow(corr_r_sig_spread)),
                         cl.cex=.5, cl.ratio = 0.1,#cl.pos = 'b', cl.offset = 1,
                         tl.cex=in_tl_size, tl.srt=30, tl.col="black",
                         pch.cex = in_tl_size, pch.col = "black") #, order="FPC"
      #dev.copy(jpeg,filename=paste0(result_dir,"/corrplot_sig_genes_in_sig_16S_corr_",in_sheet, ".jpg"), 
      #         width=700, height=1000)
      #dev.off()
      
      # genes LFC barplot

      print(ggplot(gene_corr_sig, aes(fill=pvalue, 
                                y=log2FoldChange, 
                                x=reorder(rowname,log2FoldChange)))+
        geom_bar(position="dodge", stat="identity")+
        theme_grey(base_size = 10) +
        xlab("Genes")+
        ylab("log2FoldChange") +
        ggtitle(paste("barplot genes sig lfc.",gene_thresh,in_sheet,set))+
        theme(axis.text.x = element_text(angle = 45))+
        theme(panel.grid.major = element_blank(), panel.grid.minor = element_blank(),
              panel.background = element_blank())+
        theme(plot.margin=unit(c(0,1,0,3),"cm")))
  
      # taxa LFC barplot
      print(ggplot(taxa_corr_sig, aes(fill=pvalue, 
                                    y=log2FoldChange, 
                                    x=reorder(rowname,log2FoldChange)))+
              geom_bar(position="dodge", stat="identity")+
              theme_grey(base_size = 20) +
              ggtitle(paste("barplot taxa sig lfc.",taxa_thresh,in_sheet,set))+
              theme(axis.title.y= element_blank(), panel.grid.minor = element_blank(),
                    panel.background = element_blank())+ 
              coord_flip())
      #save(taxa_corr, gene_corr, corr_r_sig_spread,corr_p_sig_spread,
      #     file=paste0(metadata_dir,"/data_4corr_",set,"_",in_sheet,".Rdata"))
    
  }  
}


merge_plot_taxa <- function(physeq_r, physeq_r_scaled_data, set_name=set, output_format="jpg",
                            legend_ncol=3, ord=F, top_taxa, hmap=T,spec_colors,
                            field,field2,field3,rank_name=c("Genus","short_title")){
  #https://micca.readthedocs.io/en/latest/phyloseq.html
  #physeq_r_scaled<-physeq_rarefied_scaled
  #physeq_r_scaled_data<-physeq_rarefied_scaled_data
  #rank_name<-c("short_title")
  #field= "Control_GDM";  field2= "trimester"
  getPalette = colorRampPalette(brewer.pal(9, "Set1"))
  sampleColor <- getPalette(max(nsamples(physeq_r)))
  #for (r in rank_names(physeq_r_scaled)){
  for (r in rank_name){
      #The option NArm set to FALSE forces the function to keep the unclassified OTUs at the phylum level. 
    #r<-"short_title"
    print(paste("r:",r))
    #scaled  
#browser()
    #physeq_r_scaled_merged <- aggregate_taxa(physeq_r_scaled, level=r)

    physeq_r_merged <- tax_glom(physeq_r, taxrank=r, NArm=T)
    
    #tax_table(physeq_r_merged)<-tax_table(physeq_r)[,1:which(rank_names(physeq_r)==r)]
    physeq_r_scaled_merged <- transform_sample_counts(physeq_r_merged, function(y) 100 * y/sum(y))
    print(physeq_r_scaled_merged)
    #abundances(physeq_r_scaled_merged)
    
    #cleaner bar plot:
    if (r!="Domain" & r!="Class"){
      #r<-"short_title"
      PhylaPalette <- getPalette(max(nrow(unique(as.data.frame(taxa_16S_df[,r])))))
#browser()
      if (ord){
        physeq_r_scaled_data<-psmelt(physeq_r_scaled_merged)
        p<-ggplot(physeq_r_scaled_data, aes(x = as.factor(Sample), y = Abundance, fill = fct_reorder(get(r), Abundance))) 
        
      } else{
        p<-phyloseq::plot_bar(physeq_r_scaled_merged, fill=r) 
      }
      if (missing(top_taxa)){
        print(p+ 
                ggtitle(paste("Abundance",r, "scaled& merged per",eval(field), "-",set_name))+
                scale_fill_manual(values = PhylaPalette) + #, guide = guide_legend(nrow=60)
                geom_bar(stat = "identity", position = "stack") + 
                theme(axis.text.x = element_text(angle = 90),
                      panel.grid.major = element_blank(), 
                      panel.grid.minor = element_blank(),
                      panel.background = element_blank(), 
                      axis.line = element_line(colour = "black"),
                      text=element_text(size = 10,  family="sans"),
                      legend.text=element_text(size=7)) +
                 guides(fill=guide_legend(ncol=legend_ncol,title=r))+
                facet_wrap(~get(field), scales="free_x", nrow=1))
        #ggsave(paste0(result_dir,"/barplot_",r,"_scaled_merged_",eval(field),"_",set_name,".pdf"), width = 8, height = 5, device = "pdf", dpi=300, units="in")
          if (output_format=="jpg"){
           dev.copy(jpeg,filename=paste0(result_dir,"/barplot_",r,"_scaled_merged_",eval(field),"_",set_name, ".jpg"), 
                   width=nrow(unique(as.data.frame(taxa_16S_df[,r])))*3+2000, height=1200)
            dev.off()
          }
      }
        if(!missing(field2)) {
        print(phyloseq::plot_bar(physeq_r_scaled_merged, fill=r, 
                                 title=paste("Abundance",r, "scaled& merged per",eval(field), "-",set_name))+ 
                scale_fill_manual(values = PhylaPalette) + 
                theme(axis.text.x = element_text(angle = 90),
                      panel.grid.major = element_blank(), 
                      panel.grid.minor = element_blank(),
                      panel.background = element_blank(), 
                      axis.line = element_line(colour = "black"),
                      text=element_text(size = 9,  family="sans"),
                      legend.text=element_text(size=5)) +
                guides(fill=guide_legend(ncol=legend_ncol))+
                facet_wrap(~get(field2), scales="free_x", nrow=1))
        if (output_format=="jpg"){
            dev.copy(jpeg,filename=paste0(result_dir,"/barplot_",r,"_scaled_merged_",eval(field2),"_",set_name, ".jpg"), 
                 width=nrow(unique(as.data.frame(taxa_16S_df[,r])))*3+1000, height=1200)
          dev.off()
        }
        }     
      
#browser()
      #plot_heatmap(physeq )
      if (length(phy_tree(physeq_r_scaled_merged,errorIfNULL =F))!=0 & hmap){
        plot_heatmap(physeq_r_scaled_merged)
        print(plot_heatmap(physeq_r_scaled_merged, taxa.label=r, method = "DPCoA", distance = "bray",
                           title=paste("Abundance",r, "scaled& merged-",set_name)))
        if (output_format=="jpg"){
          
          ggsave(paste0(result_dir,"/plot_",r,"_scaled_merged_sample_heatmap","_",set_name, ".jpg"), 
                 width=10, height=10)
          
          dev.off()
        }
      }
      physeq_r_merged_sample <- merge_samples(physeq_r_merged, field) #physeq_r_scaled_merged
      physeq_r_merged_sample_scaled <- transform_sample_counts(physeq_r_merged_sample, function(x) x*100 / sum(x))
      physeq_r_scaled_data<-psmelt(physeq_r_merged_sample_scaled)
#browser()          
      if (!missing(top_taxa)){
        top<-physeq_r_scaled_data %>%
              dplyr::select(eval(r),Abundance)   %>%
              group_by(get(r))  %>%
              dplyr::summarize(mean_abund = mean(Abundance, na.rm=TRUE)) %>%
              arrange(desc(mean_abund), .by_group = TRUE) %>%
              top_n(top_taxa,mean_abund) %>%
              dplyr::select(V1='get(r)') %>%
              pull(V1)
        
        physeq_r_scaled_data<- physeq_r_scaled_data %>%
                                dplyr::select(Sample,!!r,Abundance)   %>%
                                mutate(!!sym(r):=ifelse(!!sym(r) %in% top,!!sym(r),"Other"))   %>%
                                arrange(Sample)
        return(top)
        # openxlsx::write.xlsx(physeq_r_scaled_data,
        #                      paste0(result_dir,"/top",top_taxa,"_",r,"_data_",proj,"_",set,"_",set_name,".xlsx"),
        #                      rowNames = F, overwrite = T)
#browser()          
      }
      if (ord){
        p<-ggplot(physeq_r_scaled_data, aes(x = as.factor(Sample), y = Abundance, fill = fct_reorder(get(r), Abundance))) 
      } else{
        p<-ggplot(physeq_r_scaled_data, aes(x = as.factor(Sample), y = Abundance, fill = get(r)))
        # p<-phyloseq::plot_bar(physeq_r_merged_sample_scaled, fill=r)
        # p<-ggplot(physeq_r_scaled_data, aes(x = factor(Sample,levels=c("Child","Sibling","Father","Mother")), y = Abundance, fill=get(r))) 
      }
#browser()
      getPalette = colorRampPalette(brewer.pal(9, "Set1"))
      print(p+ 
              labs(x=eval(field),y="Relative Abundance (%)")+ 
              ggtitle(paste("Microbiota Composition Scaled" ,r,
                            ifelse(!missing(top_taxa),paste0("top",top_taxa),""),"-",set_name)) + 
              geom_bar(stat = "identity", position = "stack") + 
              #facet_wrap(~eval(field), scales = "free_x") + 
              {if(!missing(spec_colors)) scale_fill_manual(values = spec_colors) else scale_fill_manual(values = getPalette(length(unique(physeq_r_scaled_data[[r]])))) } +
              theme(axis.text.x = element_text(angle = 90),
                    panel.grid.major = element_blank(), 
                    panel.grid.minor = element_blank(),
                    panel.background = element_blank(), 
                    legend.text=element_text(size=10),
                    text=element_text(size = 14,  family="sans")) +
              guides(fill=guide_legend(ncol=legend_ncol,title=r)))
      if (output_format=="jpg"){
          dev.copy(jpeg,filename=paste0(res_dir,"/plot_",r,"_scaled_",eval(field),"_grouped_",set_name, ".jpg"), 
                   width=1600, height=1000)
          dev.off()
      }
          
        if(!missing(field2)) {
          print("field2")
          sample_data(physeq_r_scaled_merged)$cond<- paste0(sample_data(physeq_r_scaled_merged)[[field]],".",sample_data(physeq_r_scaled_merged)[[field2]])
          physeq_r_merged_sample <- merge_samples(physeq_r_scaled_merged, "cond")
          sample_data(physeq_r_merged_sample)$cond = sample_names(physeq_r_merged_sample)
          sample_data(physeq_r_merged_sample) <- data.frame(sample_data(physeq_r_merged_sample)) %>%
                                                        separate(cond,into=c(field,field2))
          print(data.frame(sample_data(physeq_r_merged_sample)))
          physeq_r_merged_sample_scaled <- transform_sample_counts(physeq_r_merged_sample, function(x) x / sum(x))
          
          print(phyloseq::plot_bar(physeq_r_merged_sample_scaled, fill=r)+ 
                  ylab(r)+ 
                  xlab(eval(field2)) +
                  ggtitle(paste("Microbiota Composition per ",eval(field),"+",eval(field2)," Scaled" ,r,"-",set_name)) + 
                  geom_bar(stat = "identity", position = "stack") + 
                  facet_wrap(~get(field), scales = "free_x") + 
                  scale_fill_manual(values = PhylaPalette) + 
                  theme(axis.text.x = element_text(angle = 90),
                        legend.text=element_text(size=5),
                        text=element_text(size = 9,  family="sans")) +
                  guides(fill=guide_legend(ncol=legend_ncol)))
          if (output_format=="jpg"){
            dev.copy(jpeg,filename=paste0(result_dir,"/plot_",r,"_scaled_",eval(field2),"_grouped_",eval(field),"_",set_name, ".jpg"), 
                     width=1600, height=1000)
            dev.off()
          }
    
          # print(ggplot(physeq_r_scaled_data_filled, 
          #              aes(x = get(field), y = Abundance, factor(get(r)),
          #                  fill = factor(get(r)))) + 
          print(phyloseq::plot_bar(physeq_r_merged_sample_scaled,fill=r)+ 
                  ylab(r)+ 
                  xlab(eval(field)) +
                  ggtitle(paste("Microbiota Composition per ",eval(field2)," Scaled" ,r,"-",set_name)) + 
                  geom_bar(stat = "identity", position = "stack") + 
                  facet_wrap(~get(field2), scales = "free_x", nrow=1) + 
                  scale_fill_manual(values = PhylaPalette) + 
                  theme(legend.text=element_text(size=5),
                        text=element_text(size = 9,  family="sans")) +
                  guides(fill=guide_legend(ncol=legend_ncol)))
          if (output_format=="jpg"){
            dev.copy(jpeg,filename=paste0(result_dir,"/plot_",r,"_scaled_",eval(field),"_grouped_",eval(field2),"_",set_name, ".jpg"), 
                     width=1600, height=1000)
            dev.off()
          }
        }
        if(!missing(field3)) {
          print("field3")
          # print(ggplot(physeq_r_scaled_data_filled, 
          #              aes(x = get(field3), y = Abundance, factor(get(r)),
          #                  fill = factor(get(r)))) + 
          sample_data(physeq_r_scaled_merged)$cond<- paste0(sample_data(physeq_r_scaled_merged)[[field]],".",
                                                            sample_data(physeq_r_scaled_merged)[[field2]],".",
                                                            sample_data(physeq_r_scaled_merged)[[field3]])
          physeq_r_merged_sample <- merge_samples(physeq_r_scaled_merged, cond)
          sample_data(physeq_r_merged_sample)$cond = sample_names(physeq_r_merged_sample)
          sample_data(physeq_r_merged_sample) <- data.frame(sample_data(physeq_r_merged_sample)) %>%
                                                      separate(cond,into=c(field,field2,field3))
          print(data.frame(sample_data(physeq_r_merged_sample)))
          physeq_r_merged_sample_scaled <- transform_sample_counts(physeq_r_merged_sample, function(x) x / sum(x))
          
          print(phyloseq::plot_bar(physeq_r_merged_sample_scaled, fill=r)+ 
                  ylab(r)+ 
                  xlab(eval(field3)) +
                  ggtitle(paste("Microbiota Composition per ",eval(field),"+",eval(field3)," Scaled" ,r,"-",set_name)) + 
                  geom_bar(stat = "identity", position = "stack") + 
                  facet_wrap(~get(field), scales = "free_x") + 
                  scale_fill_manual(values = PhylaPalette) + 
                  theme(legend.text=element_text(size=5),
                        text=element_text(size = 9,  family="sans")) +
                  guides(fill=guide_legend(ncol=legend_ncol)))
          if (output_format=="jpg"){
            dev.copy(jpeg,filename=paste0(result_dir,"/plot_",r,"_scaled_",eval(field3),"_grouped_",eval(field),"_",set_name, ".jpg"), 
                     width=1600, height=1000)
            dev.off()
          }
          # print(ggplot(physeq_r_scaled_data_filled, 
          #              aes(x = get(field), y = Abundance, factor(get(r)),
          #                  fill = factor(get(r)))) + 
          print(phyloseq::plot_bar(physeq_r_merged_sample1_scaled, fill=r)+ 
                  ylab(r)+ 
                  xlab(eval(field)) +
                  ggtitle(paste("Microbiota Composition per ",eval(field3)," Scaled" ,r,"-",set_name)) + 
                  geom_bar(stat = "identity", position = "stack") + 
                  facet_wrap(~get(field3), scales = "free_x", nrow=1) + 
                  scale_fill_manual(values = PhylaPalette) + 
                  theme(legend.text=element_text(size=5),
                        text=element_text(size = 9,  family="sans")) +
                  guides(fill=guide_legend(ncol=legend_ncol)))
          if (output_format=="jpg"){
            dev.copy(jpeg,filename=paste0(result_dir,"/plot_",r,"_scaled_",eval(field),"_grouped_",eval(field3),"_",set_name, ".jpg"), 
                     width=1600, height=1000)
            dev.off()
          }
        }  
#      }
    }
  }
}

Abundance_boxplot <- function(physeq_data,res_dir=result_dir, lvl="OTU",
                            field="TimeW",field2="PatientID"){
  #physeq_data<-physeq_anova
  #field="RatType"
  #field2="TimeGroup"
  #res_dir=result_dir
  
  wb <- openxlsx::createWorkbook()
  sheet_name <- "taxa_abundance_stats"
  addWorksheet(wb, sheet_name)
  titStyle <- createStyle(fontColour = "#006100", fgFill = "#C6EFCE")
  st_row<-1
  for (sht in unique(physeq_data$short_title)) {
    #sht<-" f__S24-7" #" g__Prevotella" 
    print(sht)
    physeq_data_p<- subset(physeq_data,short_title==sht, select=-short_title)
    print(paste("no of OTUs:", nrow(physeq_data_p)))
    
    # ANOVA-Test whether the groups, TimeW,  differ significantly from each
    if (nrow(subset(physeq_data_p,Abundance>0))>2 & nrow(physeq_data_p)<400){
      # prevent zero values
      physeq_data_p_trans<-physeq_data_p
      physeq_data_p_trans$Abundance<-sqrt (1+physeq_data_p$Abundance)
      #anova <- adonis(Abundance ~ get(field), data=physeq_data_p_trans)
      anova <- adonis(formula =Abundance ~ get(field)+ get(field2), data=physeq_data_p_trans)
      anova2 <- adonis(formula =Abundance ~ get(field2)+ get(field), data=physeq_data_p_trans)
      print(anova2)
      anova_2way <- aov(Abundance ~  get(field2)+get(field), data=physeq_data_p_trans) #
      tukey<- TukeyHSD(anova_2way)
      tukey_df<-rbind(as.data.frame(tukey$`get(field)`), as.data.frame(tukey$`get(field2)`))
      tukey_df_sig<-subset(tukey_df,tukey_df$`p adj` <0.05)
      anova_2way2 <- aov(Abundance ~  get(field)+get(field2), data=physeq_data_p_trans) #
      tukey2<- TukeyHSD(anova_2way2)
      tukey_df2<-rbind(as.data.frame(tukey$`get(field2)`), as.data.frame(tukey$`get(field)`))
      tukey_df_sig2<-subset(tukey_df,tukey_df$`p adj` <0.05)
      if (!is.na(anova$aov.tab$`Pr(>F)`[1]) | 
          !is.na(anova$aov.tab$`Pr(>F)`[2]) | 
          !is.na(anova$aov.tab$`Pr(>F)`[3]) | 
          nrow(tukey_df_sig)>0) {
        writeData(wb,sheet_name , sht, colNames = T, rowNames= T, startRow= st_row) 
        addStyle(wb, 1, style = titStyle, cols=1:10, rows=st_row:st_row, gridExpand = TRUE)
        writeData(wb,sheet_name , anova$aov.tab, colNames = T, rowNames= T, startRow= st_row+1) 
        writeData(wb,sheet_name , anova2$aov.tab, colNames = T, rowNames= T, startRow= st_row+6) 
        writeData(wb,sheet_name , tukey_df, colNames= T, rowNames= T, startRow= st_row+11) 
        writeData(wb,sheet_name , tukey_df2, colNames= T, rowNames= T, startRow= st_row+15) 
        st_row<-st_row+20
        print(ggplot(physeq_data_p, aes(x = get(field), y = Abundance, fill= get(field2))) +
                geom_point(size=2, position=position_jitter(width=0.2), alpha=0.5) +
                xlab(eval(field))+
                labs(fill = eval(field2))+
                theme(axis.text.x = element_text(angle = 30)) +
                ggtitle(paste(sht,"Abundance of",lvl,"per taxa and",field,"boxplot",
                              "\n no of OTUs:", nrow(physeq_data_p),
                              "\navova",eval(field),anova$aov.tab$`Pr(>F)`[1],
                              eval(field2),anova$aov.tab$`Pr(>F)`[2],
                              "\n",eval(field),":",eval(field2),anova$aov.tab$`Pr(>F)`[3],
                              "\n sig tukey:",paste(rownames(tukey_df_sig),collapse = ";"))) +
                geom_boxplot())
        dev.copy(jpeg,filename=paste0(res_dir,"/taxa_abundance_",field,"_boxplot_",sht,"_",lvl,".jpg"), 
                 width=600, height=800)
        dev.off()
      }
      if (!is.na(anova2$aov.tab$`Pr(>F)`[1]) |
          !is.na(anova2$aov.tab$`Pr(>F)`[2]) |
          !is.na(anova2$aov.tab$`Pr(>F)`[3]) |
          nrow(tukey_df_sig)>0) {
        print(ggplot(physeq_data_p, aes(x = get(field2), y = Abundance, fill= get(field))) +
                geom_point(size=2, position=position_jitter(width=0.2), alpha=0.5) +
                xlab(eval(field2))+
                labs(fill = eval(field))+
                theme(axis.text.x = element_text(angle = 30)) +
                ggtitle(paste(sht,"Abundance of",lvl,"per taxa and ",field2,"boxplot",
                              "\n no of OTUs:", nrow(physeq_data_p),
                              "\navova2",eval(field2),anova2$aov.tab$`Pr(>F)`[1],
                              eval(field),anova2$aov.tab$`Pr(>F)`[2],
                              "\n",eval(field2),":",eval(field),anova2$aov.tab$`Pr(>F)`[3],
                              "\n sig tukey2:",paste(rownames(tukey_df_sig2),collapse = ";"))) +
                geom_boxplot())
        dev.copy(jpeg,filename=paste0(res_dir,"/taxa_abundance_",field2,"_boxplot_",sht,"_",lvl,".jpg"), 
                 width=600, height=800)
        dev.off()
      }
      
    }
  }
  setColWidths(wb, sheet_name, cols = 1:8, widths = "auto")
  openxlsx::saveWorkbook(wb, file=paste0(res_dir,"/taxa_abundance_stats_",lvl,"_",set,classif, ".xlsx"), 
                         overwrite = TRUE)
  
  
}

sig_boxplot <- function(count_df,res_sig_list,res_dir=result_dir,stats=T,field,field2,group_field, 
                        pair_name,spec_colors,m16S=meta_16S,output_format="pdf",plot_all=T,
                        data_type="Relative",text_h=2,level="short_title",pnt_size=3){
#browser()
  if (missing(group_field)){
    taxa_count_long<- count_df %>%    #    
      rownames_to_column("taxa") %>%
      dplyr::filter(taxa %in% res_sig_list) %>%
      pivot_longer(col=-taxa,names_to = "sample_name", values_to = "Abundance")    %>%
      left_join(dplyr::select(rownames_to_column(m16S,"sample_name"),
                              c(sample_name,eval(field),eval(field2))),
                by="sample_name")
  } else {
#browser()
    taxa_count_long<- count_df %>%    #    
      rownames_to_column("taxa") %>%
      dplyr::filter(taxa %in% res_sig_list) %>%
      pivot_longer(col=-taxa,names_to = "sample_name", values_to = "Abundance")    %>%
      left_join(dplyr::select(rownames_to_column(m16S,"sample_name"),
                              c(sample_name,eval(group_field),eval(field),eval(field2))),
                by="sample_name")
  }
#browser()  
  
  sig_taxa_abund<- taxa_count_long %>%
                    dplyr::rename("new_name"=eval(field))
  
  pairs<-combn(unique(as.character(sig_taxa_abund$new_name)), 2)
  #pairs<- data.table::CJ(sig_taxa_abund$new_name, sig_taxa_abund$taxa, unique = TRUE)
  pairs_list <- as.list(as.data.frame(pairs))
  print(paste("length(pairs_list):",length(pairs_list)))
#browser()  
  
  
  if (output_format=="pdf") pdf(file=paste0(res_dir,"/Boxplots_",data_type,"_Abundance_",pair_name,"_",proj,"_",set, ".pdf"))  #,width=7, height=7
  
  if (!missing(group_field)) {
    for (grp in unique(m16S[[group_field]])){
      #grp<-unique(meta_16S[[group_field]])[2]
      sig_taxa_abund_p<- subset(sig_taxa_abund,sig_taxa_abund[[group_field]]==grp)
      if (nrow(sig_taxa_abund_p)>0){
#         if (length(pairs_list)>1){
# browser()
#           stat_res<-sig_taxa_abund_p %>%
#                       group_by(taxa)  %>%
#                       dunn_test(Abundance ~ new_name) %>%
#                       add_xy_position(x = "taxa") #%>%
#         } else {
#           stat_res<-sig_taxa_abund_p %>%
#                     group_by(taxa)  %>%
#                     wilcox_test(Abundance ~ new_name) %>%
#                     adjust_pvalue(method = "bonferroni") %>%
#                     add_significance("p.adj") %>%
#                     add_xy_position(x = "taxa") #%>%
#         }
        g_all<-ggplot(sig_taxa_abund_p, aes(x = taxa, y = Abundance)) +
          ggtitle(paste("Significant",data_type,"Abundance Boxplot","\n",pair_name,grp))+ 
          geom_boxplot(alpha = 0.10,lwd=1, aes(fill= new_name,color= new_name)) + 
          {if(!missing(spec_colors)) scale_color_manual(values = spec_colors)} +
          theme(panel.grid.major = element_blank(), 
                panel.grid.minor = element_blank(),
                panel.background = element_blank(), 
                axis.text.x = element_text(angle = 90),
                axis.line = element_line(colour = "black"),
                text=element_text(size = 8,  family="sans") )+ 
          coord_flip()
        # if (stats) g_all<- g_all+ stat_pvalue_manual(stat_res, label = "p.adj.signif", hide.ns =T, tip.length=0,
        #                                              coord.flip = TRUE)
        print(g_all)
      }
    } #for
  } else {
#brower()
    if (length(res_sig_list)>1){
      if (length(pairs_list)>1 & stats){
        stat_res<-sig_taxa_abund %>%
                  group_by(taxa)  %>%
                  dunn_test(Abundance ~ new_name) %>%
                  # adjust_pvalue(method = "bonferroni") %>%
                  # add_significance("p.adj") %>%
                  add_xy_position(x = "taxa") #%>%
        tit<-"dunn_test"
  #browser()
      } else {
        stat_res<-sig_taxa_abund %>%
                  group_by(taxa)  %>%
                  dunn_test(Abundance ~ new_name) %>%
                  #wilcox_test(Abundance ~ new_name) %>%
                  adjust_pvalue(method = "bonferroni") %>%
                  add_significance("p.adj") %>%
                  add_xy_position(x = "taxa") #%>%
        tit<-"dunn_test"
      }
      print(stat_res)
#browser()     
      if (plot_all){
        g_all<-ggplot(sig_taxa_abund, aes(x = taxa, y = Abundance)) +
          ggtitle(paste("Significant",data_type,"Abundance Boxplot","\n",pair_name))+ 
          geom_boxplot(alpha = 0.10,lwd=0.5, aes(fill= new_name,color= new_name),outlier.shape = NA) + 
          labs(y="Relative Abundance (%)") +
          {if(!missing(spec_colors)) scale_color_manual(values = spec_colors)} +
          theme(panel.grid.major = element_blank(), 
                panel.grid.minor = element_blank(),
                panel.background = element_blank(), 
                axis.text.x = element_text(angle = 90),
                axis.line = element_line(colour = "black"),
                text=element_text(size = 8,  family="sans") )+ 
          coord_flip()
        if (stats) g_all<- g_all+ stat_pvalue_manual(stat_res, label = "p.adj.signif", hide.ns =T, tip.length=0,
                                                     coord.flip = TRUE)
        print(g_all)
    }
    
  #browser()
      if (stats) {
        stat_res_sig<- stat_res %>%
          dplyr::filter(p.adj<=0.05) %>%
          dplyr::select(taxa,group1,group2,n1,n2,p,p.adj)
        tp<-table_pdf(stat_res_sig,tit=tit)
      }
    }
  }
  # if (nrow(stat_res_sig)>40){
  #   grid.arrange(tableGrob(stat_res_sig[1:(nrow(stat_res_sig)/2),],theme=ttheme_minimal(base_size = 4)))
  #   grid.arrange(tableGrob(stat_res_sig[(as.integer(nrow(stat_res_sig)/2)+1):nrow(stat_res_sig),],theme=ttheme_minimal(base_size = 4)))
  # } else if (nrow(stat_res_sig)>0){
  #   grid.arrange(tableGrob(stat_res_sig,theme=ttheme_minimal(base_size = 6)))
  # }
  
#browser()  
  
  for (st in unique(sig_taxa_abund$taxa)){
    #st<- " f__Fusobacteriaceae"
    sig_taxa_abund_p<-subset(sig_taxa_abund,taxa==st) 
    #    if (str_detect(st,"Fuso")) {
    #      print(paste("filter quantile:",quantile(sig_taxa_abund_p$Abundance,0.999)))
    #      sig_taxa_abund_p_filtered<- sig_taxa_abund_p %>%
    #        dplyr::filter(Abundance<quantile(Abundance,0.999))
    #    }
    print(st) 
    if (nrow(sig_taxa_abund_p)>0){
#browser()
      if (length(unique(sig_taxa_abund$new_name))>2){
        sig_taxa_abund_p_4dunn<-sig_taxa_abund_p #%>%
        if (!missing(group_field)) {
          #browser()
          stat_res<-sig_taxa_abund_p_4dunn %>% 
                    group_by(eval(group_field)) %>% 
                    dunn_test(Abundance ~ new_name)#%>% 
          stat_res_k<-sig_taxa_abund_p_4dunn %>% 
                    group_by(eval(group_field)) %>% 
                    kruskal_test(Abundance ~ new_name)
        } else {
          stat_res_k<-sig_taxa_abund_p_4dunn %>%
                   kruskal_test(Abundance ~ new_name)
          stat_res<-sig_taxa_abund_p_4dunn %>% 
                  dunn_test(Abundance ~ new_name)#%>% 
        }
        #print(stat_res)
        text<-tableGrob(stat_res,theme=ttheme_default(base_size = 5) )
        text$heights <- rep(unit(1/nrow(text), "null"), nrow(text))
        tit=textGrob(paste("Statistics kruskal:",stat_res_k$p,"dunn_test:"))
      }
#browser()
      g1<-ggplot(sig_taxa_abund_p, aes(x = new_name, y = Abundance, color= new_name)) +
        labs(x=field, y="Relative Abundance (%)", color=field)+
        ggtitle(paste(st,data_type,"Abundance boxplot",
                      ifelse(!stats,"",ifelse(length(levels(sig_taxa_abund_p$new_name))>2,"stats:wilcox.test","stats:kruskal.test")),
                      "\n",pair_name))+ 
        geom_boxplot(lwd=0.5, outlier.shape = NA) + #color = "grey",
        geom_point(size= pnt_size, 
                   position=position_jitter(width=0.2)) + #shape= 21,
        {if(!missing(spec_colors)) scale_color_manual(values = spec_colors)} +
        theme(panel.grid.major = element_blank(), 
              panel.grid.minor = element_blank(),
              panel.background = element_blank(), 
              # axis.text.x = element_text(angle = 30),
              axis.line = element_line(colour = "black"),
              text=element_text(size = 12,  family="sans") )+ 
        {if (!missing(group_field)) facet_grid(.~get(group_field),scales = "fixed", space = "free")} +
        {if (stats & length(levels(sig_taxa_abund_p$new_name))==2) stat_compare_means(method = "kruskal.test",size=2)} + 
        {if (stats & length(levels(sig_taxa_abund_p$new_name))>2) stat_compare_means(comparisons = pairs_list,label = "p.signif",
                                                                                     method = "wilcox.test",p.adjust.method = "bonferroni", hide.ns =T, tip.length=0)} # + Add pairwise comparisons p-value
      # stat_compare_means(method = "kruskal.test",paired = TRUE,size=2)
#browser()   
      if (length(unique(sig_taxa_abund$new_name))>2){
        
        plot(arrangeGrob(g1, tit, text,  ncol = 1, heights=c(6,0.5,text_h)))
        # grid.arrange(tableGrob(sig_taxa_abund_p))
      }else{
        plot(arrangeGrob(g1))
      }
      
    } 
    
    
  }
  if (output_format=="pdf") dev.off()
}

 taxa_sig_pairs<- function(taxa_thresh, taxa_all_results,set_name,element="taxa", cols_dend = TRUE, 
                           an_colors,special_color,bp_order,LFC_barplot=T,all_barplot=T, plot_pv=T,
                           res_dir=result_dir, anno_df=taxa_all_df, pv_labeled=T, rowname_len=70,
                           use_padj=FALSE, pthresh=0.05,corr=F,font_size=8, out_format="jpeg", short_output=F){
   #taxa_all_results<- taxa_MW_res # func_all_res  taxa_all_res metabo_all_res
   #taxa_thresh<-1  #0.58
   #res_dir<- result_dir  #result_dir #result_dir_all MB_result_dir
   #set_name<-"" #
   #corr=F ; use_padj=T
   #anno_df=meta_4MA  #taxa_all_df
   if (!short_output) print(set_name)
   wb <- openxlsx::createWorkbook()
   res_sig_all<-rownames_to_column(as.data.frame(get(taxa_all_results[1])[0,]),"short_title")
   first_res<-TRUE
   for (res in taxa_all_results){
     #res <- taxa_all_results[2]
     if (!short_output) print(res)

     sheet_name<-str_replace(res,"taxa_","")
     if (str_length(sheet_name)>31) sheet_name<-str_replace(sheet_name,"y_","")
     if (str_length(sheet_name)>31) sheet_name<-str_sub(sheet_name,1,31)
     addWorksheet(wb,sheet_name )
     
     compared<-str_split_fixed(res,"\\_",4)[,1]
     res_name<-str_split_fixed(res,"\\_",2)[,2]
#browser()     
     res_df<- as.data.frame(get(res))
     if (!short_output) print(paste("nrow res_df:",nrow(res_df)))
     if (plot_pv){
     ## Volcano plot t
       pv <- plot_volcano(res_df,paste0(set,"_16s_",compared,"_pairs_sig",
                                        ifelse(use_padj,paste0("PADJ.",pthresh), 
                                               paste0("pval_lfc.",taxa_thresh)),"_", res,"_",set_name),
                          res_dir = res_dir,
                          labeled = pv_labeled,
                          element = element,
                          output_format=out_format,
                          lfc=taxa_thresh,
                          padj_thresh=pthresh)
     }
     
     ## calculate sig taxa
     res_padj <- subset(res_df, padj <= pthresh)
     if (!short_output) print(paste(res, "padj sig:",nrow(res_padj)))
     
     res_pval <- subset(res_df, pvalue <= pthresh)
     if (!short_output) print(paste(res, "pvalue <=:",pthresh,nrow(res_pval)))
     
     if (!use_padj){
       res_sig <- as.data.frame(subset(res_pval, abs(log2FoldChange) >=taxa_thresh))
       if (!short_output) print(paste(compared,"pval sig",taxa_thresh,nrow(res_sig)))
       assign(paste0(res,"_pval_sig"),res_sig, envir = .GlobalEnv)
     } else {
       res_sig <- as.data.frame(subset(res_padj, abs(log2FoldChange) >=taxa_thresh))
       if (!short_output) print(paste(compared,"padj sig",taxa_thresh,nrow(res_sig)))
       assign(paste0(res,"_padj_sig"),res_sig, envir = .GlobalEnv)
     }
#browser()
     #print(res_sig)
     writeData(wb,sheet_name , res_sig, colNames = TRUE, rowNames = TRUE) 
     
     assign(paste0("res_",set_name,"_",res_name),res_sig, envir = .GlobalEnv)
    #4correlation
     if (corr){
       taxa_abund<-get(paste0(element,"_count_",set_name))
       if (!use_padj){
         taxa_abund_all<- taxa_abund  %>%
           rownames_to_column("short_title") %>%
           dplyr::filter(short_title %in% rownames(res_pval)) %>%
           column_to_rownames("short_title") 
       } else {
         taxa_abund_all<- taxa_abund  %>%
           rownames_to_column("short_title") %>%
           dplyr::filter(short_title %in% rownames(res_padj)) %>%
           column_to_rownames("short_title") 
       }
       #print(rownames(res_sig))
       taxa_abund_sig<- taxa_abund  %>%
                         rownames_to_column("short_title") %>%
                         dplyr::filter(short_title %in% rownames(res_sig)) %>%
                         column_to_rownames("short_title") 
       #print(rownames(taxa_abund_sig))
       
       #4corr
       res_sig_taxa<-res_sig
       res_taxa_pval<-res_pval
#browser()
       save(res_sig_taxa, taxa_abund_sig, res_taxa_pval,taxa_abund_all,
            file=paste0(metadata_dir,"/sig.",taxa_thresh,"_abund_4corr_",res,"_",set,"_",set_name, ".Rdata"))
       
     } 
     tit<-paste("barplot",compared,"sig",ifelse(use_padj,"PADJ", paste0("lfc.",taxa_thresh)),res,set,set_name)
     if (nrow(res_sig)>0){
         res_sig_all<-res_sig  %>%
               rownames_to_column("short_title") %>%
               mutate(pair=res) %>%
               bind_rows(res_sig_all)
#browser()
         if (nrow(res_sig)>1 & LFC_barplot){
       # LFC barplot
      		 g<-ggplot(rownames_to_column(res_sig), aes(fill=pvalue, 
      															y=log2FoldChange, 
      															x=reorder(substr(rowname,1,60),log2FoldChange)))+
      				 geom_bar(position="dodge", stat="identity")+
      				 #theme_grey(base_size = 20) +
      				 ggtitle(tit)+
      				 theme(axis.title.y= element_blank())+#, 
      				 coord_flip()
      		 if (out_format=="jpeg"){
      		   g<-g+
      		     theme(panel.grid.major = element_blank(), 
      		           panel.grid.minor = element_blank(),
      		           panel.background = element_blank(), 
      		           axis.line = element_line(colour = "black"),
      		           text=element_text(size = 20,  family="sans") )
      		     
      		   print(g)
      		   dev.copy(jpeg,filename=paste0(res_dir,"/barplot_",compared,"_sig_",
      										 ifelse(use_padj,"PADJ", paste0("lfc.",taxa_thresh)),"_",res,"_",set,"_",set_name, ".jpg"), 
      					width=max((max(res_sig$log2FoldChange)-min(res_sig$log2FoldChange))*20,str_length(tit)*25), 
      					height=nrow(res_sig)*45+50)
      		   dev.off()
      		 } else if (out_format=="pdf"){
      		   g<-g+
      		     theme(panel.grid.major = element_blank(), 
      		           panel.grid.minor = element_blank(),
      		           panel.background = element_blank(), 
      		           axis.line = element_line(colour = "black"),
      		           text=element_text(size = 18,  family="sans") )
      		   print(g)
      		   ggsave(paste0(res_dir,"/barplot_",compared,"_sig_",
      						 ifelse(use_padj,"PADJ", paste0("lfc.",taxa_thresh)),"_",res,"_",set,".pdf"), width = 5, height = 5, device = "pdf", dpi=300, units="in")
      		   dev.off()
      		 } else {
      		   g<-g+
      		     theme(panel.grid.major = element_blank(), 
      		           panel.grid.minor = element_blank(),
      		           panel.background = element_blank(), 
      		           axis.line = element_line(colour = "black"),
      		           text=element_text(size = font_size,  family="sans") )
      		   print(g)
      		 }
         }
		 #openxlsx::write.xlsx(res_sig,paste0(res_dir,"/",compared,"_sig_lfc.",taxa_thresh,"_",res,"_",set,".xlsx"),
		#                      rowNames = TRUE, colNames = TRUE)
     
     #plot sample-taxa heatmap
     # only the relevant cols
     if (!short_output) print(str_split(res,"_",simplify = T))
     ana_type<-ncol(str_split(res,"_",simplify = T))
#browser()
     if ((ana_type==4 | ana_type==6) & !str_detect(res,"_y_")& !str_detect(res,"_scaled")){
       vst_pair_df<-vst_all
       cond_df<-anno_df
     }else{
       vst_pair_df<-get (str_replace(res,compared,"vst"))
       # only the relevant rows
       cond_df<-get (str_replace(res,compared,"anno")) %>%
                  arrange_all()
     }
#browser()
     dds_count_sig<-dds_count_df[rownames(res_sig),] #%>%
     assign(paste0("dds_count_sig_",set_name),dds_count_sig, envir = .GlobalEnv)
     
     # heatmap for all base on the first result
#browser()       
     vst_sig<-vst_pair_df[rownames(res_sig),] %>%
                rownames_to_column() %>%
                # mutate(rowname=ifelse (rowname_len !="max",substr(rowname,1,rowname_len),rowname)) %>%
                relocate(rownames(cond_df)) %>%
                column_to_rownames()
     if (!short_output) print(paste("vst_sig",nrow(vst_sig)))
    if (nrow(vst_sig)>1 & !short_output){
       
         plot_genes_heatmap(vst_sig,
                            cond_df,
                            dbg=if (short_output) FALSE else TRUE,
                            y_name=compared,
                            paste0(set,"_",res,"_sig",ifelse(use_padj,"PADJ", paste0("lfc.",taxa_thresh)),"_",set_name), scl="row",
                            res_dir=res_dir, 
                            out_format=out_format,
                            cols_dend=cols_dend,
                            an_colors = an_colors,
                            special_color=special_color,
                            font_size = font_size,
                            plot_title = paste(set,res,"sig",ifelse(use_padj,"PADJ", paste0("lfc.",taxa_thresh)),set_name,"\nscale=row") )
       }
     }
#browser()       
     
    if (str_detect(res,"_scaled|\\.cont") & nrow(res_sig)>0){
      scaled_col<-str_remove(res,"taxa_.+?_")
      count_sig_long<- dds_count_sig %>%
                      rownames_to_column("taxa") %>%
                      pivot_longer(cols = -taxa,names_to = "sample_name", values_to = "abund")  %>%
                      pivot_wider(names_from = "taxa", values_from = "abund")  %>%
                      left_join(rownames_to_column(subset(md,select=eval(scaled_col)), "sample_name"),by= "sample_name")  %>%
                      column_to_rownames("sample_name")  #%>%
                      #t()
      corr_bacteria <- corr.test(count_sig_long, method="spearman")
      corrplot::corrplot(corr_bacteria$r, p.mat = corr_bacteria$p, method = "square",type = "lower",
                         insig = "label_sig", sig.level = c(.001, .01, .05),
                         title=paste("\nSig Bacteria-",res, "correlation", set,proj),
                         pch.cex = .9, pch.col = "white", order="hclust")
      
    }
       
    openxlsx::saveWorkbook(wb, file=paste0(res_dir,"/",proj,"_",compared,"_sig_",ifelse(use_padj,paste0("PADJ_lfc.",taxa_thresh), paste0("PVAL_lfc.",taxa_thresh)),
                                           "_",set,if (element=="taxa") classif,"_",set_name, ".xlsx"), 
                           overwrite = TRUE)
    
   } 
#browser()       
   #print("after loop")
   #print(res_sig_all)
   if (nrow(res_sig_all)>1 & ncol(res_sig_all)>1 & all_barplot & length(unique(res_sig_all$pair))>1){
     # if (!missing(bp_order)) res_sig_all$pair<- factor(res_sig_all$pair,levels = bp_order)
     hline <- data.frame(pair = unique(res_sig_all$pair), xp = rep(0,length(unique(res_sig_all$pair))))
     
     g_all<-ggplot(res_sig_all, aes(fill=pvalue, 
                                  y=log2FoldChange, 
                                  x=reorder(substr(short_title,1,60),log2FoldChange)))+
           geom_bar(position="dodge", stat="identity")+
           labs(x=paste("Significant",ifelse(element=="metabo","Metabolites",element)))+
           ggtitle(paste("barplot",compared,"sig",ifelse(use_padj,"PADJ", paste("lfc.",taxa_thresh)),set,set_name))+
           geom_hline(aes(yintercept = xp), data = hline,  colour = "red", size=1)+
           theme(axis.title.y= element_blank(),
                 axis.text=element_text(size = font_size,  family="sans") )+#, 
           {if(!missing(bp_order)) {
             facet_wrap(~factor(pair, levels = bp_order), 
                                              strip.position = "top",drop=FALSE, 
                                              scales = "fixed",nrow = 1) 
             } else {
               facet_wrap(~pair,strip.position = "top", drop=FALSE, scales = "fixed",nrow = 1)
             }
           } +
           coord_flip()
     
#browser() 
     if (out_format=="jpeg"){
       
       w<-max(max((max(res_sig_all$log2FoldChange)-min(res_sig_all$log2FoldChange))*
                    length(unique(res_sig_all$pair)) *20,str_length(tit)*25),
              max(str_length(unique(substr(res_sig_all$short_title,1,60)))))
       print(w)
       print(g_all+ theme_grey(base_size = 12) )
       dev.copy(jpeg,filename=paste0(res_dir,"/barplot_",compared,"_sig_",
                                     ifelse(use_padj,"PADJ", paste0("lfc.",taxa_thresh)),"_",set,"_",set_name, ".jpg"), 
                width=w, 
                height=nrow(res_sig_all)*30)
       dev.off()
     } else {
       print(g_all+ theme_grey(base_size = 8) )
     }
   }
  

   return(res_sig_all)
 }
 
 
 genus_contaminantion_check<- function(genus2check){
   contaminant_Alpha_proteobacteria<-c("Afipia", "Aquabacterium", "Asticcacaulis", "Aurantimonas", "Beijerinckia", "Bosea", "Bradyrhizobium", "Brevundimonas" , "Caulobacter", "Craurococcus", "Devosia", "Hoeflea", "Mesorhizobium", "Methylobacterium", "Novosphingobium", "Ochrobactrum", "Paracoccus", "Pedomicrobium", "Phyllobacterium", "Rhizobium", "Roseomonas", "Sphingobium", "Sphingomonas", "Sphingopyxis")
   contaminant_Beta_proteobacteria<-c("Acidovorax", "Azoarcus", "Azospira", "Burkholderia", "Comamonas", "Cupriavidus", "Curvibacter", "Delftia", "Duganella", "Herbaspirillum", "Janthinobacterium", "Kingella", "Leptothrix", "Limnobacter", "Massilia", "Methylophilus", "Methyloversatilis", "Oxalobacter", "Pelomonas", "Polaromonas", "Ralstonia", "Schlegelella", "Sulfuritalea", "Undibacterium", "Variovorax")
   contaminant_Gamma_proteobacteria<-c("Acinetobacter" , "Enhydrobacter", "Enterobacter", "Escherichia", "Nevskia", "Pseudomonas", "Pseudoxanthomonas", "Psychrobacter", "Stenotrophomonas", "Xanthomonas")
   contaminant_Actinobacteria<-c("Aeromicrobium", "Arthrobacter", "Beutenbergia", "Brevibacterium", "Corynebacterium", "Curtobacterium", "Dietzia", "Geodermatophilus", "Janibacter", "Kocuria", "Microbacterium", "Micrococcus", "Microlunatus", "Patulibacter", "Propionibacterium", "Rhodococcus", "Tsukamurella")
   contaminant_other<-c("Abiotrophia", "Bacillus", "Brevibacillus", "Brochothrix", "Facklamia", "Paenibacillus", "Streptococcus","Chryseobacterium", "Dyadobacter", "Flavobacterium", "Hydrotalea", "Niastella", "Olivibacter", "Pedobacter", "Wautersiella", "Deinococcus","Acidobacteria")
   if(any(genus2check$Genus %in% contaminant_Alpha_proteobacteria, na.rm = FALSE)){
     ifelse(genus2check$Genus %in% contaminant_Alpha_proteobacteria,taxa_16S_df$Genus,NA)
   } else print("contaminant_Alpha_proteobacteria not contaminanted")
   if(any(genus2check$Genus %in% contaminant_Beta_proteobacteria, na.rm = FALSE)){
     ifelse(genus2check$Genus %in% contaminant_Beta_proteobacteria,taxa_16S_df$Genus,NA)
   } else print("contaminant_Beta_proteobacteria not contaminanted")
   if(any(genus2check$Genus %in% contaminant_Gamma_proteobacteria, na.rm = FALSE)){
     ifelse(genus2check$Genus %in% contaminant_Gamma_proteobacteria,taxa_16S_df$Genus,NA)
   }else print("contaminant_Gamma_proteobacteria not contaminanted")
   if(any(genus2check$Genus %in% contaminant_Actinobacteria, na.rm = FALSE)){
     ifelse(genus2check$Genus %in% contaminant_Actinobacteria,taxa_16S_df$Genus,NA)
   }else print("contaminant_Actinobacteria not contaminanted")
   if(any(genus2check$Genus %in% contaminant_other, na.rm = FALSE)){
     ifelse(genus2check$Genus %in% contaminant_other,taxa_16S_df$Genus,NA)
   }else print("contaminant_other not contaminanted")
 }
 
 Bray_dist<- function(count,md,group,set_name="", res_dir = result_dir,plot_jpg=T,mtd="bray",directions=T,
                      spec_colors, label=T, ellipse=T, meta_pair, wb_in){
#browser()
   md_df<-data.frame(V1=md) #%>%
     # dplyr::filter(!is.na(V1))
   
   ## Bray-Curtis distances between samples
   dis <- vegdist(t(count),method = mtd, na.rm =T)
   assign("dis", dis, envir = .GlobalEnv)
   # heatmap(as.matrix(dis))
   ## bray dissimilarities is the default, noshare triggers stepacross
   dis_nmds <- metaMDS(dis)

   
#browser()
  #data.scores <- as.data.frame(scores(dis_nmds))  #Using the scores function from vegan to extract the site scores and convert to a data.frame
   
   # col_palette <- palette()[md]
   # col_palette <- hue_pal()(length(unique(md)))
   # plot(dis_nmds, type = "n", main=paste(mtd, "distances on MDS",set_name, set))
   # points(dis_nmds, col = col_palette, pch = 16)
   # if (label) text(dis_nmds$points[ ,1], dis_nmds$points[ ,2], labels=rownames(dis_nmds$points), col = col_palette, 
   #      pos=1, cex = 0.5) 
   # legend("topright", legend=unique(md), col=col_palette, pch = 16, title = group)
   # ordiellipse(ord = dis_nmds, groups = md, kind = "se", conf = .95, col = unique(col_palette), label=TRUE) 
#browser()
   
   anova_dist<-adonis2(dis ~V1 ,data=md_df, method="bray", na.action = na.omit)
   if (!missing(meta_pair)){
     meta_pair_new<-meta_pair %>%
     dplyr::rename("new_name"=eval(group))
     
#browser()
     
    anova_pair <- pairwiseAdonis::pairwise.adonis2(dis ~new_name,data=meta_pair_new, na.action = na.omit)
    anova_pair_df<-as.data.frame(anova_pair,check.names=F) %>%
      dplyr::select(ends_with("Pr(>F)")) %>%
      rename_with(~str_remove_all(.,"\\.Pr\\(\\>F\\)")) %>%
      dplyr::filter(row_number()==1) %>%
      t() %>%
      as.data.frame() %>%
      rownames_to_column("pairwise.adonis2") %>%
      mutate(pairwise.adonis2=str_replace(pairwise.adonis2,"_vs_","-")) #%>%
#browser()
    # anova_pair_sig_df<-anova_pair_df[anova_pair_df[ ,2]<0.1,] %>%
    anova_pair_sig_df<-anova_pair_df %>%
                    dplyr::filter(new_name<0.1) %>%
                    column_to_rownames("pairwise.adonis2")
    
    plot_data<- as.data.frame(dis_nmds$points) %>%
      rownames_to_column("sample_name2") %>%
      left_join(rownames_to_column(meta_pair,"sample_name2") ,by="sample_name2" ) %>%
      column_to_rownames("sample_name2")
#browser()
    print(ggplot(plot_data, aes(x=MDS1, y=MDS2))+ #sets up the plot
            geom_point(aes(MDS1, MDS2, colour = md))+ #adds site points to plot
            coord_fixed()+
            theme_classic()+ 
            {if(!missing(spec_colors)) scale_color_manual(values = spec_colors)} +
            theme(panel.background = element_rect(fill = NA, colour = "black", size = 1, linetype = "solid"))+
            theme(legend.position = "right", legend.text = element_text(size = 8), legend.title = element_text(size = 12),
                  axis.text = element_text(size = 10)) +# add legend at right of plot
            {if(label) ggrepel::geom_text_repel(aes( label = rownames(plot_data)), size=1.5)}+
            labs(title = paste(set,set_name, "Distances-bray"),colour = group) )
    if(directions){
      dis.envfit <- envfit(dis_nmds, meta_pair, permutations = 99, na.rm = TRUE)
      env.scores <- data.frame(dis.envfit$factors$centroids,var=dis.envfit$factors$var.id) %>%
                      rownames_to_column()
#browser()
      if (length(dis.envfit$factors$pvals)>0){
        env.pval <- data.frame(pval=dis.envfit$factors$pvals) %>%
                      rownames_to_column("var")
        env.scores <- merge(env.scores, env.pval) # add pvalues to dataframe
        sig.env.scrs <- subset(env.scores, pval<=0.05) #subset data to show variables significant at 0.05

        if (nrow(sig.env.scrs)>0) {
          print(ggplot(plot_data, aes(x=MDS1, y=MDS2))+ #sets up the plot
                  geom_point(aes(MDS1, MDS2, colour = md))+ #adds site points to plot
                  coord_fixed()+
                  theme_classic()+ 
                  theme(panel.background = element_rect(fill = NA, colour = "black", size = 1, linetype = "solid"))+
                  theme(legend.position = "right", legend.text = element_text(size = 8), legend.title = element_text(size = 12),
                        axis.text = element_text(size = 10)) +# add legend at right of plot
                  labs(title = paste(set,set_name, "Distances-bray"),colour = group) + 
                  geom_segment(data = sig.env.scrs, aes(x = 0, xend=NMDS1, y=0, yend=NMDS2), arrow = arrow(length = unit(0.25, "cm")), colour = "grey10", lwd=0.3) + #add vector arrows of significant env variables
                  ggrepel::geom_text_repel(data = sig.env.scrs, aes(x=NMDS1, y=NMDS2, label = rowname), cex = 2, direction = "both", segment.size = 0.25))#+ #add labels for env variables
        }
      }
    }
   }
   mod <- betadisper(dis, md, type = "centroid") #, type = "median", bias.adjust=TRUE [[1]]
   #print(mod)
#browser()
   #Implements a permutation-based test of multivariate homogeneity of group dispersions (variances) for the results of a call to betadisper
   prm<-permutest(mod, pairwise = TRUE)
   #matrix of coefficients of the linear model, with rows representing sources of variation and columns representing species;  
   #These are NOT available if you supply the distance matrix in the formula, rather than the site x species matrix
   # coefficients(anova_dist)
   
   if (!missing(wb_in)){
     posStyle <- createStyle(fontColour = "#006100", fgFill = "#C6EFCE")
     s <- createStyle(numFmt = "$ #,##0.00")
     #m<-"Observed"
     titStyle <- createStyle(fontColour = "#006100", fgFill = "#C6EFCE")
     sheet_b_anova <- paste0("Bdiv ",set_name," ",group)
     addWorksheet(wb_in, sheet_b_anova)
     writeData(wb_in,sheet_b_anova,group, startRow= 1) 
     addStyle(wb_in, sheet_b_anova, style = titStyle, cols=1:10, rows=1:1, gridExpand = TRUE)
     #browser()
     writeData(wb_in,sheet_b_anova, anova_dist, colNames = T, rowNames= T) 
     writeData(wb_in,sheet_b_anova, "permutation test of multivariate homogeneity of group dispersions (variances)", colNames = T, rowNames= T, startRow = 10) 
     writeData(wb_in,sheet_b_anova, data.frame(prm$tab), colNames = T, rowNames= T, startRow = 11) 
     writeData(wb_in,sheet_b_anova, data.frame(prm$pairwise), colNames = T, rowNames= T, startRow = 15) 
     if (!missing(meta_pair)) writeData(wb_in,sheet_b_anova, anova_pair_df, colNames = T, rowNames= T, startCol = 10) 
   }
   ## Calculate multivariate dispersions. Both permdisp and betadisper accept a single "grouping" factor 
   
   # tests if centroid distances are significantly different from each other
   anova_mod<-anova(mod)
   ## Permutation test for F
   #densityplot(permustats(permutest(mod, pairwise = TRUE, permutations = 99)), scales = list(x = list(relation = "free")))
   
   ## Tukey's Honest Significant Differences
   mod_tukey <- TukeyHSD(mod)
   tukey_df<-as.data.frame(mod_tukey$group)
   tukey_df_sig<-subset(tukey_df,tukey_df$`p adj` <0.1)
   ##Differences in mean levels of the group
   #plot(mod_tukey)
#browser()
   plot(mod, ellipse = ellipse, hull = FALSE, conf = 0.90, xlab=group, cex.main=0.8,
        ylab=paste(mtd,"distance"),
        main=paste(mtd,"distances between samples",set_name,
                   "\nStat",round(anova_dist$`Pr(>F)`[1],digits = 3),
                   ifelse(!missing(meta_pair),
                          paste("\nsig pairs:",paste0(rownames(anova_pair_sig_df)," (",round(anova_pair_sig_df$new_name,digits = 2),") ", collapse = ";"))),""))
        # 90% data ellipse
  
   if (label)      text( mod$vectors[,1:2],labels = rownames(mod$vectors), cex = 0.5,pos = 1)
   if (plot_jpg){
     dev.copy(jpeg,filename=paste0(res_dir,"/",mtd,"_Dist_",group,"_",set,"_",set_name, ".jpg"))
     dev.off()
   }
#browser()
   # boxplot(mod,xlab=group, ylab=paste(mtd,"distance to centroid", col=group),
   #           las=2,cex.lab=1,cex.axis=.7,
   #         main=paste("Distances to centroid",set_name,                 
   #                    "\nAnova",round(anova_mod$`Pr(>F)`[1],digits = 2),
   #                    paste(" sig tukey:",paste0(rownames(tukey_df_sig)," (",round(tukey_df_sig$`p adj`,digits = 2),")+ ", collapse = ";")))) +
   #   {if(!missing(spec_colors)) scale_color_manual(values = spec_colors)} 
   #   
   # if (plot_jpg){
   #   dev.copy(jpeg,filename=paste0(res_dir,"/",mtd,"_Dist_",group,"_boxplot_",set,"_",set_name, ".jpg"))
   #   dev.off()
   # }
   
   #------------------------Density ridgeline plots
   #https://cran.r-project.org/web/packages/ggridges/vignettes/introduction.html
   #https://r-charts.com/distribution/ggridges/ 
   dist_2centroid<-as.data.frame(cbind(group=as.character(mod$group), 
                                       distances = mod$distances)) %>%
                    mutate(group=as.factor(group),distances=as.numeric(distances))
   print(ggplot(dist_2centroid, aes(y = distances, x = group, color=group)) +
           geom_boxplot()+
           xlab(paste(mtd,"distance to centroid")) +
           ylab(eval(group))+ 
           {if(!missing(spec_colors)) scale_color_manual(values = spec_colors)} +
           ggtitle(paste("Distances between samples",set_name,                 
                         "\nAnova",round(anova_mod$`Pr(>F)`[1],digits = 2),
                         paste(" sig tukey:",paste0(rownames(tukey_df_sig)," (",round(tukey_df_sig$`p adj`,digits = 2),") ", collapse = ";")))) +
           theme(legend.position = "none"))
   if (plot_jpg){
     dev.copy(jpeg,filename=paste0(res_dir,"/",mtd,"_Dist_",group,"_boxplot_",set,"_",set_name, ".jpg"))
     dev.off()
   }
   

  print(ggplot(dist_2centroid, aes(x = distances, y = group, fill=group)) +
           geom_density_ridges(alpha = 0.5,jittered_points = TRUE, point_alpha=1,point_shape=21)+
           theme_ridges() + 
           ylab(paste(mtd,"distance to centroid")) +
           xlab(eval(group))+ 
          {if(!missing(spec_colors)) scale_fill_manual(values = spec_colors)} +
          ggtitle(paste("Distances between samples",set_name,                 
                         "\nAnova",round(anova_mod$`Pr(>F)`[1],digits = 2),
                         paste(" sig tukey:",paste0(rownames(tukey_df_sig)," (",round(tukey_df_sig$`p adj`,digits = 2),") ", collapse = ";")))) +
           theme(legend.position = "none"))
   if (plot_jpg){
     dev.copy(jpeg,filename=paste0(res_dir,"/",mtd,"_Dist_",group,"_Ridgeline_",set,"_",set_name, ".jpg"))
     dev.off()
   }
  #------------ Samples PCA  
  count_pca <- prcomp(t(count), center = TRUE,scale. = TRUE)
  #pc_directions <- data.frame(count_pca$rotation)
  count_pca_4plot <-data.frame(count_pca$x) %>%
    rownames_to_column("sample_name2") %>%
    left_join(rownames_to_column(meta_pair,"sample_name2") ,by="sample_name2" ) #%>%
  pca.var <- round(count_pca$sdev^2/sum(count_pca$sdev^2)*100, 1)
  
  
  #Samples PCA
  print(ggplot(count_pca_4plot, aes(x=PC1, y=PC2,col= get(group))) + 
        labs(x=paste("PC1:",pca.var[1],"% variance"),
             y=paste("PC2:",pca.var[2],"% variance"),
             title='Samples PCA') + 
        {if(!missing(spec_colors)) scale_color_manual(values = spec_colors)} +
        {if(label) ggrepel::geom_text_repel(aes( label = sample_name2), size=1.5, max.overlaps=Inf)}+
        theme(aspect.ratio=1,plot.title = element_text(size = 16, face = "bold"),
                panel.grid.major = element_blank(),
                panel.grid.minor = element_blank(),
                panel.background = element_blank(),
                text = element_text(size = 16))+
        geom_point(size = 2))
  
 }

 
 
 
 
 Adiver<- function(wb_in=wb, phy_rarefied,field1,field2,field3,color_field2=TRUE,fill_field3=TRUE, plus_f3=T,
                   set_name="all",PD=TRUE,sheet_a= NULL, posthoc=TRUE,spec_colors,strata_col, uselog=T,
                   plot_jpg=T, inc_plot=T){
   
   #phy_rarefied<-physeq_rarefied
   #field1<- "Control_GDM" #RatType TimeW Condition
   #field2<- "trimester" #TimeGroup PatientID Wasp
   #field3<- "woman" #RatID batch Stage
   #color_field2=T ;fill_field3=F ; plus_f3=T
   print(field1)
   posthoc_res<-character()
   set.seed(42)
   posStyle <- createStyle(fontColour = "#006100", fgFill = "#C6EFCE")
   s <- createStyle(numFmt = "$ #,##0.00")
   st_row<-1
   #m<-"Observed"
   titStyle <- createStyle(fontColour = "#006100", fgFill = "#C6EFCE")
   #sheet_a_divers <- paste0("Adivers ",sheet_a)
#   sheet_wilcox <- "alpha_pairwise_wilcox"
   sheet_a_anova <- paste0("Adiv ",sheet_a)
   addWorksheet(wb_in, sheet_a_anova)
   
   #-------------faithPD- species richness (SR) and phylogenetic diversity (PD) ------
   lmer_a <- NULL
   
   if (PD){
     alpha_diversity_pd<-estimate_pd(phy_rarefied)
     pairwise.wilcox.test_PD <- NULL
     PD_posthoc_sig <-NULL
     lmer_PD <- NULL
#browser()     
     if(!missing(field3)) {
       if (!plus_f3){
         
         if (!missing(strata_col)) {
#browser()
           alpha_divers_pd_meta<- alpha_diversity_pd %>%
                         rownames_to_column("rowname") %>%
                         left_join(rownames_to_column(meta_16S,"rowname"),by="rowname") %>%
                         dplyr::select(rowname,PD,SR,field1=eval(field1),field2=eval(field2),
                                       field3=eval(field3),strt=eval(strata_col)) %>%
                         mutate_if(is.integer,as.factor)  %>%
                         mutate(log_PD=log(PD)) %>%
                         column_to_rownames("rowname")
#browser()     
           stat_PD_call<-paste("log(PD) ~ ",field1,"*",field2,"*",field3,"strata=",strata_col,"using lmerTest::lmer+anova")
           model_PD<- lmerTest::lmer(log_PD ~ field1*field2*field3+ (1 | strt), data=alpha_divers_pd_meta)
           lmer_PD <- as.data.frame(summary(model_PD)$coefficients)%>%
             rownames_to_column("fields") %>%
             mutate(fields=str_replace_all(fields,"field1",eval(field1))) %>%
             mutate(fields=str_replace_all(fields,"field2",eval(field2))) %>%
             mutate(fields=str_replace_all(fields,"field3",eval(field3))) #%>%
           
           stat_PD <- anova(model_PD, ddf="Kenward-Roger") %>%
             rownames_to_column("fields") %>%
             mutate(fields=str_replace_all(fields,"field1",eval(field1))) %>%
             mutate(fields=str_replace_all(fields,"field2",eval(field2))) %>%
             mutate(fields=str_replace_all(fields,"field3",eval(field3))) #%>%
             
           if (posthoc){
             #   posthoc_res<-alpha_divers_pd_meta %>% 
             #     group_by(strt) %>% 
             #     dunn_test(PD ~ comb)#%>% 
#browser()     
             ph_model<-emmeans(model_PD,~ field1 * field2* field3)   
            posthoc_res<-as.data.frame(summary(pairs(ph_model, simple = "each", combine = TRUE,adjust = "BH")))
             PD_posthoc_sig<-subset(posthoc_res,posthoc_res$p.value <=0.05)
             
           }
         } else { #no strata
#browser()
           alpha_divers_pd_meta<- alpha_diversity_pd %>%
                           rownames_to_column("rowname") %>%
                           left_join(rownames_to_column(meta_16S,"rowname"),by="rowname") %>%
                           dplyr::select(rowname,PD,SR,field1=eval(field1),field2=eval(field2),field3=eval(field3)) %>%
                           mutate_if(is.integer,as.factor)  %>%
                           mutate(log_PD=log(PD)) %>%
                           column_to_rownames("rowname")
           model_PD <- summary(lm(log_PD ~ field1*field2*field3, data=alpha_divers_pd_meta))
           stat_PD <- as.data.frame(model_PD$coefficients) %>%
             rownames_to_column("fields") %>%
             mutate(fields=str_replace_all(fields,"field1",eval(field1))) %>%
             mutate(fields=str_replace_all(fields,"field2",eval(field2))) %>%
             mutate(fields=str_replace_all(fields,"field3",eval(field3))) %>%
             dplyr::rename('Pr(>F)'='Pr(>|t|)')
           stat_PD_call<-paste("log(PD) ~ ",field1,"*",field2,"*",field3,"using lm")
           if (posthoc){
             # posthoc_res<-alpha_divers_pd_meta %>% 
             #   dunn_test(PD ~ comb)#%>% 
             anova_2way <- aov(log_PD ~ field1*field2*field3, data=alpha_divers_pd_meta) #
             tukey<- TukeyHSD(anova_2way)
             posthoc_res<-as.data.frame(rbind(tukey$field1,
                                              tukey$field2,
                                              tukey$field3,
                                              tukey$'field1:field2',
                                              tukey$'field1:field3',
                                              tukey$'field2:field3'))
             PD_posthoc_sig<-subset(posthoc_res,posthoc_res$'p adj' <=0.05)
             
           }
         }
       } else{ # if (plus_f3)
          print(paste("plus_f3",sheet_a_anova))
          stat_PD <- adonis(formula =PD ~ get(field1)*get(field2)+get(field3), data=alpha_divers_pd_meta)
          stat_PD_call<-paste("PD ~ ",field1,"*",field2,"+",field3)
          anova_2way <- aov(PD ~ get(field1)*get(field2)+get(field3), data=alpha_divers_pd_meta) #
          tukey<- TukeyHSD(anova_2way)
          tukey_df<-as.data.frame(rbind(tukey$`get(field1)`,
                                        tukey$`get(field2)`,
                                        tukey$`get(field3)`,
                                        tukey$`get(field1):get(field2)`,
                                        tukey$`get(field2):get(field3)`))
          PD_posthoc_sig<-subset(tukey_df,tukey_df$`p adj` <0.05)
          print(PD_posthoc_sig)

       } 
       print(stat_PD)
       if(color_field2) {
         if(fill_field3) {
           g<-ggplot(alpha_divers_pd_meta, aes(x=field1, y=log_PD, color=field2, fill=field3))+
             geom_boxplot()+
             scale_color_discrete(name = eval(field2)) +
            scale_fill_manual(name = field3,values = rainbow(length(levels(alpha_divers_pd_meta$field3))))
         } else{
           g<-ggplot(alpha_divers_pd_meta, aes(x=field1, y=log_PD, color=field2))+
             geom_boxplot()+
             scale_color_discrete(name = field2 )
          }
        } else{
          print("1-field1 only")
           g<-ggplot(alpha_divers_pd_meta, aes(x=field1, y=log_PD,color=field1)+
                       geom_boxplot(lwd=1)+
                       geom_point(position = position_dodge(width = 0.5), size=2)+
                       stat_compare_means(method = "kruskal.test"))
        }
       if (plus_f3){
         print(paste("plus_f3 plot",sheet_a_anova))
#browser()         
         g<- g+ggtitle(paste("FaithPD-phylogenetic diversity (PD) (log)", sheet_a,
                             "\n",stat_PD_call,
                             "\n",eval(field1), stat_PD$'Pr(>F)'[2],
                             "\n",eval(field2),stat_PD$'Pr(>F)'[3],
                             "\n",eval(field3),stat_PD$'Pr(>F)'[4],
                             "\n",eval(field1),":",eval(field2),stat_PD$'Pr(>F)'[4],
                             "\nPosthoc sig",paste(rownames(PD_posthoc_sig), collapse = ";") ))
       }else{
         g<- g+ggtitle(paste("FaithPD-phylogenetic diversity (PD) (log)", sheet_a,
                             "\n",stat_PD_call,
                             "\nStats",eval(field1), stat_PD$'Pr(>F)'[2],
                             "\n",eval(field2),stat_PD$'Pr(>F)'[3],
                             "\n",eval(field3),stat_PD$'Pr(>F)'[4],
                             "\n",eval(field1),":",eval(field2),stat_PD$'Pr(>F)'[5],
                             "\n",eval(field1),":",eval(field3),stat_PD$'Pr(>F)'[6],
                             "\n",eval(field1),":",eval(field2),":",eval(field3),stat_PD$'Pr(>F)'[7],
                             "\nPosthoc sig",paste(rownames(PD_posthoc_sig), collapse = ";")  ))
       }
       print(g+ 
               xlab(eval(field1))+
               theme_bw() + 
               theme(axis.text.x = element_text(angle = 90),
                     text = element_text(size = 12))+
               guides(color=guide_legend(title=eval(field2)))+
               guides(fill=guide_legend(title=eval(field3)))+
               #stat_compare_means(method = "wilcox.test",p.adjust.method = "bonferroni"))
               stat_compare_means(method = "kruskal.test"))
      if (plot_jpg) {
         ggsave(paste0(result_dir,"/FaithPD_PD_",eval(field1),"_boxplot_",proj,"_",set,"_",sheet_a,".jpg"), 
              width=7, height=5)
          dev.off()
       }
     } else if(!missing(field2)){
     print(paste("PD 2*field:",sheet_a_anova))
#browser()
     if (!missing(strata_col)) {
       alpha_divers_pd_meta<- alpha_diversity_pd %>%
               rownames_to_column("rowname") %>%
               left_join(rownames_to_column(meta_16S,"rowname"),by="rowname") %>%
               dplyr::select(rowname,PD,SR,field1=eval(field1),field2=eval(field2),strt=eval(strata_col)) %>%
               dplyr::filter( str_length(field1)>0  & !is.na(field1) & field1!="na" &
                                str_length(field2)>0  & !is.na(field2) & field2!="na") %>%
               mutate(log_PD=log(PD)) %>%
               mutate(comb=paste0(field1,".",field2)) %>%
               column_to_rownames("rowname")
#browser()       
       stat_PD_call<-paste("log(PD) ~ ",field1,"*",field2,"strata=",strata_col,"using lmerTest::lmer+anova")
       model_PD<- lmerTest::lmer(log_PD ~ field1*field2+ (1 | strt), data=alpha_divers_pd_meta)
       lmer_PD <- as.data.frame(summary(model_PD)$coefficients)%>%
         rownames_to_column("fields") %>%
         mutate(fields=str_replace_all(fields,"field1",eval(field1))) %>%
         mutate(fields=str_replace_all(fields,"field2",eval(field2))) #%>%

       stat_PD <- anova(model_PD, ddf="Kenward-Roger") %>%
                     rownames_to_column("fields") %>%
                     mutate(fields=str_replace_all(fields,"field1",eval(field1))) %>%
                     mutate(fields=str_replace_all(fields,"field2",eval(field2))) #%>%
         
       if (posthoc){
           #   posthoc_res<-alpha_divers_pd_meta %>% 
           #     group_by(strt) %>% 
           #     dunn_test(PD ~ comb)#%>% 
           ph_model<-emmeans(model_PD,~ field1 * field2)   
           posthoc_res<-as.data.frame(summary(pairs(ph_model, simple = "each", combine = TRUE,adjust = "BH")))#%>% 
             #rename_with(~str_remove(., 'simple.contrasts.for.field1.')) 
           PD_posthoc_sig<-subset(posthoc_res,posthoc_res$p.value <=0.05)
           
       }
     } else {
#browser()
       alpha_divers_pd_meta<- alpha_diversity_pd %>%
                 rownames_to_column("rowname") %>%
                 left_join(rownames_to_column(meta_16S,"rowname"),by="rowname") %>%
                 dplyr::select(rowname,PD,SR,field1=eval(field1),field2=eval(field2)) %>%
                 dplyr::filter(str_length(field1)>0  & !is.na(field1) & field1!="na" &
                                  str_length(field2)>0  & !is.na(field2)& field2!="na") %>%
                 mutate(comb=paste0(field1,".",field2)) %>%
                 mutate(log_PD=log(PD)) %>%
                 column_to_rownames("rowname")
       model_PD <- summary(lm(log_PD ~ field1*field2, data=alpha_divers_pd_meta))
       stat_PD <- as.data.frame(model_PD$coefficients) %>%
                     rownames_to_column("fields") %>%
                     mutate(fields=str_replace_all(fields,"field1",eval(field1))) %>%
                     mutate(fields=str_replace_all(fields,"field2",eval(field2))) %>%
                    dplyr::rename('Pr(>F)'='Pr(>|t|)')
       stat_PD_call<-paste("log(PD) ~ ",field1,"*",field2,"using lm")
       if (posthoc){
         # posthoc_res<-alpha_divers_pd_meta %>% 
         #   dunn_test(PD ~ comb)#%>% 
         anova_2way <- aov(log_PD ~ field1*field2, data=alpha_divers_pd_meta) #
         tukey<- TukeyHSD(anova_2way)
         posthoc_res<-as.data.frame(rbind(tukey$field1,
                                       tukey$field2,
                                       tukey$'field1:field2'))
         PD_posthoc_sig<-subset(posthoc_res,posthoc_res$'p adj' <=0.05)
         
       }
     }
#browser()
     print(stat_PD)
     print(stat_PD_call)
     if (!posthoc) PD_posthoc_sig<-NULL

     
     print(PD_posthoc_sig)
#browser()
     # plot_stats <- alpha_divers_pd_meta %>%
     #       group_by(field2) %>%
     #       wilcox_test(log_PD ~ field1) %>%
     #       adjust_pvalue(method = "bonferroni") %>%
     #       add_significance("p.adj") %>%
     #       add_xy_position(x = "field1", dodge = 0.8)
     
    # plot_stats <- posthoc_res %>%
    #         add_xy_position( dodge = 0.8) %>%
    #         separate(group1,into = c("g1field1","g1field2"),sep = "\\.",remove = F) %>%
    #         separate(group2,into = c("g2field1","g2field2"),sep = "\\.",remove = F)
    # plot_stats1<- subset(plot_stats,g1field2==g2field2) %>%
    #               mutate(field2=g1field2)
    # 
    # plot_stats2<- subset(plot_stats,g1field1==g2field1) %>%
    #               mutate(field1=g1field1)
    
#browser()
     if (inc_plot){
       g<-ggplot(alpha_divers_pd_meta, aes(x=field1, y=log_PD, color=field2)) +
               ggtitle(paste("FaithPD-phylogenetic diversity (PD) (log)", sheet_a, 
                             "\n",stat_PD_call,
                             "\n",field1, round(stat_PD$'Pr(>F)'[1],digits = 2),
                             "\n",field2,round(stat_PD$'Pr(>F)'[2],digits = 2),
                             "\n",field1,":",field2,round(stat_PD$'Pr(>F)'[3],digits = 2),
                             "\nPosthoc sig",paste(rownames(PD_posthoc_sig), collapse = ";") )) +  
               xlab(eval(field1))+
               geom_boxplot()+
               theme_bw() + 
               {if(!missing(spec_colors)) scale_color_manual(values = spec_colors)} +
               guides(color=guide_legend(title=eval(field2)))+
               theme(axis.text.x = element_text(angle = 90),
                     text = element_text(size = 12)) #+
               # facet_grid(.~field2, scales="free_x") +
               # stat_pvalue_manual(plot_stats1, label = "{p.adj}")
               #stat_compare_means(method = "wilcox.test",p.adjust.method = "bonferroni")
               # stat_compare_means(method = "wilcox.test",
               #                    mapping=aes(label = format.pval(..p.adj.., digits = 3)))
               #stat_compare_means(method = "kruskal.test")
       
       g2<-ggplot(alpha_divers_pd_meta, aes(x=field2, y=log_PD, color=field1)) +
             ggtitle(paste("FaithPD-phylogenetic diversity (PD) (log)", sheet_a, 
                           "\n",stat_PD_call,
                           "\n",eval(field1), round(stat_PD$'Pr(>F)'[1],digits = 2),
                           "\n",eval(field2),round(stat_PD$'Pr(>F)'[2],digits = 2),
                           "\n",eval(field1),":",eval(field2),round(stat_PD$'Pr(>F)'[3],digits = 2),
                           "\nPosthoc sig",paste(rownames(PD_posthoc_sig), collapse = ";") )) +  
             xlab(eval(field2))+
             geom_boxplot()+
             theme_bw() + 
             {if(!missing(spec_colors)) scale_color_manual(values = spec_colors)} +
             guides(color=guide_legend(title=eval(field1)))+
             theme(axis.text.x = element_text(angle = 90),
                   text = element_text(size = 12)) #+
             # stat_compare_means(method = "wilcox.test",
             #                    mapping=aes(label = format.pval(..p.adj.., digits = 3)))
             #stat_compare_means(method = "kruskal.test")
       
       print(g)
       if (plot_jpg) {
         ggsave(paste0(result_dir,"/FaithPD_PD_",eval(field1),"_",eval(field2),"_boxplot_",proj,"_",set,"_",sheet_a,".jpg"), 
                width=7, height=5)
         dev.off()
       }      
       print(g2)
       if (plot_jpg) {
         ggsave(paste0(result_dir,"/FaithPD_PD_",eval(field2),"_",eval(field1),"_boxplot_",proj,"_",set,"_",sheet_a,".jpg"), 
                width=7, height=5)
         dev.off()
       }  
     }# inc_plot
     }else{
       print(paste("2-field1 only", sheet_a_anova))
#browser()
       a_pd_meta<- alpha_diversity_pd %>%
             rownames_to_column("rowname") %>%
             left_join(rownames_to_column(meta_16S,"rowname"),by="rowname") %>%
              mutate(log_PD=log(PD)) #%>%
         if (!missing(strata_col)) {
         
          alpha_divers_pd_meta<- a_pd_meta %>%
                      dplyr::select(rowname,PD,log_PD,SR,field1=eval(field1),strt=eval(strata_col)) %>%
                      dplyr::filter( str_length(field1)>0  & !is.na(field1) ) %>%
                      column_to_rownames("rowname")
          print(field1)
          # perm <- how(nperm = 1000)
          # setBlocks(perm) <- with(alpha_divers_pd_meta, strt)
          # stat_PD <- adonis(formula =PD ~ get(field1), data=alpha_divers_pd_meta, permutations = perm)
#browser()
          if (uselog) {
            model_PD<- lmerTest::lmer(log_PD ~ field1+ (1 | strt), data=alpha_divers_pd_meta)
            stat_PD_call<-paste("log(PD) ~ ",field1,"strata=",strata_col,"using lmerTest::lmer+anova")
          }else{
            model_PD<- lmerTest::lmer(PD ~ field1+ (1 | strt), data=alpha_divers_pd_meta)
            stat_PD_call<-paste("PD ~ ",field1,"strata=",strata_col,"using lmerTest::lmer+anova")
          }
          lmer_PD <- as.data.frame(summary(model_PD)$coefficients)%>%
            rownames_to_column("fields") %>%
            mutate(fields=str_replace_all(fields,"field1",eval(field1))) #%>%
          
          stat_PD <- anova(model_PD, ddf="Kenward-Roger") %>%
                        rownames_to_column("fields") %>%
                        mutate(fields=str_replace_all(fields,"field1",eval(field1))) #%>%


          if (posthoc){
            # posthoc_res<-alpha_divers_pd_meta %>% 
            #     group_by(strt) %>% 
            #     dunn_test(PD ~ field1)#%>% 
            ph_model<-emmeans(model_PD,~ field1 )   
            posthoc_res<-as.data.frame(summary(pairs(ph_model, simple = "each", combine = TRUE,adjust = "BH")))#%>% 
                          #rename_with(~str_remove(., 'simple.contrasts.for.field1.')) 
            PD_posthoc_sig<-subset(posthoc_res,posthoc_res$p.value <=0.05)
          }
      } else{ #missing(strata_col)
        alpha_divers_pd_meta<- a_pd_meta %>%
                              dplyr::select(rowname,PD,log_PD,SR,field1=eval(field1)) %>%
                              dplyr::filter( str_length(field1)>0  & !is.na(field1) &field1!="na" ) %>%
                              column_to_rownames("rowname")
          print(paste("No strata, field1:",field1))
          if (uselog) {
            stat_PD <- kruskal_test(formula =log_PD ~ field1, data=alpha_divers_pd_meta)%>%
                      dplyr::rename('Pr(>F)'="p")
            stat_PD_call<-paste("log(PD) ~ ",field1)
          }else{
            stat_PD <- kruskal_test(formula =PD ~ field1, data=alpha_divers_pd_meta)%>%
              dplyr::rename('Pr(>F)'="p")
            stat_PD_call<-paste("PD ~ ",field1)
          }
        if (posthoc){
            anova_2way <- aov(log_PD ~ field1, data=alpha_divers_pd_meta) #
            tukey<- TukeyHSD(anova_2way)
            posthoc_res<-as.data.frame(rbind(tukey$field1))
            PD_posthoc_sig<-subset(posthoc_res,posthoc_res$'p adj' <=0.05)
          }
      }
      print(stat_PD)
      print(stat_PD_call)

      # print("pairwise.wilcox.test_PD")
      # pairwise.wilcox.test_PD <- pairwise.wilcox.test(alpha_divers_pd_meta$PD,alpha_divers_pd_meta[[field1]],p.adjust.method="none" )
      # print(pairwise.wilcox.test_PD)
#browser()
      # plot_stats <- posthoc_res %>%
      #       mutate(p.adj = format.pval(p.adj, digits = 3)) %>%
      #       add_xy_position(x = "field1", dodge = 0.8)
      if (inc_plot){
#browser()
        if (uselog) g<-ggplot(a_pd_meta, aes(x=get(field1), y=log_PD, color=get(field1))) else g<-ggplot(a_pd_meta, aes(x=get(field1), y=PD, color=get(field1)))
        print( g+
                 ggtitle(paste(ifelse(uselog,"log of",""),"FaithPD-phylogenetic diversity (PD)", sheet_a,
                               "\n",stat_PD_call,
                               "\n",eval(field1), stat_PD$'Pr(>F)'[1],
                               "\nPosthoc sig",paste(rownames(PD_posthoc_sig), collapse = ";\n"))) +  
                xlab(eval(field1))+
                 geom_boxplot(lwd=1)+
                 geom_point(position = position_dodge(width = 0.5),size=0.5)+
                theme_bw() + 
                {if(!missing(spec_colors)) scale_color_manual(values = spec_colors)} +
                guides(color=guide_legend(title=eval(field1)))+
                theme(axis.text.x = element_text(angle = 90),
                      panel.border = element_rect(size=2),
                      panel.grid.major = element_blank(),
                      panel.grid.minor = element_blank(),
                      panel.background = element_blank(),
                      text = element_text(size = 10))) #+
                #stat_pvalue_manual(plot_stats, label = "{p.adj}"))
                #stat_compare_means(method = "wilcox.test",p.adjust.method = "bonferroni")) 
                # stat_compare_means(method = "kruskal.test")) 
        if (plot_jpg) {
          ggsave(paste0(result_dir,"/FaithPD_PD_",eval(field1),"_boxplot_",proj,"_",set,"_",sheet_a,".jpg"), 
                width=7, height=5)
          dev.off()
        }
      }#inc_plot
     }
     writeData(wb_in,sheet_a_anova,stat_PD_call,
               colNames=T, rowNames=T, startRow= 1) 
     addStyle(wb_in, sheet_a_anova, style = titStyle, cols=1:10, rows=1:1, gridExpand = TRUE)
     st_row=st_row+1
#browser()
     if (!is.null(lmer_PD)){
       writeData(wb_in,sheet_a_anova, lmer_PD, colNames = T, rowNames= T, startRow= st_row) 
       st_row=st_row+nrow(lmer_PD)+1
     }
     writeData(wb_in,sheet_a_anova, stat_PD, colNames = T, rowNames= T, startRow= st_row) 
     st_row=st_row+nrow(stat_PD)+2
     
#browser()
     if (posthoc){
       
       #if (nrow(PD_posthoc_sig)>0){
         writeData(wb_in,sheet_a_anova,"PD posthoc",
                   colNames=T, rowNames=T, startRow= 1,startCol= 10) 
         writeData(wb_in,sheet_a_anova, posthoc_res, colNames = T, rowNames= T, 
                   startRow= 2, startCol= 10) 
       # } else {
       #   writeData(wb_in,sheet_a_anova,paste(pairwise.wilcox.test_PD$method,pairwise.wilcox.test_PD$data.name),
       #             colNames=T, rowNames=T, startRow= 1,startCol= 10) 
       #   writeData(wb_in,sheet_a_anova, pairwise.wilcox.test_PD$p.value, colNames = T, rowNames= T, 
       #             startRow= 2, startCol= 10) 
       # }
     }
#browser()
     print(st_row)
     st_row<-max(st_row,nrow(posthoc_res))+2
 } 
   

  ###------------------------alpha_diversity----------------------------------
   print("alpha_diversity")
   ##Test whether the observed number of ASVs differs significantly between groups
#browser()
   measures <- c("Observed",  "Shannon")

   alpha_diversity = estimate_richness(phy_rarefied,measures=measures) 
   pairwise.wilcox.test_measures <- NULL

   #alpha_diversity$ShannonEvenness = rich$Shannon / log(rich$Observed)
   #rownames(alpha_diversity) = gsub("X", "", rownames(alpha_diversity))
   #writeData(wb_in,sheet_a_divers , alpha_diversity, colNames = TRUE) 
   #setColWidths(wb_in, sheet_a_divers, cols = 1:8, widths = "auto")
   
   #measures <- c("Observed", "Chao1", "ACE", "Shannon", "Simpson", "InvSimpson", "Fisher")
   print(measures)
   if (inc_plot){
     if (missing(field2)){
       print(plot_richness(phy_rarefied, x=eval(field1), measures=measures, color =eval(field1)) + 
               xlab(eval(field1))+
               geom_boxplot()+
               ggtitle(paste("Richness box per",eval(field1),sheet_a,"-",set_name))+ 
               geom_point(position = position_dodge(width = 0.5))+
               #stat_compare_means(method = "wilcox.test",p.adjust.method = "bonferroni"))
               stat_compare_means(method = "kruskal.test"))
       if (plot_jpg) {
         dev.copy(jpeg,filename=paste0(result_dir,"/Alpha_",sheet_a_anova,"_boxplot_",set_name, ".jpg"), 
                width=800, height=400)
        dev.off()
       }
     } else{
       print(plot_richness(phy_rarefied, x=eval(field1), measures=measures, color=eval(field2)) + 
               xlab(eval(field1))+
               geom_boxplot()+
               ggtitle(paste("Richness box per",eval(field1),sheet_a,"-",set_name))+ 
               geom_point(position = position_dodge(width = 0.5))+
               stat_compare_means(method = "kruskal.test"))
       if (plot_jpg) {
         dev.copy(jpeg,filename=paste0(result_dir,"/Alpha_",sheet_a_anova,"_boxplot_",set_name, ".jpg"), 
                width=400, height=400)
          dev.off()
       }
       print(plot_richness(phy_rarefied, x=eval(field2), measures=measures, color=eval(field1)) + 
               xlab(eval(field2))+
               geom_boxplot()+
               ggtitle(paste("Richness box per",eval(field2),sheet_a,"-",set_name))+ 
               geom_point(position = position_dodge(width = 0.5))+
               stat_compare_means(method = "kruskal.test"))
       if (plot_jpg) {
         dev.copy(jpeg,filename=paste0(result_dir,"/Alpha_",sheet_a_anova,"_boxplot_",set_name, ".jpg"), 
                width=800, height=400)
        dev.off()
       }
       if(!missing(field3)){
         print(field3)
         print(plot_richness(phy_rarefied, x=eval(field1), measures=measures, color=eval(field3)) + 
                 xlab(eval(field2))+
                 geom_boxplot()+
                 ggtitle(paste("Richness box per",eval(field2),sheet_a,"-",set_name))+ 
                 geom_point(position = position_dodge(width = 0.5))+
                 stat_compare_means(method = "kruskal.test"))
         if (plot_jpg) {
           dev.copy(jpeg,filename=paste0(result_dir,"/Alpha_",eval(field1),"_",eval(field3),"_boxplot_",set_name, ".jpg"), 
                  width=1000, height=500)
          dev.off()
         }
         print(plot_richness(phy_rarefied, x=eval(field2), measures=measures, color=eval(field3)) + 
                 xlab(eval(field2))+
                 geom_boxplot()+
                 ggtitle(paste("Richness box per",eval(field2),sheet_a,"-",set_name))+ 
                 geom_point(position = position_dodge(width = 0.5))+
                 stat_compare_means(method = "kruskal.test"))
         if (plot_jpg) {
           dev.copy(jpeg,filename=paste0(result_dir,"/Alpha_",sheet_a_anova,"_boxplot_",set_name, ".jpg"), 
                  width=800, height=400)
          dev.off()
         }
       }
     }
   }#inc_plot
   print(st_row)
   st_row<-max(st_row,(nrow(posthoc_res)))+2 #+nrow(SR_tukey_df_sig)
   #st_row<-12
#browser()
   md<-sample_data(phy_rarefied)
   alpha_divers_meta<- alpha_diversity %>%
                         rownames_to_column("rowname") %>%
                         #mutate(rowname=str_replace_all(rowname,"\\.","\\-")) %>%
                         mutate(rowname=str_remove(rowname, "^X"))  %>%
                         left_join(rownames_to_column(meta_16S,"rowname"),by="rowname") %>%
                         column_to_rownames("rowname")
   print("measures loop")
#   wilcox <- data.frame(measure=character(),field1_pv=double(), field2_pv=double())
   for (m in measures){
     #m<-"Shannon"
#     wilcox_field1 <- pairwise.wilcox.test(alpha_divers_meta[[m]], alpha_divers_meta[[field1]] )

     if(missing(field2)) {
       print("alpha field1")
       if (!missing(strata_col)) {
         alpha_divers_meta_m<- alpha_divers_meta %>%
                                   dplyr::select(m=eval(m),field1=eval(field1),strt=eval(strata_col)) %>%
                                   dplyr::filter( str_length(field1)>0  & !is.na(field1)) %>%
                                   mutate(log_m=log(m)) #%>%
#browser()       
         model_a<- lmerTest::lmer(log_m ~ field1+ (1 | strt), data=alpha_divers_meta_m)
         lmer_a <- as.data.frame(summary(model_a)$coefficients)
         stat_a <- anova(model_a, ddf="Kenward-Roger") %>%
                     rownames_to_column("fields") %>%
                     mutate(fields=str_replace_all(fields,"field1",eval(field1))) #%>%

         anova_call<-paste("log(",m,") ~ ",field1,"strata=",strata_col,"using lmerTest::lmer+anova")
       } else {
#browser()
         alpha_divers_meta_m<- alpha_divers_meta %>%
                                  dplyr::select(m=eval(m),field1=eval(field1)) %>%
                                  dplyr::filter( str_length(field1)>0  & !is.na(field1)) #%>%
         stat_a <- kruskal_test(formula =m ~ field1, data=alpha_divers_meta_m)
         anova_call<-paste(m,"~",field1,"using kruskal_test")
         }
       pairwise.wilcox.test_measures <- pairwise.wilcox.test(alpha_diversity[[m]],md[[field1]],p.adjust.method="none" )
     }else{
        if(missing(field3)) {
          print("alpha 2*field")
#browser()
           
           if (!missing(strata_col)) {
#browser()       
             alpha_divers_meta_m<- alpha_divers_meta %>%
                                  dplyr::select(m=eval(m),field1=eval(field1),field2=eval(field2),strt=eval(strata_col)) %>%
                                  dplyr::filter( str_length(field1)>0  & !is.na(field1) & str_length(field2)>0  & !is.na(field2)) %>%
                                  mutate(log_m=log(m)) #%>%
               
             # perm <- how(nperm = 1000)
             # setBlocks(perm) <- with(alpha_divers_meta_m, get(strata_col))
             model_a<- lmerTest::lmer(log_m ~ field1*field2+ (1 | strt), data=alpha_divers_meta_m)
             lmer_a <- as.data.frame(summary(model_a)$coefficients)%>%
                         rownames_to_column("fields") %>%
                         mutate(fields=str_replace_all(fields,"field1",eval(field1)),
                                fields=str_replace_all(fields,"field2",eval(field2))) #%>%
             stat_a <- anova(model_a, ddf="Kenward-Roger") %>%
                         rownames_to_column("fields") %>%
                         mutate(fields=str_replace_all(fields,"field1",eval(field1)),
                                fields=str_replace_all(fields,"field2",eval(field2))) #%>%
             anova_call<-paste("log(",m,") ~",field1,"*",field2,"strata=",strata_col,"using lmerTest::lmer+anova")
           } else { #missing(strata_col)
#browser()
             alpha_divers_meta_m<- alpha_divers_meta %>%
                                      dplyr::select(m=eval(m),field1=eval(field1),field2=eval(field2)) %>%
                                      dplyr::filter( str_length(field1)>0  & !is.na(field1) & str_length(field2)>0  & !is.na(field2)) %>%
                                      mutate(log_m=log(m)) #%>%
             stat_a <- as.data.frame(summary(lm(log_m ~ field1*field2, data=alpha_divers_meta_m))$coefficients)
             anova_call<-paste("log(",m,") ~",field1,"*",field2,"using lm")
           }
       } else {
         print("alpha 3*field")
#browser()
         if (!missing(strata_col)) {
#browser()
           alpha_divers_meta_m<- alpha_divers_meta %>%
                                   dplyr::select(m=eval(m),field1=eval(field1),field2=eval(field2),field3=eval(field3),strt=eval(strata_col)) %>%
                                   dplyr::filter( str_length(field1)>0  & !is.na(field1) & 
                                                    str_length(field2)>0  & !is.na(field2) & 
                                                    str_length(field3)>0  & !is.na(field3)) %>%
                                   mutate(log_m=log(m)) #%>%
           if (plus_f3){
             model_a<- lmerTest::lmer(log_m ~ field1*field2+field3+ (1 | strt), data=alpha_divers_meta_m)
             lmer_a <- as.data.frame(summary(model_a)$coefficients)%>%
               rownames_to_column("fields") %>%
               mutate(fields=str_replace_all(fields,"field1",eval(field1)),
                      fields=str_replace_all(fields,"field2",eval(field2)),
                      fields=str_replace_all(fields,"field3",eval(field3))) #%>%
             stat_a <- anova(model_a, ddf="Kenward-Roger") %>%
               rownames_to_column("fields") %>%
               mutate(fields=str_replace_all(fields,"field1",eval(field1)),
                      fields=str_replace_all(fields,"field2",eval(field2)),
                      fields=str_replace_all(fields,"field3",eval(field3))) #%>%
             anova_call<-paste("log(",m,") ~",field1,"*",field2,"+",field3,"strata=",strata_col,"using lmerTest::lmer+anova")
           } else {
             model_a<- lmerTest::lmer(log_m ~ field1*field2*field3+ (1 | strt), data=alpha_divers_meta_m)
             lmer_a <- as.data.frame(summary(model_a)$coefficients)%>%
                         rownames_to_column("fields") %>%
                         mutate(fields=str_replace_all(fields,"field1",eval(field1)),
                                fields=str_replace_all(fields,"field2",eval(field2)),
                                fields=str_replace_all(fields,"field3",eval(field3))) #%>%
             stat_a <- anova(model_a, ddf="Kenward-Roger") %>%
                         rownames_to_column("fields") %>%
                         mutate(fields=str_replace_all(fields,"field1",eval(field1)),
                                fields=str_replace_all(fields,"field2",eval(field2)),
                                fields=str_replace_all(fields,"field3",eval(field3))) #%>%
             anova_call<-paste("log(",m,") ~",field1,"*",field2,"*",field3,"strata=",strata_col,"using lmerTest::lmer+anova")
           }
         } else { #missing(strata_col)
           alpha_divers_meta_m<- alpha_divers_meta %>%
                     dplyr::select(m=eval(m),field1=eval(field1),field2=eval(field2),field3=eval(field3)) %>%
                     dplyr::filter( str_length(field1)>0  & !is.na(field1) & 
                                      str_length(field2)>0  & !is.na(field2) & 
                                      str_length(field3)>0  & !is.na(field3)) %>%
                     mutate(log_m=log(m)) #%>%
           if (plus_f3){
             stat_a <- as.data.frame(summary(lm(log_m ~ field1*field2+field3, data=alpha_divers_meta_m))$coefficients)
             anova_call<-paste("log(",m,") ~",field1,"*",field2,"+",field3,"using lm")
           } else {
#browser()
             stat_a <- as.data.frame(summary(lm(log_m ~ field1*field2*field3, data=alpha_divers_meta_m))$coefficients)
             anova_call<-paste("log(",m,") ~",field1,"*",field2,"*",field3,"using lm")
             
           }
         }
       }
     }
#browser()
     print(stat_a)
     writeData(wb_in,sheet_a_anova,anova_call,
               colNames=T, rowNames=T, startRow= st_row) 
     addStyle(wb_in, sheet_a_anova, style = titStyle, cols=1:10, rows=st_row:st_row, gridExpand = TRUE)
     if (!is.null(lmer_a)){
       writeData(wb_in,sheet_a_anova, lmer_a, colNames = T, rowNames= T, startRow= st_row+1) 
       st_row=st_row+nrow(lmer_a)+2
     }
     writeData(wb_in,sheet_a_anova,paste(pairwise.wilcox.test_measures$method,pairwise.wilcox.test_measures$data.name),
               colNames=T, rowNames=T, startRow= st_row,startCol= 10) 
     writeData(wb_in,sheet_a_anova, stat_a, colNames = T, rowNames= T, startRow= st_row+1) 
     st_row=st_row+nrow(stat_a)+2
     writeData(wb_in,sheet_a_anova, pairwise.wilcox.test_measures$p.value, colNames = T, rowNames= T, 
               startRow= st_row, startCol= 10) 
     st_row<-st_row+max(nrow(stat_a),nrow(pairwise.wilcox.test_measures$p.value))+2
     
   }
#browser()
   if (PD) return(alpha_divers_pd_meta)
 }

 
 Bdiver<- function(wb_in=wb, phy_rarefied,field1,field2,field3, dist_in,plus_f3=T,Bdiver_dist=T,
                   ord_meths, ellipse=T, keep_dist=F, to_label,anova_in,anova_pair,spec_colors,
                   set_name="all",   dst= "unifrac" ,weight=T ,sheet_b= NULL,plot_jpg=T){
   #phy_rarefied<-physeq_rarefied_MW
   #field1<- "Condition" #RatType TimeW Condition
   #field2<- "Wasp" #TimeGroup PatientID Wasp
   #field3<- "Stage" #RatID batch Stage
   # plus_f3=F
   #dst<- "unifrac" #bray unifrac
   set.seed(42)
   
   sheet_b_divers <- paste(set_name,sheet_b)
   addWorksheet(wb_in, sheet_b_divers)
   st_row<-1
   print(sheet_b_divers)
   titStyle <- createStyle(fontColour = "#006100", fgFill = "#C6EFCE")
   
   # PCoA plot using the weighted/un UniFrac/bray as distance
   theme_set(theme_bw())
   # methods for distance
   #str(phy_tree(phy_rarefied))
#browser()
   phy_tree(phy_rarefied) <- ape::multi2di(phy_tree(phy_rarefied))
   #dist_methods <- c("bray")
   ##  distances between samples
   if (missing(anova_in))   {
     dist_in <- phyloseq::distance(phy_rarefied, method=set_name, weighted=weight) #dst
     md_rarefied<-data.frame(sample_data(phy_rarefied))
   }
   
   # ANOVA-Test whether the groups, Condition,  differ significantly from each
   if (missing(field2)){
     anova_call<-paste(sheet_b_divers,": dist ~ ",field1)
#browser()
    dist_df<- as.data.frame(as.matrix(dist_in))
  
     # dist_df<- as.data.frame(reshape2::melt(as.matrix(dist_in))) %>%
     #             dplyr::filter(Var1!=Var2) %>%
     #             mutate(Var1=as.factor(Var1)) %>%
     #             mutate(Var2=as.factor(Var2)) %>%
     #             left_join(rownames_to_column(dplyr::select(meta_16S,field1),"Var1"), by="Var1") %>%
     #             left_join(rownames_to_column(dplyr::select(meta_16S,field1),"Var2"), by="Var2") %>%
     #             mutate(cond1=get(paste0(field1,".x"))) %>%
     #             mutate(cond2=get(paste0(field1,".y"))) %>%
     #             mutate(cond=as.factor(paste0(cond1, "_",cond2)) )
#      if (missing(anova_in)) anova_in <- adonis(dist_in ~ md_rarefied[[field1]], data=md_rarefied)
      if (missing(anova_in)) anova_in <- adonis(dist_in ~ get(field1), data=md_rarefied)
#browser()
    if (scale_type(sample_data(phy_rarefied)[[eval(field1)]])!="continuous"){
       #Dispersion test-the average distance of group members to the group centroid in multivariate space (generated by a distance matrix)
       dispr <- vegan::betadisper(dist_in, sample_data(phy_rarefied)[[eval(field1)]])
       anova_dispr<-anova(dispr)
       
       print(plot(dispr, main = paste("Ordination Centroids Distance", sheet_b_divers), 
                  sub = paste("Anova betadisper",anova_dispr[1,]$`Pr(>F)`)))
       # print(anova_in)
    } else {
#browser()
      if (as.numeric(as.data.frame(anova_in)[1,5])<0.1){
        dist_meta<- as.data.frame(reshape2::melt(as.matrix(dist_in))) %>%
                    left_join(rownames_to_column(subset(md_rarefied,select=field1),"Var1"),by="Var1") %>%
                    left_join(rownames_to_column(subset(md_rarefied,select=field1),"Var2"),by="Var2") %>%
                    dplyr::select(field1.x=eval(paste0(field1,".x")),field1.y=eval(paste0(field1,".y")),value,Var1,Var2) %>%
                    mutate(field1_diff=field1.x-field1.y) %>%
                    dplyr::filter(field1_diff>0) #%>%
        print(ggplot(dist_meta, aes(x = field1_diff, y = value,color = value))+
                # Scatter plot with regression line
                ylab("Distances")+
                xlab(paste(field1,"Difference")) +
                geom_point()+
                geom_smooth(aes(color =value, fill = value),
                            method = "lm") +
                stat_cor())#+ #method = "spearman" #correlation coefficient, meaning R
      }
    }
   }  else if (missing(field3)){
#browser()     
     if (missing(anova_in))    anova_in <- adonis2(dist_in ~ sample_data(phy_rarefied)[[field1]] *
                                                    sample_data(phy_rarefied)[[field2]] )
     anova_call<-paste(sheet_b_divers,": dist ~ ",field1,"*",field2)
     dist_df<- as.data.frame(as.matrix(dist_in))
     # dist_df<- as.data.frame(melt(as.matrix(dist_in))) %>%
     #             dplyr::filter(Var1!=Var2) %>%
     #             left_join(rownames_to_column(dplyr::select(meta_16S,field1,field2),"Var1"), by="Var1") %>%
     #             left_join(rownames_to_column(dplyr::select(meta_16S,field1,field2),"Var2"), by="Var2") %>%
     #             mutate(cond1=paste0(get(paste0(field1,".x")),"_",get(paste0(field2,".x")))) %>%
     #             mutate(cond2=paste0(get(paste0(field1,".y")),"_",get(paste0(field2,".y")))) %>%
     #             mutate(cond=as.factor(paste0(cond1, "_",cond2)) )
   } else {
     if (plus_f3){
       print("plus_f3")
       if (missing(anova_in))    anova_in <- adonis2(dist_in ~ sample_data(phy_rarefied)[[field1]] *
                                         sample_data(phy_rarefied)[[field2]] +
                                         sample_data(phy_rarefied)[[field3]])
       anova_call<-paste(sheet_b_divers,": dist ~ ",field1,"*",field2,"+",field3)
     } else {
       if (missing(anova_in))    anova_in <- adonis2(dist_in ~ sample_data(phy_rarefied)[[field1]] *
                                         sample_data(phy_rarefied)[[field2]] *
                                         sample_data(phy_rarefied)[[field3]])
       anova_call<-paste(sheet_b_divers,": dist ~ ",field1,"*",field2,"*",field3)
     }
     dist_df<- as.data.frame(as.matrix(dist_in))
     # dist_df<- as.data.frame(melt(as.matrix(dist_in))) %>%
     #           dplyr::filter(Var1!=Var2) %>%
     #           left_join(rownames_to_column(dplyr::select(meta_16S,field1,field2,field3),"Var1"), by="Var1") %>%
     #           left_join(rownames_to_column(dplyr::select(meta_16S,field1,field2,field3),"Var2"), by="Var2") %>%
     #           mutate(cond1=paste0(get(paste0(field1,".x")),"_",get(paste0(field2,".x")),"_",get(paste0(field3,".x")))) %>%
     #           mutate(cond2=paste0(get(paste0(field1,".y")),"_",get(paste0(field2,".y")),"_",get(paste0(field3,".y")))) %>%
     #           mutate(cond=as.factor(paste0(cond1, "_",cond2)) )
   }
   #print(anova_in)
#browser()  
   writeData(wb_in,sheet_b_divers,startRow = st_row,
             #rownames_to_column(anova_in$aov.tab,anova_call),
             rownames_to_column(anova_in,anova_call),
             colNames = TRUE) 
   if (!missing(anova_pair))   {
     writeData(wb_in,sheet_b_divers,startRow = st_row, startCol = 10,
                                         colNames = T, rowNames = T,
                                         as.data.frame(t(as.data.frame(anova_pair))) ) 
#browser()
     # mean_dist_df<- as.data.frame(reshape2::melt(as.matrix(dist_in))) %>%
     #             separate(Var1,into = c(NA,NA,"Condition1"),sep = "\\.", remove = F,extra = "merge") %>%
     #             separate(Var2,into = c(NA,NA,"Condition2"),sep = "\\.", remove = F,extra = "merge") %>%
     #             group_by(Condition1,Condition2) %>% 
     #             dplyr::summarise(mean_dist = mean(value))  #%>%
     # 
     # 
     # anova_pair_df<-as.data.frame(anova_pair) %>%
     #               dplyr::select(ends_with("Pr..F.")) %>%
     #               rename_with(~str_remove(.,".Pr..F.")) %>%
     #               dplyr::filter(row_number()==1) %>%
     #               t() %>%
     #               as.data.frame() %>%
     #               dplyr::select("a_paired"=condition) %>%
     #               mutate(a_sig=case_when(a_paired<=0.001 ~ "***",
     #                                      a_paired<=0.01 ~ "**", 
     #                                      a_paired<=0.05 ~ "*",
     #                                      TRUE ~ "")) %>%
     #               rownames_to_column("condition") %>%
     #               separate(condition,into = c("Condition1","Condition2"),sep = "_vs_")
     # 
     # 
     # 
     # dist_anova<- mean_dist_df %>%
     #               mutate(Distances=if_else(as.character(Condition1)>as.character(Condition2),mean_dist,
     #                                        if_else(as.character(Condition1)==as.character(Condition2),0,100))) %>%
     #               dplyr::filter(Distances!=100) %>%
     #               left_join(anova_pair_df,by=c("Condition1","Condition2")) %>%
     #               left_join(anova_pair_df,by=c("Condition1"="Condition2","Condition2"="Condition1")) %>%
     #               mutate(sig=  if_else(is.na(a_sig.x),a_sig.y,a_sig.x)) #%>%
     # 
     # 
     # print(ggplot(dist_anova, aes(x=Condition1, y=Condition2, fill=Distances))+
     #         geom_tile(color="white")+
     #         geom_text(aes(label = sig), color = "black", size = 4) +
     #         theme_minimal()+
     #         theme(text = element_text(size = 12),
     #               panel.grid.major = element_blank(), 
     #               panel.grid.minor = element_blank(),
     #               panel.background = element_blank(),
     #               axis.text.x = element_text(angle = 45, hjust = 1))+ 
     #         ggtitle(paste("Mean Dist",set_name,proj,set,"heatmap"))+
     #         scale_fill_gradient(low = "#FF0000", high = "#FFFFFF",name="Mean Distances"))
     # 
     # if (plot_jpg) {
     #   ggsave(paste0(result_dir,"/mean_dist_",set_name,"_",proj,"_",set,"_heatmap",".jpg"), 
     #          width=7, height=5.2)
     #   dev.off()
     # }
     
   }
   
   
   # beta diversity boxplots-distribution of distances
   st_row<-st_row+8
   # anova3_2way <- aov(value ~  cond1+cond2, data=dist_df) #
   # #anova3_2way <- summary(aov(as.matrix(dist_in) ~  sample_data(phy_rarefied)[[eval(field1)]])) #
   # tukey<- TukeyHSD(anova3_2way)
   # tukey_df<-as.data.frame(tukey$cond2)
   # tukey_df_sig<-subset(tukey_df,tukey_df$`p adj` <0.05)
   # print(ggplot(dist_df, aes(x=as.factor(cond1), y=value, color=as.factor(cond1))) +
   #         geom_boxplot()+
   #         xlab(eval(field1)) +
   #         theme_bw()+
   #         facet_wrap(~cond2, scales="free_x")+
   #         #geom_jitter(shape=16, position=position_jitter(0.2))+
   #         theme(legend.position = "none", 
   #               text = element_text(size = 12),
   #               axis.text.x = element_text(angle = 45, hjust = 1))+
   #         ggtitle(paste(sheet_b_divers, "Group significance plot",
   #                      # "\nAnova2way",anova2$aov.tab$`Pr(>F)`[1],
   #                       "\n sig tukey:",paste(rownames(tukey_df_sig), collapse = ";")))+
   #         stat_compare_means(method = "wilcox.test",p.adjust.method = "bonferroni"))
   # if (plot_jpg) {
   #   ggsave(paste0(result_dir,"/boxplot_dist_",proj,"_",set,"_",sheet_b_divers,".jpg"), 
   #        width=7, height=5)
   #   dev.off()
   # }
   if (keep_dist) assign(paste0("dist_df_",str_replace_all(sheet_b_divers," ","_")), dist_df, envir = .GlobalEnv)
#browser()   
   gc()
  #---------------------posthoc------------------------------------
   if (Bdiver_dist){
     for (cnd in unique(dist_df$cond1)){
       #cnd<-"G_MW_CTRL-G"
       dist_cnd<-subset(dist_df,cond1==cnd ) #& cond1!=cond2
       #deal with zeros
       dist_cnd$value<-sqrt (1+dist_cnd$value)
       
       anova2 <- adonis(value ~ cond2, data = dist_cnd)
       anova_2way <- aov(value ~  cond2, data=dist_cnd) #
       tukey<- TukeyHSD(anova_2way)
       tukey_df<-as.data.frame(tukey$cond2)
       tukey_df_sig<-subset(tukey_df,tukey_df$`p adj` <0.05)
       
       writeData(wb_in,sheet_b_divers,paste(cnd,as.character(anova2$call[2])),
                 colNames=T, rowNames=T, startRow= st_row) 
       addStyle(wb_in, sheet_b_divers, style = titStyle, cols=1:10, rows=st_row:st_row, gridExpand = TRUE)
       writeData(wb_in,sheet_b_divers, anova2$aov.tab, colNames = T, rowNames= T, startRow= st_row+1) 
       writeData(wb_in,sheet_b_divers, paste(rownames(tukey_df_sig), collapse = ";  "),
                 colNames = T, rowNames= T, startRow= st_row+5) 
       st_row<-st_row+8
       
     }
   }
   setColWidths(wb_in, sheet_b_divers, cols = 1:1, widths = 50)
   setColWidths(wb_in, sheet_b_divers, cols = 2:10, widths = "auto")
#browser()   
   #-----------------------plot ordination--------------------------------
   if (missing(ord_meths)) ord_meths = c( "CCA", "RDA", "DPCoA", "NMDS", "PCoA") #"DCA", "MDS",
   #ord_meths =  "CCA"
   for (mth in ord_meths){
     #  for (dst in dist_methods){
     #mth<- "PCoA"
     
     print(paste("method:", mth, "dist method: ", set_name))
#browser(expr =(mth=="NMDS") )
     if (mth!="DPCoA") ord <- ordinate(phy_rarefied, method=mth, distance=dist_in, weighted=weight) else ord <- ordinate(phy_rarefied, method=mth, distance=dist_in)
#browser()
     #plots
     result = tryCatch({
       if (!missing(field2)){
         if (!missing(to_label)) {
           g<-plot_ordination(phy_rarefied, ord, color=eval(field1), shape = eval(field2), label=eval(to_label)) 
          }else{
            if (scale_type(sample_data(phy_rarefied)[[eval(field2)]])!="continuous"){
              g<-plot_ordination(phy_rarefied, ord, color=eval(field1), shape = eval(field2))
            }else{
              if (scale_type(sample_data(phy_rarefied)[[eval(field1)]])!="continuous"){
                g<-plot_ordination(phy_rarefied, ord, color=eval(field2), shape = eval(field1))
              }else{
                g<-plot_ordination(phy_rarefied, ord, color=eval(field2))
              }
            }
          }
          g<-g+ggtitle(paste("Distance", sheet_b_divers,mth,"Ordination",
                           "\nAnova",eval(field1), round(as.data.frame(anova_in)[1,5],digits = 2),
                           "\nAnova",eval(field2),round(as.data.frame(anova_in)[2,5],digits = 2),
                           "\nAnova",eval(field1),":",eval(field2),round(as.data.frame(anova_in)[3,5],digits = 2)))
                           
       
       } else{
         print("only field1")
         if (!missing(to_label)) g<-plot_ordination(phy_rarefied, ord, color=eval(field1), label=eval(to_label)) + 
                                    geom_text( mapping = aes(label = get(to_label)), size = 5, vjust = 1.5) else
                                g<-plot_ordination(phy_rarefied, ord, color=eval(field1)) 
         g<-g+  
           ggtitle(paste("Distance", sheet_b_divers,mth,"Ordination",
                         "\nAnova",eval(field1), as.data.frame(anova_in)[1,5]))#+ 
#browser()
       }
       g<-g+  
             #geom_label_repel(aes(label=genelabels),color="black", max.overlaps=Inf, size =2, 
            #                  box.padding = unit(0.2, "lines"))+
            theme(aspect.ratio=1,plot.title = element_text(size = 16, face = "bold"),
                  panel.grid.major = element_blank(),
                  panel.grid.minor = element_blank(),
                  panel.background = element_blank(),
                  text = element_text(size = 16))+
           {if(!missing(spec_colors)) scale_color_manual(values = spec_colors)} +
           {if((ellipse & scale_type(sample_data(phy_rarefied)[[eval(field1)]])!="continuous")) ggforce::geom_mark_ellipse()} +
           guides(shape = guide_legend(order = 0),
                    color  = guide_legend(order = 1))+
            geom_point(size = 5) 
        print(g)
       
       if (plot_jpg) {
         dev.copy(jpeg,filename=paste0(result_dir,"/Beta_",mth,"_",sheet_b_divers, "_plot.jpg"), 
                width=500, height=500)
        dev.off()
       }
       
       if (!missing(field3)){
         g2<-plot_ordination(physeq_rarefied, ord, color=eval(field1), shape = eval(field3)) + #, label="SampleID"
                 theme(aspect.ratio=1,plot.title = element_text(size = 16, face = "bold")) +
                 geom_point(size = 5) +
                 ggtitle(paste( "Distance", sheet_b_divers,mth,"Ordination"))+ 
                 scale_shape_manual(values = c(20,18))
         if (ellipse){
           print(g2+ggforce::geom_mark_ellipse())
           #print(g2+stat_ellipse() +coord_fixed())#+ #geom = "circle")
         }else{
           print(g2)
         }
         if (plot_jpg) {
           dev.copy(jpeg,filename=paste0(result_dir,"/Beta_",mth,"_",sheet_b_divers, "_plot2.jpg"), 
                  width=600, height=600)
          dev.off()
         }
       }
     },
     error = function (condition) {
       print(paste("ERROR in method:", mth, "set_name: ", set_name))
     })
   }
   # print(plot_ordination(phy_rarefied, ord, type="taxa", color="Phylum",  
   #                       title="taxa distance"))
   sig_anova<-as.data.frame(subset(anova_in,`Pr(>F)`<0.05))
   #sig_anova<-as.data.frame(subset(anova$aov.tab,`Pr(>F)`<0.05))
   print(paste("no of sig anova:",nrow(sig_anova)))
   if (nrow(sig_anova)>0){return(TRUE)}else{return(FALSE)}
 }
 
dist_heatmap<- function(phy_rarefied,field,anova_pair_cond_w,dist_in,mthd="unifrac", 
                        set_name="",pdf=T,
                        w=T,md_in,res_dir=result_dir){
#browser()  
  if(missing(md_in)){
    md_in<-data.frame(sample_data(phy_rarefied)) #%>%
              #arrange(get(field))
  }
  md<-md_in%>%
      dplyr::rename("cond"=eval(field))
  #dist_in <- phyloseq::distance(phy_rarefied, method=mthd, weighted=w) #
  if (missing(anova_pair_cond_w)){
    assign("dist_4heatmap",dist_in,envir = .GlobalEnv)
    anova_pair_cond_w <- pairwiseAdonis::pairwise.adonis2(dist_4heatmap ~ cond,data=md)
  }
  print(anova_pair_cond_w)
#browser()                   
  
 dist_df<- as.data.frame(reshape2::melt(as.matrix(dist_in))) %>%
             left_join(rownames_to_column(subset(md,select=cond),"Var1"),by="Var1") %>%
             left_join(rownames_to_column(subset(md,select=cond),"Var2"),by="Var2") %>%
             group_by(cond.x,cond.y) %>% 
             dplyr::summarise(mean_dist = mean(value))  %>%
             dplyr::select(cond1=cond.x,cond2=cond.y,mean_dist)
 
#browser()                   
 anova_pair_df<-as.data.frame(anova_pair_cond_w,check.names=F) %>%
                   dplyr::select(ends_with("Pr(>F)")) %>%
                   rename_with(~str_remove_all(.,"\\.Pr\\(\\>F\\)")) %>%
                   dplyr::filter(row_number()==1) %>%
                   t() %>%
                   as.data.frame() %>%
                   dplyr::select("a_paired"=eval(field)) %>%
                   # dplyr::select("a_paired"=cond) %>%
                   mutate(a_sig=case_when(a_paired<=0.001 ~ "***",
                                          a_paired<=0.01 ~ "**", 
                                          a_paired<=0.05 ~ "*",
                                          TRUE ~ "")) %>%
                   rownames_to_column("comp") %>%
                   separate(comp,into = c("cond1","cond2"),sep = "_vs_")
 
 
 dist_anova<- dist_df %>%
                 mutate(Distances=if_else(as.character(cond1)>as.character(cond2),mean_dist,
                                          if_else(as.character(cond1)==as.character(cond2),0,100))) %>%
                 dplyr::filter(Distances!=100) %>%
                 left_join(anova_pair_df,by=c("cond1","cond2")) %>%
                 left_join(anova_pair_df,by=c("cond1"="cond2","cond2"="cond1")) %>%
                 mutate(sig=  if_else(is.na(a_sig.x),a_sig.y,a_sig.x)) #%>%
                 # arrange(factor(cond1, levels = levels(md_in$AD_severity)),
                 #         factor(cond2, levels = levels(md_in$AD_severity)))
 
 if (pdf) pdf(file=paste0(res_dir,"/Dist_",mthd,"_",ifelse(w,"W","UNW"),"_",field,"_",
                 ifelse(!is.na(set_name), paste0(set_name,"_"),"_"),proj,"_",set,"_heatmap",".pdf"))
#browser()                   
 
 print(ggplot(dist_anova, aes(x=cond1, y=cond2, fill=Distances))+
   xlab(field)+
   ylab(field)+
   geom_tile(color="white")+
   geom_text(aes(label = sig), color = "black", size = 4) +
   theme_minimal()+
   theme(text = element_text(size = 12),
         panel.grid.major = element_blank(), 
         panel.grid.minor = element_blank(),
         panel.background = element_blank(),
         axis.text.x = element_text(angle = 45, hjust = 1))+ 
   ggtitle(paste(field, "Distances ",mthd,ifelse(w ,"W","UNW"),"mean heatmap-",proj,set))+
   scale_fill_gradient(low = "#FF0000", high = "#FFFFFF",name="Mean Distances"))
 
 # ggsave(paste0(res_dir,"/dist_",mthd,"_",ifelse(w,"W","UNW"),"_",field,"_",proj,"_",set,"_heatmap",".jpg"), 
 #        width=7, height=5.2)
 if (pdf) dev.off()
 return(md)
}
 
 
 deseq_part<- function(md,taxa_count,field,value, to_inc=T,dds_design,res_for,res_for2,set_name,exact=F,
                       sfactor=T, FDR_method="BH",dbg=T, woResults=F,
                       res_dir=result_dir, element="taxa",
                       to_label=T,filter_id=F,pca_text=T,out_format="jpeg"){
   #value  <-"GDM"   # "CTRL-G"
   #field<-"Control_GDM"   #Condition"
   #dds_design<-" ~ woman + trimester"   #formula(" ~ Stage") 
   #res_for<-"trimester"   #Stage"
   #set_name<-GDM_only
#browser()   
   
   if (field!="dummy"){
     if (to_inc)  {
#browser()
       if (!exact){
         md_part<-md %>%
                   dplyr::filter(str_detect(get(field),value) & str_length(md[[res_for]])>0  & !is.na(get(res_for)) & get(res_for)!="na" & get(res_for)!="NA") %>%
                   dplyr::select(-eval(field))  %>%
                   droplevels()
       }else {
         md_part<-md %>%
                    dplyr::filter(get(field)==value & str_length(md[[res_for]])>0  & !is.na(get(res_for)) & get(res_for)!="na" & get(res_for)!="NA") %>%
                    dplyr::select(-eval(field)) %>%
           droplevels()
       }
     } else {
       if (!exact){
         md_part<-md %>%
                   dplyr::filter(!str_detect(get(field),value))  %>%
         droplevels()
       }else{
         md_part<-md %>%
           dplyr::filter(get(field)!=value)  %>%
           droplevels()
       }
     }
   }else{
     md_part<-md #%>%
   }
#browser()   
   
   if (scale_type(md_part[[res_for]])!="continuous" & nlevels(md_part[[res_for]])==0)    md_part[[res_for]]<-factor(md_part[[res_for]],levels = unique(md_part[[res_for]]))
   if (dbg)
     if (missing(res_for2)){
       print(table(md_part[[res_for]]))
     } else {
       print(table(md_part[[res_for]],md_part[[res_for2]]))
     }
#browser()   
   assign(paste0("md_",set_name), md_part, envir = .GlobalEnv)
   taxa_count_part<-taxa_count %>%
                    dplyr::select(rownames(md_part)) #%>% # filter zero cols
   #select_if(purrr::negate(function(col) is.numeric(col) && sum(col) >0))
#browser()   
   assign(paste0(element,"_count_",set_name), taxa_count_part, envir = .GlobalEnv)

   dds_taxa_part <- DESeqDataSetFromMatrix(countData = taxa_count_part,
                                         colData = md_part,
                                         design=dds_design)
#browser()  
   if (dbg) {    
     dds_taxa_part = DESeq(dds_taxa_part)
    }  else {
    # dds_taxa_part = DESeq(dds_taxa_part, parallel=TRUE, BPPARAM=SnowParam(parallel::detectCores()-1), quiet = T)
    dds_taxa_part = DESeq(dds_taxa_part, quiet = T)
    }
#no normalization. Alternative library size estimators can be supplied using the assignment function sizeFactors<-
   if (!sfactor) {
     s<-sizeFactors(dds_taxa_part)
     s[1:length(s)]<-1
     sizeFactors(dds_taxa_part)<-s
   }
   if (dbg) print(resultsNames(dds_taxa_part))
   if (dbg) print(sizeFactors(dds_taxa_part))
#browser()   
   assign(paste0("dds_",set_name),dds_taxa_part, envir = .GlobalEnv)
   assign("dds_count_df",as.data.frame(counts(dds_taxa_part, normalized=TRUE)), envir = .GlobalEnv)
   assign(paste0(element,"_norm_abund_",set_name), dds_count_df, envir = .GlobalEnv)
   #for ImmQuant
   if ( field=="dummy")  { #element=="gene" &
     write.table(rownames_to_column(dds_count_df,"gene"),
                 paste0(res_dir,"/../RNAseq_normalized_",set_name,"_",set,".txt"),
               row.names = F, col.names = TRUE,sep = "\t", quote = F)
     assign("vst_all", as.data.frame(assay(varianceStabilizingTransformation(dds_taxa_part, blind=FALSE))),envir = .GlobalEnv)
   } 
   assign(paste0("vst_",set_name), as.data.frame(assay(varianceStabilizingTransformation(dds_taxa_part, blind=FALSE))),envir = .GlobalEnv)

   
   if (!woResults){
#browser()
     if (length(resultsNames(dds_taxa_part))>1 & scale_type(md_part[[res_for]])!="continuous") {
       list1<-rev(levels(md_part[[res_for]]))
       if (dbg) print(list1)
       list2<-list1
       res_all<-character()
       res_all_df<-character()
       for (r1 in list1){
         list2<-list2[list2 != r1]
         for (r2 in list2){
           if (r1!=r2){
             #r1<-"CTRL-G"; r2<-"CTRL-O"
             #The factor level given last is the base level for the comparison.
             #print(paste("res_for,r1,r2",res_for,r1,r2))
  #browser()
           taxa_res_part<-results(dds_taxa_part,contrast=c(res_for,r1,r2), pAdjustMethod=FDR_method)
             if (filter_id) taxa_res_part<-subset(taxa_res_part,!startsWith(rownames(taxa_res_part),"id_"))
             # DESeq2::summary(taxa_res_part)
             md_part2<-md_part %>%
                      dplyr::filter(get(res_for)==r1 | get(res_for)==r2)
  #browser()
             vst_part<-varianceStabilizingTransformation (dds_taxa_part, blind=FALSE) %>%
                         assay() %>%
                         as.data.frame(stringsAsFactors = FALSE) %>%
                         #rownames_to_column %>%
                         #mutate(rowname=substr(rowname,1,75)) %>%
                         #column_to_rownames %>%
                         dplyr::select(rownames(md_part2))
             if (to_inc) {name<-paste0("_",value,"_",r1,"_vs_",r2)} else{
              name<-paste0("_exclude-",value,"_",r1,"_vs_",r2)
             }
             assign(paste0(element,name),taxa_res_part, envir = .GlobalEnv)
             assign(paste0("vst",name), vst_part, envir = .GlobalEnv)
  #browser()
             if (missing(res_for2)) {
  #browser()
               anno_df<-subset(md_part2,select=eval(res_for)) %>% 
                         rownames_to_column() %>% 
                         arrange(get(res_for)) %>% 
                         column_to_rownames()
               assign(paste0("anno",name), anno_df, envir = .GlobalEnv)
               if ( field=="dummy") assign("anno_df", anno_df, envir = .GlobalEnv)
             } else {
               anno_df<-subset(md_part2,select=c(eval(res_for),eval(res_for2)))
               assign(paste0("anno",name), anno_df, envir = .GlobalEnv)
             }
             res_all<-c(res_all,paste0(element,name))
             if (dbg) print(paste("res_all:",res_all))
             taxa_res<-as.data.frame(taxa_res_part) %>%
                        rownames_to_column("Gene_name") %>%
                        mutate(Comparison=name)
             if (dbg) print(name)
             res_all_df<-rbind(res_all_df,taxa_res)
           }
         }
       }
     } else{
  #browser()
       if (length(resultsNames(dds_taxa_part))>1) {
         name<-paste0("_",value,"_",res_for)
         taxa_res_part<-results(dds_taxa_part,name = res_for)
       } else{
         name<-paste0("_",value,"_Intercept")
         taxa_res_part<-results(dds_taxa_part)
       }
       if (dbg) DESeq2::summary(taxa_res_part)
       vst_part<-varianceStabilizingTransformation (dds_taxa_part, blind=FALSE) %>%
         assay() %>%
         as.data.frame(stringsAsFactors = FALSE) #%>%
       assign(paste0(element,name),taxa_res_part, envir = .GlobalEnv)
       assign(paste0("vst",name), vst_part, envir = .GlobalEnv)
       assign(paste0("anno",name), subset(md_part,select=eval(res_for)), 
                envir = .GlobalEnv)
       res_all<-paste0(element,name)
       if (dbg) print(paste("res_all:",res_all))
       taxa_res<-as.data.frame(taxa_res_part) %>%
                 rownames_to_column("Gene_name") %>%
                 mutate(Comparison=name)
       if (dbg) print(name)
       res_all_df<-taxa_res
     }
  
     openxlsx::write.xlsx(res_all_df,
                          paste0(res_dir,"/",element,"_results_",proj,"_",set,"_",set_name,".xlsx"),
                          rowNames = F, overwrite = T)
     assign("res_all_df",res_all_df,envir = .GlobalEnv)
   } #woResults
   
   #boxplot(log10(assays(dds_taxa_part)[["cooks"]]), range=0, las=2,
  #         main=paste("boxplot",element," Cook’s distances - test for outliers",
  #         "\n",set_name),
  #         ylab="Cook’s distances")
  #dev.copy(jpeg,filename=paste0(res_dir,"/plot_",element,"_Cooks_distances_",set_name, ".jpg"), 
  #        width=600, height=600)
  #dev.off()
   
   
   norm_count<-as.data.frame(counts(dds_taxa_part, normalized=TRUE))
   openxlsx::write.xlsx(norm_count,paste0(res_dir,"/Norm_count_",element,"_",set_name,".xlsx"),rowNames=T,overwrite = T)
   if (dbg) print (paste("no of results:",length(resultsNames(dds_taxa_part))))
   if (length(resultsNames(dds_taxa_part))>1 & pca_text) {
# browser()
     pc<- plot_PCA(dds_taxa_part,group=if (missing(res_for2)) res_for else c(res_for,res_for2), 
                   set_name=paste0(element,"_",set_name,"_for_",res_for),out_format=out_format,
                   dds_res=DESeq2::summary(taxa_res_part), res_dir = res_dir,to_label = to_label)
   } else {
     pc<- plot_PCA(dds_taxa_part,group=if (missing(res_for2)) res_for else c(res_for,res_for2), 
                   set_name=paste0("res_",element,"_",set_name,"_for_",res_for),out_format=out_format,
                   res_dir = res_dir,to_label = to_label)
     if (length(resultsNames(dds_taxa_part))<=1) res_all<-NULL
   }
   #ph<- plot_dds_heatmap(dds_taxa_part,group=res_for, res_dir= res_dir,
  #                       set_name=paste0("res_",element,"_",set_name,"_for_",res_for))
#browser()
   return(res_all)
 }
 
 
 prep_count_levels<- function(phy_rarefied_data,taxa_level=c("short_title", "GenusAgg"),
                              add_one=F,rowSums_thresh=0,colSums_thresh=0, rel=F,set_name){
   #phy_rarefied_data<-physeq_rarefied_data                      
   
   for (tl in taxa_level){
     #tl<-"short_title"
     print(tl)
#browser()     
     
     taxa_count<-phy_rarefied_data %>%
       dplyr::select(Sample,Abundance,taxa=eval(tl)) %>%
       mutate(taxa=str_trim(taxa))  %>% 
       dplyr::filter(Sample %in% rownames(meta_16S)) %>%
       dplyr::group_by(Sample,taxa) %>% 
       dplyr::summarise(total_abund = sum(Abundance))  %>%
       pivot_wider(names_from =Sample, values_from =total_abund) %>%
       dplyr::filter(taxa!="k__Bacteria" & 
                       taxa!="d__Bacteria" & 
                       str_length(taxa)>0 &
                       !is.na(taxa)) %>%
       replace(is.na(.), 0) %>% 
       column_to_rownames("taxa")
     # filter zero rows
     print(paste("nrow:",nrow(taxa_count)))
     print(paste("ncol:",ncol(taxa_count)))
#browser()
     print(head(table(rowSums(taxa_count)),10))
     print(head(table(colSums(taxa_count)),10))
     taxa_count <- taxa_count[rowSums(taxa_count != 0) > rowSums_thresh, ]
     taxa_count <- taxa_count[, colSums(taxa_count != 0) > colSums_thresh]
     #estimateSizeFactorsForMatrix: every row contains at least one zero, cannot compute log geometric means
     if (add_one)   {
       taxa_count<-taxa_count  %>% 
         mutate(multiply.100(.)) %>%
         replace(.==0, 1) #%>% 
     }
     print(paste("nrow:",nrow(taxa_count)))
     print(paste("ncol:",ncol(taxa_count)))
     assign(paste0("taxa_count_",eval(tl),if (rel) "_rel",if (!missing(set_name)) paste0("_",set_name)),taxa_count,envir = .GlobalEnv)
   }
   return(taxa_level)
   
 } 

 multiply.10 <- function(x, na.rm=FALSE) (x*10)
 multiply.100 <- function(x, na.rm=FALSE) (x*100)
 
 
 get_clusters <- function( count_df,tit,element="taxa",bootstrap=500,res_dir=result_dir,k,kmax=10,
                           rectan=0,set_name=paste0(proj,"_",set)) {
#browser()   
   count_dst <- vegdist( count_df)
   dst <- as.matrix(count_dst)
   hclst<-hclust(count_dst, method="ward.D2")
   summary(hclst)
   # Gap statistic: # nboot = 50 to keep the function speedy. 
   # recommended value: nboot= 500 for your analysis.
#browser()
   set.seed(123)
   if (missing(k)){
     gap <- clusGap(dst, kmeans,  K.max=kmax, B=bootstrap)
     gap_k <- maxSE(gap$Tab[, "gap"], gap$Tab[, "SE.sim"], method="Tibs2001SEmax")
   }else{
     gap_k<-k
   }
   #http://www.sthda.com/english/wiki/beautiful-dendrogram-visualizations-in-r-5-must-known-methods-unsupervised-machine-learning
#browser()
   count_cluster <- cutree(hclst, k = gap_k)# cut tree into gap_k clusters
   print(table(count_cluster))
   count_cluster_df<- as.data.frame(count_cluster) 
   if (gap_k>1){
     openxlsx::write.xlsx (rownames_to_column(count_cluster_df, "feature id"),
                           paste0(res_dir,"/",tit,"_k.",gap_k,"_",set_name,".xlsx"),
                           sheetName ="count_cluster_df" ,
                           rowNames = F, colNames = T, append = F,overwrite = T) 
     count_cluster_wide<- count_cluster_df %>%
       rownames_to_column( "rowname") %>%
       dplyr::group_by(count_cluster) %>% 
       dplyr::summarise(rowname_str = paste0(str_trim(rowname), collapse = ",")) 
     
     write.table(count_cluster_wide, 
                 file = paste0(res_dir,"/",tit,"_csv_k.",gap_k,"_",set_name,".tsv"), row.names=F, sep="\t")
     assign(paste0("count_cluster"), count_cluster, envir = .GlobalEnv)
     assign(paste0("count_dst"), count_dst, envir = .GlobalEnv)
  
  #browser()   
     anova_cluster <- as.data.frame(adonis2(count_dst ~ count_cluster))
     dst_cluster<- as.data.frame(cbind(dst,count_cluster))
     anova_pair_cluster <- as.data.frame(pairwiseAdonis::pairwise.adonis2(count_dst ~ count_cluster,data=dst_cluster)) %>%
       dplyr::filter(row_number()==1) %>%
       dplyr::select(contains("Pr..F.")) %>%
       t() %>%
       as.data.frame() #%>%
     
     dispr <- vegan::betadisper(count_dst, count_cluster)
     anova_dispr<-as.data.frame(anova(dispr))[1,]$"Pr(>F)"
     dispar_cluster<-merge(as.data.frame(dispr$distances), count_cluster_df, by="row.names")%>%
       mutate(count_cluster= as.factor(count_cluster))
     
     #https://scienceparkstudygroup.github.io/microbiome-lesson/04-alpha-diversity/index.html
     richness <- count_df %>%
       mutate_if(is.numeric,as.integer) %>%
       estimateR() %>%
       as.data.frame() %>%
       rownames_to_column("index") %>%
       dplyr::filter(index=="S.chao1") %>%
       column_to_rownames("index") %>%
       t
     cluster_richness<-merge(count_cluster_df,richness, by="row.names") %>%
       mutate(count_cluster= as.factor(count_cluster))
  #browser()   
     
    #-----------------------------Plot clusters ------------------------
     #gap_k=6
     
     pdf(file = paste0(res_dir,"/",tit,"_k=",gap_k,"_",set_name, ".pdf"))
     f1<-fviz_nbclust(dst, kmeans, method = "wss") +
       #geom_vline(xintercept = 4, linetype = 2)+
       labs(subtitle = "Elbow method")
     # Silhouette method
     f2<-fviz_nbclust(dst, kmeans, method = "silhouette")+
       labs(subtitle = "Silhouette method")
  #browser()
     if (missing(k)){
       f3<-fviz_gap_stat(gap)
       labs(subtitle = "Gap statistic method")  
       print((f1 + f2) / (f3))
     }else{
       print((f1 + f2))
     }
     # Put the labels at the same height: hang = -1
     plot(hclst, cex.axis=0.5,hang = -1, cex = 0.5)
     # draw dendogram with red borders around the clusters
     rect.hclust(hclst, k=gap_k, border="red")
     abline(h = gap_k, col = "blue")
     
     kmeans_cluster <- kmeans(dst, gap_k)
     kmeans_cluster$centers
     print(fviz_cluster(kmeans_cluster, data = dst, labelsize = 5, main = paste0("kmeans k=",gap_k)) )
     
     print(clusplot(dst, kmeans_cluster$cluster, color=TRUE, shade=TRUE, labels=4, lines=0, cex.txt=10))
     print(ggplot( cluster_richness, aes(x=count_cluster, y=S.chao1,fill=count_cluster)) +
       geom_boxplot()+
       geom_jitter(shape=16, position=position_jitter(0.2))+
       stat_compare_means(method = "kruskal.test") +
       ggtitle("Cluster Richness Boxplot"))
     
     print(ggplot( dispar_cluster, aes(x=count_cluster, y=dispr$distances,color=count_cluster)) +
       #geom_boxplot()+
       geom_jitter(shape=16, position=position_jitter(0.2))+
       ggtitle(paste("Cluster Dispar Boxplot anova:",anova_dispr)))
     
     
     t1<-tableGrob(anova_cluster,theme=ttheme_default(base_size = 6))
     t2<-tableGrob(anova_pair_cluster,theme=ttheme_default(base_size = 5))
     grid.arrange(t1, t2, top=textGrob("Distances Anova"))+
       theme_minimal()
  #browser()   
     # pgc<-plot_gene_cor(as.data.frame(dst),set_name=set_name,rectan=gap_k,mth="spearman",
     #                    res_dir=result_dir,element="Taxa",output_format="",sig=F)
  
     dev.off()
   }
   return(count_cluster_df)
 } 
 
 
identify_clusters<- function( scaled_data=physeq_rarefied_scaled_data,
                              rareified_scaled=physeq_rarefied_scaled,
                              data_cluster=gene_cluster_df,
                              fctr_col=factor_col,
                              cond,
                              set_name,
                              element="taxa",
                              corr=F,
                              md=meta,
                              res_dir=result_dir) {
#browser()   
  if (element=="taxa") {
    taxa_lvl<-prep_count_levels(scaled_data,
                                 taxa_level=c("GenusAgg"),
                                 rel=T)
     
     taxa_rarefied_scaled<-as.data.frame(tax_table(rareified_scaled))
     taxa_rarefied_scaled$genus_cluster <-
       unlist(c(taxa_cluster_df[match(taxa_rarefied_scaled$GenusAgg, rownames(taxa_cluster_df)), "count_cluster"]))
     tax_table(rareified_scaled)<-tax_table(as.matrix(taxa_rarefied_scaled))
  } else {
    taxa_count_GenusAgg_rel<-scaled_data
  }
#browser()
  GenusAgg_rel_clustered<-taxa_count_GenusAgg_rel %>%
                       rownames_to_column("GenusAgg") %>%
                       left_join(rownames_to_column(data_cluster,"GenusAgg"), by="GenusAgg") %>%
                       mutate(cluster= as.factor(count_cluster)) %>%
                       dplyr::select(-count_cluster) %>%
                       arrange(cluster) %>%
                       column_to_rownames("GenusAgg")
#browser()  
  if (element=="taxa") {
     clinic_clustered_rel <- GenusAgg_rel_clustered %>%
                       rownames_to_column("GenusAgg") %>%
                       pivot_longer(-c(GenusAgg,cluster),names_to = "Sample", values_to = "rel_abund") %>%
                       left_join(scaled_data, by=c("GenusAgg","Sample")) %>%
                       dplyr::select("cluster", "rel_abund",all_of(fctr_col)) %>%
                       #dplyr::filter(rel_abund!=0) %>%
                       distinct()
  }else{
    clinic_clustered_rel <- GenusAgg_rel_clustered %>%
                      rownames_to_column("GenusAgg") %>%
                      pivot_longer(-c(GenusAgg,cluster),names_to = "Sample2", values_to = "rel_abund") %>%
                      left_join(rownames_to_column(md,"Sample2"), by=c("Sample2")) %>%
                      dplyr::select("cluster", "rel_abund",all_of(fctr_col)) %>%
                      #dplyr::filter(rel_abund!=0) %>%
                      distinct()
  }
   
   clinic_clustered_rel_val <- clinic_clustered_rel %>%
                           mutate_if(is.integer,as.factor)%>%
                           pivot_longer(-c(cluster,rel_abund),names_to = "clinic_var", values_to = "clinic_val") %>%
                           distinct()
#browser()
   clinic_clustered_freq <- clinic_clustered_rel_val %>%
                       dplyr::filter(rel_abund!=0) %>%
                       group_by(cluster, clinic_var, clinic_val) %>%
                       mutate(count = n_distinct(rel_abund)) %>%
                       distinct(cluster, clinic_var, clinic_val, count) %>%
                       group_by(cluster, clinic_var) %>%
                       mutate(freq = count / sum(count)) %>%
                       mutate(clinic_val = as.factor(paste0(clinic_var,";",clinic_val))) %>%
                       #dplyr::select(-"clinic_var") %>%
                       arrange(cluster,clinic_val)
   clinic_clustered_rel_median <- clinic_clustered_rel_val %>%
                       group_by(cluster, clinic_var, clinic_val) %>%
                       mutate(rel_median = median(rel_abund)) %>%
                       distinct(cluster, clinic_var, clinic_val, rel_median) %>%
                       group_by(cluster, clinic_var) %>%
                       #mutate(freq = count / sum(count)) %>%
                       mutate(clinic_val = as.factor(paste0(clinic_var,";",clinic_val))) %>%
                       #dplyr::select(-"clinic_var") %>%
                       arrange(cluster,clinic_val)
   clinic_clustered_rel_val<-clinic_clustered_rel_val%>%
               mutate(clinic_val = as.factor(paste0(clinic_var,";",clinic_val)))
#browser()
#https://rdrr.io/cran/janitor/man/chisq.test.html
  # freq_cond <- tabyl(clinic_clustered_rel_val, clinic_val,cluster) # freq of values in each cluster like table
  # freq_cond <- as.data.frame(freq_cond)%>%
  #   column_to_rownames("clinic_val")
  stat_rel_d_sig<-character()
  stat_freq_d_sig<-character()
  for (cv in unique(clinic_clustered_freq$clinic_var)){
    #cv<-unique(clinic_clustered_freq$clinic_var)[3]
    print(cv)
    # http://www.sthda.com/english/wiki/comparing-proportions-in-r
    clinic_clustered_count_p<-subset(clinic_clustered_freq,clinic_var==cv)  %>%
            dplyr::select(-c(clinic_var,freq)) %>%
            pivot_wider(names_from = "clinic_val",values_from = "count" ) %>%
            replace(is.na(.), 0) %>% 
            column_to_rownames("cluster")
    clinic_clustered_count_p$clinic_var<-NULL
    stat_freq<-chisq.test(clinic_clustered_count_p)
    stat_freq_ph<-as.data.frame(chisq.posthoc.test(clinic_clustered_count_p, method = "bonferroni")) %>% 
                  dplyr::filter(Value=="p values")

    clinic_clustered_rel_p<-subset(clinic_clustered_rel_val,clinic_var==cv) %>%
                          droplevels()
    #counts<-tabyl(clinic_clustered_rel_p, clinic_val,cluster)
    
    stat_rel<-clinic_clustered_rel_p %>% 
      group_by(cluster ) %>% 
      kruskal_test(rel_abund ~ clinic_val)
    # if(nrow(subset(stat_rel,p<=0.05))>0 | cv==cond){
      grid.arrange(textGrob(paste("Stats",cv)),
                   tableGrob(stat_rel,theme=ttheme_minimal(base_size = 6)),
                   tableGrob(clinic_clustered_count_p,theme=ttheme_minimal(base_size = 6)),
                   tableGrob(stat_freq_ph,theme=ttheme_minimal(base_size = 6)),
                   ncol=1)
    # }
    
    stat_rel_d<-clinic_clustered_rel_p %>% 
      group_by(cluster ) %>% 
      dunn_test(rel_abund ~ clinic_val)#%>% 
    
    stat_rel_d_sig<-rbind(stat_rel_d_sig,stat_rel_d%>% dplyr::filter(p.adj<0.1))
  }
   prop.table(table(clinic_clustered_freq$cluster, clinic_clustered_freq$clinic_val), 2) * 100
   prop.table(table(clinic_clustered_rel_median$cluster, clinic_clustered_rel_median$clinic_val), 2) * 100
   
   #-----------4corr of eigengenes==first principal component (eigengene) in each cluster
   trait<-with(subset(md, select=eval(cond)), model.matrix(as.formula(paste0("~",cond," + 0"))))
   rownames(trait) <- rownames(md)
   dim(trait)
   cluster_4cor<- t(subset(GenusAgg_rel_clustered,select=-cluster))
   dim(cluster_4cor)
#browser()   
   cluster_eigengene<- WGCNA::moduleEigengenes(cluster_4cor,GenusAgg_rel_clustered$cluster,scale = F)$eigengenes
   cluster_trait_cor<- cor(cluster_eigengene,trait , use = "p")
   cluster_trait_cor_pval <- corPvalueStudent(cluster_trait_cor, nrow(trait))
   
   #------------------plot
   
   #---------stack bar only cond
#browser()
  if (element=="taxa"){
      b1<-phyloseq::plot_bar(rareified_scaled, fill = "genus_cluster")+
        geom_bar(aes(color = genus_cluster, fill = genus_cluster), stat = "identity", position = "stack") +
       labs(x = "", y = "Relative Abundance\n") +
       #ggtitle(paste(set_name,"Clusters chi:", chi_freq_cond))+
       theme(panel.background = element_blank(),
             axis.text.x=element_blank(),
             axis.ticks.x=element_blank())
     print(b1)
     if (!missing(cond)) print(b1+  facet_wrap(~ get(cond), scales = "free") )
  }
   
   #----------heatmap clinic
   print(ggplot(clinic_clustered_freq, aes(x = cluster, y = clinic_val)) +
           geom_tile(aes(fill = freq), alpha = 0.85)+
           labs(title = "Distribution of  characteristics across clusters-prevalence", x = "Cluster number", y = NULL) +
           scale_fill_gradient2(low = "darkslategray1", mid = "yellow", high = "turquoise4"))
#browser()   
   clinic_clustered_rel_median<- clinic_clustered_rel_median %>%
                                mutate(rel_median_scaled=scale(rel_median))
   print(ggplot(clinic_clustered_rel_median, aes(x = cluster, y = clinic_val)) +
           geom_tile(aes(fill = rel_median_scaled), alpha = 0.85)+
           labs(title = "Distribution of characteristics across clusters-median relative abundance", x = "Cluster number", y = NULL) +
           scale_fill_gradient2(low = "darkslategray1", mid = "yellow", high = "turquoise4"))
   
   print(paste("nrow(stat_rel_d_sig):",nrow(stat_rel_d_sig)))
#browser()
   tp<-table_pdf(stat_rel_d_sig,tit="dunn_test sig")
   
   
   #----------heatmap cor to eigengenes
     
   corrplot::corrplot(cluster_trait_cor, p.mat = cluster_trait_cor_pval, method = "square",#type = "lower",
                                  insig = "label_sig", sig.level = c(.001, .01, .05),
                                  title=paste("\ncorrelation of characteristics with clusters' eigengenes", set_name),
                                  pch.cex = 1.2, pch.col = "black", mar=c(0,0,2,0)) #, order="hclust"
   cluster_trait_cor_4plot<- as.data.frame(cluster_trait_cor) %>%
                               rownames_to_column("ME") %>%
                               mutate(cluster=str_remove(ME,"ME")) %>%
                               dplyr::select(-ME) %>%
                               pivot_longer(-cluster,values_to = "r_cldyer_eigengene", names_to = "clinic_val")
#browser()
  colnames(cluster_trait_cor) <- paste(colnames(cluster_trait_cor), 'r', sep = '_')
  colnames(cluster_trait_cor_pval) <- paste(colnames(cluster_trait_cor_pval), 'p', sep = '_')
  cluster_trait_cor_res<- cbind(cluster_trait_cor,cluster_trait_cor_pval)
   tp<-table_pdf(cluster_trait_cor_res,tit="correlation of characteristics with clusters' eigengenes")
   
   # if (nrow(stat_rel_d_sig)>40){
   #   grid.arrange(tableGrob(stat_rel_d_sig[1:(nrow(stat_rel_d_sig)/2),],theme=ttheme_minimal(base_size = 4)))
   #   grid.arrange(tableGrob(stat_rel_d_sig[(as.integer(nrow(stat_rel_d_sig)/2)+1):nrow(stat_rel_d_sig),],theme=ttheme_minimal(base_size = 4)))
   # } else if (nrow(stat_rel_d_sig)>0){
   #   grid.arrange(tableGrob(stat_rel_d_sig,theme=ttheme_minimal(base_size = 6)))
   # }
   print(ggplot(clinic_clustered_freq, aes(y=freq)) +
           labs(x=NULL, y="Proportion [%]") + theme_bw()+ 
           geom_bar(aes(x=clinic_val, fill=cluster), 
                    position="fill", stat="identity") +
           ggtitle(paste(set,"cluster frequency per clinic")) +
           theme(legend.text=element_text(size=6),
                 axis.text.x = element_text(angle = -30, hjust = 0, vjust=0.5))  +
           guides(fill =guide_legend(ncol=1)) +
           facet_wrap(~clinic_var, scales = "free_x")) #+ 
   
   print(ggplot(clinic_clustered_rel_median, aes(y=rel_median)) +
           labs(x=NULL, y="Median relative abundance") + theme_bw()+ 
           geom_bar(aes(x=clinic_val, fill=cluster), 
                    position="fill", stat="identity") +
           ggtitle(paste(set,"cluster median relative per clinic")) +
           theme(legend.text=element_text(size=6),
                 axis.text.x = element_text(angle = -30, hjust = 0, vjust=0.5))  +
           guides(fill =guide_legend(ncol=1)) +
           facet_wrap(~clinic_var, scales = "free_x")) #+ 
   
   getPalette <- colorRampPalette(brewer.pal(9, "Set1"))
   color_no<-length(unique(clinic_clustered_freq$clinic_val))
   print(ggplot(clinic_clustered_freq, aes(x=cluster,y=freq,group = seq(1, nrow(clinic_clustered_freq)),
                                               fill=clinic_val)) +
           labs(x=NULL, y="Proportion [%]") + theme_bw()+ 
           geom_bar(position="fill", stat="identity") +
           scale_fill_manual(values =getPalette(color_no)[order(sample(1:color_no, color_no))])+
           #scale_fill_manual(values =colorRampPalette(brewer.pal(12, "Accent"))(length(unique(clinic_clustered_freq$clinic_val))))+
           ggtitle(paste(set,"clinic frequency per cluster")) +
           theme(legend.text=element_text(size=6),
                 axis.text.x = element_text(angle = 30))  +
           guides(fill =guide_legend(ncol=2)) +
           facet_wrap(~clinic_var, scales = "free_x")) #+ 
#browser()   
   
   print(ggplot(clinic_clustered_rel_median, aes(x=cluster,y=rel_median,group = seq(1, nrow(clinic_clustered_rel_median)),
                                               fill=clinic_val)) +
           labs(x=NULL, y=ifelse (element=="taxa","Median relative abundance [%]", "Median [%]")) + 
                  theme_bw()+ 
           geom_bar(position="fill", stat="identity") +
           scale_fill_manual(values =getPalette(color_no)[order(sample2(1:color_no, color_no))])+
           #scale_fill_manual(values =colorRampPalette(brewer.pal(12, "Accent"))(length(unique(clinic_clustered_freq$clinic_val))))+
           ggtitle(paste(set,ifelse (element=="taxa","clinic median relative per cluster","clinic median per cluster"))) +
           theme(legend.text=element_text(size=6),
                 axis.text.x = element_text(angle = 30))  +
           guides(fill =guide_legend(ncol=2)) +
           facet_wrap(~clinic_var, scales = "free_x")) #+ 

   if (element=="taxa"){
     distW <- phyloseq::distance(rareified_scaled, method="unifrac", weighted=T) #
     ord <- ordinate(rareified_scaled, method="MDS", distance=distW, weighted=T)
     print(plot_ordination(rareified_scaled, ord, type="taxa", color="genus_cluster", title="taxa")+
             ggforce::geom_mark_ellipse())
   }

   # --------------------- Correlations taxa within cluster
   if (corr) {
     p_thresh<-0.05
  #browser()   
     for (clst in levels(GenusAgg_rel_clustered$cluster)) {
       #clst<-levels(GenusAgg_rel_clustered$cluster)[1]
       print(clst)
       taxa_4corr_mtx_part<- subset(GenusAgg_rel_clustered,cluster==clst,select=-cluster) %>%
                 rownames_to_column("taxa") %>%
                 mutate(taxa=str_trim(taxa),
                        taxa=str_replace_all(taxa,"(\\W)","."))  %>% # handle[]
                 column_to_rownames("taxa") %>%
                 as.matrix() %>%
                 t
       #correlate, default is adjust="holm"
       corr_taxa <- corr.test(taxa_4corr_mtx_part, method="spearman", adjust="BH") 
       
       
       #filter only taxa that passed thresh
       corr_taxa_p <-corr_taxa$p%>%
         as.data.frame%>%
         select_if(~ any(.<=p_thresh))%>%
         filter_all(any_vars(. <=p_thresh))%>%
         as.matrix()
       
       corr_taxa_r<-corr_taxa$r%>%
         as.data.frame%>%
         rownames_to_column("taxa") %>%
         dplyr::filter(taxa %in% rownames(corr_taxa_p))%>%
         dplyr::select(taxa,colnames(corr_taxa_p)) %>%
         column_to_rownames("taxa") %>%
         as.matrix()
       if (ncol(corr_taxa_r)>0){
         qgraph(corr_taxa_r, minimum = 0.25, cut = 0.4,title=paste0("Taxa Network for C",clst))
         # q_df<-centralityTable(q)
         corrplot(corr_taxa_r,p.mat =corr_taxa_p, method = "square", type="upper",
                  title = paste("\n clinic-taxa correlation-subset C",clst,set,proj),
                  insig = "label_sig", sig.level = c(.001, .01, .05), pch.cex = 0.5, pch.col = "black",
                  col = colorRampPalette(c("blue","white","red"))(100), #cex.col = 0.4, 
                  tl.srt=45, tl.col="black",tl.cex = 0.3, tl.offset =0.3,
                  cl.cex=0.5,cl.ratio = 0.1,cl.offset = 0.1, order="hclust")# , addrect = 5
       }
     }
   }
   
   
   return()
}

top_bacteria <- function(phylo, phylo_data, min_samples=30,field,rel=F,specific,spec_colors,PCOA=T,set_name="") {
  
  if (rel){
    taxa_lvl<-prep_count_levels(phylo_data,taxa_level=c("GenusAgg"),rel = T)
    t_count<-taxa_count_GenusAgg_rel
  } else{
    taxa_lvl<-prep_count_levels(phylo_data,taxa_level=c("GenusAgg"))
    t_count<-taxa_count_GenusAgg
  }
  
  # sample_variables(phylo)
  # rank_names(phylo)
  #unique(as.data.frame(tax_table(physeq_rarefied))$GenusAgg)
  if (missing(specific)){
    genus_list<-t_count %>%
              dplyr::filter(rowSums(.>0) > min_samples) %>%
              rownames() 
    pdf(file=paste0(result_dir,"/Top_prevalence_bacteria_PCoA_",field,"_",proj,"_",set,classif, ".pdf"))
  }else{
#browser()
    genus_df<-t_count %>%
              rownames_to_column("taxa") %>%
              dplyr::filter(str_detect(taxa,specific))%>%
              dplyr::select(where(~ any(. != 0))) #%>%
    sample_no<-ncol(genus_df)
    genus_list<-c(genus_df$taxa) 
    pdf(file=paste0(result_dir,"/",specific,"_",sample_no,"_bacteria_PCoA_",field,"_",set_name,"_",proj,"_",set,classif, ".pdf"))
  }
#browser()
  
  genus_list<-str_trim(genus_list)
  print(length(genus_list))
  distW <- phyloseq::distance(phylo, method="unifrac", weighted=T) #
  ord <- ordinate(phylo, method="DPCoA", distance=distW) #, weighted=weight

  for (bctr in genus_list){
    #bctr<-genus_list[4]
    print(bctr)
    bacteria_df<-phylo_data %>%
      dplyr::filter(str_detect(GenusAgg,str_replace_all(bctr,"(\\W)","\\\\\\1"))) %>% # deal with []
      dplyr::select(Abundance,Sample,cond=eval(field)) %>%
      group_by(Sample,cond) %>%
      dplyr::summarise(Abundance = sum(Abundance))  %>%
      arrange(rownames(sample_data(phylo)),cond)
    #setdiff(bacteria_df$Sample,rownames(sample_data(phylo)))
    #setdiff(rownames(sample_data(phylo)),bacteria_df$Sample)
    #colnames(otu_df) = meta_16S[match(colnames(otu_df), meta_16S$Sample), "sample_name"]
    sample_data(phylo)$bacteria <- unlist(c(bacteria_df[match(rownames(sample_data(phylo)), bacteria_df$Sample), "Abundance"]))
    #sample_variables(phylo)
#browser()
    print(ggplot( bacteria_df, aes(x=cond, y=Abundance,color=cond)) +
            geom_boxplot()+
            ylab(paste(bctr,ifelse(rel,"Relative",""),"Abundance"))+
            theme(axis.text.x = element_text(angle = 90))+
            geom_point()+
            {if(!missing(spec_colors)) scale_color_manual(values = spec_colors)} +
            ggtitle(paste(ifelse(rel,"Relative",""),"Abundance",field, set_name,"boxplot",bctr))+
            theme(panel.grid.major = element_blank(), 
                  panel.grid.minor = element_blank(),
                  panel.background = element_blank(), 
                  # axis.text.x = element_text(angle = 30),
                  axis.line = element_line(colour = "black"),
                  text=element_text(size = 12,  family="sans") )+ 
            scale_y_continuous(expand = expansion(mult = c(0,.2)))+
            stat_compare_means(method = "kruskal.test",paired = TRUE,size=5))
    
    print(plot_ordination(phylo, ord, color="bacteria",shape=eval(field) )+
            #scale_fill_gradient(aes(fill=sample_data(phylo)$bacteria))+
            scale_shape_manual(values=1:nlevels(meta_16S[[field]])) +
            guides(color=guide_legend(title=paste("%",bctr))))#+ 
    #geom_text(mapping = aes(label = Sample), size = 2)) #+
  }
  dev.off()
  return(genus_list)
}
 
table_pdf <- function( df,total_rows_per_page = 30,start_row = 1,tit="" ) {
   
  
  
  if(total_rows_per_page > nrow(df)){
    end_row = nrow(df)
  }else {
    end_row = total_rows_per_page 
  }    
  
  for(i in 1:ceiling(nrow(df)/total_rows_per_page)){
    
    #grid.newpage()   
    
    grid.arrange(tableGrob(df[start_row:end_row, ],theme=ttheme_minimal(base_size = 6)),
                 top=textGrob(tit))#, show.rownames = FALSE
    
    start_row = end_row + 1
    
    if((total_rows_per_page + end_row) < nrow(df)){
      
      end_row = total_rows_per_page + end_row
      
    }else {
      
      end_row = nrow(df)
    }    
  }
}

Adiver_vegan<- function(wb_in=wb, count_df,md, field1,mtd="shannon",set_name=""){
#browser()
  shannon_div <- data.frame(shannon_di=vegan::diversity(t(count_df),index=mtd)) %>%
                  rownames_to_column("sample_name2") %>%
                  left_join(rownames_to_column(md,"sample_name2"),by="sample_name2")  %>%
                    dplyr::select(sample_name2,shannon_di,field1=eval(field1)) #%>%
    
  
  stat_div <- kruskal_test(formula =shannon_di ~ field1, data=shannon_div) 
  pairwise.wilcox_a <- pairwise.wilcox.test(shannon_div$shannon_di,shannon_div$field1,p.adjust.method="none" ) 
  a_posthoc_sig<- as.data.frame(pairwise.wilcox_a$p.value) %>%
                    rownames_to_column("set1") %>%
                    pivot_longer(-set1,names_to = "set2",values_to = "p.val") %>%
                    dplyr::filter(!is.na(p.val) & p.val<0.1)
  
  print(ggplot(shannon_div, aes(x=field1, y=shannon_di, color=field1)) +
          xlab(field1) +
          ggtitle(paste(mtd,"diversity", proj,set,set_name,
                        "\nStats",field1, stat_div$p[1],
                        "\nPosthoc sig",paste(paste0(a_posthoc_sig$set1,"-",a_posthoc_sig$set2),collapse= ";"))) +  
          geom_boxplot()+
          geom_point(position = position_dodge(width = 0.5))+
          theme_bw() + 
          theme(axis.text.x = element_text(angle = 90),
                text = element_text(size = 9))) #+
  posStyle <- createStyle(fontColour = "#006100", fgFill = "#C6EFCE")
  s <- createStyle(numFmt = "$ #,##0.00")
  #m<-"Observed"
  titStyle <- createStyle(fontColour = "#006100", fgFill = "#C6EFCE")
  sheet_a_anova <- paste0("Adiv ",set_name," ",field1)
  addWorksheet(wb_in, sheet_a_anova)
  writeData(wb_in,sheet_a_anova,field1, startRow= 1) 
  addStyle(wb_in, sheet_a_anova, style = titStyle, cols=1:10, rows=1:1, gridExpand = TRUE)
  #browser()
  writeData(wb_in,sheet_a_anova, stat_div, colNames = T, rowNames= T) 
  writeData(wb_in,sheet_a_anova, "pairwise.wilcox.test", colNames = T, rowNames= T, startCol = 10) 
  writeData(wb_in,sheet_a_anova, pairwise.wilcox_a$p.value, colNames = T, rowNames= T, startCol = 10, startRow = 2) 
  
}
                  
samples_cluster<-function(count_df, md, fld) {
  #-----hclust clustering
  #fld <- "Condition"
#browser()
  count_df_logged <- t(log10(count_df+1)) #%>% #log transform data.
  
  count_dst <- vegdist( count_df_logged)
  dst <- as.matrix(count_dst)
  hclst<-hclust(count_dst, method="ward.D2")
  summary(hclst)
  ddata <- dendro_data(hclst, type="rectangle")
  ddata$labels <- merge(ddata$labels, rownames_to_column(md,"label"))
#browser()
  print(ggplot() +
    geom_segment(data = segment(ddata), aes(x=x, y=y, xend=xend, yend=yend))+
    geom_segment(data = ddata$segments %>%
                   filter(yend == 0) %>%
                   left_join(ddata$labels, by = "x"), aes(x=x, y=y.x, xend=xend, yend=yend, color = .data[[fld]])) +
    geom_text(data = label(ddata), aes(x=x, y=y, label=label, colour = .data[[fld]], hjust=0), size=2.5) +
    scale_y_reverse(expand=c(0.2, 0)) +
    theme_dendro() +
    coord_flip() +
    ggtitle(paste(proj,set,run,"- bray Dissimilarity ward.D2 Clustering of samples (Colored by",fld,")")))
}


 separate_paires <- function( md,taxa_count,res_for) {
   list1<-rev(levels(md[[res_for]]))
   print(list1)
   list2<-list1
   res_all<-character()
   res_all_df<-character()
   for (r1 in list1){
     list2<-list2[list2 != r1]
     for (r2 in list2){
       if (r1!=r2){
browser()   
         md_part<-md %>%
                  dplyr::filter(get(res_for)==r1 | get(res_for)==r2)
         taxa_count_part<-taxa_count %>%
                  dplyr::select(rownames(md_part)) #%>% # filter zero rows?
         assign(paste0("taxa_count_",R1,"_",), taxa_count_part, envir = .GlobalEnv)
         
       } #if (r1!=r2)
     } #for (r2 in list2)
   } #for (r1 in list1)
 }
   
 
 metaphlanToPhyloseq <- function(
   tax,
   metadat=NULL,
   simplenames=TRUE,
   roundtointeger=FALSE,
   split="|"){
   ## tax is a matrix or data.frame with the table of taxonomic abundances, rows are taxa, columns are samples
   ## metadat is an optional data.frame of specimen metadata, rows are samples, columns are variables
   ## if simplenames=TRUE, use only the most detailed level of taxa names in the final object
   ## if roundtointeger=TRUE, values will be rounded to the nearest integer
   xnames = rownames(tax)
   shortnames = gsub(paste0(".+\\", split), "", xnames)
   if(simplenames){
     rownames(tax) = shortnames
   }
   if(roundtointeger){
     tax = round(tax * 1e4)
   }
   x2 = strsplit(xnames, split=split, fixed=TRUE)
   taxmat = matrix(NA, ncol=max(sapply(x2, length)), nrow=length(x2))
   colnames(taxmat) = c("Kingdom", "Phylum", "Class", "Order", "Family", "Genus", "Species", "Strain")[1:ncol(taxmat)]
   rownames(taxmat) = rownames(tax)
   for (i in 1:nrow(taxmat)){
     taxmat[i, 1:length(x2[[i]])] <- x2[[i]]
   }
   #taxmat = gsub("[a-z]__", "", taxmat)
   #taxmat = phyloseq::tax_table(taxmat)
   #otutab = phyloseq::otu_table(tax, taxa_are_rows=TRUE)
   colnames(tax)[match(metadat$BarcodeID, str_split_fixed(colnames(tax),"_",3)[,2])] = rownames(metadat)
   assign("Taxa" , taxmat,envir = .GlobalEnv)
   
#browser()
   TAX = tax_table(as.matrix(taxmat))
   OTU = otu_table(as.matrix(tax), taxa_are_rows = TRUE)
   META <- sample_data(metadat)
   head(sample_names(OTU))
   head(sample_names(META))
   
   if(is.null(metadat)){
     res = phyloseq::phyloseq(taxmat, otutab)
   }else{
     res = phyloseq::phyloseq(TAX, OTU, META)
   }
   return(res)
 }
 
 
 #' Centered log-ratio transformation
 #'
 #' @param m Count matrix with features in rows and samples (cells) in columns
 #' @return matrix
 #' @export
 clr <- function(m) {
   apply(m, 2, function(x) {
     log1p(x = x/(exp(x = sum(log1p(x = x[x > 0]), na.rm = TRUE)/length(x = x))))
   })
 }
 
 
 
```
