## Supplementary material for "An experimental approach towards untangling the role of nature versus nurture in shaping the microbiome of social insects": R code: RNAseq_functions.R

```
###########################################RNAseq_functions.R#########################
my_plot_PCA <- function(dds, set_name,intgroup, be,res_dir=DE_result_dir, out="jpg"){
  
  vstcounts_exp <- varianceStabilizingTransformation (dds, blind=TRUE)
  print(DESeq2::plotPCA(vstcounts_exp, intgroup=intgroup) + 
          labs(title=paste(set_name,'PCA'))+
          geom_label_repel(aes(label = dds@colData@rownames), box.padding   = 0.35, point.padding = 0.5, segment.color = 'grey50'))
  if (out=="jpg"){
    dev.copy(jpeg,filename=paste0(res_dir,"/PCA_",set_name, ".jpg"), 
             width=1000, height=1200)
  } else{
    dev.copy2pdf(device = quartz, file =paste0(res_dir,"/PCA_",set_name, ".pdf"))
  }
  dev.off()
  
  if (!missing(be)){
    print(paste("be:",be))
    # remove batch effect
    assay(vstcounts_exp) <- 
        limma::removeBatchEffect(assay(vstcounts_exp), vstcounts_exp@colData[,be])
    print(DESeq2::plotPCA(vstcounts_exp, intgroup=intgroup) + 
            labs(title=paste(set_name,'PCA- BE removed')) +
            geom_label_repel(aes(label = dds@colData@rownames), box.padding   = 0.35, point.padding = 0.1, segment.color = 'grey50'))
    if (out=="jpg"){
      dev.copy(jpeg,filename=paste0(res_dir,"/PCA_nbe_",set_name,".jpg"), 
             width=1000, height=1200)
    } else{
      dev.copy2pdf(device = quartz, file =paste0(res_dir,"/PCA_nbe_",set_name, ".pdf"))
    }
    dev.off()
    return(vstcounts_exp)
  }
}

plot_samples_dist_heatmap <- function(dds, vst_nbe, set_name,res_dir=DE_result_dir){
  
  colors <- colorRampPalette( rev(brewer.pal(9, "Blues")) )(255)
  
  vst <- varianceStabilizingTransformation (dds, blind=FALSE)
  sampleDists <- DESeq2::dist(t(assay(vst)))
  
  pheatmap(as.matrix(sampleDists),
           clustering_distance_rows=sampleDists,
           clustering_distance_cols=sampleDists,
           col=colors, main="Samples Distance VST Heatmap")
  dev.copy(jpeg,filename=paste0(res_dir,"/heatmap_",set_name,"_vst_samples_dists", ".jpg"), 
           width=1000, height=1200)
  dev.off()
  
  # remove batch effect
  if (!missing(vst_nbe)){
    print("vst_nbe")
    sampleDists <- dist(t(assay(vst_nbe)))
    
    pheatmap(as.matrix(sampleDists),
             clustering_distance_rows=sampleDists,
             clustering_distance_cols=sampleDists,
             col=colors, main="Samples Distance VST Heatmap - batch effect removed")
    dev.copy(jpeg,filename=paste0(res_dir,"/heatmap_",set_name,"_vst_samples_dists_nbe", ".jpg"), 
             width=1000, height=1200)
    dev.off()
    
    # vst is now the normalized log2-transformed data
    pheatmap(cor(assay(vst_nbe)), main="Samples Correlation VST Heatmap - batch effect removed")
    dev.copy(jpeg,filename=paste0(res_dir,"/heatmap_",set_name,"_vst_samples_cor_nbe", ".jpg"), 
             width=1000, height=1200)
    dev.off()
    
  }
 # rld <- rlog(dds)
#  sampleDists <- dist( t( assay(rld) ) )
#  pheatmap(as.matrix(sampleDists),
#           clustering_distance_rows=sampleDists,
#           clustering_distance_cols=sampleDists,
#           col=colors, main="Samples Distance RLD Heatmap")
#  dev.copy(jpeg,filename=paste0(DE_result_dir,"/heatmap_",set_name,"_rld_samples_dists", ".jpg"), 
#           width=800, height=1000)
#  dev.off()
  
  
}

plot_genes_heatmap <- function(rld,
                               anno_df = NA,
                               set_name="",
                               anno_row,
                               cluster, 
                               dbg = TRUE,
                               cols_dend = TRUE, 
                               res_dir=DE_result_dir,
                               y_name="genes", 
                               gap_c = NULL, 
                               gap_r = NULL,
                               set_colnames=TRUE,
                               special_color,
                               an_colors,
                               scl="row",
                               plot_title,
                               font_size=8,
                               out_format="jpeg"){
  if (missing(plot_title)) plot_title<-set_name
  #colors<-colorRampPalette(rev(heat.colors(50)))(255)
  suerat_palette<- Custom_palette(low = "#313695" , high = "#A50026", mid = "#FFFFBF", k = 50)
  suerat_palette2<- Custom_palette(low = "magenta", high = "yellow", mid = "black", k = 50)
  suerat_palette3<- Custom_palette(low = "white", high = "black", mid = NULL, k = 50)
  my_palette <- colorRampPalette(c("blue", "white", "red"))(n = 50)
  my_colors<-colorRampPalette(rev(brewer.pal(n = 8, name="RdYlBu")))(100)
#browser()
  gene_no <-nrow(rld)
  sample_no <-ncol(rld)
  if (!missing(special_color)){my_colors=special_color}
  if (dbg) print(paste("out_format:",out_format))
  if (out_format=="pdf"){

    w<-max((sample_no*0.25)+str_length(rownames(rld))*0.3)
    h<-(gene_no*0.05)+1.50
    pdf(file=paste0(res_dir,"/heatmap_",set_name,"_", y_name,".pdf"),width=w, height=h)  
    }
  #anno_df <- as.data.frame(colnames(rld))
#browser()
  if (missing(anno_row)){
    if (!cols_dend){
      if (dbg) print("missing anno_row, cols_dend=F")
      if (dbg) print(paste("gap_c:",gap_c,"gap_r",gap_r))
      any(is.na(rld) )
      rld[sapply(rld, is.infinite)]
      rld[sapply(rld, is.nan)]
#browser()
      p_title<-paste(plot_title,y_name,"(",nrow(rld),")")
      pheatmap(rld, fontsize=font_size,annotation_col=anno_df, scale = scl,
               fontsize_number = 0.6 * fontsize, col=my_palette, fontsize_row = font_size*0.8,
               gaps_col =  gap_c,gaps_row =  gap_r,
               cluster_cols = FALSE, show_colnames= set_colnames,
               main = p_title)
      #heatmap.2(as.matrix(rld),scale="row", 
      #          trace="none", Rowv=T, Colv=F ,col= colors,
      #          main=paste(plot_title,y_name,nrow(rld),")-zscore"))
      
    }else{
      if (dbg) print("missing anno_row, cols_dend=T")
#browser()
      p_title<-paste(plot_title,y_name,"(",nrow(rld),")")
      if (missing(an_colors)){
        # anno_colors = rainbow(length(anno_df))
        pheatmap(rld, fontsize=font_size,annotation_col=anno_df, scale = scl,
                 fontsize_number = 0.5 * fontsize, fontsize_row = font_size*0.8,col=my_colors,
                 main = p_title)
      } else  {
          pheatmap(rld, fontsize=font_size,annotation_col=anno_df, scale = scl, annotation_colors = an_colors,
                   fontsize_number = 0.5 * fontsize, fontsize_row = font_size*0.8,col=my_colors,
                   main = p_title)
      }
    }
      
  } else if (missing(cluster)){
    if (dbg) print("anno_row, no cluster")
#browser()
    p_title<-paste(plot_title,y_name,"(",nrow(rld),")")
    rld <- rld[rownames(anno_row), ]
    pheatmap(rld, annotation_col=anno_df, scale = scl,
             annotation_row=anno_row, cluster_rows = F,
             fontsize=8,fontsize_row = 5, fontsize_number = 0.5*fontsize,fontsize_col = 8,
             col=my_colors, cluster_cols = cols_dend, show_colnames= set_colnames,
             main = p_title)
  } else {
    print("anno_row+cluster")
    if (missing(an_colors)){
      anno_colors = rainbow(length(cluster))
      names(anno_colors) <- cluster
      anno_colors <- list(FlowSOM = anno_colors)}
    else  {
      names(an_colors) <- cluster
      anno_colors <- list(FlowSOM = an_colors)
    }
    
    print(anno_colors)
    if (cols_dend){
      anno_row<-anno_row %>% arrange(across())
      rld<-rld[match(rownames(anno_row), rownames(rld)), ]
#browser()
      if (dbg) print("cols_dend=T")
      p_title<-paste(plot_title,set_name,y_name,"(",nrow(rld),")")
      pheatmap(rld, fontsize=font_size, 
             scale = scl,annotation_col=anno_df,
             annotation_row=anno_row, cluster_rows = FALSE, annotation_colors = anno_colors,
             fontsize_number = 0.6 * font_size, fontsize_row = font_size, col=my_colors, show_colnames= set_colnames,
             main = p_title)
    } else{
      #order by column name b/a
      if (dbg) print("cols_dend=F")
      #rld<-rld[ ,order(substr(names(rld),6,6))]
      rld<-rld[match(rownames(anno_row), rownames(rld)), ]
      p_title<-paste(plot_title,set_name,y_name,"(",nrow(rld),")")
      pheatmap(rld,  
               scale = scl,annotation_col=anno_df,cluster_cols = FALSE,
               annotation_row=anno_row, cluster_rows = FALSE, annotation_colors = anno_colors,
               fontsize=font_size,fontsize_row = 0.6 * font_size, fontsize_number = 0.6 * font_size,fontsize_col = font_size,
               gaps_col =  gap_c, gaps_row =  gap_r, # border_color = "black",
               #cellheight = 10, cellwidth = 20,
               col=my_colors, show_colnames= set_colnames,
               main = p_title)
    }
  }
#browser()
  if (out_format=="jpeg"){
    if (gene_no<20){
      wid<-max((sample_no*25)+str_length(rownames(rld))*3,str_length(p_title))*3
      dev.copy(jpeg,filename=paste0(res_dir,"/heatmap_",set_name,"_", y_name,".jpg"), 
               width=wid, height=(gene_no*30)+300)
    }else if (gene_no<50){
      wid<-max((sample_no*25)+str_length(rownames(rld))*5,str_length(p_title))*2
      dev.copy(jpeg,filename=paste0(res_dir,"/heatmap_",set_name,"_", y_name,".jpg"), 
               width=wid, height=(gene_no*10)+170)
    }else if (gene_no<250){
      wid<-max((sample_no*28)+str_length(rownames(rld))*5,str_length(p_title))*2
      dev.copy(jpeg,filename=paste0(res_dir,"/heatmap_",set_name,"_", y_name,".jpg"), 
               width=wid, height=(gene_no*14)+150)
    }else {
      wid<-max((sample_no*45)+str_length(rownames(rld))*5,str_length(p_title))*4
      dev.copy(jpeg,filename=paste0(res_dir,"/heatmap_",set_name,"_", y_name,".jpg"), 
               width=wid, height=(gene_no*14)+150)
    }
  }
  if (out_format!="")   dev.off()
}

plot_sample_cor <- function(df,set_name,rectan=2,corr_mth="spearman",mtd="bray",
                            res_dir=DE_result_dir,show_sample_name=T,anno_df,
                          output_format="jpg"){
#browser()
  df_t<-df %>%
        t %>%
        as.data.frame(stringsAsFactors=FALSE)#%>%
  print(paste("dim df:",dim(df_t)))
#browser()
  sample_dist <- vegdist(df_t,method = mtd, na.rm =T)
  # sample_dist[lower.tri(sample_dist)] <- NA
  
  colors <- colorRampPalette( rev(brewer.pal(9, "Blues")) )(255)
  if (missing(anno_df)) {
    pheatmap(as.matrix(sample_dist),
             clustering_distance_rows=sample_dist,
             clustering_distance_cols=sample_dist,
             show_colnames=show_sample_name,
             show_rownames=show_sample_name,
             # na_col="white",
             # border_color ="white",
             col=colors, main=paste("Samples Distance Heatmap",set_name))
  } else {
    pheatmap(as.matrix(sample_dist),
             clustering_distance_rows=sample_dist,
             clustering_distance_cols=sample_dist,
             show_colnames=show_sample_name,
             show_rownames=show_sample_name,
             annotation_col=anno_df,   
             col=colors, main=paste("Samples Distance Heatmap",set_name))
  }
  
  print("before corr")
  
  
  
  # cor_mat <- corr.test(df_t, method=corr_mth)
  # if (rectan==0){
  #     corrplot::corrplot(cor_mat_sig$r, p.mat = cor_mat_sig$p, method = "square",type = "lower",
  #                        insig = "label_sig", sig.level = c(.001, .01, .05),
  #                        title=paste("\nsample correlation", set_name),
  #                        pch.cex = .4, pch.col = "white", order="hclust")
  # }else {
  #     corrplot::corrplot(cor_mat_sig$r, p.mat = cor_mat_sig$p, method = "square",#type = "lower",
  #                        insig = "label_sig", sig.level = c(.001, .01, .05), tl.cex=.5,
  #                        title=paste("\nsample correlation", set_name),
  #                        pch.cex = ., pch.col = "white", order="hclust", addrect = rectan)
  # }
  if (output_format=="jpg"){
    w<-ncol(df)*30+max(str_length(rownames(df)))
    print(w)
    dev.copy(jpeg,filename=paste0(res_dir,"/correlation_",mth,"_",set_name, ".jpg"), 
             width=w, height=w)
    dev.off()
  }
}


plot_volcano <- function(res,set_name, res_dir=DE_result_dir, labeled=TRUE, element="gene",
                         lfc=1, taxa=taxa_16S_df, label_list,label_list_name="pos",
                         label_list2, label_list2_name="neg",
                         output_format="jpg",vtitle, padj_thresh=0.05,
                         quantile_val=0){
  #res<-res_treat
  #set_name<-proj
  #lfc=1
  #label_list<-spec_genes
  #res_dir<-DE_result_dir
  print(paste("output_format:",output_format))
  if(missing(vtitle)) vtitle<-set_name
  res_4plot <- as.data.frame(res) %>% 
    rownames_to_column(var="gene") %>%    #save rowname as col
    mutate(genelabels = "") %>%           #add empty col
    arrange(pvalue) %>%                   #order by pvalue
    mutate(sig = pvalue <= 0.05 & abs(log2FoldChange) >= lfc) # add logical vector where TRUE values

  # add short name to OTUs
  sig_no <- nrow(subset(res_4plot,sig == TRUE))
#browser()  
  if (!labeled & missing(label_list)) {
    print("!labeled & missing(label_list)")
    print(ggplot(res_4plot) +
            geom_point(aes(x = log2FoldChange, y = -log10(pvalue), colour = sig)) +
            ggtitle(paste(set, set_name,"\n significant","(LFC>=",lfc,")",element,"number:",sig_no)) +
            xlab("log2 fold change") + 
            ylab("-log10 p-value") +
            geom_text_repel(aes(x = log2FoldChange, y = -log10(pvalue),label = genelabels)) +
            theme(legend.position = "none",
                  text = element_text(size = 12),
                  plot.title = element_text(size = rel(1.5), hjust = 0.5),
                  axis.title = element_text(size = rel(1.25)))  )
    if (output_format=="jpg"){
      dev.copy(jpeg,filename=paste0(res_dir,"/volcano_sig_","_",set_name,".jpg"), 
               width=600, height=600)
      dev.off()
    }
  } else {
    print(paste("no of sig:", sig_no))
    if (missing(label_list)) {
#browser()
      res_4plot<- res_4plot %>%
                    mutate(genelabels=ifelse(res_4plot$padj<=padj_thresh & abs(log2FoldChange) >= lfc,res_4plot$gene,NA))
      #res_4plot$genelabels <- ifelse(res_4plot$padj<=padj_thresh & abs(log2FoldChange) >= lfc,res_4plot$gene,NA)
      #res_4plot$genelabels <- ifelse(res_4plot$padj<=0.05,res_4plot$gene,NA)																			
      padj_no<-nrow(subset(res_4plot,!is.na(res_4plot$genelabels)))
      print(paste("no of genes padj<=",padj_thresh,":",padj_no))
    } else {
      if (missing(label_list2)){
        res_4plot<-res_4plot %>%
                    mutate(genelabels = ifelse(gene %in% label_list,gene,NA)) %>%
                    mutate(sig = ifelse(gene %in% label_list,"spec",sig)) %>%
                    arrange(sig,genelabels)
      } else {
        res_4plot<-res_4plot %>%
                    mutate(genelabels = ifelse(gene %in% label_list,gene,
                                               ifelse(gene %in% label_list2,gene,NA))) %>%
                    mutate(sig = ifelse(gene %in% label_list,label_list_name,
                                        ifelse(gene %in% label_list2,label_list2_name,sig))) %>%
                    arrange(sig,genelabels)
      }

    }   
#browser()
    gg<-ggplot(res_4plot) +
        geom_point(aes(x = log2FoldChange, y = -log10(pvalue), colour = sig))  + 
        scale_color_manual(values=c("#b5b5b5","#CA2259","#258CB5","#25b531"))+
        #scale_color_manual(values=c("#999999", "#d10420", "#56B4E9"))+
        xlab("log2 fold change") + 
        ylab("-log10 p-value") +
        #geom_label_repel(aes(x= log2FoldChange, y= -log10(pvalue),label = genelabels),
        #                 min.segment.length= 1, max.overlaps=100) +
        theme(#legend.position = "top",
          plot.title = element_text(size = rel(1.3), hjust = 0.5),
          axis.title = element_text(size = rel(1.2)),
          panel.grid.major = element_blank(), 
          panel.grid.minor = element_blank(),
          panel.background = element_blank(), 
          axis.line = element_line(colour = "black"),
          text=element_text(size = 8,  family="sans") ) 
    
    if (labeled){
      print ("labled")
		gg<-gg +
		  geom_label_repel(aes(x= log2FoldChange, y= -log10(pvalue),label = genelabels),size = 1.8,box.padding=0.1,
							   min.segment.length= 1, max.overlaps=100) 
    } else {
      print ("lable_list but not labled")
     
    }
    if (output_format=="pdf") {
      res_4plot<-dplyr::filter(res_4plot,log2FoldChange!=min(log2FoldChange))
      gg<-ggplot() +
        geom_point(data = res_4plot[res_4plot$sig != 'spec',], aes(x = log2FoldChange, y = -log10(pvalue), colour = sig), size = 2)  + 
        geom_point(data = res_4plot[res_4plot$sig == 'spec',], aes(x = log2FoldChange, y = -log10(pvalue), colour = sig), size = 3)  + 
        scale_color_manual(values=c("#b5b5b5","#CA2259","#258CB5"))+
        xlab("log2 fold change") + 
        ylab("-log10 p-value") + xlim(c(-6,6)) +
        geom_text_repel(data = res_4plot[res_4plot$sig == 'spec',], aes(x = log2FoldChange, y = -log10(pvalue),label = gene), 
                        min.segment.length = unit(0, 'lines'),max.overlaps=Inf,
                        point.padding = 0.5) +
        theme(legend.position = "none",
              plot.title = element_text(size = rel(1.5), hjust = 0.5),
              axis.title = element_text(size = rel(1.25)),
              panel.grid.major = element_blank(), 
              panel.grid.minor = element_blank(),
              panel.background = element_blank(), 
              axis.line = element_line(colour = "black"),
              text=element_text(size = 16,  family="sans"))
      
      
      ggsave(paste0(res_dir,"/volcano_labeled_sig_",set_name,".pdf"), plot = gg,
             width = 7.5, height = 5, device = "pdf", dpi=300, units="in")
      #save(res_4plot, file=paste0(home_dir,"/volcano_labeled_4Shlomi", ".Rdata"))
      
    }  else  {
#browser()
      if (missing(label_list)) {
        print(gg +
                ggtitle(paste(set, vtitle, "\n significant ","(LFC>=",lfc,") number:",sig_no,
                              "\n Label on PADJ<= ",padj_thresh,":",padj_no)))
        if (output_format=="jpg"){
          dev.copy(jpeg,filename=paste0(res_dir,"/volcano_labeled_sig_",set_name, ".jpg"), 
                   width=800, height=1000)
          dev.off()
        }
      }else{
        if (missing(label_list2)) {
          print(gg +
                  ggtitle(paste(set, vtitle, "\n significant ","(LFC>=",lfc,") number:",sig_no,
                                "\n Label on Specific genes (",length(label_list),")")))
        }else {
          print(gg +
                  ggtitle(paste(set, vtitle, "\n significant ","(LFC>=",lfc,") number:",sig_no,
                                "\n Label on positive genes (",length(label_list),"), \nnegative (",length(label_list2),")")))
          
        }
        if (output_format=="tiff"){
          ggsave(paste0(res_dir,"/volcano_labeled_spec_",set_name, ".tif"), width = 6, height = 5, device = "tiff", dpi=300, units="in")
        }else if (output_format=="jpg"){
    			dev.copy(jpeg,filename=paste0(res_dir,"/volcano_labeled_spec_",set_name, ".jpg"), 
    					 width=800, height=1000)
    			dev.off()
        }
      }
      
      
    }   
    
  }
}

plot_k <- function(data,set,max_k=40, res_dir=DE_result_dir){
  #elbow plot to Determining Optimal Clusters
  
  set.seed(123)
  # Set maximum cluster 
  data_scaled <- scale(data, center = TRUE, scale = TRUE)
#browser()
  wss <- (nrow(data_scaled)-1)*sum(apply(data_scaled,2,var))
  for (i in 2:max_k) wss[i] <- sum(kmeans(data_scaled,
                                          centers=i)$withinss)
  plot(1:max_k, wss, type="b", xlab="Number of Clusters",
       ylab="Within groups sum of squares")
  dev.copy(jpeg,filename=paste0(res_dir,"/elbow_",set, ".jpg"), 
           width=1000, height=700)
  dev.off()
  
}


plot_pval_hist <- function(res,set_name){
  
  colori <- c(`not passed`="khaki", `passed`="powderblue")
  
  use <- res$baseMean > metadata(res)$filterThreshold
  h_no_use <- hist(res$pvalue[!use], breaks=0:50/50, plot=FALSE)
  h_use <- hist(res$pvalue[use], breaks=0:50/50, plot=FALSE)

  barplot(height = rbind(h_no_use$counts, h_use$counts), beside = FALSE,
          col = colori, space = 0, ylab="frequency",
          main = paste("p-values results of DEseq  threshold:",
                       round(metadata(res)$filterThreshold,digit=2),
                       "\n passed:",length(res$pvalue[use]),
                       "not passed" ,length(res$pvalue[!use]) )) 
  text(x = c(0, length(h_no_use$counts)), y = 0, label = paste(c(0,1)), adj = c(0.5,1.7), xpd=NA)
  legend("top", fill=rev(colori), legend=rev(names(colori)))
  dev.copy(jpeg,filename=paste0(DE_result_dir,"/pval_hist_",set_name,".jpg"), 
           width=600, height=600)
}

plot_gene_cor <- function(df,set_name,rectan=2,mth="spearman",res_dir=DE_result_dir,element="genes",
                          sig=T,output_format="jpg"){
  #df<-spec_mat
  #filter out some not interesting genes
  df<-df %>%
        rownames_to_column("gene") %>%
        dplyr::filter(!str_detect(gene,"^Gm") & !str_detect(gene,"Rik$")) %>%
        column_to_rownames("gene") %>%
        t %>%
        as.data.frame(stringsAsFactors=FALSE)#%>%
  print(paste("nrow df:",nrow(df)))
  cor_mat_sig <- corr.test(df, method=mth)
  if (rectan==0){
    if(sig){
      corrplot::corrplot(cor_mat_sig$r, p.mat = cor_mat_sig$p, method = "square",#type = "lower",
                         insig = "label_sig", sig.level = c(.001, .01, .05),
                         title=paste("\nsignificant",element,"correlation", set_name),
                         pch.cex = .4, pch.col = "white", order="hclust")
    }else{
      corrplot::corrplot(cor_mat_sig$r,  method = "square",#type = "lower",p.mat = cor_mat_sig$p,
                         # insig = "label_sig", sig.level = c(.001, .01, .05),pch.cex = .4, pch.col = "white",
                         title=paste("\nsignificant",element,"correlation", set_name),
                          order="hclust")
    }
  }else {
    if(sig){
      corrplot::corrplot(cor_mat_sig$r, p.mat = cor_mat_sig$p, method = "square",#type = "lower",
                         insig = "label_sig", sig.level = c(.001, .01, .05), tl.cex=.5,
                         title=paste("\nsignificant",element,"correlation", set_name),
                         pch.cex = ., pch.col = "white", order="hclust", addrect = rectan)
    }else{
      corrplot::corrplot(cor_mat_sig$r,  method = "square",#type = "lower",p.mat = cor_mat_sig$p,
                         # insig = "label_sig", sig.level = c(.001, .01, .05), pch.cex = .4, pch.col = "white",
                         tl.cex=.2,
                         title=paste("\nsignificant",element,"correlation", set_name),
                         order="hclust", addrect = rectan)
    }
  }
  if (output_format=="jpg"){
    w<-ncol(df)*30+max(str_length(rownames(df)))
    print(w)
    dev.copy(jpeg,filename=paste0(res_dir,"/correlation_sig_",element,"_",mth,"_",set_name, ".jpg"), 
             width=w, height=w)
    dev.off()
  }
}

#---------------------------------------all vs CTRL 
all_vs_ref <- function(ref,ref_col,design_formula, set_name,md=meta,count_data=sample_data,all_set=NULL,
                       res_dir=home_dir){
  #browser()  
  dds <- DESeqDataSetFromMatrix(countData = count_data,
                                colData = md,
                                design= design_formula ) 
  
  #reference on CTRL
  dds[[ref_col]] <- relevel(dds[[ref_col]], ref = ref) 
  dds <- DESeq(dds)
  resultsNames(dds)
  res_all_vs_ref<- results(dds)
  assign(paste0("res_all",all_set,"_vs_",ref),res_all_vs_ref,envir = .GlobalEnv)
  DESeq2::summary(res_all_vs_ref)
  assign("vst_all", as.data.frame(assay(varianceStabilizingTransformation(dds, blind=FALSE))),envir = .GlobalEnv)
  assign("dds_count_df" , as.data.frame(counts(dds, normalized=TRUE)),envir = .GlobalEnv)
  dds_count<- dds_count_df %>%
    rownames_to_column("gene")
  if (res_dir=="home_dir"){
    openxlsx::write.xlsx (dds_count, paste0(res_dir,"/RNAseq_normalized_",set_name,"_",set,".xlsx"),
                          rowNames = TRUE, colNames = TRUE, overwrite = T) 
    
    #for ImmQuant
    write.table(dds_count, paste0(res_dir,"/RNAseq_normalized_",set_name,"_",set,".txt"),
                row.names = F, col.names = TRUE,sep = "\t", quote = F)
  } else {
    openxlsx::write.xlsx (dds_count, paste0(res_dir,"/normalized_count_",set_name,"_",set,".xlsx"),
                          rowNames = TRUE, colNames = TRUE, overwrite = T) 
  }
  assign("cond_df", subset(md,select=eval(ref_col)),envir = .GlobalEnv) 
  gap_cols <- match(unique(cond_df[,ref_col]), cond_df[,ref_col])-1
  assign("gap_cols" , gap_cols[2:length(gap_cols)],envir = .GlobalEnv) 
  return(paste0("res_all",all_set,"_vs_",ref))
  
}


RNA_sig_pairs <- function(RNA_thresh,RNA_all_results,use_pval=T,element="gene",labeled=T,font_size=8,
                          res_dir=DE_result_dir,meta_dir=home_dir,mean_name_part=2, organism_name="mouse",
                          out_format="jpeg",col_order,padj_thresh=0.05,mean_anno_col,run,corr=F){
    
  wb_RNA <- openxlsx::createWorkbook()
  if (missing(run)) run<-set else run<-paste0(set,"_",run)
  first_res<-TRUE
  all_sig_genes<-character()
  all_sig_genes_df<-character()
  for (res in RNA_all_results){
    #res <- RNA_all_results[2]
    print(res)
    res_df<- as.data.frame(get(res))
#browser()    
    ## Volcano plot t
    pv <- plot_volcano(res_df,paste0(if (use_pval) "PVAL_" else "PADJ_",set,"_", res),output_format=out_format,
                       lfc = RNA_thresh, labeled=labeled, padj_thresh=padj_thresh,res_dir = res_dir)
    
    res_padj_sig <- subset(res_df, padj <= padj_thresh)
    print(paste(res, "padj sig:",nrow(res_padj_sig)))
    
    res_sig <- subset(res_df, pvalue <= 0.05)
    print(paste(res, "pvalue sig:",nrow(res_sig)))
    
    gene_res_padj_sig<-subset(res_padj_sig, abs(log2FoldChange) >=RNA_thresh)
    assign(paste0(res,"_padj_sig.",RNA_thresh), gene_res_padj_sig, envir = .GlobalEnv)
    print(paste("res_padj_sig_lfc.",RNA_thresh,nrow(gene_res_padj_sig)))
    
    # significats threshold 
    gene_res_pval_sig <- subset(res_sig, abs(log2FoldChange) >=RNA_thresh)
    assign(paste0(res,"_sig.",RNA_thresh), gene_res_pval_sig, envir = .GlobalEnv)
    print(paste("res_pvalue_sig_lfc.",RNA_thresh,nrow(gene_res_pval_sig)))
    
    if (use_pval) res_lfc<-gene_res_pval_sig else res_lfc<-gene_res_padj_sig
    #aggregate all sig genes from all the pairs
    # if(!str_detect(res,"all") & str_detect(res,"CTRL")) all_pairs_genes<- unique(c(all_pairs_genes,rownames(res_lfc)))
    all_sig_genes<- unique(c(all_sig_genes,rownames(res_lfc)))
    all_sig_genes_df<- rbind(all_sig_genes_df,cbind(res_lfc,Comparison=res))
    
    # Up padj significats threshold 
    gene_res_padj_sig_up <- subset(res_padj_sig, log2FoldChange >=RNA_thresh)
    assign(paste0(res,"_padj_up.",RNA_thresh), gene_res_padj_sig_up, envir = .GlobalEnv)
    print(paste("res_padj_up.",RNA_thresh,nrow(gene_res_padj_sig_up)))
    
    # Down padj significats threshold 
    gene_res_padj_sig_down <- subset(res_padj_sig, log2FoldChange <=-RNA_thresh)
    assign(paste0(res,"_padj_down.",RNA_thresh), gene_res_padj_sig_down, envir = .GlobalEnv)
    print(paste("res_padj_down.",RNA_thresh,nrow(gene_res_padj_sig_down)))
    
    # Up pval significats threshold 
    gene_res_pval_sig_up <- subset(res_sig, log2FoldChange >=RNA_thresh)
    assign(paste0(res,"_up.",RNA_thresh), gene_res_pval_sig_up, envir = .GlobalEnv)
    print(paste("res_pvalue_up.",RNA_thresh,nrow(gene_res_pval_sig_up)))
    
    # Down pval significats threshold 
    gene_res_pval_sig_down <- subset(res_sig, log2FoldChange <=-RNA_thresh)
    assign(paste0(res,"_down.",RNA_thresh), gene_res_pval_sig_down, envir = .GlobalEnv)
    print(paste("res_pvalue_down.",RNA_thresh,nrow(gene_res_pval_sig_down)))
    #save sig.1 to file for correlations with 16S
    #write.table(rownames(get(paste0(res,"_sig.",RNA_thresh))), 
    #            file = paste0(DE_result_dir,"/",set,"_",res,"_sig.",RNA_thresh,".txt"),  
    #            sep = "\t", row.names = F, col.names = FALSE, quote = FALSE)
    
#browser()    
    
    #------------------pairs up & down genes  
    ana_type<-ncol(str_split(res,"_",simplify = T))
    if ((ana_type==4 | ana_type==6) & !str_detect(res,"_y_")){
      st<-str_split_fixed(res,"\\_",4)[,2]
      nd<-str_split_fixed(res,"\\_",4)[,4]
    }else{
      st<-str_split_fixed(res,"\\_",5)[,3]
      nd<-str_split_fixed(res,"\\_",5)[,5]
    }
    #rld_pair_df<- get(paste0("rld_",nd))
    
    dds_count_sig<- dds_count_df %>%
        rownames_to_column("gene") %>%
        dplyr::filter(gene %in% rownames(res_lfc)) %>%
        column_to_rownames("gene")
#browser()    
    # keep the normalyzed counts of significant genes
    if (!str_detect(res,"all")){
      vst_pair_df<-get (str_replace(res,element,"vst"))
      anno_df<-get (str_replace(res,element,"anno"))
      
      #vst_pair_df<-vst_pair_df[, str_starts(colnames(vst_pair_df),st)|
      #                           str_starts(colnames(vst_pair_df),nd)]
      #dds_count_sig<-dds_count_sig[, str_starts(colnames(dds_count_sig),st)|
      #                               str_starts(colnames(dds_count_sig),nd)]
      # only the relevant rows
      #anno_df<-cond_df[str_starts(rownames(cond_df),st)|
      #                   str_starts(rownames(cond_df),nd),,drop = FALSE]
    }else{
      vst_pair_df<-vst_all
      anno_df<-cond_df
    }
    assign(paste0("dds_count_sig_",st,"_vs_",nd), dds_count_sig, envir = .GlobalEnv)
    
    # only the relevant cols
    print(paste("dds_count_sig:",nrow(dds_count_sig),ncol(dds_count_sig)))
#browser()    
    res_lfc <- res_lfc %>%
                  rownames_to_column("gene") %>%
                  arrange(lfcSE) %>%
                  column_to_rownames("gene")
    gap_cols <- match(unique(anno_df[,1]), anno_df[,1])-1
    gap_cols <-gap_cols[2:length(gap_cols)]
    if (corr) save(dds_count_sig, gene_res_pval_sig,res_lfc,
         file=paste0(meta_dir,"/",run,"_",res,if (use_pval)"_PVAL" else "_PADJ",
                     "_sig.",RNA_thresh,".Rdata"))
    
    #res_sig<-get(paste0(res,"_sig_",thresh))
    sig_rld<-vst_pair_df %>%
          rownames_to_column("gene") %>%
          dplyr::filter(gene %in% rownames(res_lfc)) %>%
          column_to_rownames("gene")
    
    if (use_pval) {
      up <- rownames(gene_res_pval_sig_up)
      down <- rownames(gene_res_pval_sig_down)
    }else{
      up <- rownames(gene_res_padj_sig_up)
      down <- rownames(gene_res_padj_sig_down)
    }
    assign(paste0(res,"_sig_vst.",RNA_thresh), sig_rld, envir = .GlobalEnv)
    print(paste("sig_vst:",nrow(sig_rld)))
    #save(sig_rld,anno_df,gap_cols,vst_pair_df,
    #     file=paste0(meta_dir,"/vst_",run,"_",res,"_sig.",RNA_thresh,".Rdata"))
    if (nrow(sig_rld)>0){
      # up genes 
      print(paste("up",length(up)))
#browser()      
      if (length(up)>1)      up_rld<- up_or_down_genes(res ,if (use_pval) "up_pval" else "up_padj", 
                                                       organism_name=organism_name, 
                                                       up,RNA_thresh, vst_pair_df, res_dir = res_dir, meta_dir=meta_dir,
                                                     anno_df, wb=wb_RNA, res_df=res_df,
                                                     fontsize=font_size,out_format=out_format)
      # down genes 
      print(paste("down",length(down)))
      if (length(down)>1)      down_rld<- up_or_down_genes(res ,if (use_pval) "down_pval" else "down_padj", 
                                                           organism_name=organism_name, 
                                                           down,RNA_thresh, vst_pair_df, res_dir = res_dir,meta_dir=meta_dir,
                                                         anno_df, wb=wb_RNA, res_df=res_df,
                                                         fontsize=font_size,out_format=out_format)
      if (length(up)>1 & length(down)>1){
        all_sig <- rbind(down_rld,up_rld)
        nrow(all_sig)
        nrow(up_rld)
        nrow(down_rld)
        reg_df<- data.frame("up_down" =c(rep("up",nrow(up_rld)),rep("down",nrow(down_rld)))) %>%
          arrange(up_down)
        rownames(reg_df)<-row.names(all_sig)
#browser()      
        plot_genes_heatmap(all_sig,anno_df,
                           out_format = out_format,
                           font_size=font_size,
                           paste0(set,"_",res,"_up_down_sig_", 
                                  if(use_pval) "pval" else "padj",".",RNA_thresh),reg_df,res_dir = res_dir)
        
      
        if(str_detect(res,"all")){
          print("mean heatmap")
#browser()      
          all_sig_mean_df <- all_sig %>%
            rownames_to_column("gene")  %>%
            #dplyr::filter(!str_detect(gene,"^Gm" ) &  !str_detect(gene,"Rik$" ))%>%
            pivot_longer(cols=-gene) %>%
            separate(name, c("a", "b"),  "_", extra = "drop") %>%
            dplyr::select(c(gene,id=ifelse(mean_name_part==1,"a","b"),value)) %>%
            #group_by(id,gene) %>%
            dplyr::group_by(gene,id) %>%
            #mutate(mean=mean(value,na.rm=TRUE))
            dplyr::summarize(Mean = mean(value, na.rm=TRUE)) %>%
            #dplyr::select(-value) %>%
            #distinct()  %>%
            pivot_wider(names_from = id, values_from=Mean)   %>%
            mutate(gene=substr(gene,1,60)) %>%
            ungroup() %>%
            arrange(rownames(reg_df)) %>%
            column_to_rownames("gene")
          all_sig_mean_df <- all_sig_mean_df[rownames(reg_df), ]             # Order data based on one column
          
          if (!missing(col_order)) all_sig_mean_df <-all_sig_mean_df %>% dplyr::relocate(col_order)
            
            gap_rows<-head(as.numeric(cumsum(table(reg_df$up_down))), -1)
          if (out_format=="pdf"){
  #browser()      
            #pdf(file=paste0(DE_result_dir,"/heatmap_",set,"_",res,"_up_down_sig_",
            #                                           if(use_pval) "pval" else "padj",".",RNA_thresh,"_mean","_genes",".pdf"))#,width=7, height=7)  
            save_pheatmap_pdf <- function(x, filename, width=7, height=7) {
              stopifnot(!missing(x))
              stopifnot(!missing(filename))
              pdf(filename, width=width, height=height)
              grid::grid.newpage()
              grid::grid.draw(x$gtable)
              dev.off()
            }
            p_title<-paste(set,res,"_up_down_sig",
                                  if(use_pval) "pval" else "padj",".",RNA_thresh,"mean","genes(",nrow(all_sig_mean_df),")")
            p1<-pheatmap(all_sig_mean_df, scale = "row", #col=my_colors,annotation_row=reg_df,
                     show_colnames= T,
                     fontsize=12,fontsize_row = 4, fontsize_number = 14,fontsize_col = 12,
                     cluster_cols = F, cluster_rows = T,
                     main = p_title)
            save_pheatmap_pdf(p1, paste0(res_dir,"/heatmap_",set,"_",res,"_up_down_sig_",
                                         if(use_pval) "pval" else "padj",".",RNA_thresh,"_mean","_genes",".pdf"))
          } else {
            if (!missing(mean_anno_col)){
#browser()
              plot_genes_heatmap(rld=all_sig_mean_df,
                                 anno_df=mean_anno_col, #gap_c=gap_cols,
                                 #cols_dend =FALSE,
                                 #scl="none",
                                 anno_row=reg_df,
                                 gap_r = gap_rows,
                                 set_colnames=T,
                                 out_format = out_format,
                                 res_dir = res_dir,
                                 set_name=paste0(set,"_",res,"_up_down_sig_",
                                                 if(use_pval) "pval" else "padj",".",RNA_thresh,"_mean"))
            }else{
              plot_genes_heatmap(rld=all_sig_mean_df,
                               anno_df=NA, #gap_c=gap_cols,
                               cols_dend =FALSE,
                               #scl="none",
                               anno_row=reg_df,
                               gap_r = gap_rows,
                               set_colnames=T,
                               out_format = out_format,
                               res_dir = res_dir,
                               set_name=paste0(set,"_",res,"_up_down_sig_",
                                               if(use_pval) "pval" else "padj",".",RNA_thresh,"_mean"))
            }#mean_anno_col
          }
        }
      }
      }
    }
  #}
#browser()
  assign(paste0("all_sig_genes.",RNA_thresh,"_",run), all_sig_genes, envir = .GlobalEnv)
  assign(paste0("all_sig_genes_df.",RNA_thresh,"_",run), all_sig_genes_df, envir = .GlobalEnv)
  openxlsx::saveWorkbook(wb_RNA, 
                         file=paste0(meta_dir,"/",run,if (use_pval)"_PVAL" else "_PADJ",
                                     "_sig_gene_annotation_",RNA_thresh,".xlsx"), overwrite = TRUE)
}
  
  
  
compare_sig_genes<-function(set_name=NULL,list1,label_list1, res_dir=DE_result_dir,thresh_in=thresh_pair,
                            element="genes",fig_LFC_thresh, use_pval=use_pval, pthresh=0.05){
  pdf(file=paste0(res_dir,"/venn_compare_sig_",element,"_",set_name, ".pdf"),width=7, height=7) 
  wb_comp <- openxlsx::createWorkbook()
  list2<-list1
  for (res1 in list1){
    list2<-list2[list2 != res1]
    for (res2 in list2){
      if (res1!=res2){
        print(paste(res1,res2))
#browser()        
        par(mfrow=c(2,3))
        if (element=="genes" | element=="proteo"){
          res1_sig_df<-as.data.frame(get(paste0(res1,"_sig.",thresh_in)))
          res2_sig_df<-as.data.frame(get(paste0(res2,"_sig.",thresh_in)))

          venn(c( setNames(list(rownames(res1_sig_df)),
                           paste0(res1,"_sig.",thresh_in)), 
                  setNames(list(rownames(res2_sig_df)),
                           paste0(res2,"_sig.",thresh_in)) ))
          
          venn(c( setNames(list(rownames(get(paste0(res1,"_up.",thresh_in)))),
                           paste0(res1,"_up.",thresh_in)), 
                  setNames(list(rownames(get(paste0(res2,"_up.",thresh_in)))),
                           paste0(res2,"_up.",thresh_in)) ))
          venn(c( setNames(list(rownames(get(paste0(res1,"_up.",thresh_in)))),
                           paste0(res1,"_up.",thresh_in)), 
                  setNames(list(rownames(get(paste0(res2,"_down.",thresh_in)))),
                           paste0(res2,"_down.",thresh_in)) ))
          venn(c( setNames(list(rownames(get(paste0(res1,"_down.",thresh_in)))),
                           paste0(res1,"_down.",thresh_in)), 
                  setNames(list(rownames(get(paste0(res2,"_up.",thresh_in)))),
                           paste0(res2,"_up.",thresh_in)) ))
          venn(c( setNames(list(rownames(get(paste0(res1,"_down.",thresh_in)))),
                           paste0(res1,"_down.",thresh_in)), 
                  setNames(list(rownames(get(paste0(res2,"_down.",thresh_in)))),
                           paste0(res2,"_down.",thresh_in)) ))
        } else if (element=="metabo" | element=="taxa"){
          res1_sig_df<-as.data.frame(get(paste0(res1,if(use_padj) "_padj_sig" else "_pval_sig")))
          res2_sig_df<-as.data.frame(get(paste0(res2,if(use_padj) "_padj_sig" else "_pval_sig")))
        
          venn(c( setNames(list(rownames(res1_sig_df)),
                           paste0(res1,"_sig.",thresh_in)), 
                  setNames(list(rownames(res2_sig_df)),
                           paste0(res2,"_sig.",thresh_in)) ))
          
          venn(c( setNames(list(rownames(subset(res1_sig_df,log2FoldChange>0))),
                           paste0(res1,"_up.",thresh_in)), 
                  setNames(list(rownames(subset(res2_sig_df,log2FoldChange>0))),
                           paste0(res2,"_up.",thresh_in)) ))
          venn(c( setNames(list(rownames(subset(res1_sig_df,log2FoldChange>0))),
                           paste0(res1,"_up.",thresh)), 
                  setNames(list(rownames(subset(res2_sig_df,log2FoldChange<0))),
                           paste0(res2,"_down.",thresh_in)) ))
          venn(c( setNames(list(rownames(subset(res1_sig_df,log2FoldChange<0))),
                           paste0(res1,"_down.",thresh_in)), 
                  setNames(list(rownames(subset(res2_sig_df,log2FoldChange>0))),
                           paste0(res2,"_up.",thresh_in)) ))
          venn(c( setNames(list(rownames(subset(res1_sig_df,log2FoldChange<0))),
                           paste0(res1,"_down.",thresh_in)), 
                  setNames(list(rownames(subset(res2_sig_df,log2FoldChange<0))),
                           paste0(res2,"_down.",thresh_in)) ))
        }
#browser()        
        res1_df<-as.data.frame(get(res1)) #%>%
                    # remove_rownames() %>%
                    # column_to_rownames("gene")
        res2_df<-as.data.frame(get(res2)) #%>%
                    # remove_rownames() %>%
                    # column_to_rownames("gene")
        if (!missing(fig_LFC_thresh)){
          res1_df<-res1_df %>%
                      dplyr::filter(abs(log2FoldChange)<fig_LFC_thresh)
          res2_df<-res2_df %>%
                      dplyr::filter(abs(log2FoldChange)<fig_LFC_thresh)
        }
        res1_df_lfc<- subset(res1_df,select="log2FoldChange")
        res2_df_lfc<- subset(res2_df,select="log2FoldChange") 
        res_df_both <- merge(res1_df_lfc, res2_df_lfc, by="row.names") %>%
          column_to_rownames("Row.names") %>%
          setnames(c(paste0("lfc_",res1),paste0("lfc_",res2)))
        if(missing(label_list1))  label_list1<-intersect(rownames(res1_sig_df),rownames(res2_sig_df))
        label_list2<-c("null")
#browser()        
        
        lfc_merged<- rownames_to_column(res1_df,"gene") %>%
          full_join(rownames_to_column(res2_df,"gene"),by="gene") %>%
          dplyr::rename(log2FoldChange1=log2FoldChange.x, 
                        log2FoldChange2=log2FoldChange.y,
                        pval1=ifelse(use_pval,"pvalue.x","padj.x"),
                        pval2=ifelse(use_pval,"pvalue.y","padj.y"))  %>%
          mutate_at(vars(!contains("gene")),funs(replace(., is.na(.), 0)))  %>%
          # add sig, "darkorange2" both, green-H, magenta-Drug, grey-none
          mutate(significant=ifelse(abs(log2FoldChange1) >=thresh_in & abs(log2FoldChange2) >= thresh_in &
                                      pval2<pthresh & pval1<pthresh, "both", 
                                    ifelse(abs(log2FoldChange2) >=thresh_in & pval2<pthresh,eval(res2), 
                                           ifelse(abs(log2FoldChange1) >=thresh_in & pval1<pthresh,eval(res1), "none"))))  %>%
          # add label to both sig
          mutate(genelabels = ifelse(gene %in% label_list1, gene,
                                     ifelse(gene %in% label_list2,
                                            gene,NA))) %>%
          column_to_rownames("gene")
#browser()        
        par(mfrow=c(1,1))
        print(ggscatter(lfc_merged,x="log2FoldChange1", y="log2FoldChange2", 
                        #ylim = c(-5,5),
                        main = paste("Scatter plot LFC of",res1,"and",res2,
                                     ifelse(missing(label_list1),"\nBoth sig labels","")),
                        shape  = 20, 
                        size  = ifelse(lfc_merged$significant=="none",0.2,2.5), 
                        frame = FALSE,
                        #label = "genelabels", repel = TRUE, font.label=c(7, "bold", "black"),  label.rectangle = T,
                        color = "significant", palette = c("darkorange2", "green", "magenta", "grey"))+
                xlab(str_replace(res1,"gene_y_","LFC "))+
                ylab(str_replace(res2,"gene_y_","LFC "))+
                geom_label_repel(aes(label=genelabels),color="black", max.overlaps=Inf, size =2, 
                                 box.padding = unit(0.2, "lines"))+
                geom_hline(yintercept=0) +
                geom_vline(xintercept=0) )#+ 
        
        #plot(res_df_both[,1], res_df_both[,2], 
        #     main = paste("Scatter plot LFC of ",res1,"and LFC of",res2),
        #     xlab = paste("LFC",res1), ylab = paste("LFC",res2),
        #     pch = 20, cex = .1, frame = FALSE)
        #lines(x = c(min(res_df_both[,1]),max(res_df_both[,1])), 
        #      y = c(min(res_df_both[,2]),max(res_df_both[,2])))
        
        
        
#browser()
        sheet_name<-str_replace_all(paste0(res1,"+",res2),"gene_","")
        sheet_name<-str_replace_all(sheet_name,"metabo_","")
        sheet_name<-str_replace_all(sheet_name,"y_","")
        if (str_length(sheet_name)>31) sheet_name<-substr(sheet_name,1,31)
        addWorksheet(wb_comp,sheet_name )
        comm_list<-intersect(rownames(res1_sig_df),
                             rownames(res2_sig_df))
        if (element=="genes"){
          anno_gene_list <- anno_genes(comm_list) %>%
            dplyr::rename("gene"="external_gene_name")%>%
            left_join(rownames_to_column(res_df_both,"gene"),by="gene")
          writeData(wb_comp,sheet_name , anno_gene_list, colNames = TRUE, startRow = 1) 
        } else{
#browser()
          comm_df<-data.frame(gene=comm_list)%>%
                    left_join(rownames_to_column(res_df_both,"gene"),by="gene")
          writeData(wb_comp,sheet_name , comm_df, colNames = TRUE, startRow = 1) 
          posStyle <- createStyle(fontColour = "#006100", bgFill = "#C6EFCE")
          negStyle <- createStyle(fontColour = "#9C0006", bgFill = "#FFC7CE")
          conditionalFormatting(wb_comp, sheet_name,
                                cols = 2:3,
                                rows = 2:nrow(comm_df), rule = "AND($B2<0,$C2>0)", style = posStyle)
          conditionalFormatting(wb_comp, sheet_name,
                                cols = 2:3,
                                rows = 2:nrow(comm_df), rule = "AND($B2>0,$C2<0)", style = negStyle)
        }
        setColWidths(wb_comp, sheet_name, cols = 1:10, widths = "auto")
        
        sheet_name<-str_replace_all(paste0(res1,"-",res2),"gene_","")
        sheet_name<-str_replace_all(sheet_name,"metabo_","")
        sheet_name<-str_replace_all(sheet_name,"y_","")
        if (str_length(sheet_name)>31) sheet_name<-substr(sheet_name,1,31)
        addWorksheet(wb_comp,sheet_name )
        only_res1<-setdiff(rownames(res1_sig_df),
                             rownames(res2_sig_df))
        if (element=="genes"){
          anno_gene_list <- anno_genes(only_res1) %>%
                        dplyr::rename("gene"="external_gene_name")%>%
                        left_join(rownames_to_column(res1_df_lfc,"gene"),by="gene")
          writeData(wb_comp,sheet_name , anno_gene_list, colNames = TRUE, startRow = 1) 
        } else{
          #browser()
          only_res1_df<-data.frame(gene=only_res1)%>%
                    left_join(rownames_to_column(res1_df_lfc,"gene"),by="gene")
          writeData(wb_comp,sheet_name , only_res1_df, colNames = TRUE, startRow = 1) 
        }
        setColWidths(wb_comp, sheet_name, cols = 1:10, widths = "auto")
#browser()        
        sheet_name<-str_replace_all(paste0(res2,"-",res1),"gene_","")
        sheet_name<-str_replace_all(sheet_name,"metabo_","")
        sheet_name<-str_replace_all(sheet_name,"y_","")
        if (str_length(sheet_name)>31) sheet_name<-substr(sheet_name,1,31)
        addWorksheet(wb_comp,sheet_name )
        only_res2<-setdiff(rownames(res2_sig_df),
                             rownames(res1_sig_df))
        if (element=="genes"){
          anno_gene_list <- anno_genes(only_res2) %>%
                          dplyr::rename("gene"="external_gene_name")%>%
                          left_join(rownames_to_column(res2_df_lfc,"gene"),by="gene")
          writeData(wb_comp,sheet_name , anno_gene_list, colNames = TRUE, startRow = 1) 
        } else{
#browser()
          only_res2_df<-data.frame(gene=only_res2)%>%
                        left_join(rownames_to_column(res2_df_lfc,"gene"),by="gene")
          writeData(wb_comp,sheet_name , only_res2_df, colNames = TRUE, startRow = 1) 
        }
        setColWidths(wb_comp, sheet_name, cols = 1:10, widths = "auto")
        
      }
    }
  }
  dev.off()
  openxlsx::saveWorkbook(wb_comp, 
                         file=paste0(res_dir,"/",set_name,"_common_sig.",thresh_in,"_",element,".xlsx"),                        overwrite = TRUE)
}
  
compare_all_genes<-function(set_name=NULL,list1,label_list1, res_dir=DE_result_dir,thresh_in=thresh_pair,
                            element="genes",fig_LFC_thresh){
  pdf(file=paste0(res_dir,"/venn_compare_all_",element,"_",set_name, ".pdf"),width=7, height=7) 
  wb_comp <- openxlsx::createWorkbook()
  list2<-list1
  for (res1 in list1){
    list2<-list2[list2 != res1]
    for (res2 in list2){
      if (res1!=res2){
        print(paste(res1,res2))
#browser()        
        par(mfrow=c(2,3))
        res1_df<-subset(as.data.frame(get(res1)),pvalue<=0.05)
        res2_df<-subset(as.data.frame(get(res2)),pvalue<=0.05)
        res1_df_up<-subset(res1_df,log2FoldChange>0)
        res1_df_down<-subset(res1_df,log2FoldChange<0)
        res2_df_up<-subset(res2_df,log2FoldChange>0)
        res2_df_down<-subset(res2_df,log2FoldChange<0)

        venn(c( setNames(list(rownames(res1_df)),res1), 
                setNames(list(rownames(res2_df)),res2) ))
        
        venn(c( setNames(list(rownames(res1_df_up)),
                         paste0(res1,"_up")), 
                setNames(list(rownames(res2_df_up)),
                         paste0(res2,"_up")) ))
        venn(c( setNames(list(rownames(res1_df_up)),
                         paste0(res1,"_up")), 
                setNames(list(rownames(res2_df_down)),
                         paste0(res2,"_down")) ))
        venn(c( setNames(list(rownames(res1_df_down)),
                         paste0(res1,"_down")), 
                setNames(list(rownames(res2_df_up)),
                         paste0(res2,"_up")) ))
        venn(c( setNames(list(rownames(res1_df_down)),
                         paste0(res1,"_down")), 
                setNames(list(rownames(res2_df_down)),
                         paste0(res2,"_down")) ))
        #browser()        
        if (!missing(fig_LFC_thresh)){
          res1_df<-res1_df %>%
            dplyr::filter(abs(log2FoldChange)<fig_LFC_thresh)
          res2_df<-res2_df %>%
            dplyr::filter(abs(log2FoldChange)<fig_LFC_thresh)
        }
        res1_df_lfc<- subset(res1_df,select="log2FoldChange")
        res2_df_lfc<- subset(res2_df,select="log2FoldChange") 
        res_df_both <- merge(res1_df_lfc, res2_df_lfc, by="row.names") %>%
          column_to_rownames("Row.names") %>%
          setnames(c(paste0("lfc_",res1),paste0("lfc_",res2)))
        if(missing(label_list1))  label_list1<-intersect(rownames(res1_df),rownames(res2_df))
        label_list2<-c("null")
        #browser()        
        
        lfc_merged<- rownames_to_column(res1_df,"gene") %>%
          full_join(rownames_to_column(res2_df,"gene"),by="gene") %>%
          dplyr::rename(log2FoldChange1=log2FoldChange.x, 
                        log2FoldChange2=log2FoldChange.y,
                        pval1=pvalue.x,
                        pval2=pvalue.y)  %>%
          mutate_at(vars(!contains("gene")),funs(replace(., is.na(.), 0)))  %>%
          mutate(significant=ifelse(abs(log2FoldChange1) >=thresh_in & abs(log2FoldChange2) >= thresh_in &
                                      pval2<0.05 & pval1<0.05, "both", 
                                    ifelse(abs(log2FoldChange2) >=thresh_in & pval2<0.05,eval(res2), 
                                           ifelse(abs(log2FoldChange1) >=thresh_in & pval1<0.05,eval(res1), "none"))))  %>%
          # add label to both sig
          mutate(genelabels = ifelse(gene %in% label_list1, gene,
                                     ifelse(gene %in% label_list2,
                                            gene,NA))) %>%
          column_to_rownames("gene")
        #browser()        
        par(mfrow=c(1,1))
        print(ggscatter(lfc_merged,x="log2FoldChange1", y="log2FoldChange2", 
                        #ylim = c(-5,5),
                        main = paste("Scatter plot LFC of",res1,"and",res2,
                                     ifelse(missing(label_list1),"\nBoth labels","")),
                        shape  = 20, 
                        size  = ifelse(lfc_merged$significant=="none",0.2,2.5), 
                        frame = FALSE,
                        #label = "genelabels", repel = TRUE, font.label=c(7, "bold", "black"),  label.rectangle = T,
                        color = "significant", palette = c("darkorange2", "green", "magenta", "grey"))+
                xlab(str_replace(res1,"gene_y_","LFC "))+
                ylab(str_replace(res2,"gene_y_","LFC "))+
                geom_label_repel(aes(label=genelabels),color="black", max.overlaps=Inf, size =2, 
                                 box.padding = unit(0.2, "lines"))+
                geom_hline(yintercept=0) +
                geom_vline(xintercept=0) )#+ 
        
        #plot(res_df_both[,1], res_df_both[,2], 
        #     main = paste("Scatter plot LFC of ",res1,"and LFC of",res2),
        #     xlab = paste("LFC",res1), ylab = paste("LFC",res2),
        #     pch = 20, cex = .1, frame = FALSE)
        #lines(x = c(min(res_df_both[,1]),max(res_df_both[,1])), 
        #      y = c(min(res_df_both[,2]),max(res_df_both[,2])))
        
        
        
        #browser()
        sheet_name<-str_replace_all(paste0(res1,"+",res2),"gene_","")
        sheet_name<-str_replace_all(sheet_name,"metabo_","")
        sheet_name<-str_replace_all(sheet_name,"y_","")
        if (str_length(sheet_name)>31) sheet_name<-substr(sheet_name,1,31)
        addWorksheet(wb_comp,sheet_name )
        comm_list<-intersect(rownames(res1_df),
                             rownames(res2_df))
        if (element=="genes"){
          anno_gene_list <- anno_genes(comm_list) %>%
            dplyr::rename("gene"="external_gene_name")%>%
            left_join(rownames_to_column(res_df_both,"gene"),by="gene")
          writeData(wb_comp,sheet_name , anno_gene_list, colNames = TRUE, startRow = 1) 
        } else{
          #browser()
          comm_df<-data.frame(gene=comm_list)%>%
            left_join(rownames_to_column(res_df_both,"gene"),by="gene")
          writeData(wb_comp,sheet_name , comm_df, colNames = TRUE, startRow = 1) 
          posStyle <- createStyle(fontColour = "#006100", bgFill = "#C6EFCE")
          negStyle <- createStyle(fontColour = "#9C0006", bgFill = "#FFC7CE")
          conditionalFormatting(wb_comp, sheet_name,
                                cols = 2:3,
                                rows = 2:nrow(comm_df), rule = "AND($B2<0,$C2>0)", style = posStyle)
          conditionalFormatting(wb_comp, sheet_name,
                                cols = 2:3,
                                rows = 2:nrow(comm_df), rule = "AND($B2>0,$C2<0)", style = negStyle)
        }
        setColWidths(wb_comp, sheet_name, cols = 1:10, widths = "auto")
        
        sheet_name<-str_replace_all(paste0(res1,"-",res2),"gene_","")
        sheet_name<-str_replace_all(sheet_name,"metabo_","")
        sheet_name<-str_replace_all(sheet_name,"y_","")
        if (str_length(sheet_name)>31) sheet_name<-substr(sheet_name,1,31)
        addWorksheet(wb_comp,sheet_name )
        only_res1<-setdiff(rownames(res1_df),
                           rownames(res2_df))
        if (element=="genes"){
          anno_gene_list <- anno_genes(only_res1) %>%
            dplyr::rename("gene"="external_gene_name")%>%
            left_join(rownames_to_column(res1_df_lfc,"gene"),by="gene")
          writeData(wb_comp,sheet_name , anno_gene_list, colNames = TRUE, startRow = 1) 
        } else{
          #browser()
          only_res1_df<-data.frame(gene=only_res1)%>%
            left_join(rownames_to_column(res1_df_lfc,"gene"),by="gene")
          writeData(wb_comp,sheet_name , only_res1_df, colNames = TRUE, startRow = 1) 
        }
        setColWidths(wb_comp, sheet_name, cols = 1:10, widths = "auto")
        #browser()        
        sheet_name<-str_replace_all(paste0(res2,"-",res1),"gene_","")
        sheet_name<-str_replace_all(sheet_name,"metabo_","")
        sheet_name<-str_replace_all(sheet_name,"y_","")
        if (str_length(sheet_name)>31) sheet_name<-substr(sheet_name,1,31)
        addWorksheet(wb_comp,sheet_name )
        only_res2<-setdiff(rownames(res2_df),
                           rownames(res1_df))
        if (element=="genes"){
          anno_gene_list <- anno_genes(only_res2) %>%
            dplyr::rename("gene"="external_gene_name")%>%
            left_join(rownames_to_column(res2_df_lfc,"gene"),by="gene")
          writeData(wb_comp,sheet_name , anno_gene_list, colNames = TRUE, startRow = 1) 
        } else{
          #browser()
          only_res2_df<-data.frame(gene=only_res2)%>%
            left_join(rownames_to_column(res2_df_lfc,"gene"),by="gene")
          writeData(wb_comp,sheet_name , only_res2_df, colNames = TRUE, startRow = 1) 
        }
        setColWidths(wb_comp, sheet_name, cols = 1:10, widths = "auto")
        
      }
    }
  }
  dev.off()
  openxlsx::saveWorkbook(wb_comp, 
                         file=paste0(res_dir,"/",set_name,"_common_all_",element,".xlsx"),                        overwrite = TRUE)
}


LFC2LFC<-function(comparisons,res_file_in=res_file,res_sig,set_name=NULL,res_dir=DE_result_dir,thresh_in=thresh_pair,
                  element="genes",use_pval_in=use_pval, out_format="PDF", venn_no=1, quantile_val=0){
  
  adjust<- ifelse(use_pval_in,"pvalue","padj")
  if (out_format=="PDF") pdf(file=paste0(res_dir,"/",element,"_compare_lfc_",adjust,".",thresh_in,"_",set, ".pdf"),width=7, height=8) 
#browser()
  if (out_format=="HTML"){
    cat('\n\n### Compare significants counts\n\n')
  }  
  
#Make venn plot
  if (venn_no==1){
    print(ggVennDiagram(res_sig, label_alpha = 0,set_size=3)+
            theme(legend.position = "none")+
            ggtitle(paste("Compare",adjust,"significant ",element,"(thresh",thresh_in,")")) +
            scale_x_continuous(expand = expansion(mult = .2))+
            #scale_fill_gradientn(colours = terrain.colors(7))
            #paletteer::scale_fill_paletteer_c("viridis::plasma")
            scale_fill_distiller(palette = "RdBu"))
  }else if (venn_no==2){
    g1<-ggVennDiagram(res_sig[1:3], label_alpha = 0,set_size=3)+
            theme(legend.position = "none")+
            ggtitle(paste("Compare",adjust,"significant ",element,"(thresh",thresh_in,")")) +
            scale_x_continuous(expand = expansion(mult = .3))+
            #scale_fill_gradientn(colours = terrain.colors(7))
            #paletteer::scale_fill_paletteer_c("viridis::plasma")
            scale_fill_distiller(palette = "RdBu")
    if (out_format=="HTML"){
      fig_subchunkify(g1, 5, 5)
    } else {
      print(g1)
    }
    
    g1<-ggVennDiagram(res_sig[4:6], label_alpha = 0,set_size=3)+
            theme(legend.position = "none")+
            ggtitle(paste("Compare",adjust,"significant ",element,"(thresh",thresh_in,")")) +
            scale_x_continuous(expand = expansion(mult = .3))+
            #scale_fill_gradientn(colours = terrain.colors(7))
            #paletteer::scale_fill_paletteer_c("viridis::plasma")
            scale_fill_distiller(palette = "RdBu")
    if (out_format=="HTML"){
      fig_subchunkify(g1, 5, 5)
    } else {
      print(g1)
    }
  }
  
  if (out_format=="HTML"){
    cat('\n\n### Compare LFC pairwise\n\n')
  }  

  res_df_both_all<- list()
  for (i in 1:nrow(comparisons)){
    #i<-1
    res1<-as.character(comparisons[i,1])
    res2<-as.character(comparisons[i,2])
    compare_name<-paste0(res1,"-",res2)
    if (out_format!="HTML") print(paste(res1,"   ",res2))
#browser()  
    res1_df<-subset(res_file_in,Comparison==res1,select=-Comparison) %>%
            dplyr::filter(log2FoldChange>quantile(log2FoldChange, quantile_val,na.rm=T) & 
                            log2FoldChange<quantile(log2FoldChange, (1-quantile_val),na.rm=T)) %>%
            remove_rownames() %>%
            column_to_rownames("Gene_name")

    res2_df<-subset(res_file_in,Comparison==res2,select=-Comparison) %>%
            dplyr::filter(log2FoldChange>quantile(log2FoldChange, quantile_val,na.rm=T) & 
                            log2FoldChange<quantile(log2FoldChange, (1-quantile_val),na.rm=T)) %>%
            remove_rownames() %>%
            column_to_rownames("Gene_name")
#browser()  
    res_df_both <- merge(subset(res1_df,select=c("log2FoldChange",adjust)),
                         subset(res2_df, select=c("log2FoldChange",adjust)),
                         by="row.names",all =T) %>%
      setnames(c("gene","lfc1","p1","lfc2","p2")) %>%
      mutate_at(vars(!contains("gene")),funs(replace(., is.na(.), 0)))  %>%
      # add sig, "darkorange2" both, green-Chow, magenta-HFD, grey-none
      mutate(Significant=ifelse(abs(lfc1) >=thresh_in & 
                                  abs(lfc2) >= thresh_in &
                                  p1<0.05 & p2<0.05, "both", 
                                ifelse(abs(lfc1) >=thresh_in & p1<0.05,res1, 
                                       ifelse(abs(lfc2) >=thresh_in & p2<0.05,res2, "none"))),
             dif=abs(lfc1-lfc2),
             # genelabels = ifelse(abs(dif)>=2 ,gene,NA)) %>%
             genelabels = ifelse((sign(lfc1)!= sign(lfc2) & Significant=="both") & #abs(dif)>2
                                   Significant!="none" ,gene,NA)) %>%  #  label to opposite
      column_to_rownames("gene")
    both_no<-nrow(subset(res_df_both,Significant=="both"))
    both_opp_no<-nrow(subset(res_df_both,!is.na(genelabels)))
#browser()
    my_pal<-c("darkorange2", "green", "magenta", "grey")
    names(my_pal)<- c(res1,res2,"both","none")
    print(ggscatter(res_df_both,x="lfc1", y="lfc2", 
                    # ylim = c(-5,5),
                    main = paste("Compare",element,"LFC",compare_name,"Scatter plot ",
                                 "\nSignificant both (",both_no,")",
                                 "\nLabels on significant both- opposite direction(",both_opp_no,")"),
          
                    shape  = 20, 
                    size  = ifelse(res_df_both$Significant=="none",0.2,2), 
                    frame = FALSE,
                    label = "genelabels", repel = TRUE, font.label=5,  label.rectangle = T,
                    color = "Significant", palette = my_pal)+
            labs(x=paste("LFC",res1),y=paste("LFC",res2)) +
            tune::coord_obs_pred()+
            geom_hline(yintercept=0) +
            geom_vline(xintercept=0)) #+ 
    #geom_text_repel()
    
    res_df_both_sig<-subset(res_df_both,Significant!="none") %>%
      arrange(Significant)
    # pheatmap(subset(res_df_both_sig,select=c("lfc1","lfc2")),
    #          annotation_row=subset(res_df_both_sig,select="Significant"), cluster_rows = F,
    #          cluster_cols = F,fontsize_row = 2,fontsize=8)
    
    # venn(c( setNames(list(rownames(subset(res_df_both_sig,Significant!=res2))),
    #                  paste0(res1," sig.",thresh_in)), 
    #         setNames(list(rownames(subset(res_df_both_sig,Significant!=res1))),
    #                  paste0(res2," sig.",thresh_in)) ))
    res_df_both_all[[i]]<-res_df_both %>%
      setnames(c(paste0("lfc_",res1),paste0("p_",res1),
                 paste0("lfc_",res2),paste0("p_",res2),
                 "Significant","dif","genelabels")) %>%
      arrange(Significant,desc(dif)) %>%
      rownames_to_column(compare_name)
    names(res_df_both_all)[i]<-str_replace_all(compare_name,"sirt6","TG")
    names(res_df_both_all)[i]<-str_sub(str_replace_all(str_replace_all(compare_name,"Low","L"),"High","H"),1,31)
  }
  res_df_both_all[[i+1]]<-Reduce(intersect, res_sig)
  names(res_df_both_all)[i+1]<-"sig_common2all"
  
  openxlsx::write.xlsx(res_df_both_all,
                       file=paste0(res_dir,"/",element,"_compare_lfc_",adjust,".",thresh_in,"_",set,".xlsx"),overwrite=T)
  if (out_format=="PDF") dev.off()

}


  prepare_GSEA_rnk <- function(res_df,rnk_file,human,mouse,LFC=F,res_dir=home_dir){
    #res_df<-res_treat_df
    if (!LFC){
      res_df$fcSign <- sign(res_df$log2FoldChange)
      res_df$logP=ifelse(is.infinite(-log10(res_df$pvalue)), 0, -log10(res_df$pvalue))
      res_df$metric=res_df$logP/res_df$fcSign
    }else {
      res_df$metric=res_df$log2FoldChange
    }
    res_df$mouse_gene<-rownames(res_df)
#browser()
    print(paste("nrow(res_df):",nrow(res_df)))
    #print(res_df)
    
     #convert genes from mouse to human for GSEA
    Mouse2HumanTable <- homologene::mouse2human(unlist(c(res_df$mouse_gene)))
    # Mouse2HumanTable <- Mouse2Human(MouseGenes = res_df$mouse_gene,human=human,mouse=mouse)
    print(paste("nrow(Mouse2HumanTable):",nrow(Mouse2HumanTable)))
#browser()
    res_df$human_gene<- 
      Mouse2HumanTable[match(res_df$mouse_gene, Mouse2HumanTable$mouseGene), "humanGene"] 
    print(paste("nrow(res_df):",nrow(res_df)))
    
    
    #human_GSEA<- subset(res_df,!(is.na(human_gene)| is.na(metric)), select= c("human_gene", "metric"))
    human_GSEA<- subset(res_df,!is.na(human_gene)& !is.na(metric)& str_length(human_gene)>0, 
                        select= c("human_gene", "metric"))
    #human_GSEA_LFC<- subset(res_df,!is.na(human_gene)& !is.na(log2FoldChange)& str_length(human_gene)>0, 
    #                    select= c("human_gene", "log2FoldChange"))
    print(paste("nrow(human_GSEA):",nrow(human_GSEA), 
                "unique nrow(human_GSEA):", length(unique(human_GSEA$human_gene))))
    #save rnk file for GSEA 
    write.table(human_GSEA, file = paste0(res_dir,rnk_file,".rnk"), sep = "\t", 
                row.names = F, col.names = FALSE, quote = FALSE)
    #write.table(human_GSEA_LFC, file = paste0(home_dir,rnk_file,"_LFC.rnk"), sep = "\t", 
    #            row.names = F, col.names = FALSE, quote = FALSE)
}
  
Mouse2Human <- function(MouseGenes,human_mart,mouse_mart){
  if (missing(human_mart)){
      human_mart = useMart("ensembl", dataset = "hsapiens_gene_ensembl", host =  "https://www.ensembl.org")
      mouse_mart = useMart("ensembl", dataset = "mmusculus_gene_ensembl", host =  "https://www.ensembl.org")
    }
    print(paste("in Mouse2Human MouseGenes:",length(MouseGenes)))
#browser()
    genesMousetoHuman = getLDS(attributes = c("external_gene_name","mgi_symbol"), 
                               filters = "external_gene_name", 
                               values = MouseGenes , 
                               mart = mouse_mart, 
                               attributesL = c("ensembl_gene_id", "hgnc_symbol"), 
                               martL = human_mart, 
                               uniqueRows = TRUE)
    
    colnames(genesMousetoHuman) <- c("Mouse.Gene_ID", "MGI", "Human.Gene_ID", "HGNC")
    print(paste("in Mouse2Human nrow(genesMousetoHuman):",nrow(genesMousetoHuman)))
    
    return(genesMousetoHuman) 
    
  }
  
Human2Mouse <- function(HumanGenes,in_mouse_ensembl,in_human_ensembl){
  if (missing(in_human_ensembl)){
    in_mouse_ensembl<- useEnsembl(biomart = "genes", 
                               dataset = "mmusculus_gene_ensembl")
    in_human_ensembl<- useEnsembl(biomart = "genes", 
                               dataset = "hsapiens_gene_ensembl")
  }
#browser()
  print(paste("in HumanGenes HumanGenes:",length(HumanGenes)))
  genesHuman2Mouse = getLDS(attributes = c("external_gene_name"), #
                             filters = "external_gene_name", 
                             values = HumanGenes, # pw_genes$SYMBOL, 
                             mart = in_human_ensembl, 
                             attributesL = c("external_gene_name"), #,"mgi_symbol"
                             martL = in_mouse_ensembl, 
                             uniqueRows = TRUE)
  
#browser()
  colnames(genesHuman2Mouse) <- c( "Mouse.Gene_ID","Human.Gene_ID" )
  print(paste("in Human2Mouse nrow(genesHuman2Mouse):",nrow(genesHuman2Mouse)))
  
  return(genesHuman2Mouse$Mouse.Gene_ID) 
  
}

GSEA_results_file <- function(file_type="tsv",set_name,sub_dir_prefix,fdr=0.05){
  file_names <-list.files(home_dir, 
                          pattern= glob2rx(paste0("*gsea_report_for_na*.",file_type,"$")),
                          recursive = TRUE)
  if (!missing(sub_dir_prefix))       file_names <-file_names[str_detect(file_names, sub_dir_prefix)]
  #browser()
  duplicated(file_names)
  file_names[duplicated(file_names)]
  file_names <-as.data.frame(file_names) #%>%
  #dplyr::filter(str_detect(.,set))
  
  wb <- createWorkbook()
  for (fn in file_names$file_names) {
    #fn<- "CS_vs_CTRL_C2.GseaPreranked.1592844211821/gsea_report_for_na_neg_1592844211821.xls"
    print(fn)
    underscore_loc<-as.numeric(as.data.frame(str_locate(fn, ".GseaPreranked"))[1])
    de_set<-paste(substring(fn,1,underscore_loc-1), ifelse(str_detect(fn,"neg"),"neg","pos"))
    print(de_set)
    GSEA_xls<-as.data.frame(read.delim(paste0(home_dir,fn)))%>%
               mutate(ROW = row_number())
    
    sheet_name <-de_set
#browser()
    if (str_length(sheet_name)>31) sheet_name <-str_sub(de_set,9,str_length(de_set)-2)
    addWorksheet(wb, sheet_name)
    writeData(wb,sheet_name , subset(GSEA_xls,FDR.q.val<=fdr), rowNames = TRUE) 
    setColWidths(wb, sheet_name, cols = 1:15, widths = "auto")
    
  }
  saveWorkbook(wb, file=paste0(home_dir,"/GSEA_all_results_FDR.",fdr,"_",proj,"_",set,
                               ifelse(missing(set_name),"",paste0("_",set_name)),".xlsx"), 
               overwrite = TRUE)
  
}  

GSEA_results_file_long <- function(file_type="tsv",set_name,sub_dir_prefix,fdr=0.05){
#browser()
    file_names <-list.files(home_dir, 
                          pattern= glob2rx(paste0("*gsea_report_for_na*.",file_type,"$")),
                          recursive = TRUE)
  
    if (!missing(sub_dir_prefix))       file_names <-file_names[str_detect(file_names, sub_dir_prefix)]

    duplicated(file_names)
    file_names[duplicated(file_names)]
    file_names <-as.data.frame(file_names) #%>%
      #dplyr::filter(str_detect(.,set))
    
    wb <- createWorkbook()
    row_no<-1
    sheet_name <-"all_GSEA"
    addWorksheet(wb, sheet_name)
    posStyle <- createStyle(fontColour = "#006100", fgFill = "#C6EFCE")
    for (fn in file_names$file_names) {
      #fn<- "CS_vs_CTRL_C2.GseaPreranked.1592844211821/gsea_report_for_na_neg_1592844211821.xls"
      print(fn)
      underscore_loc<-as.numeric(as.data.frame(str_locate(fn, ".GseaPreranked"))[1])
      de_set<-paste(substring(fn,1,underscore_loc-1), ifelse(str_detect(fn,"neg"),"neg","pos"))
      print(de_set)
      GSEA_xls<-as.data.frame(read.delim(paste0(home_dir,fn)))%>%
                  mutate(ROW = row_number())
      
      #sheet_name <-de_set
      #browser()
      #if (str_length(sheet_name)>31) sheet_name <-str_sub(de_set,9,str_length(de_set)-2)
      
      writeData(wb,sheet_name , de_set, rowNames = TRUE, startRow = row_no) 
      if (nrow(subset(GSEA_xls,FDR.q.val<=fdr))>0) writeData(wb,sheet_name , subset(GSEA_xls,FDR.q.val<=fdr), rowNames = TRUE, startRow = row_no+1) 
      setColWidths(wb, sheet_name, cols = 1:15, widths = "auto")
      addStyle(wb,sheet_name, style =posStyle, rows = row_no, cols = 1:1, gridExpand = TRUE)
      row_no<-row_no+nrow(subset(GSEA_xls,FDR.q.val<=fdr))+4
    }
    saveWorkbook(wb, file=paste0(home_dir,"/GSEA_all_results_FDR.",fdr,"_",proj,"_",sub_dir_prefix,
                                 ifelse(missing(set_name),"",paste0("_",set_name)),".xlsx"), 
                                 overwrite = TRUE)
    
}  
  
  GSEA_terms_4words <- function(file_type="tsv",set_name,sub_dir_prefix,fdr=0.05){
 #browser()
    GSEA_word_list <- openxlsx::read.xlsx(paste0(home_dir,"../../GSEA_word_list.xlsx"))
    file_names <-list.files(home_dir, 
                              pattern= glob2rx(paste0("*gsea_report_for_na*.",file_type,"$")),
                              recursive = TRUE)
    if (!missing(sub_dir_prefix))       file_names <-file_names[str_detect(file_names, paste0("^",sub_dir_prefix))]
    duplicated(file_names)
    file_names[duplicated(file_names)]
    
    # only WT files
    #file_names<- subset(file_names,str_detect(file_names,"WT"))
    term_data<-data.frame("NAME"=character(),"NES"=numeric(),"FDR q-val"=numeric())
    for (fn in file_names) {
      #fn<-file_names[1] 
      print(fn)
      fn_list<-unlist(str_split(fn,"_"))
      #underscore_df<-as.data.frame(str_locate_all(fn, "_"))
      #underscore_loc<-as.numeric(underscore_df[nrow(underscore_df)-5,1])
      #de_set<-substring(fn,9,underscore_loc-1)
      de_set<-paste0(fn_list[1],"_",fn_list[2],"_",fn_list[3],"_",fn_list[4])
      print(de_set)
      GSEA_xls<-as.data.frame(read.delim(paste0(home_dir,fn))) 
      
      for (word in unique(GSEA_word_list$word)){
        #   term<-"GO_CARTILAGE_DEVELOPMENT"
        word_found<-GSEA_xls %>%
          dplyr::filter(str_detect(NAME,word) & FDR.q.val<=fdr) %>% 
          dplyr::select("NAME","NES","FDR.q.val") %>%
          mutate(NES=as.numeric(NES))
        term_data<- rbind(term_data, 
                          cbind(word_found,
                                data.frame(word = rep(word,nrow(word_found))),
                                data.frame(de_set = rep(de_set,nrow(word_found)))))
      }
    }
    nrow(term_data)
    
    # concatinate words per term 
    term_data_uni <- aggregate(word~ NAME+NES+FDR.q.val+de_set, term_data, 
                               FUN=paste, collapse= " ")
    nrow(term_data_uni)
#browser()
    
    # 1 row per term
    term_data_cast <- data.table::dcast(setDT(term_data_uni),NAME+word~de_set,fun.aggregate= mean,
                                        value.var=c("NES","FDR.q.val"))
    nrow(term_data_cast)
    term_data_cast<- term_data_cast %>% replace(is.na(.), NA)
    print(term_data_cast)
#browser()
    
    save(term_data_uni,term_data_cast, file=paste0(home_dir,"/term_data_4GSEA_FDR.",fdr,"_",set,
                                                   ifelse(missing(set_name),"",paste0("_",set_name)), ".Rdata"))
    
    #term_data_cast$NAME[duplicated(term_data_cast$NAME)]
    
    sheet_name <-"Terms"
    wb <- createWorkbook()
    addWorksheet(wb, sheet_name)
    posStyle <- createStyle(fontColour = "#006100", fgFill = "#C6EFCE")
    to_color<- which( !is.na(term_data_cast$NES_VHOM_vs_CTRL) &
                        !is.na(term_data_cast$NES_VHOS_vs_CTRL) &
                        !is.na(term_data_cast$NES_CS_vs_CTRL))
    writeData(wb,sheet_name , term_data_cast, rowNames = F) 
    addStyle(wb,sheet_name, style =posStyle, rows = to_color+1, 
             cols = 1:ncol(term_data_cast)+1, gridExpand = TRUE)
    setColWidths(wb, sheet_name, cols = 3:9, widths = "auto")
    saveWorkbook(wb, file=paste0(home_dir,"GSEA_suggested_terms_FDR.",fdr,"_",set,
                                 ifelse(missing(set_name),"",paste0("_",set_name)),".xlsx"), 
                 overwrite = TRUE)
    
  }
  
  GSEA_add_categories <- function(de_set_order="all", category_order, per_cat=10, fdr=0.05, add_heatmap=F,
                                  font_size=6, GSEA_category,subs,
                                  w=5200, h=6000,set_name=NULL,comment="",output="jpg",save_data=F){
    #per_cat=30
    #subs<-c("MOVEMENT","MORPHOGENESIS")
    #subs<-c("GASTRO","MUSCLE")
    #subs=c("IMMUNE","IMMUNE-CYTOKINE")
#browser()
    load(paste0(home_dir,"/term_data_4GSEA_FDR.",fdr,"_",set,
                ifelse(missing(set_name),"",paste0("_",set_name)), ".Rdata"),verbose = T)
    print(nrow(term_data_uni))
    #GSEA_category <- read_excel(paste0(home_dir,"GSEA_category2.xlsx"))
    #just for demonstration
    if (missing(GSEA_category)) GSEA_category <- read_excel(paste0(home_dir,"GSEA_terms_category_FDR.",fdr,"_",set,
                                       ifelse(missing(set_name),"",paste0("_",set_name)),".xlsx"))
    GSEA_category <- GSEA_category %>%
      dplyr::select(NAME,category) %>%
      dplyr::filter(category!="NA")
    
    if (!missing(subs)) {
        GSEA_category <- GSEA_category %>%
                            dplyr::filter(str_detect(category,paste(subs,collapse = '|'))) %>%
                            arrange(category)
    }
    if (missing(category_order)) if (missing(subs)) {
      category_order<-sort(unique(GSEA_category$category))
	  } else{
        category_order<-subs
        GSEA_category <- GSEA_category %>%
				                mutate(category=factor(category, levels = category_order)) 
      }
    
    print(table(GSEA_category$category)  )
    print(setdiff(GSEA_category$NAME,term_data_uni$NAME))
    term_data_uni<- merge(term_data_uni, GSEA_category, by="NAME") 
#browser()   
    if (add_heatmap){
      term_data_cast_cat<- merge(term_data_cast, GSEA_category, by="NAME")
      print(table(term_data_uni$category)  )
      print(table(term_data_cast_cat$category)  )
    }
    
    term_data_uni<-term_data_uni %>% 
      mutate(category=factor(category, levels = category_order)) %>%
      mutate(NAME=as.factor(NAME)) %>%
      arrange(category)
    
#browser()    
    
    if (!("all" %in% de_set_order)){
      term_data_uni_4plot <- term_data_uni %>%
                              dplyr::filter(de_set %in% de_set_order) %>%
                              mutate(NAME=str_pad(NAME,max(str_length(NAME))))
    } else {
      de_set_order<-unique(term_data_uni$de_set)
      term_data_uni_4plot <- term_data_uni %>%
                                mutate(NAME=str_pad(NAME,max(str_length(NAME))))# %>%
      assign("term_data_uni_4plot",term_data_uni_4plot,envir = .GlobalEnv)
    }
    print(table(term_data_uni_4plot$category)  )
    
    #add artificial rows to empty de_set
#browser()    
    if (nrow(term_data_uni_4plot)>0){
      for (cat in levels(term_data_uni_4plot$category)){
        term_cat<-subset(term_data_uni_4plot, category==cat)
        empty_de_set<-setdiff(de_set_order,unique(term_cat$de_set))
        if (length(empty_de_set)>0) 
          if  (!("all" %in% empty_de_set))
            for (ds in empty_de_set){
              term_data_uni_4plot<-rbind(term_data_uni_4plot,
                                         cbind(NAME=term_cat$NAME[1],NES=0,FDR.q.val=0,de_set=ds,word="NA",category=cat))%>%
                mutate(NES=as.numeric(NES)) %>%
                mutate(FDR.q.val=as.numeric(FDR.q.val)) %>%
                arrange(category)
            }
      }
    
#browser()       
      if (!missing(subs)) {
        set_name<-paste0(set_name,"_",paste(subs,collapse = '_'))
      }
#browser()  
    if (output=="jpg"){
      pg <- plot_GSEA(term_data_uni_4plot, name_font=60, per_cat=per_cat)
    dev.copy(jpeg,filename=paste0(DE_result_dir,"/GSEA_barplot2_FDR.",fdr,"_",set,"_",set_name,"_",comment, ".jpg"), 
                   width=w, height=h)
      dev.off()
    }else if (output=="pdf"){
      pg <- plot_GSEA(term_data_uni_4plot, name_font=3.5, per_cat=per_cat)
      dev.copy2pdf(device = quartz, file =paste0(DE_result_dir,"/GSEA_barplot2_",set,"_",set_name,"_",comment,".pdf"),
                 width = 6.5, height = 6)
        dev.off()
    } else {
      pg <- plot_GSEA(term_data_uni_4plot, name_font=font_size, per_cat=per_cat)
    }
    if (save_data) save(term_data_uni,term_data_cast_cat, 
                        file=paste0(home_dir,"/term_data_categ_4GSEA_",set,"_",set_name,"_",comment,"FDR.",fdr,".Rdata"))
      #openxlsx::write.xlsx (term_data_cast, paste0(home_dir,"/GSEA_terms_cat_",set,"_all",".xlsx"), row.names = F, col.names = T) 
        
       
       #-------------------------heatmap
  #browser()
    if (add_heatmap){
       term_data_cast_4plot<- term_data_cast_cat %>%
                                dplyr::filter(category %in% GSEA_category$category)  %>%
                                mutate(category=factor(category, levels = category_order)) %>%
                                arrange(category) %>%
                                dplyr::select(NAME,starts_with("NES"))  %>%
                                setNames(str_replace_all(names(.),"NES_", ""))  #%>%
       if (de_set_order!="all") term_data_cast_4plot<-term_data_cast_4plot%>%
                                dplyr::select(NAME,de_set_order)  %>%
                                relocate((de_set_order))  #%>%    #rev
       term_data_cast_4plot<-term_data_cast_4plot%>%
                                column_to_rownames("NAME")  %>%
                                mutate_all(~replace(., is.na(.), 0))
       print(nrow(term_data_cast_4plot))
       if(ncol(term_data_cast_4plot)>1)  {
         term_data_cast_4plot<-as.data.frame(term_data_cast_4plot[rowSums(term_data_cast_4plot != 0) >0, ] )
         print(nrow(term_data_cast_4plot))
         
         GSEA_category<-GSEA_category %>%
                          #rownames_to_column("NAME")  %>%
                          dplyr::filter(NAME %in% rownames(term_data_cast_4plot))  #%>%
                          #column_to_rownames("NAME")  #%>%
         print(nrow(GSEA_category))
         
#browser()
         anno_r<-column_to_rownames(GSEA_category,"NAME")
         gap_rows<-head(as.numeric(cumsum(table(anno_r$category))), -1)
         plot_genes_heatmap(rld=term_data_cast_4plot,
                            anno_df=NA, #gap_c=gap_cols,
                            cols_dend =FALSE,
                            scl="none",
                            anno_row=anno_r,
                            gap_r = gap_rows,
                            cluster=unique(GSEA_category$category),
                            plot_title= paste("GSEA",proj,set,set_name,comment),
                            set_colnames=T,
                            y_name="Pathways",
                            set_name=paste0("GSEA_",proj,set,"_",set_name,"_",comment))
       }
       
    }
    }
  }    
plot_GSEA <- function(path_data,per_cat=10, name_font=60){
        #path_data<-term_data_uni
      #path_data<-term_data_uni_subset
      gg<-1
      my_plots = list()
      h_all<-numeric()
      cat_uni<-levels(path_data$category)
      print(paste("no of categiries:",length(levels(path_data$category))))
      print(table(path_data$category)  )
      for (cat in cat_uni){
        #cat<-cat_uni[1]
        print(cat)
  #browser()      
        path_data_names<-subset(path_data, category==cat) %>%
            group_by(de_set)  %>%
            arrange(desc(abs(NES))) %>%
            head(per_cat)
        path_data_p<-path_data %>%
                        dplyr::filter(category==cat & NAME %in% path_data_names$NAME)
        print(paste("no of NAMEs:",nrow(path_data_p)))
  #browser()
        my_plots[[gg]]<- ggplot(path_data_p,aes(fill=FDR.q.val, y=NES, x=NAME))+
          geom_bar(position="dodge", stat="identity")+
          theme_grey(base_size = name_font) +
          #coord_cartesian(ylim = Ylims) +
          scale_y_continuous(limits=c(ifelse(min(path_data$NES)<0,min(path_data$NES),0), 
                                      ifelse(max(path_data$NES)>0,max(path_data$NES),0))) +
          scale_fill_continuous(limits=c(min(path_data$FDR.q.val), max(path_data$FDR.q.val)))+
          labs(fill = "FDR qvalue")+
          theme(axis.title.y= element_blank(), 
  			  axis.text.y= element_text(size = name_font,  family="sans"),#)+ 
  			  legend.position = "bottom",
  			  plot.title = element_text(size = rel(1.5), hjust = 0.5),
  			  axis.title = element_text(size = rel(1.25)),
  			  panel.grid.major = element_blank(), 
  			  panel.grid.minor = element_blank(),
  			  panel.background = element_blank())+#, 
  			  #axis.line = element_line(colour = "black"))+
          facet_wrap(~de_set, strip.position = "top",drop=FALSE, scales = "fixed",nrow = 1)+
          scale_color_manual(values=c("#5B1031","#455F4A","#7A5E2E","#144756"))+
          coord_flip()
        #last one shall have y axis
        if(cat!= last(cat_uni)){
          my_plots[[gg]]<-my_plots[[gg]]+ theme(axis.ticks.x= element_blank(),
                                                axis.title.x= element_blank(), 
                                                axis.text.x= element_blank())
        } else {
          my_plots[[gg]]<-my_plots[[gg]]+ theme(axis.title.x= element_text(size = name_font),
                                                axis.text.x= element_text(size = name_font))
        }
        # first shall include titles and space for them
        if(cat!= cat_uni[1] ){ #| length(unique(path_data_p$de_set))==1
          my_plots[[gg]]<-my_plots[[gg]]+ theme(strip.background = element_blank() ,
                                                strip.text.x = element_text(size=0))
        } else {
          my_plots[[gg]]<-my_plots[[gg]]+ theme(strip.text.x = element_text(size=name_font))
        }
        
        #add cat as text. gpar Alpha channel for transparency
        my_plots[[gg+1]] <- grobTree(rectGrob(gp=gpar(fill="grey90", alpha=0.5)),
                                     textGrob(cat,gp=gpar(fontsize=name_font+1)))
        
        gg<-gg+2
        h_all<- c(h_all,nrow(path_data_p)/nrow(path_data)*200)
      }
      #calculate plot hight
      print(length(h_all))
      print(h_all)
  #browser()    
      print(c((length(unique(path_data$de_set))*150)+max(str_length(path_data$NAME),na.rm =T),max(str_length(cat_uni))*10))
      print(wrap_plots(my_plots,ncol=2,heights = round(h_all*35), 
                 widths=c((length(unique(path_data$de_set))*150)+max(str_length(path_data$NAME),na.rm =T),max(str_length(cat_uni))*10)) +
        plot_layout(guides = 'collect')& theme(legend.position = 'bottom'))
      #save(path_data, file=paste0(home_dir,"/GSEA_barplot_4Shlomi", ".Rdata"))
      return(my_plots)
    }
    
GSEA_top<- function(top_no=20, meta_dir=DE_result_dir, neg_top=F){

  # if (shape!="set")  {
  # GSEA_topX<- term_data_uni %>%
  #   dplyr::select(-word)  %>%
  #   group_by(de_set)  %>%
  #   arrange(desc(abs(NES)), .by_group = TRUE) %>%
  #   top_n(top_no,wt = abs(NES))
  # }else {
#browser()
  GSEA_topX<- term_data_uni %>%
      dplyr::select(-word)  %>%
      mutate(tissue=str_split_fixed(de_set,"_",2)[,1],
             comp=str_split_fixed(de_set,"_",2)[,2])  %>%
      group_by(comp)  %>%
      arrange(desc(abs(NES))) %>% #   , .by_group = TRUE) %>%
      top_n(top_no,wt = abs(NES))
  # }
#browser()
  table(GSEA_topX$comp)
  openxlsx::write.xlsx(GSEA_topX,
                       paste0(meta_dir,"/GSEA_top",top_no,"_",set,".xlsx"),
                       rowNames = F, colNames = TRUE) 
  GSEA_topX_cat<- GSEA_topX %>%
    left_join(subset(GSEA_category,select=c(NAME,category)),by="NAME") %>%
    mutate(NAME=str_sub(NAME,1,80),
           category=as.factor(category))
  
  mycolors <- distinctColorPalette(length(levels(GSEA_topX_cat$category)))
  names(mycolors) <- sample(levels(GSEA_topX_cat$category))
  # mycolors <- list(category = mycolors)
  
  pdf(file=paste0(DE_result_dir,"/GSEA_top",top_no,"_dotplot_",set,"_",run,"_","_FDR.",GSEA_FDR,".pdf"),
      paper = "USr",width=11, height=8.5) 
  for (ds in unique(GSEA_topX_cat$comp) ){
    print(ds)
    if (neg_top) {
      g<-ggplot(subset(GSEA_topX_cat,comp==ds),aes(x=reorder(NAME,-NES),y=NES,color=category, shape=tissue))
    }else{
      g<-ggplot(subset(GSEA_topX_cat,comp==ds),aes(x=reorder(NAME,NES),y=NES,color=category, shape=tissue))
    }
#browser()
    print(g + 
            geom_point(aes(size=FDR.q.val))+
            scale_size(trans = 'reverse') +
            scale_color_manual(values=mycolors)+
            labs(x="", title=ds)+
            theme(axis.text=element_text(size=8), #axis.title=element_text(size=18,face="bold"),
                  panel.grid.major = element_blank(), panel.grid.minor = element_blank()) +
            coord_flip())
  }
  dev.off()
}  
  

GSEA_genes_4pathway<- function(pw, anno_field, meta_dir=DE_result_dir, anno_all){
  
  #pw<-c("HALLMARK_INTERFERON_ALPHA_RESPONSE")
  # pathway<-pathways[1]
  print(pw)
  pdf(file=paste0(meta_dir,"/GSEA_pathway_","genes_",pw,"_",set,".pdf"))
  
  file_names <-list.files(home_dir, 
                          pattern= paste0(pw,".tsv"),
                          recursive = TRUE)
  if (length(file_names)!=0){
    pw_genes_all_res<-character()
    for (fn in file_names){
      #fn<-file_names[3]
      print (fn)
      pw_genes<- as.data.frame(read.delim(paste0(home_dir,fn))) %>%
                    dplyr::filter(CORE.ENRICHMENT=="Yes") %>%
                    pull(SYMBOL)
      print(paste("excel enriched genes:",length(pw_genes)))
      #convert genes from mouse to human for GSEA
      pw_mouse_gene <- homologene::human2mouse(pw_genes) %>%
                        pull(mouseGene)
      print(paste("length(pw_mouse_gene):",length(pw_mouse_gene)))
      
      pw_genes_all_res<-unique(c(pw_genes_all_res,pw_mouse_gene))
      
      #underscore_loc<-as.numeric(as.data.frame(str_locate_all(fn, "_"))[3,1])
      set<-str_split_fixed(fn,"_",2)[1]
      st<-str_split_fixed(fn,"\\_",3)[,2]
      nd <- str_split_fixed(fn,"\\_",5)[,4]
      de_set<-paste0(st,"_vs_",nd)
      print(de_set)
      
      res_name<-paste0(  "y_",de_set)
      res_pair_df<-get (paste0("gene_", res_name))
      vst_pair_df<-get (paste0("vst_", res_name))
      anno_df<-get (paste0("anno_",res_name))
      
      #        path_genes_rld <- subset( vst_pair_df,
      #                                  str_to_upper(rownames(vst_pair_df)) %in% pw_genes$SYMBOL)
      path_genes_vst <- subset( vst_pair_df, rownames(vst_pair_df) %in% pw_mouse_gene)%>%
                          relocate(rownames(anno_df))
      print(paste("enriched genes found in set:",nrow(path_genes_vst)))
      if (nrow(path_genes_vst)>1){
        #order the cols
        #path_genes_vst_pair<-
        #    path_genes_vst_pair[ ,order(match(colnames(path_genes_vst_pair),rownames(meta))) ]
        #path_genes_vst<-
        #    path_genes_vst[ ,order(match(colnames(path_genes_vst),rownames(meta))) ]
        #plot heatmap only pair
        #plot_genes_heatmap(path_genes_vst_pair,anno_df, #gap_c=gap_cols,
        pv <- plot_volcano(res_pair_df,output_format="",
                           set_name=paste(substr(pw,1,50),"genes on", res_name,set,proj),
                           vtitle=paste(substr(pw,1,50),"genes","\non", res_name,set,proj),
                           lfc= thresh_pair,label_list = rownames(path_genes_vst))
        
        plot_genes_heatmap(path_genes_vst,anno_df, #gap_c=gap_cols,
                           paste0(proj,"_pairs_ ",set,"_",de_set,"_",pw),
                           cols_dend =FALSE,
                           plot_title= paste(proj,set,de_set,"- Pathway\n",pw),
                           out_format="",
                           set_colnames=FALSE)
        #         } else {print("too little sig genes")}
      }else {print("too little genes")}
    }
  } else {print("path empty")}

  if (missing(anno_all)) anno_all<- subset(meta,select=eval(anno_field))
#browser()
  gap_cols <- match(unique(anno_all[[anno_field]]), anno_all[[anno_field]])-1
  gap_cols <-gap_cols[2:length(gap_cols)]
  print(gap_cols)
  print(paste("length(pw_genes_all_res:",length(pw_genes_all_res)))
  pw_all_genes_vst <- subset( vst_all, rownames(vst_all) %in% pw_genes_all_res)%>%
                      relocate(rownames(anno_all))
  
  plot_genes_heatmap(pw_all_genes_vst,anno_all, gap_c=gap_cols,
                     paste0(proj,"_pairs_ ",set,"_",de_set,"_",pw),
                     cols_dend =FALSE,
                     plot_title= paste(proj,set,"all genes- Pathway\n",pw),
                     out_format="",
                     set_colnames=FALSE)
  dev.off()
  
}

GSEA_genes_4category<- function(categ="ENERGY", gene_res_name=gene_res, anno_field,fnts=8,
                                vst_in=vst_all, md_in=meta, set_name,
                                sig_only=F, meta_dir=DE_result_dir){
  
  # categ<-"ENERGY" # METABOLISM LIPID NEURO ENERGY INFLAMMATION
  # sig_only<-F
  GSEA_colname<-"SYMBOL"
  file_type="tsv"
  
  if (!exists("meta")) {
    load(paste0(home_dir,"/",proj,"_",set,"_data", ".Rdata"),verbose=T)
  }
  
  GSEA_category_cat <- read_excel(paste0(home_dir,"GSEA_terms_category_FDR.",GSEA_FDR,"_",set,"_",run,".xlsx")) %>%
                        dplyr::filter(category==categ)
  
  quantile_val<-0.0005
  
  
  pdf(file=paste0(meta_dir,"/GSEA_categ_",if (sig_only) "sig_","genes_",categ,"_",set,".pdf"))
  wb_comp <- openxlsx::createWorkbook()
  all_categ_genes_unique<-character()
  all_categ_genes_sig<-character()
  for (res in gene_res_name){
    #res<-res_part[1]
    print(res)
    sheet_name<-res
    addWorksheet(wb_comp,sheet_name )
    
    res_df<- as.data.frame(get(res))
    vst_pair_df<- get(str_replace(res,"gene","vst"))
    anno_df<- get(str_replace(res,"gene","anno"))
    sig_vst<- as.data.frame(get(paste0(res,"_sig.",thresh_pair)))
    #load(paste0(home_dir,"/vst_",set,"_Batch_Condition_",res,"_sig.",thresh_pair,".Rdata"),verbose = T)
    res_sig_list<-rownames(sig_vst)
    quantile_df<-res_df %>%
      dplyr::filter(log2FoldChange>quantile(log2FoldChange, quantile_val) & 
                      log2FoldChange<quantile(log2FoldChange, (1-quantile_val)))
    nrow(quantile_df)
    if(sig_only){
      sig_vst<-sig_vst%>%
        dplyr::relocate(rownames(anno_df))
      nrow(sig_vst)
      pgh_categ_genes<-pathway_gene_heatmap(rld_mtx=sig_vst,anno_df=anno_df,res_name=res,inc_sig=T,sub_dir_prefix=paste0(set,"_"),
                                            res_sig=res_sig_list,res_df=quantile_df,terms=GSEA_category_cat,
                                            file_type=file_type,categ=categ,output_format = "")
    } else {
      vst_pair_df<-vst_pair_df%>%
        dplyr::relocate(rownames(anno_df))
      pgh_categ_genes<-pathway_gene_heatmap(rld_mtx=vst_pair_df,anno_df=anno_df,res_name=res,
                                            inc_sig=F,sub_dir_prefix=paste0(set,"_"),term_df=GSEA_category_cat,
                                            file_type=file_type,categ=categ, #res_df=quantile_df,
                                            output_format = "") 
    }
    if (nrow(pgh_categ_genes)>0){
      # pgh_categ_genes<-pgh_categ_genes%>%
      #                   dplyr::filter(!is.na(pw_mouse_gene))
      writeData(wb_comp,sheet_name , pgh_categ_genes, colNames = TRUE, startRow = 1) 
      categ_genes_unique<-unique(pgh_categ_genes$pw_mouse_gene)
      categ_genes_unique_df<-as.data.frame(categ_genes_unique)%>%
        purrr::set_names("unique_categ_genes")%>%
        left_join(rownames_to_column(res_df,"gene")%>%
                    dplyr::select(gene,log2FoldChange,pvalue),by=c("unique_categ_genes"="gene")) 
      if (!sig_only){
        categ_genes_unique_df<- categ_genes_unique_df %>%
          mutate(sig=ifelse(unique_categ_genes %in% res_sig_list,"sig","")) %>%
          arrange(desc(sig),unique_categ_genes)
      }
      writeData(wb_comp,sheet_name , categ_genes_unique_df, colNames = TRUE, startCol = 4) 
      all_categ_genes_unique<-unique(c(all_categ_genes_unique,categ_genes_unique_df$unique_categ_genes))
      print(length(all_categ_genes_unique))
      all_categ_genes_sig<-unique(c(all_categ_genes_sig,
                                    as.character(unlist(subset(categ_genes_unique_df,sig=="sig",select=unique_categ_genes)))))
    } #nrow(pgh_categ_genes)>0
  }
  dev.off()
  setColWidths(wb_comp, sheet_name, cols = 1:10, widths = "auto")
  openxlsx::saveWorkbook(wb_comp, 
                         file=paste0(meta_dir,"/Category_",if (sig_only) "sig",
                                     "_genes_",categ,"_",set,".xlsx"),                        
                         overwrite = TRUE)
  
  
  pdf(file=paste0(meta_dir,"/",ifelse(missing(set_name),"ALL",set_name),"_GSEA_categ_",if (sig_only) "sig_","genes_",categ,"_",set,".pdf"))
  categ_genes_vst <- subset( vst_in,rownames(vst_in) %in% all_categ_genes_unique) 
  anno_df<- subset(md_in,select=eval(anno_field))
#browser()
  if (nrow(categ_genes_vst)>0){
    categ_genes_vst<- categ_genes_vst[ ,order(match(colnames(categ_genes_vst),rownames(anno_df))) ]
    plot_genes_heatmap(categ_genes_vst,anno_df, #gap_c=gap_cols,
                       paste(proj,set,categ),
                       plot_title = paste(proj,set,"Total Genes categ",categ),
                       cols_dend =FALSE,
                       font_size = fnts,
                       set_colnames=T,
                       out_format = "")
  }
  #-----only sig
  categ_genes_sig_vst <- subset( vst_in,rownames(vst_in) %in% all_categ_genes_sig) 
  if (nrow(categ_genes_sig_vst)>0){
    categ_genes_sig_vst<- categ_genes_sig_vst[ ,order(match(colnames(categ_genes_sig_vst),rownames(anno_df))) ]
    assign(paste0(categ,"_genes_sig_vst"),categ_genes_sig_vst,envir = .GlobalEnv)
    plot_genes_heatmap(categ_genes_sig_vst,anno_df, #gap_c=gap_cols,
                       paste(proj,set,categ),
                       plot_title = paste(proj,set,"Total sig Genes categ",categ),
                       cols_dend =FALSE,
                       set_colnames=T,
                       out_format = "")
    plot_genes_heatmap(categ_genes_sig_vst,anno_df, #gap_c=gap_cols,
                       paste(proj,set,categ),
                       plot_title = paste(proj,set,"Total sig Genes categ",categ),
                       cols_dend =FALSE,
                       set_colnames=T,
                       out_format = "")
    
  }
  
  dev.off()
  return(categ_genes_vst)
}


pathway_gene_heatmap <- function(rld_mtx,anno_df,res_name,inc_sig=F,res_sig,res_df,term_df,
                                 file_type="tsv",sub_dir_prefix,categ,min_genes=3,output_format="jpg"){
  pair_name<-str_replace(res_name,"res_","")
  pair_name<-str_replace(pair_name,"gene_y_","")
  print(pair_name)
  
  file_names_all <-list.files(home_dir, 
                              pattern= glob2rx(paste0("*.",file_type)), #xls
                              recursive = TRUE)
  #browser()
  if (!missing(sub_dir_prefix))       file_names_all <-file_names_all[startsWith(file_names_all, sub_dir_prefix)]
  
  if (file_type=="tsv") GSEA_colname<-"SYMBOL" else GSEA_colname<-"PROBE"
  #browser()
  #check if pathway is significant
  if (str_detect(pair_name,"all")){
    sig_path<- term_data_cast$NAME
    file_names<-file_names_all[str_detect(file_names_all,paste(sig_path,collapse = "|"))]
  }else {
    sig_path<-str_trim(subset(term_data_uni,str_detect(de_set,pair_name),"NAME")$NAME)
    file_names<-file_names_all[str_detect(file_names_all,paste(sig_path,collapse = "|")) & 
                                 str_detect(file_names_all,pair_name)]
  }
  #browser()
  if (!missing(categ)){
    GSEA_category_cat_set <-term_df %>%
      dplyr::select(NAME,category,starts_with("NES")) %>%
      pivot_longer(cols= starts_with("NES"), names_to ="pair",values_to =  "NES",values_drop_na = TRUE) %>%
      dplyr::filter(str_detect(pair,pair_name)) #%>%
    if (nrow(GSEA_category_cat_set)>0) {
      file_names<-file_names[str_detect(file_names,paste(GSEA_category_cat_set$NAME,collapse = "|"))]
    } else {
      file_names<-character()
    }
    # file_names[str_detect(file_names,"GOBP_ATP_BIOSYNTHETIC_PROCESS")]
  }
  print(paste("no of files:",length(file_names)))
  print(paste("no of sig path:",length(sig_path)))
  #browser()    
  categ_genes<-data.frame(pw=character(),gene=character())
  for (fn in file_names){
    # fn<-file_names[1]
    print(fn)
    pathway<-str_replace(basename(fn),paste0(".",file_type),"")
    pw_xls<-as.data.frame(read.delim(paste0(home_dir,fn)))
    print(paste("excel lines:", nrow(pw_xls)))
    pw_genes<- subset(pw_xls, CORE.ENRICHMENT=="Yes", eval(GSEA_colname)) #PROBE
    print(paste("excel enriched genes:",nrow(pw_genes)))
    
    #browser()
    #convert genes from mouse to human for GSEA
    # pw_mouse_gene <- Human2Mouse(HumanGenes = unlist(c(pw_genes[1])), human ,mouse)
    pw_mouse_gene <- homologene::human2mouse(unlist(c(pw_genes[1])))$mouseGene
    # pw_mouse_gene <- convert_human_to_mouse_symbols(unlist(c(pw_genes[1])))
    print(paste("length(pw_mouse_gene):",length(pw_mouse_gene)))
    #browser()      
    #collect all genes in category
    # categ_genes<- unique(rbind(categ_genes,cbind(pathway,pw_genes)))
    if (length(pw_mouse_gene)>0) categ_genes<- unique(rbind(categ_genes,cbind(pathway,pw_mouse_gene)))
    print(paste("nrow(categ_genes):",nrow(categ_genes)))
    
    pw_genes_rld <- subset( rld_mtx,rownames(rld_mtx) %in% pw_mouse_gene) #PROBE
    print(paste("enriched genes found in set:",nrow(pw_genes_rld)))
    #pw_genes_sig <- subset( res_sig,str_to_upper(rownames(res_sig)) %in% pw_genes[,eval(GSEA_colname)]) #PROBE
    #nrow(pw_genes_sig)
    if (nrow(pw_genes_rld)>min_genes & missing(categ)){
      pw_genes_rld<-
        pw_genes_rld[ ,order(match(colnames(pw_genes_rld),rownames(anno_df))) ]
      #if (inc_sig) {
      #browser()    
      if(!missing(res_df)){
        pv <- plot_volcano(res_df,
                           set_name=paste(substr(pathway,1,50),"genes on", res_name,set,proj),
                           vtitle=paste(substr(pathway,1,50),"genes","\non", res_name,set,proj),
                           lfc= thresh_pair,label_list = rownames(pw_genes_sig))
      }
      
      plot_genes_heatmap(pw_genes_rld,anno_df, #gap_c=gap_cols,
                         paste(proj,set,res_name," SIG- Pathway",substr(pathway,1,70)),
                         plot_title = paste(proj,set,res_name,"SIG- \nPathway",pathway),
                         cols_dend =FALSE,
                         set_colnames=T,
                         out_format = output_format)
      #if (!str_detect(res_name,"all")) {
      
      #tit<-paste(proj,set,res_name,"LFC SIG-Pathway \n",substr(pathway,1,70))
      #print(ggplot(rownames_to_column(pw_genes_rld), aes(fill=pvalue, 
      #                                                         y=log2FoldChange, 
      #                                                         x=reorder(rowname,log2FoldChange)))+
      #        geom_bar(position="dodge", stat="identity")+
      #        #theme_grey(base_size = 20) +
      #        theme(panel.grid.major = element_blank(), 
      #              panel.grid.minor = element_blank(),
      #              panel.background = element_blank(), 
      #              axis.line = element_line(colour = "black"),
      #              text=element_text(size = 18,  family="sans") )+ 
      #        ggtitle(tit)+
      #        theme(axis.title.y= element_blank())+#, 
      #        coord_flip())
      
      #w<-max((max(pw_genes_rld$log2FoldChange)-min(res_sig$log2FoldChange))*3,str_length(tit)*3) 
      #h<-if (nrow(pw_genes_rld)<10) nrow(pw_genes_rld)*20 else nrow(pw_genes_rld)*10
      #if (output_format=="jpg") ggsave(paste0(DE_result_dir,"/barplot_SIG_Pathway_",substr(pathway,1,70),"_",proj,set,res_name,".jpg"), 
      #       width = w, height = h,  dpi=300, units="mm",limitsize = FALSE)
      #}
      #}else{
      #plot_genes_heatmap(pw_genes_rld,anno_df, gap_c=gap_cols,
      #                   paste(proj,set,res_name,"- Pathway",substr(pathway,1,70)),
      #                   plot_title = paste(proj,set,res_name,"- \nPathway",pathway),
      #                   cols_dend =FALSE,
      #                   set_colnames=T,
      #                   out_format = output_format)
      #}
    }else {print("too little genes/categ level")}
  }
  #save(categ_genes,file=paste0(home_dir,"/Top_sig_pathways_genes_4corr_",set,".Rdata"))
  
  #browser()
  pw_genes_rld <- subset( rld_mtx,rownames(rld_mtx) %in% unique(categ_genes$pw_mouse_gene)) #PROBE
  if (nrow(pw_genes_rld)>min_genes){
    pw_genes_rld<-pw_genes_rld[ ,order(match(colnames(pw_genes_rld),rownames(anno_df))) ]
    plot_genes_heatmap(pw_genes_rld,anno_df, #gap_c=gap_cols,
                       paste(proj,set,res_name,if (!missing(categ)) {paste("categ",categ)}),
                       plot_title = paste(proj,set,res_name,"Total",if (!missing(categ)) paste("categ",categ)),
                       cols_dend =FALSE,
                       set_colnames=T,
                       out_format = output_format)
  }
  #browser()
  return(categ_genes)
}


  up_or_down_genes<- function(res_name ,up_or_down, gene_list,thresh, meta_dir=home_dir,out_format="jpeg",
                              organism_name="mouse",
                              rld_mtx,anno_df,wb,res_df,fontsize=8,run,res_dir = DE_result_dir){
    #gene_list<-up
    #res_name<-res
    #up_or_down<-"up"
#browser()
    if(missing(run)) run<-set
    print(length(gene_list))
    gene_list_rld <- rld_mtx%>%
      as.data.frame(stringAsFactor=F)  %>%
      rownames_to_column("gene")  %>%
      dplyr::filter(gene %in% gene_list)  %>%
      mutate(gene=substr(gene,1,60)) %>%
      column_to_rownames("gene")
    gene_list_rld <- gene_list_rld - rowMeans(gene_list_rld)
#browser()    
    # plot_genes_heatmap(gene_list_rld,anno_df,paste0(run,"_",res_name,"_sig_",up_or_down,".",thresh), 
    #                    res_dir = res_dir, font_size = fontsize, out_format=out_format)
    assign(paste0(res_name,"_",up_or_down,"_",thresh),gene_list, envir = .GlobalEnv)
    
    sheet_name <- str_remove(paste0(res_name,".",up_or_down,".",thresh ),"gene_y_")
    print(sheet_name)
    if (str_length(sheet_name)>31) sheet_name<-substr(sheet_name,1,31)
    addWorksheet(wb, sheet_name)
    anno_gene_list <- anno_genes(gene_list,meta_dir = meta_dir,org=organism_name) %>%
      dplyr::rename("gene"="external_gene_name")%>%
      left_join(rownames_to_column(res_df,"gene"),by="gene")
    writeData(wb,sheet_name , anno_gene_list, colNames = TRUE, startRow = 1) 
    setColWidths(wb, sheet_name, cols = 2:10, widths = "auto")
    return(gene_list_rld)
  }
  
  anno_genes<- function(gene_list, org="mouse",meta_dir=home_dir){
   # if (!exists(paste0(meta_dir,"../../mouse.Rdata"))) {
    #if (!exists("mouse_ensembl")) mouse_ensembl = useDataset("mmusculus_gene_ensembl",mart=useMart("ensembl"))
    #save(mouse_ensembl,file=paste0(meta_dir,"../../mouse.Rdata"))
    #save(human_ensembl,file=paste0(meta_dir,"../../human.Rdata"))
    #save(dm_ensembl ,file=paste0(meta_dir,"../../dm.Rdata"))
    if (org== "mouse" ){
      if (!exists("mouse_ensembl"))  {load(paste0(meta_dir,"/../../mouse.Rdata"),verbose = T)}
      ens<-mouse_ensembl
    } else if (org== "human" ){
      if (!exists("human_ensembl"))  {load(paste0(meta_dir,"/../../human.Rdata"),verbose = T)}
      ens<-human_ensembl
    } else if (org== "dm" ){
      if (!exists("dm_ensembl"))  {load(paste0(meta_dir,"/../../dm.Rdata"),verbose = T)}
      ens<-dm_ensembl
    }
    #listAttributes(mart=mouse_ensembl)
#browser()    
    if (org!= "dm" ){
      result = tryCatch({
        anno_data<-
            getBM(attributes=c('external_gene_name','description','gene_biotype'), useCache = FALSE,
                  #filters = 'mgi_symbol', 
                  filters = 'external_gene_name', 
                  values = gene_list, 
                  mart = ens)
      },
      error = function (condition) {
        anno_data<-data.frame(external_gene_name=gene_list)
      })
  
  } else {
      dmdb <- org.Dm.eg.db
      result<-select(dmdb, keys = gene_list, keytype = "UNIPROT",
                   columns = "ONTOLOGY")  %>% #c("SYMBOL", "UNIPROT", "GENENAME","ONTOLOGY")) %>%
        group_by(UNIPROT) %>%
        summarise(ONTOLOGY=paste(ONTOLOGY,collapse=',')) %>%
        dplyr::select(external_gene_name="UNIPROT","ONTOLOGY")
    
  }
    #return(anno_data)
    if (nrow(result)==0) result<-data.frame(external_gene_name=gene_list)
    return(result)
}
  
  

find_cells <- function(sample_data=sample_data,anno_field, pair_stat=F,line_field,set_name="sample_data",clust_cols=F){
#browser()  
  sample_data_human<-sample_data
  Mouse2HumanTable <- Mouse2Human(MouseGenes = rownames(sample_data),human=human,mouse=mouse)
  # sample_data_human$human_gene <- convert_mouse_to_human_symbols(unlist(c(rownames(sample_data))))
  print(paste("length(Mouse2Human):",sum(!is.na(sample_data_human$human_gene ))))
  sample_data_human$human_gene<-Mouse2HumanTable[match(rownames(sample_data_human), Mouse2HumanTable$MGI), "HGNC"]
  #sample_data_human$human_gene<-Mouse2HumanTable[match(sample_data_human$gene, Mouse2HumanTable$MGI), "HGNC"] 
  sample_data_human<- sample_data_human %>%
    dplyr::filter(!is.na(human_gene) & str_length(human_gene)>0) %>%
    #dplyr::select(-gene) %>%
    remove_rownames() %>%
    relocate(human_gene,everything())
  
  
  print(paste("nrow(sample_data_human):",nrow(sample_data_human)))
  sample_data_human_mean <- sample_data_human %>%
    group_by(human_gene) %>%
    summarise_if(is.numeric, mean) %>%
    column_to_rownames("human_gene") %>%
    dplyr::select(rownames(meta))
  print(paste("nrow(sample_data_human_mean):",nrow(sample_data_human_mean)))
  
  
  cells<-xCellAnalysis(sample_data_human_mean) 
  
  
  cells_df <-as.data.frame(cells) %>%
    dplyr::filter(rowSums(.) > 0.001) #%>%
  #browser() 
  anno_df<-subset(meta,select=eval(anno_field))
  print(pheatmap(cells_df, annotation_col=anno_df,cluster_cols=clust_cols,scale = "row"))
  dev.copy(jpeg,filename=paste0(DE_result_dir,"/xCell_",set_name,"_",set,"_",proj, "_heatmap.jpg"), 
           width=600, height=700)
  dev.off()
  
  if (!missing(line_field)) meta_part<-subset(meta,select=c(eval(anno_field),eval(line_field))) else meta_part<-anno_df
  
  cells_long <- cells_df %>%
    rownames_to_column("cell") %>%
    pivot_longer(cols = -cell,names_to = "sample_name", values_to = "rank") %>%
    left_join(rownames_to_column(meta_part,"sample_name"),by= "sample_name" )
  
  #browser()
  pairs<-combn(unique(as.character(cells_long[[anno_field]])), 2)
  pairs_list <- as.list(as.data.frame(pairs))
  pdf(file=paste0(DE_result_dir,"/xCell_",set_name,"_",set,"_",proj, "_boxplot.pdf"),width=7, height=7) 
  for (cl in rownames(cells_df)){
    print(cl)
    g<-ggplot(subset(cells_long,cell==cl), aes(x = get(anno_field), y = rank, fill=get(anno_field))) +
      xlab(eval(anno_field))+
      labs(fill = eval(anno_field))+
      ggtitle(paste("Cells boxplot: ",cl))+ 
      geom_boxplot( alpha = 0.10,lwd=1, colour = "grey") +
      geom_point( size= 2, shape= 21, position=position_jitter(width=0.2)) + #aes(fill= cell),
      facet_wrap(~cell ,ncol = 7,scales = "fixed")+
      theme(panel.grid.major = element_blank(), 
            panel.grid.minor = element_blank(),
            panel.background = element_blank(), 
            axis.text.x = element_text(angle = 30),
            axis.line = element_line(colour = "black"),
            text=element_text(size = 12,  family="sans") )
    
    if (pair_stat) {
      g<-g+stat_compare_means(comparisons = pairs_list,method = "wilcox.test",p.adjust.method = "bonferroni")
    } else {
      g<-g+stat_compare_means(method = "kruskal.test",paired = TRUE,size=5)
    }
    if (!missing(line_field)) g<-g+ geom_line(aes(group=get(line_field)),position=position_jitter(width=0.2)) #,color=get(line_field)
    
    print(g)
  }
  dev.off()
  return(cells_long)
}


my_dimReduction<- function(data, markers = NULL,method = "tsne", tsneSeed = 42){
    data <- as.matrix(data)
    rnames <- row.names(data)
    out_dim <- 2
    if (is.null(markers)) {markers <- colnames(data)}
    marker_filtered_data <- data[, markers]
    print(ncol(marker_filtered_data))
    cat("  Running t-SNE...with seed", tsneSeed)
    set.seed(tsneSeed) # Set a seed if you want reproducible results
    tsne_out <- Rtsne(marker_filtered_data, initial_dims = ncol(marker_filtered_data), 
                      dims = 2, 
                      check_duplicates = FALSE, 
                      pca = TRUE)
    mapped <- tsne_out$Y
    if(!is.null(mapped)){
      if(ncol(mapped) < out_dim){
        out_dim <- ncol(mapped)
        message("Run ",method," for dimensional reduction, out dimension coerced to ",out_dim)
      }
      mapped <- mapped[ ,c(1:out_dim)]
      colnames(mapped) <- paste(method, c(1:out_dim), sep = "_")
      rownames(mapped) <- rnames
    }
    cat("  DONE\n")
    return(mapped)
    
  }
  
  
  cluster_function <- function(df, df_tsne, k, set,clusterColorName="Set1", output_format="jpg"){
    #df <- fcs_markers_immune4cluster_equal
    #df_tsne <- fcs_tsne_immune
    #k<-selected_k
    #set<-paste0("immune_",threshold,"_",cells_number)
    
    set.seed(123)
    cluster_FlowSOM <- my_cluster(xdata = df, ydata= df_tsne, method = "FlowSOM", FlowSOM_k = k)
    
    data_FlowSOM <- as.data.frame(cbind(as.matrix(df), df_tsne, FlowSOM=cluster_FlowSOM))
    print(paste("clusterColorName:",clusterColorName))
    #pdf(file=paste0(result_dir,"/FlowSOM_cluster_marker_",set,"_", k,"_",day,"_",clusterColorName, ".pdf"),width=7, height=7)  
    #  print(my_clusterPlot(data=data_FlowSOM, xlab="tsne_1", ylab="tsne_2", cluster="FlowSOM",
    #                       sampleLabel = FALSE, addLabel = TRUE, 
    #                       title = paste("FlowSOM marker",set, "k=",k)))
    print(my_clusterPlot(data=data_FlowSOM, xlab="tsne_1", ylab="tsne_2", cluster="FlowSOM",
                           sampleLabel = FALSE, #addLabel = FALSE, 
                           title = paste("FlowSOM marker",set, "k=",k),
                           clusterColorName=clusterColorName))
    
    if (output_format=="jpg") {
      dev.copy(jpeg,filename=paste0(DE_result_dir,"/FlowSOM_cluster_marker_",set,"_", k,"_", ".jpg"), 
             width=1000, height=700)
      dev.off()
    } 

    return(data_FlowSOM)
  }
  
  my_cluster<- function(xdata = NULL, ydata= NULL, method= c("Rphenograph", "FlowSOM"),
                        FlowSOM_k=30, Rphenograph_k=30, flowSeed = 123){
 
    xdata <- as.matrix(xdata)
    print(method)
    method = match.arg(method)
    if(method == "NULL"){
      return(NULL)
    }
    switch(method, 
           Rphenograph = {
             cat("  Running PhenoGraph...")
             clusters <- as.numeric(membership(Rphenograph(xdata, k=Rphenograph_k)))
           },
           FlowSOM = {
             cat("  Running FlowSOM...")
             set.seed(flowSeed)
             clusters <- FlowSOM_integrate2cytofkit(xdata, FlowSOM_k, flowSeed = flowSeed)
           })
    
    if( length(clusters) != ifelse(is.null(ydata), nrow(xdata), nrow(ydata)) ){
      message("Cluster is not complete, cluster failed, try other cluster method(s)!")
      return(NULL)
    }else{
      if(!is.null(xdata) && !is.null(row.names(xdata))){
        names(clusters) <- row.names(xdata)
      }else if(!is.null(ydata) && !is.null(row.names(ydata))){
        names(clusters) <- row.names(ydata)
      }
      cat(" DONE!\n")
      return(clusters)
    }
    print(length(cluster))
    return(cluster)
    #}
  }
  
  FlowSOM_integrate2cytofkit <- function(xdata, k, flowSeed = NULL, ...){
    cat("    Building SOM...\n")
    xdata <- as.matrix(xdata)
    
    ord <- tryCatch({
      map <- SOM(xdata, silent = TRUE,  ...) #
      cat("    Meta clustering to", k, "clusters...\n")
      metaClusters <- suppressMessages(metaClustering_consensus(map$codes, k = k, seed = flowSeed))
      cluster <- metaClusters[map$mapping[,1]]
    }, error=function(cond) {
      message("Run FlowSOM failed")
      message("Here's the error message:")
      message(cond)
      return(NULL)
    }) 
    
    if(is.null(ord)){
      cluster <- NULL
    }else{
      if(length(ord) != nrow(xdata)){
        message("Run FlowSOM failed!")
        return(NULL)
      }
      cluster <- ord
    }
    
    return(cluster)
  }
  
  my_clusterPlot <- function(data, xlab, ylab, cluster, sample, title = "cluster", 
                             type = 1, point_size = NULL, addLabel=TRUE, 
                             labelSize=8, sampleLabel=TRUE, 
                             labelRepel = FALSE, fixCoord=TRUE, clusterColorName="Set1") {
    if(!is.data.frame(data))
      data <- as.data.frame(data)
    
    if(missing(sample)){
      sample <- "sample"
      data$sample <- "data"
    }
    
    paraCheck <- c(xlab, ylab, cluster, sample) %in% colnames(data)
    if(any(!paraCheck)){
      stop("Undefined parameters found: ",
           c(xlab, ylab, cluster, sample)[!paraCheck])
    }
    
    data[[cluster]] <- as.factor(data[[cluster]])
    data[[sample]] <- as.factor(data[[sample]])
    cluster_num <- length(unique(data[[cluster]]))
    sample_num <- length(unique(data[[sample]]))
    col_legend_row <- ceiling(cluster_num/8)
    size_legend_row <- ceiling(sample_num/4)
    grid_row_num <- round(sqrt(sample_num))
    if (sample_num >= 8) {
      shape_value <- LETTERS[1:sample_num]
    } else {
      shape_value <- c(1:sample_num) + 15
    }
    if (is.null(point_size)) {
      point_size <- ifelse(nrow(data) > 10000, 0.85, 3)
    }
    if (clusterColorName !="hila"){
      getPalette <- colorRampPalette(brewer.pal(9, clusterColorName))
      clusterColor <- getPalette(max(cluster_num))
      
    } else {
      clusterColor <- c("#4a8ff7","#f56733","#fca030", "#f7e045", "#bed160", "#808080", "#8bb559","#72a37f", "#549e88", "#60b1d1", "#325363", "#035496",  "#325cbf", "#6e5399", "#775c82", "#bf91b7", "#e8b7d1", "#bababa", "#e590f0", "#8c6f4f", "#999267", "#64566e")       }

    if(type == 1){
      if(sampleLabel){
        cp <- ggplot(data, aes_string(x = xlab, y = ylab, colour = cluster, shape = sample)) + 
          geom_point(size = point_size) + scale_shape_manual(values = shape_value) + 
          scale_colour_manual(values = clusterColor) + 
          xlab(xlab) + ylab(ylab) + ggtitle(paste(title, "Scatter Plot", sep = " ")) + 
          theme_bw() + theme(legend.position = "bottom") + 
          theme(axis.text=element_text(size=14), axis.title=element_text(size=18,face="bold"),
                panel.grid.major = element_blank(), panel.grid.minor = element_blank()) +
          guides(colour = guide_legend(nrow = col_legend_row, override.aes = list(size = 4)), 
                 shape = guide_legend(nrow = size_legend_row, override.aes = list(size = 4)))
      }else{
        cp <- ggplot(data, aes_string(x = xlab, y = ylab, colour = cluster)) + 
          geom_point(size = point_size) + scale_shape_manual(values = shape_value) + 
          scale_colour_manual(values = clusterColor) + 
          xlab(xlab) + ylab(ylab) + ggtitle(paste(title, "Scatter Plot", sep = " ")) +
          theme_classic()+ 
          #theme_bw() + 
          theme(legend.position = "bottom") + 
          theme(axis.text=element_text(size=14), axis.title=element_text(size=18,face="bold"),
                panel.grid.major = element_blank(), panel.grid.minor = element_blank()) + #,
          #      axis.line = element_line(colour = "black")) +
          guides(colour = guide_legend(nrow = col_legend_row, override.aes = list(size = 4))) #
      }
      
      if(addLabel){
        edata <- data[ ,c(xlab, ylab, cluster)]
        colnames(edata) <- c('x', "y", "z")
        center <- aggregate(cbind(x,y) ~ z, data = edata, median)
        
        if(labelRepel && !sampleLabel){
          cp <- cp + geom_text_repel(data=center, aes_string(x = "x", y = "y", label = "z"),
                                     size = labelSize, fontface = 'bold', color = "black",
                                     box.padding = unit(0.5, 'lines'),
                                     point.padding = unit(1.6, 'lines'),
                                     segment.color = '#555555',
                                     segment.size = 0.5,
                                     arrow = arrow(length = unit(0.02, 'npc')))
        }else{
          cp <- cp + annotate("text", label = center[,1], x=center[,2], y = center[,3],
                              size = labelSize, colour = "black")
        }
      }
      
    }else if (type == 2){
      cp <- ggplot(data, aes_string(x = xlab, y = ylab, colour = cluster)) +
        facet_wrap(~sample, nrow = grid_row_num, scales = "fixed") + 
        geom_point(size = point_size - 0.05 * sample_num) + 
        scale_colour_manual(values = clusterColor) + 
        xlab(xlab) + ylab(ylab) + ggtitle(paste(title, "Grid Plot", sep = " ")) + 
        theme_bw() + theme(legend.position = "bottom") + 
        theme(axis.text=element_text(size=14), axis.title=element_text(size=18,face="bold")) +
        guides(colour = guide_legend(nrow = col_legend_row, override.aes = list(size = 4)), 
               shape = guide_legend(nrow = size_legend_row, override.aes = list(size = 4)))
      
      if(addLabel){
        edata <- data[ ,c(xlab, ylab, cluster)]
        colnames(edata) <- c('x', "y", "z")
        center <- aggregate(cbind(x,y) ~ z, data = edata, median) 
        
        if(labelRepel && !sampleLabel){
          cp <- cp + geom_text_repel(data=center, aes_string(x = "x", y = "y", label = "z"),
                                     size = labelSize, fontface = 'bold', color = "black",
                                     box.padding = unit(0.5, 'lines'),
                                     point.padding = unit(1.6, 'lines'),
                                     segment.color = '#555555',
                                     segment.size = 0.5,
                                     arrow = arrow(length = unit(0.02, 'npc')))
        }else{
          cp <- cp + geom_text(data=center, aes_string(x = "x", y = "y", label = "z"), 
                               size = labelSize, colour = "black") 
        }
      }
      
    }else{ 
      stop("Undefined type, only 1 or 2.") 
    }
    
    if(fixCoord){
      cp <- cp + coord_fixed()
    }
    
    return(cp)
    
  }
  
  Custom_palette <- function(
    low = "white",
    high = "red",
    mid = NULL,
    k = 50
  ) {
    low <- col2rgb(col = low) / 255
    high <- col2rgb(col = high) / 255
    if (is.null(x = mid)) {
      r <- seq(from = low[1], to = high[1], len = k)
      g <- seq(from = low[2], to = high[2], len = k)
      b <- seq(from = low[3], to = high[3], len = k)
    } else {
      k2 <- round(x = k / 2)
      mid <- col2rgb(col = mid) / 255
      r <- c(
        seq(from = low[1], to = mid[1], len = k2),
        seq(from = mid[1], to = high[1], len = k2)
      )
      g <- c(
        seq(from = low[2], to = mid[2], len = k2),
        seq(from = mid[2], to = high[2],len = k2)
      )
      b <- c(
        seq(from = low[3], to = mid[3], len = k2),
        seq(from = mid[3], to = high[3], len = k2)
      )
    }
    return(rgb(red = r, green = g, blue = b))
  }
  
  appendRData <- function(robj, filename) {
    
    tmpEnv <- new.env()
    
    if (!file.exists(filename)) warning("'file' does not exist. A new file will be created.")
    savedObjects <- load(file = filename, envir = tmpEnv, verbose = TRUE)
    
    # quick check for name collisions
    stopifnot(!(deparse(substitute(robj)) %in% savedObjects))
    
    save(list = c(savedObjects, deparse(substitute(robj))),
         file = filename,
         envir = tmpEnv)
  }
```
